# Targeting dormant ovarian cancer cells *in vitro* and in an *in vivo* model of platinum resistance

**DOI:** 10.1101/716464

**Authors:** Zhiqing Huang, Eiji Kondoh, Zachary Visco, Tsukasa Baba, Noriomi Matsumura, Emma Dolan, Regina S. Whitaker, Ikuo Konishi, Shingo Fujii, Andrew Berchuck, Susan K. Murphy

## Abstract

**Objective:** Ovarian cancer cells often exist *in vivo* as multicellular spheroids. Spheroid formation *in vitro* has been used to enrich for cancer stem cell populations from primary tumors. Such spheroids exhibit drug resistance and slow proliferation, suggesting involvement in disease recurrence. Our objectives were to characterize cancer spheroid phenotypes, determine gene expression profiles associated with spheroid forming capacity and to evaluate the responsiveness of spheroids to commonly used and novel therapeutic agents.

**Methods:** Tumorigenic potential was assessed using anchorage independent growth assays in 24 cell lines. Spheroids from cell lines (N=12) and from primary cancers (N=8) were grown on non-adherent tissue culture plates in serum-free media. Cell proliferation was measured using MTT assays and Ki67 immunostaining. Affymetrix HT U133A gene expression data was used to identify differentially expressed genes based on spheroid forming capacity. Matched monolayers and spheroids (N=7 pairs) were tested for response to cisplatin, paclitaxel and 7-hydroxystaurosporine (UCN-01) while mitochondrial inhibition was performed using oligomycin. Xenograft tumors from intraperitoneal injection of CAOV2-GFP/LUC ovarian cancer cells into nude mice were treated with carboplatin to reduce tumor burden followed by secondary treatment with carboplatin, UCN-01, or Oltipraz. Tumor formation and response was monitored using live imaging.

**Results:** Of 12 cell lines with increased anchorage-independent growth, 8 also formed spheroids under serum-free spheroid culture conditions. Spheroids showed reduced proliferation (p<0.0001) and Ki67 immunostaining (8% versus 87%) relative to monolayer cells. Spheroid forming capacity was associated with increased mitochondrial pathway activity (p ≤ 0.001). The mitochondrial inhibitors, UCN-01 and Oligomycin, demonstrated effectiveness against spheroids, while spheroids were refractory to cisplatin and paclitaxel. By live *in vivo* imaging, ovarian cancer xenograft tumors were reduced after primary treatment with carboplatin. Continued treatment with carboplatin was accompanied by an increase in tumor signal while there was little or no increase in tumor signal observed with subsequent treatment with UCN-01 or Oltipraz.

**Conclusions:** Our findings suggest that the mitochondrial pathway in spheroids may be an important therapeutic target in preventing disease recurrence.

## INTRODUCTION

Ovarian cancer is the fifth leading cause of cancer death among women in the United States with the highest mortality rate of all gynecologic cancers ^1^. Although 70% of ovarian cancer patients achieve complete clinical remission following cytoreductive surgery and treatment with a platinum/taxane-based chemotherapeutic regimen, the majority suffer relapse with disease that has acquired or will develop chemoresistance ^2–5^. New therapies aimed at reducing disease recurrence are urgently needed in order to improve survival.

Cancer stem-like cells (CSLCs) that are able to enter into a state of dormancy and later regenerate tumors may provide a causal explanation for ovarian cancer relapse after initially successful treatments ^4, 6^. Furthermore, cells with CSLC characteristics are naturally resistant to chemotherapeutic agents, in part due to expression of drug transporters and a slow rate of proliferation ^7^. In patients with advanced ovarian cancer, ascites fluid often accumulates in the peritoneal cavity and can become enriched with tumor cells that aggregate together to form multilayered spheroids ^8^. These spheroids contain CSLCs ^9^ and exhibit resistance to chemotherapy ^4, 10^. They are also capable of embedding into other tissues and this feature may be responsible for the “grains of sand” texture in the peritoneal cavity in women with advanced stage serous ovarian cancer at the time of initial cytoreductive surgery ^11^. The embedded cells may enter into a state of dormancy, thus rendering them resistant to commonly use chemotherapeutics that target actively proliferating cells. These dormant cells may later awaken to initiate growth and recurrent disease once conditions are favorable. Thus, targeting these spheroids may offer a plausible approach to achieve long-term remission of women with advanced ovarian cancer. To this end, we grew ovarian cancer cells on non-adherent plates using serum-free spheroid culture conditions, which results in aggregation of cells with spheroid-forming capacity into spheroids ^12–14^. We then sought to identify novel candidate drugs that are effective at targeting these spheroids.

In the present study, spheroid-forming ovarian cancer cells indeed exhibited characteristics of cellular dormancy, a condition of greatly reduced growth and/or quiescence that is associated with drug resistance and disease recurrence ^4, 15^. We previously reported that the protein kinase C inhibitor, UCN-01 (7-hydroxystaurosporine), had higher efficacy against slow proliferating and/or quiescent ovarian cancer cells ^16^. Here, we assessed the efficacy of UCN-01 against ovarian cancer spheroids and compared these results to the efficacy of cisplatin and paclitaxel. In addition, analysis of gene expression profiles in spheroid-forming ovarian cancer cell lines versus cell lines that do not form spheroids suggested that the activity of mitochondrial pathway genes contributes to spheroid forming capacity. Mitochondria are known to play a central role in regulating cellular metabolism and produce adenosine triphosphate (ATP) through the citric acid cycle ^17^. Oligomycin, an inhibitor of mitochondrial ATP synthase ^18^, repressed spheroid-formation in a dose-dependent manner. In a mouse xenograft model we found that UCN-01 or mitochondrial inhibitor Oltipraz prolonged the disease-free interval following carboplatin treatment. This markedly contrasted with results from continued carboplatin treatment, in which the mice showed a rapid increase in tumor volume. Collectively, these results show that cells cultured under serum-free spheroid culture conditions are induced into cellular dormancy, and further suggest that the mitochondrial pathway is a promising target for eradication of these dormant ovarian cancer cells.

## RESULTS

Twenty two cancer cell lines were used to perform anchorage-independent growth assays, the first bioassay developed for CSLCs ^19^. Average numbers of colonies formed per microscopic field that were >100 µm in diameter for these cell lines ranged from 0 to 7.15 (Table 1). The cell lines CAOV3, DOV13, and OVCA429 did not form any colonies. OVCA420 only formed colonies <100 µm in diameter and TYK-nu, SKOV8, 41M, OVCA432, OVCAR3, and FUOV1 formed a small number of colonies (>0 but ≤0.1 colonies/microscopic field). Nine cell lines with ≥2.9 colonies/microscopic field were categorized as having high tumorigenic potential, including PEO1, OVCAR8, OVARY1847, OV90, HEYA8, OVCAR2, CAOV2, HEY, and HEYC2.

**Table 1.**
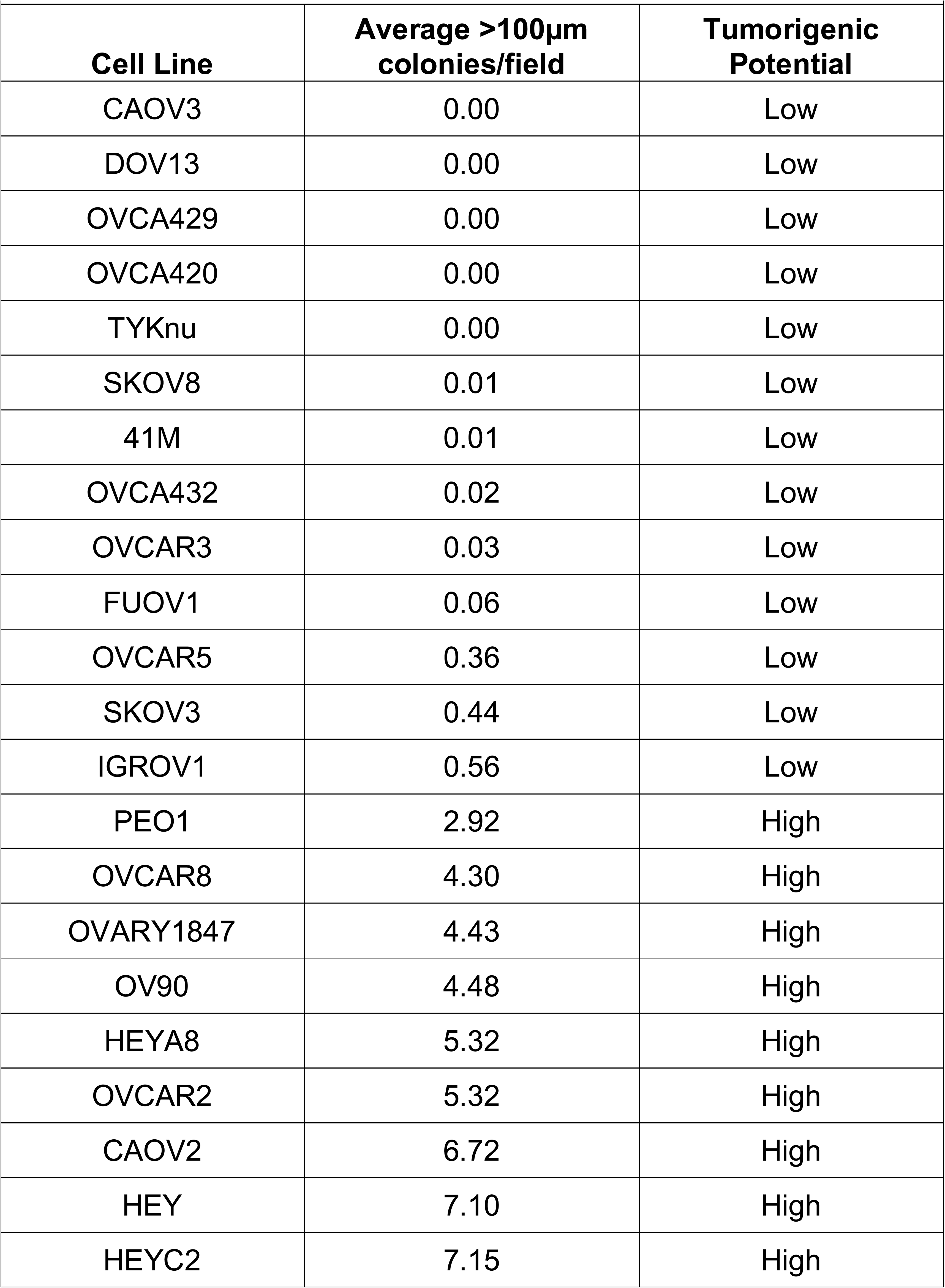
Anchorage-independent growth

| **Table 1. Anchorage-independent growth** |  |  |
| --- | --- | --- |
| **Cell Line** | **Average >100µm colonies/field** | **Tumorigenic Potential** |
| CAOV3 | 0.00 | Low |
| DOV13 | 0.00 | Low |
| OVCA429 | 0.00 | Low |
| OVCA420 | 0.00 | Low |
| TYKnu | 0.00 | Low |
| SKOV8 | 0.01 | Low |
| 41M | 0.01 | Low |
| OVCA432 | 0.02 | Low |
| OVCAR3 | 0.03 | Low |
| FUOV1 | 0.06 | Low |
| OVCAR5 | 0.36 | Low |
| SKOV3 | 0.44 | Low |
| IGROV1 | 0.56 | Low |
| PEO1 | 2.92 | High |
| OVCAR8 | 4.30 | High |
| OVARY1847 | 4.43 | High |
| OV90 | 4.48 | High |
| HEYA8 | 5.32 | High |
| OVCAR2 | 5.32 | High |
| CAOV2 | 6.72 | High |
| HEY | 7.10 | High |
| HEYC2 | 7.15 | High |

Non-adherent spheroid culture conditions have been used to enrich for CSLCs from tumor tissue of a variety of malignancies ^12–14^. Therefore, we tested the ability of the nine cell lines with increased tumorigenic potential to form spheroids using a non-adherent spheroid assay. The numbers of spheroids formed by each of the cell lines and the ability of the spheroids to undergo serial passaging are shown in **Table S1.** Five cell lines were able to undergo passage more than five times and were determined to have increased capacity to form spheroids under serum-free spheroid culture conditions. On the other hand, OVARY1847, OV90, PEO1 and OVCAR8 formed either no or few spheroids, and most were not able to survive beyond two passages (data not shown). Representative monolayer cells and spheroids of HEYA8 are presented in Figure 1A. Primary ovarian cancers were also used for non-adherent spheroid assays. A number of spheroids were obtained initially from seven of eight tumors, but only few spheroids, if any, were formed following five passages. Representative spheroids for primary ovarian cancers are also shown in Figure 1A.

**Figure 1.**
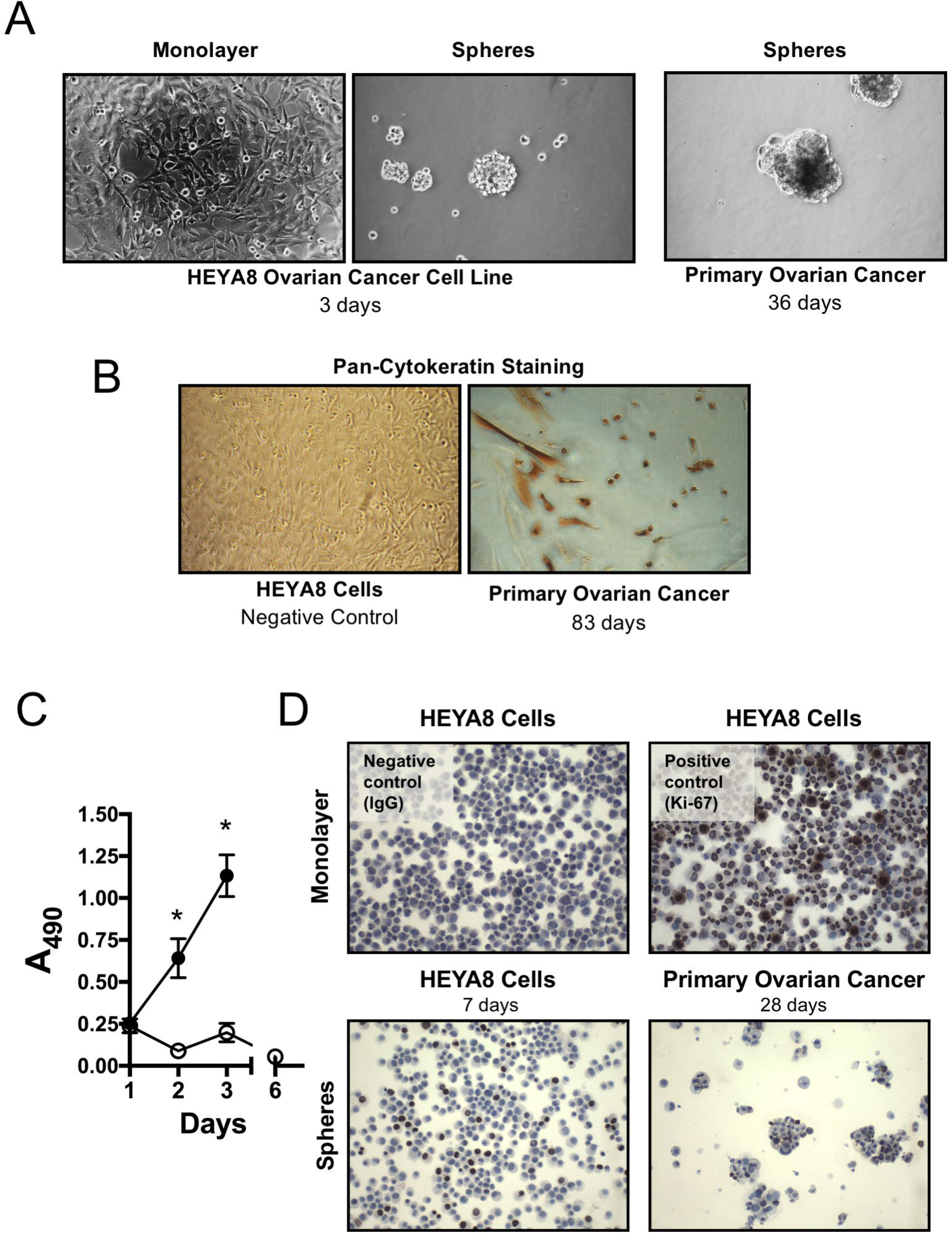
Ovarian cancer cell monolayer cells and corresponding spheroids. (A) Morphology (100X objective) of monolayer cells and spheroids. Left panel shows HEYA8 cells growing as a monolayer under regular culture conditions and as spheroids under serum-free spheroid culture conditions, both three days after seeding. Spheroids from a primary ovarian cancer are also shown on the 36^th^ day, just before the 5th passage in serum-free spheroid culture. (B) Pan-cytokeratin staining of surviving cells from a primary ovarian cancer (83rd day after initiation of serum-free spheroid culture) that adhered to tissue culture-treated dishes (right panel, 100X objective). Left panel shows IgG negative control using HEYA8 cells (100X objective (C) Suppression of HEYA8 cell proliferation in serum-free spheroid culture conditions. Y-axis corresponds to the absorbance at 490 nm (A490). Filled circles, monolayer culture, open circles, spheroid culture. Mean +/- SEM for five replicates shown *, *P*<0.0001 (D) Ki-67 staining of monolayer cells (upper panel, x10) and spheroids (lower panel, 100X objective). In serum-free spheroid culture, a small number of HEYA8 cells (7^th^ day) and a primary ovarian cancer (28^th^ day; just before 4^th^ passage) were Ki-67 positive. Negative (upper left, 100X objective) and positive (upper right, 100X objective) controls for Ki-67 staining used HEYA8 monolayer cells.

Sustainable spheroid formation beyond ten passages was not achieved for the ovarian cancer cell lines tested nor for spheroids derived from primary ovarian cancers. However, a small number of single cells remained viable in serum-free spheroid culture even three months after initiation of the culture. Surviving primary ovarian cancer cells in serum-free spheroid culture conditions for 83, 90, and 98 days were tentatively transferred into tissue culture-treated dishes. They subsequently adhered to the dishes and resumed growth. Representative staining with a pan-cytokeratin marker is shown in Figure 1B. In order to investigate the ability of the cells within the spheroids to self-renew, a characteristic of stem cells, single cell cloning was performed by serial dilution of spheroid-forming cells under serum-free spheroid culture conditions. No secondary spheroids were formed within the three-month duration of the experiment (data not shown).

Serial cell proliferation assays demonstrated that HEYA8 ovarian cancer monolayer cells exhibit rapid growth and became confluent three days after plating, while the growth of HEYA8 cells in serum-free spheroid culture was strongly repressed (unpaired t-test, *P*<0.0001 on days 2 and 3; Figure 1C). The proportion of proliferation marker Ki-67 positive cells was decreased in the spheroid-forming cells of HEYA8 (7^th^ day in serum-free spheroid culture), as compared to HEYA8 grown as monolayer cells (8% versus 87%; Figure 1D). Spheroids from a primary tumor (28^th^ day; just before 4^th^ passage) also exhibited a relatively small population of Ki-67 positive cells (Figure 1D).

The kinase C inhibitor, 7-hydroxystaurosporine (UCN-01; 25nM, 50nM, and 100nM) exhibited an enhanced antiproliferative effect against HEYA8 cells in serum-free spheroid culture compared with HEYA8 monolayer cells at each tested dose (unpaired t-test, *P*=0.0016, *P*=0.016, and *P*=0.009, respectively; Figure 2). The response to cisplatin (25µM, 50µM, and 100µM) did not quite reach statistical significance when comparing the spheroids versus monolayer cells but showed a less antiproliferative effect against spheroids (unpaired t-test, *P*=0.47, *P*=0.08, *P*=0.15, respectively; Figure 2). Paclitaxel treatment (10nM, 50nM, 100nM) did not inhibit, but rather increased cell survival in spheroids (unpaired t-test, *P*=0.003, *P*=0.0002, and *P*=0.001, respectively; Figure 2). Similar results were obtained for the six additional cell lines tested (**Figure S1**).

**Figure 2.**
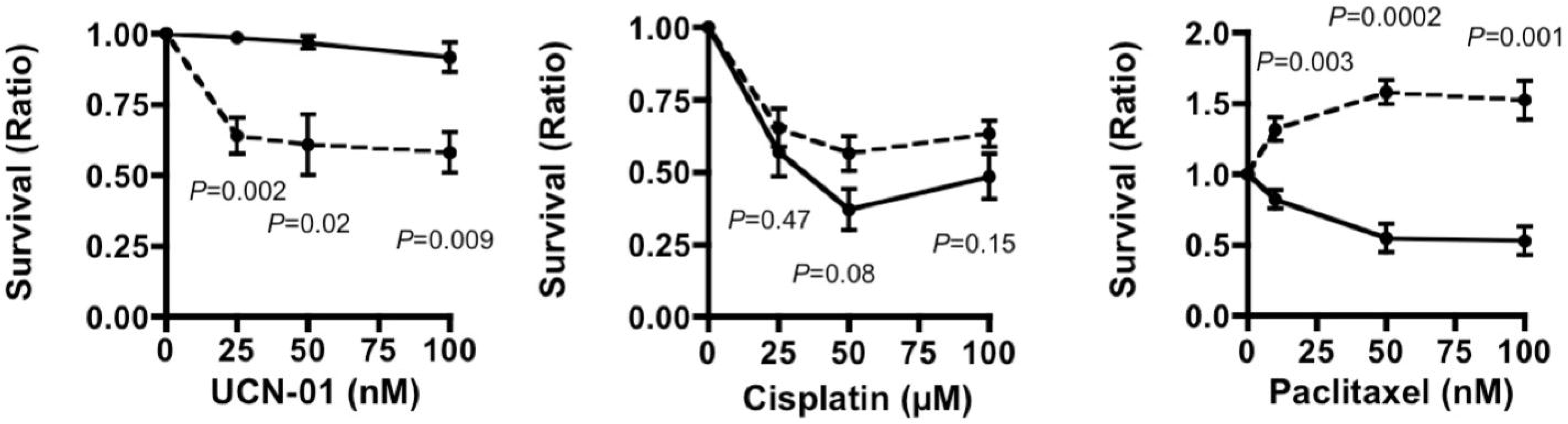
HEYA8 cell monolayers and spheroids exhibit divergent drug sensitivity. UCN-01 (left) exhibits higher efficacy against spheroids relative to monolayer cells, which contrasts with cisplatin (middle) and paclitaxel (right), which are more effective against monolayer cells. Solid and dashed lines correspond to monolayer cells and spheroids, respectively. Shown are the mean +/- SEM.

Gene expression profiling was performed to identify genes and/or pathways that distinguish spheroid-forming cells (N=5) from the cell lines that do not have spheroid-forming capacity (N=4). Of 22,277 probes on the Affymetrix U133 HTA gene expression microarray platform, 1,358 probes (1,066 named genes) were significantly increased in cell lines that have high spheroid forming capacity under serum-free spheroid culture conditions (**Table S2**), while 1,441 probes, representing 1,114 named genes, were significantly increased in cell lines with low potential to form spheroids (**Table S3**). A heat map of all 2,799 probes with significant differences in expression between the spheroid forming and non-spheroid forming groups is shown in Figure 3. Functional annotation clustering analysis of the 1,066 genes whose expression is increased in spheroid forming cells showed the most significant enrichment for genes involved in mitochondrial function (137 genes, 11.6-fold enrichment; FDR=2.3e^-19^) and DNA damage/repair (46 genes, 5.9-fold enrichment; FDR=1.2e^-5^) (Table 2; full list in **Table S4**). For the 1,114 genes that are more highly expressed in the cells with low spheroid-forming capacity, metal binding (5.6-fold enrichment; 268 genes, p=1.8e^-10^, FDR=2.6e^-7^) and cell junctions (4.2-fold enrichment; 67 genes, p=4.3e^-7^, FDR=6.1e^-4^) were the most significant annotations (Table 2; full list in **Table S5**).

**Figure 3.**
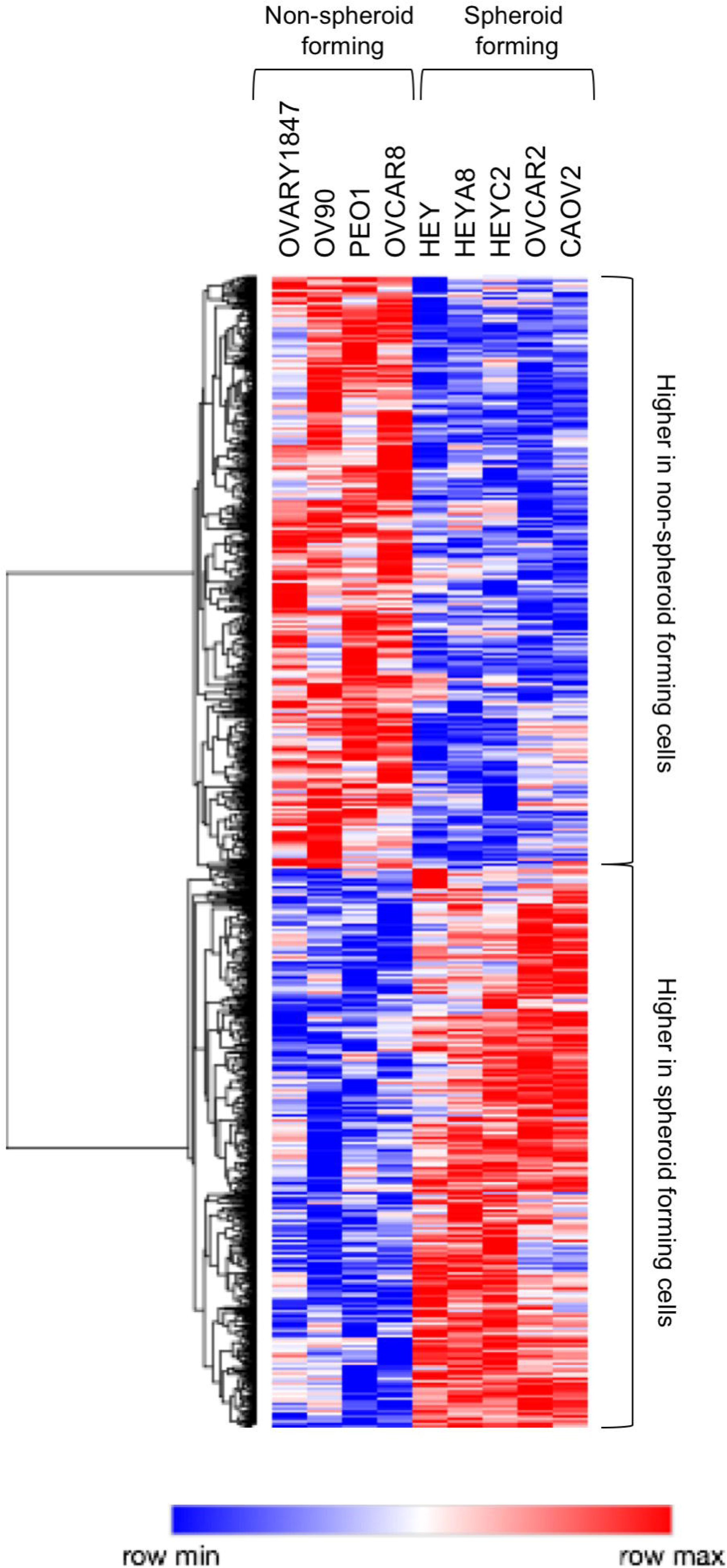
Differentially expressed genes between cells having high and low spheroid-forming ability in serum-free spheroid culture. Unsupervised clustering using 2,799 probes that exhibit differential gene expression between groups. In the heat map, each row represents a gene, and each column corresponds to a cell line. Red, high relative expression; blue, low relative expression.

**Table 2.** Functional annotations for genes discriminating high and low spheroid-forming capacity

|  |  |  |  |
| --- | --- | --- | --- |
| Enriched in cells with high sphere potential |  |  |  |
| Category | Enrichment Score | Term | FDR |
| **Cluster 1** | **11.6** |  |  |
| UP_KEYWORDS |  | Mitochondrion | 2.3e-19 |
| UP_KEYWORDS |  | Transit peptide | 7.5e-09 |
| UP_SEQ_FEATURE |  | Transit peptide:Mitochondrion | 8.4e-07 |
| **Cluster 2** | **5.9** |  |  |
| UP_KEYWORDS |  | DNA damage | 1.2e-05 |
| UP_KEYWORDS |  | DNA repair | 0.003 |
| **Cluster 3** | **5.6** |  |  |
| GOTERM_BP_DIRECT |  | GO:0006412~translation | 3.8e-07 |
| GOTERM_CC_DIRECT |  | GO:0005840~ribosome | 2.8e-06 |
| UP_KEYWORDS |  | Ribosomal protein | 4.7e-06 |
| UP_KEYWORDS |  | Ribonucleoprotein | 5.47E-06 |
| GOTERM_BP_DIRECT |  | GO:0006364~rRNA processing | 1.49E-05 |
| GOTERM_BP_DIRECT |  | GO:0006413~translational initiation | 3.66E-05 |
| GOTERM_MF_DIRECT |  | GO:0003735~structural constituent of ribosome | 7.97E-05 |
| KEGG_PATHWAY |  | hsa03010:Ribosome | 0.006 |
| GOTERM_BP_DIRECT |  | GO:0000184~nuclear-transcribed mRNA catabolic process, nonsense-mediated decay | 0.009 |
| GOTERM_CC_DIRECT |  | GO:0022625~cytosolic large ribosomal subunit | 0.02 |
| Enriched in cells with low sphere potential |  |  |  |
| Category | Enrichment Score | Term | FDR |
| **Cluster 1** | **5.6** |  |  |
| UP_KEYWORDS |  | Metal-binding | 2.6e-07 |
| UP_KEYWORDS |  | Zinc-finger | 5.2e-04 |
| **Cluster 2** | **4.2** |  |  |
| UP_KEYWORDS |  | Cell junction | 6.1e-04 |
| **Cluster 3** | **3.6** |  |  |
| UP_KEYWORDS |  | Transcription | 0.001 |
| UP_KEYWORDS |  | Transcription regulation | 0.004 |

Since the mitochondrial pathway was identified as a possible vulnerable target for eradicating spheroids (Table 2), we tested the impact of Oligomycin on spheroids. Oligomycin inhibits mitochondrial function and electron transport by blocking the proton channel for ATP-synthase. Oligomycin treatment only modestly inhibited growth of both HEYA8 cells grown as monolayers and spheroids at 5 µg/ml but reduced cell survival 81.4% and 70.4%, respectively, at 25 µg/ml (p < 0.0001 for both; Figure 4A), Cell-cell contact within spheroids was diminished at 5 µg/ml and spheroids were nearly completely dissociated at 25 µg/ml Oligomycin (Figure 4B).

**Figure 4.**
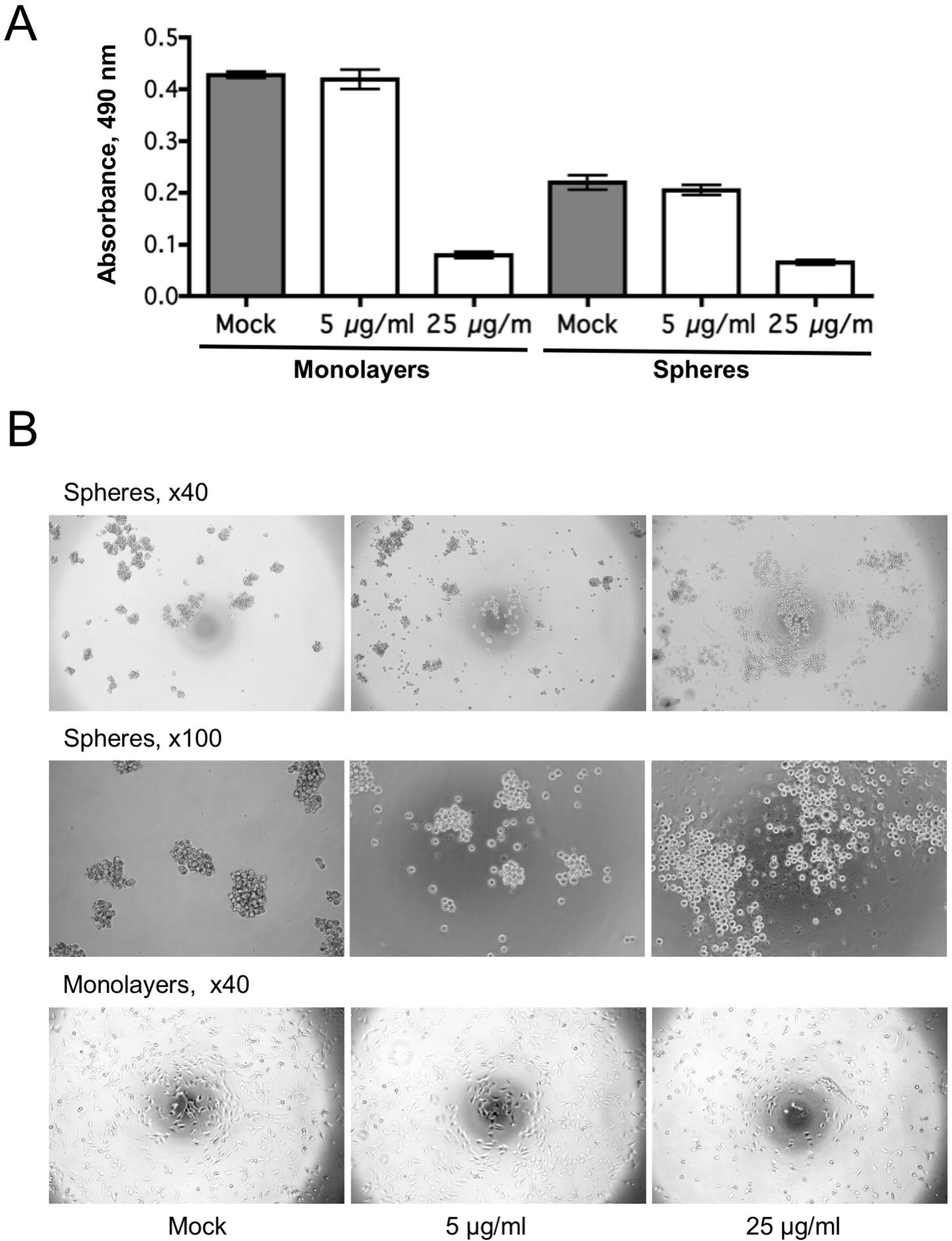
The mitochondrial inhibitor Oligomycin inhibits spheroid formation under serum-free spheroid culture conditions. (A) Relatively similar efficacy of Oligomycin toward HEYA8 monolayer cells and spheroids, with reduced cell survival at 25 µg/ml (p<0.0001 for both). (B) Spheroid formation is suppressed by Oligomycin. Upper (4X objective) and middle (10X objective) panel demonstrates that spheroid forming capacity was reduced by Oligomycin treatment (left, Mock; middle, 5 µg/ml; right, 25 µg/ml). Lower panel (4X objective) shows the corresponding monolayer cells.

Patients with platinum resistant/refractory ovarian cancer have poor clinical prognosis ^20^. Platinum-resistant ovarian cancer tends to relapse within the first six months after the end of first-line chemotherapy (carboplatin and/or paclitaxel) ^21^. We have shown that UCN-01 and Oltipraz function preferentially on slowly proliferating cells *in vitro*. Previous studies have shown that slow-cycling subpopulations of quiescent cancer cells include cancer stem-like cells and contribute to chemoresistance and recurrence ^22^. In order to test our hypothesis that UCN-01 and Oltipraz would be effective in treating platinum resistant/refractory ovarian cancer, we employed a novel mouse model of carboplatin resistance.

We used *in vivo* live imaging to visualize tumor growth and regression in our mouse model. We first generated stable cell lines that express luciferase (LUC) and green fluorescent protein (GFP) in CAOV2 ovarian cancer cells. The LUC/GFP positive cells (CAOV2-GFP/LUC) were enriched by flow cytometry for GFP positive signals. Forty mice were injected intraperitoneally with 3.5X10^5^ CAOV2-GFP/LUC cells. After seven days, 26 mice had visible tumors by *in vivo* live imaging with a mean of 3.2X10^7^ total photon flux. Six of these mice received no treatment (Arm 5). The remaining 20 of these tumor-bearing mice were treated once every four days with carboplatin at 80 mg/kg for a total of three rounds of treatment, until tumor signal was reduced to an average of 50% total photon flux (total photon flux from 3.2X10^7^ to 1.5X10^7^). The mice were then assigned to arms for the secondary treatment as indicated in Figure 5A. The secondary treatment regimen is described in detail in Materials and Methods. The dosing regimen for each mouse is shown in Table 3 and the average total photon flux from each experimental arm is summarized in **Table S6**. The mice receiving carboplatin followed by additional carboplatin (Arm 1) or no additional treatment (Arm 4) showed decreased tumor size from the primary carboplatin treatment, but tumor growth resumed after treatment was terminated (Figure 5A). We found that mice experienced a reduction in tumor burden after three rounds of treatments with carboplatin, but this improvement was reversed with further carboplatin treatment due to apparent acquired resistance. Mice in the experimental arms that received carboplatin primary treatment followed by UCN-01 (Arm 2) or Oltipraz (Arm 3) showed little to no evidence of tumor re-growth (Figure 5A **and** 5B). As expected, mice that received no treatment showed progressive tumor growth throughout the study and reached humane endpoints sooner than all other mice. Continued carboplatin treatment in Arm 4 showed some benefit relative to no treatment or no continued treatment (**Table S6**, Arm 5). At necropsy, the majority of mice receiving secondary UCN-01 or Oltipraz treatment were found to have no evidence of remaining disease, which confirmed our observations with live imaging. Similar to tumor distribution in ovarian cancer patients, most tumors were located on the peritoneum or omentum in the mice with detectable disease at necropsy, with fewer tumors attached to the fallopian tubes, ovaries, or intestines. Several mice in Arm 5 (no treatment) and Arm 1 (carboplatin followed by no treatment) developed malignant ascites. Overall, these data indicate that prolonged use of carboplatin *in vivo* can result in development of chemoresistance, whereas drugs that target slow proliferating cancer cells (like UCN-01 and oltipraz) may delay disease recurrence.

**Figure 5.**
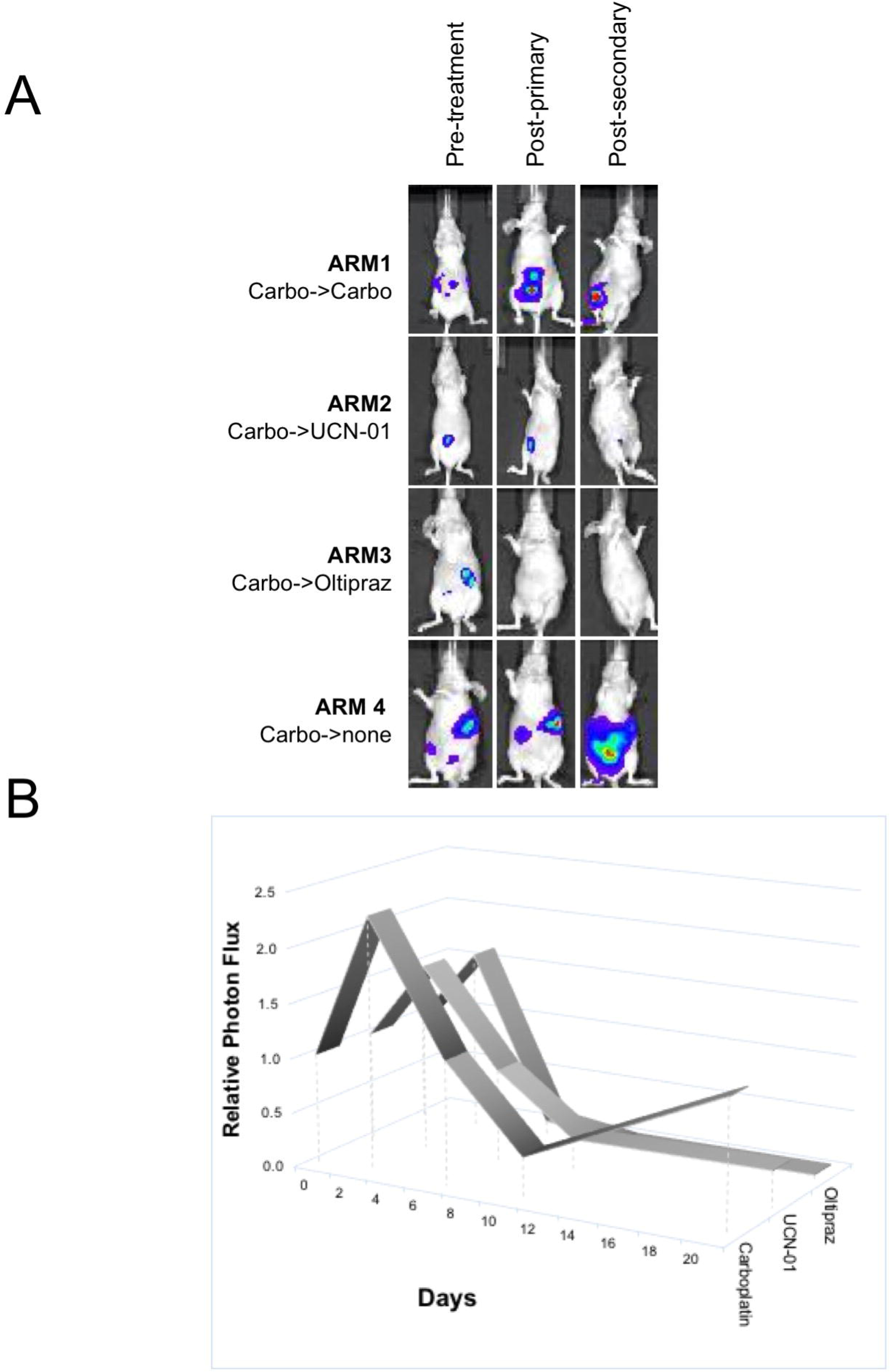
Delayed tumor regrowth *in vivo* with UCN-01 or Oltipraz as secondary treatment. (A) Representative live imaging of tumors in Arms 1-4. Arm 1: carboplatin primary followed by carboplatin secondary treatment; Arm 2: carboplatin primary followed by UCN-01 secondary treatment; Arm 3: carboplatin primary followed by Oltipraz secondary treatment; Arm 4: carboplatin primary followed by no secondary treatment; Arm 5 (not shown): no primary and no secondary treatment. (B) Average relative signal from tumors (photon flux) in Arms 1-3, normalized to the average tumor size at the onset of primary treatment (day 0), shown over the course of the study (days, x-axis).

**Table 3.**
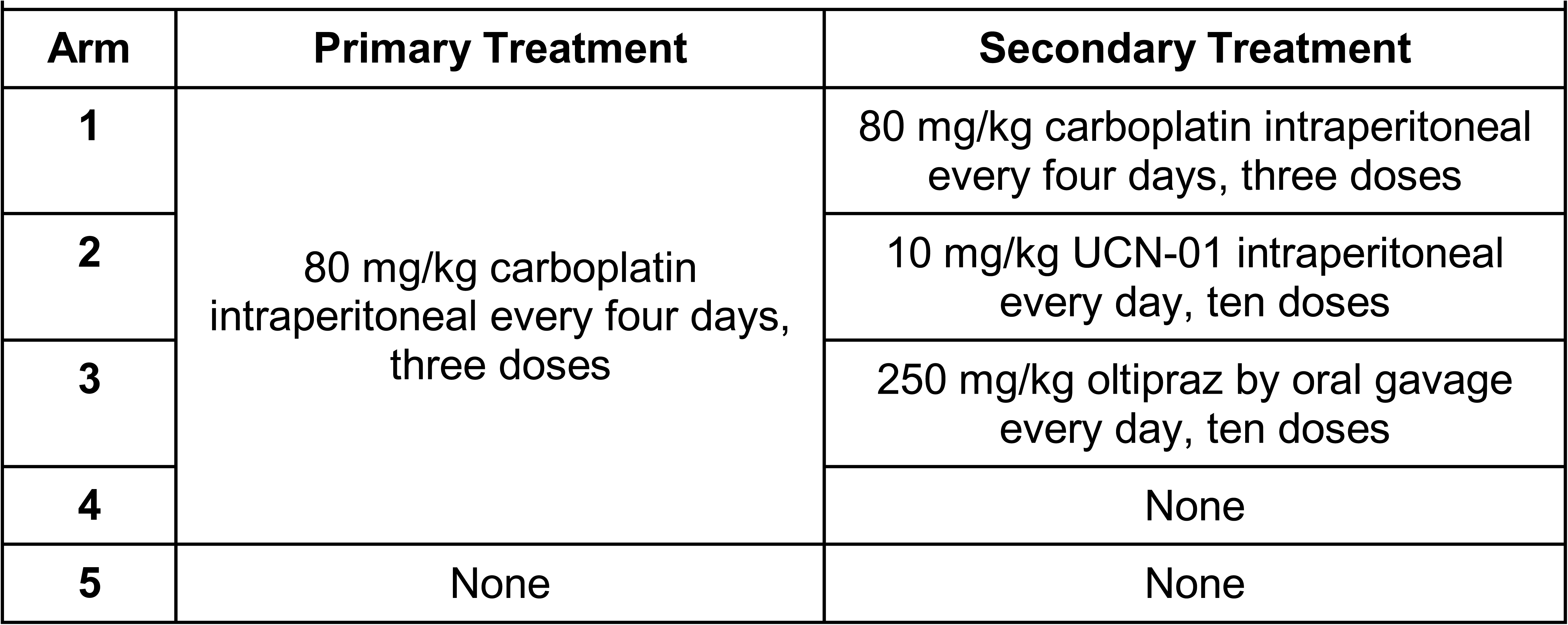
Preclinical Study Dosing Regimen

## DISCUSSION

Nearly inevitable relapse is a major problem in treating patients with advanced ovarian cancer ^5^. The concept of cancer cell dormancy has been put forward in hematological malignancies, breast and prostate cancer as an explanation for disease recurrence ^15, 23, 24^. According to this postulate, dormancy provides a source of latent cancer cells that resist conventional chemotherapy, since such drugs are primarily effective against rapidly proliferating cancer cells ^25^. Ovarian cancer patients don’t typically receive treatment between the completion of their surgery and primary chemotherapy and the point at which a recurrence is detected. This leaves a time frame during which residual cancer cells could modify their genetic and/or epigenetic profiles in order to regrow when systemic and/or local microenvironment conditions become favorable ^26^. Several *in vitro* models of dormancy have been reported ^27, 28^, but the molecular features accompanying tumor cell dormancy are poorly understood ^15, 23^. We showed that ovarian cancer cells that grow in an anchorage-independent manner in serum-free media are able to enter into a dormant state and can exit this state as many as three months later following transfer into regular culture conditions. The cellular morphology of dormant cells was very different from the same cells grown in regular culture, and this change in phenotype may involve gene mutations and/or epigenetic modifications ^29–31^. The peritoneal cavity is a common site of disease recurrence in ovarian cancer ^32^. Our findings support the notion that a specialized subpopulation of ovarian cancer cells remain viable as dormant cells in the peritoneal cavity, possibly for years, and ultimately cause recurrent disease. The serum-free spheroid culture model can be employed to clarify the signaling pathways required for the survival or reactivation of dormant cancer cells, although a major limitation is that this model may not accurately reflect the state of these cells *in vivo* ^33^.

Whether or not dormant cancer cells possess CSLC features is currently a topic of debate ^15, 34, 35^. In breast cancer, a majority of disseminated tumor cells persist in a dormant state in the bone marrow and are distinguished by having a CD44^+^/ CD24^-/low^ cell surface marker profile ^34^. We sought to enrich for ovarian CSLCs using the serum-free culture model reported previously ^14^, but failed to reproduce sustainable spheroids with self-renewal capability. This shortcoming may be because the cancer cells investigated did not contain CSLCs as previously discussed ^12^. CD44 is present on CSLCs of a number of cancer types, including breast, prostate, colon, and ovarian cancer ^14, 36–39^ as well as in cell lines ^12, 40^. In our study, CD44 expression was strongly and positively correlated with the capacity for anchorage-independent colony formation and the ability of spheroids to undergo passaging, although CD44 expression was not correlated with spheroid numbers. CD24, a cell surface marker that is absent on breast CSLCs, here discriminated the capacity for spheroid formation among the cancer cell lines. These data also suggests a relationship between CSCL marker expression and the ability to enter into a dormant state. It is well known that spheroid formation is determined by both intrinsic properties of cells and by the cellular microenvironment ^41^. Accordingly, we found that OVARY1847, OV90, PEO1, and OVCAR8 spheroid cells did not survive over a prolonged period in serum-free spheroid culture, while they could be expanded in regular media (**Table S1**). These findings suggest that the microenvironment indeed plays an important role in the selection of a specific population of cancer cells that enter into cellular dormancy.

Drug resistance remains a major problem in the treatment of recurrent ovarian cancer. Spheroid models have been used primarily as an *in vitro* model for three-dimensional solid tumors to investigate mechanisms of drug resistance ^42^. Spheroid cells are known to become refractory to conventional chemotherapeutic agents, especially to paclitaxel when compared to monolayer cells ^43, 44^. Therefore, eradication of spheroids within ascites may be quite clinically relevant to decrease the high rate of ovarian cancer recurrence. We previously reported that UCN-01 had higher efficacy against slow-proliferating and/or quiescent cancer cells ^16^, and that UCN-01 exhibits higher efficacy against ovarian cancer cells grown as spheroids. This finding is in stark contrast to the increased resistance exhibited by the spheroids to carboplatin and paclitaxel.

UCN-01 has been evaluated in clinical trials for various malignancies, including ovarian cancer ^45–49^. In the ovarian cancer trail, UCN-01 was delivered with topotecan in the recurrent setting, when tumor growth was already evident. In order to delay or prevent recurrent disease, it is likely important to deliver drugs like UCN-01 or Oltipraz when the cells are in a slow-proliferating or dormant state, before the cells begin recurrent growth. This approach has not yet been formally tested in clinical trials with an aim of targeting the residual dormant cancer cell population. Targeting this residual cell population may be a relevant strategy for improving survival of women with high grade invasive ovarian cancer.

Bioinformatic analyses of the genes showing differential expression between the spheroid-forming and non-spheroid forming ovarian cancer cell lines showed that genes involved in mitochondria functions were elevated and important to the survival of the spheroids in serum-free spheroid culture conditions. UCN-01 is an inhibitor of protein kinase C and cyclin-dependent kinases, and is also known to induce mitochondrial damage and apoptosis ^50, 51^. In support of the microarray data analysis, another mitochondrial inhibitor, Oligomycin, reduced cell survival in both monolayer cells and spheroids and also inhibited the ability of the cells in serum-free spheroid culture conditions to maintain their cell-cell contacts and spheroid phenotype. These findings are compatible with another report demonstrating that metabolism-related genes are up-regulated in compact spheroids rather than loose cellular aggregates ^52^. Such structural change induced by mitochondrial inhibitors may improve efficacy of conventional chemotherapeutic drugs by allowing their penetration and enhancing their distribution within spheroids. Since compounds toxic to mitochondria have been developed through clinical trials ^53^, it may be pertinent to investigate the role of the mitochondrial pathway in dormant cancer cells to determine targetability.

Our results from the xenograft mouse studies suggest that UCN01 and Oltipraz, and similar drugs that target slower proliferating cells, may have efficacy against residual ovarian cancer cells when delivered after completion of the primary chemotherapeutic regimen and prior to emergence of recurrent disease. Continued carboplatin treatment resulted in tumor regrowth, contrasting with the results from treating with UCN-01 and Oltipraz, in which tumor cells remaining following completion of the carboplatin regimen were apparently not able to initiate growth in the mice, at least during the duration of the experiment. It is likely important that treatment to target the residual cancer cells commenced soon after the completion of the primary chemotherapy. This may be a critically important but underappreciated point since the numbers of residual cancer cells are at a nadir, and presumably more easily targeted. The prior Phase I and II clinical trials in which UCN-01 was tested for efficacy against recurrent ovarian cancer initiated treatment after the recurrent disease was evident ^54, 55^. The conclusion was that although well tolerated, UCN-01 (and topotecan, given together in those studies) did not show anti-tumor activity. However, after the disease recurs, the cancer cells are again rapidly dividing, and according to our results here and those we have previously published ^16^, these actively proliferating cells would have shown resistance to UCN-01. This may explain the failure to achieve a response in those initial trials.

In conclusion, our results support that serum-free spheroid culture can be used as a model to examine the mechanisms of cancer cell dormancy. Ovarian cancer spheroid formation is associated with expression of the cancer stem-like cell markers, CD44 and CD133. The capacity for cells to form spheroids and enter into a dormant state is associated with mitochondrial gene functions, and the mitochondrial inhibitors, UCN-01 and Oligomycin, caused spheroid dissociation while spheroids were highly refractory to cisplatin and paclitaxel. Unlike continued tumor growth observed with prolonged treatment with carboplatin in our xenograft mouse model, UCN-01 and Oltipraz were inhibitory to tumor regrowth subsequent to carboplatin treatment. Dormant cancer cells that survive initial chemotherapy are likely to be the source of recurrent disease, and we contend that these cells need to be targeted *prior to* the onset of recurrent disease. Our findings suggest that targeting aspects of mitochondrial function may be a reasonable strategy for decreasing the high recurrence rate of advanced stage ovarian cancers.

## MATERIALS AND METHODS

### Cells

Twenty-five ovarian cancer cell lines were grown in RPMI-1640 medium with L-glutamine (Sigma-Aldrich; St. Louis, MO) supplemented with 10% fetal bovine serum in a humidified incubator with 5% CO_2_ at 37°C. Cell lines were genetically authenticated with each expansion at the Duke University DNA Analysis Facility to confirm identity of newly-prepared freezer stocks.

### Anchorage-independent growth

Assays for colony formation in soft agar were performed as described ^56^. Briefly, 2X RPMI media was prepared from powder and supplemented with fetal bovine serum and antibiotics (Invitrogen; Carlsbad, CA). A 1% agarose solution was made with the RPMI media using low-melting-temperature agarose (Invitrogen). One ml of 0.5% agarose was placed into each well of six-well tissue culture dishes and overlaid with 1 ml of 0.33% agarose prepared in 1x RPMI and containing 2 x 10^4^ cells. After three weeks incubation at 37°C in a humidified chamber with 5% CO_2_, colonies larger than 100 µm in diameter were counted. The colony number formed for each cell line was determined by averaging the number of colonies >100 µm that were counted in 10-20 microscopic fields at 100X magnification. Data were analyzed using Graphpad Prism software with a p value less than 0.05 considered significant.

### Spheroids

Single dissociated ovarian cancer cells derived from cell lines were plated at 500 cells/ml in 2 ml serum-free DMEM-F12 (GIBCO; Invitrogen) supplemented with 5 µg/ml insulin (SAFC Biosciences; Lenexa, KS), 20 ng/ml human recombinant EGF (Invitrogen), 10 ng/ml bFGF (R&D Systems; Minneapolis, MN), and 0.4% BSA using 6-well ultra-low attachment plates (Corning Life Sciences; Lowell, MA). After 7-10 days in culture, spheroids larger than 50 µm in diameter were counted. To assess the ability of spheroids to undergo serial passaging, dissociated ovarian cancer cells from spheroids were plated at 5X10^3^ cells/ml. Cells grown in serum-free spheroid culture were enzymatically dissociated by incubation in a trypsin-EDTA solution (0.025%) for ten minutes and underwent passaging every 7-10 days. Fresh medium (500 µl per well) was added twice a week. Cell viability was measured at each passage using trypan blue staining. The experiments were performed at least in triplicate.

### Malignant human ovarian tissues

Tumors were obtained from eight patients diagnosed with advanced ovarian cancer at the time of initial cytoreductive surgery at Duke University Medical Center, after receiving approval from the Duke University Institutional Review Board and following provision of written informed consent. Fresh tumors were minced and suspended in serum-free media with growth factors as described above. To further dissociate the tumors, 300 units per ml of both collagenase (Sigma-Aldrich) and hyaluronidase (EMD Biosciences; Gibbstown, NJ) were added followed by overnight incubation in a humidified chamber with 5% CO_2_. The cell suspension was filtered using a 40 µm cell strainer and red blood cells were removed using Histopaque 1077 (Sigma-Aldrich). The cells were resuspended in the media described above and plated on ultra-low attachment plates. To determine the ability of the cells within spheroids to undergo self-renewal, spheroids generated by eight ovarian cancer cell lines and seven primary tumors were dissociated following three to five passages. Cells were plated at single cell dilution into 96-well ultra-low attachment plates under serum-free spheroid culture conditions described above. Microscopic visualization confirmed the presence of one cell per well in the 96-well plates.

### Proliferation activity

HEYA8 cells (3X10^3^ or 6X10^3^) were seeded into individual wells of 96-well tissue culture plates in regular media or into individual wells of 96-well ultra-low attachment plates in serum-free spheroid culture media, respectively. To compare cell proliferation activity between these two conditions, a proliferation rate was calculated using the data obtained from MTT colorimetric assays (CellTiter 96® AQueous One Solution Cell Proliferation Assay; Promega; Madison, WI). Data from five replicates for at each time point were analyzed using Graphpad Prism software with a p value less than 0.05 considered significant.

For Ki67 staining, plates were collected after seven days of culture and fixed with 1:1 acetone:methanol. The cells were incubated with a monoclonal antibody to human Ki-67 antigen, clone MIB-1 (diluted 1:100; DAKO, Carpinteria, CA) at 4°C overnight. Immunodetection was carried out using the streptavidin-biotin based Multi-Link Super Sensitive Detection System (4plus Universal HPR Detection System from Biocare Medical; Concord, CA).

To investigate whether quiescent cancer cells in serum-free spheroid culture can resume growth in regular media, cells from cell lines and three primary tumors that underwent serial passage in serum-free spheroid culture were collected and plated on tissue-culture treated plates with regular media. Immunostaining with a pan-cytokeratin cocktail antibody, AE1/AE3 (Dako), was used to confirm the epithelial nature of the cancer cells. The Mouse Super Sensitive TM Negative Control, clone HK408-5R (BioGenex; San Ramon, CA) was used as a negative control.

### Drug sensitivity assays

UCN-01 was kindly provided by the National Cancer Institute. Cisplatin, paclitaxel and oligomycin were purchased from Sigma-Aldrich. Cells in regular media were seeded at 3X10^3^ cells per well into tissue culture-treated plates, and cells in serum-free sphere culture were plated at 6X10^3^ cells per well in 96-well ultra-low attachment plates. Drugs were added 24 hours later, and the cells were incubated in the presence of the drugs for an additional 24 hours (Oligomycin) or 48 hours (all other drugs). Six wells were used for each drug concentration. The cell survival ratio was determined by quantifying the reduction in growth compared with mock-treated cells at 48 hours using a standard MTT colorimetric assay (Promega AQueous One). The assay was repeated four times.

### Microarray data

We analyzed our previously generated Affymetrix HG-U133A DNA microarray data for ovarian cancer cell lines (Gene Expression Omnibus: GSE25429). Data analysis was performed using Partek Genomics Suite software (St.Louis, MO). Affymetrix CEL files were normalized using the Robust Multichip Average method ^57^. To investigate gene expression profiles that allow cancer cells to survive serum-free spheroid culture conditions, we identified differentially expressed genes between cell lines that have low (n=4) versus high (n=5) capacity to survive under these conditions. We used student t tests with an unadjusted p value cutoff of 0.05, which resulted in the identification of 2,799 probes. Supervised clustering was performed as previously described ^58^. Bioinformatics analyses of the genes differentially expressed between cell lines with low and high potential to survive serum-free spheroid culture conditions was done using DAVID 6.8 ^59, 60^.

### CAOV2-GFP/LUC cells

CAOV2 ovarian cancer cells were transduced with the dual reporter lentivector pGreenFire1 (pGF1-CMV; positive control) according to the protocol from the manufacturer (System Biosciences; Palo Alto, CA). This vector encodes the copepod green fluorescent protein (GFP) and firefly luciferase (LUC) under a constitutively active CMV promoter, and contains the puromycin selectable marker. Expression of GFP enables detection and isolation of cells harboring the construct and expression of LUC allows for monitoring of tumor growth and response using non-invasive live imaging. CAOV2-GFP/LUC cells were sorted for GFP positivity by flow cytometry. The GFP positive cells were expanded in cell culture prior to injection into nude mice.

### Xenograft tumors

The Duke University Institutional Animal Care & Use Committee approved this research. To generate xenograft tumors, 3.5X10^5^ CAOV2-GFP/LUC cells were injected intra-peritoneally (IP) into each of 40 healthy six to seven-week old female athymic NCr-nu/nu mice (NCI-Frederick). Live imaging was performed following administration of isoflurane using an XGI-8 Gas Anesthesia System. Tumor formation and regression with reatment were monitored using the IVIS 100 Optical Imaging System (Xenogen, Caliper Life Sciences) in the Duke Cancer Institute’s Optical Molecular Imaging and Analysis Core. Living Image 2.6.1 (Caliper Life Sciences) software was used for quantitative analysis. Tumor flux measurements, defined as photon/sec/cm^2^/steradian, were the raw output values from the luminescent imaging data.

Following establishment of a measurable tumor, treatment with carboplatin was performed every four days for three weeks at 80 mg/kg of mouse body weight. Tumor signal was monitored by IVIS imaging. Twenty-six mice were responsive to the carboplatin treatment and were randomly grouped for secondary treatment (Table 3). Arm 1 mice (N=4, Carbo/None) received no further treatment. Arm 2 mice (N=5, Carbo/UCN-01) received UCN-01 treatment at 10 mg/kg IP for five days, no treatment for two days, then another cycle of UCN-01 every day for another five days, totaling 10 doses. Arm 3 mice (N=6, Carbo/Oltipraz) received Oltipraz at 250 mg/kg via oral gavage every day for six days, no treatment for one day, then another cycle of Oltipraz every day for six more days, totaling 12 doses. Arm 4 mice (N=5, Carbo/Carbo) received continue carboplatin treatment at 80 mg/kg IP every 4 days for 2 weeks. Arm 5 mice (N=6, None/None) received no treatment from the beginning to the end of the study. Mice were imaged weekly during the secondary treatment period.

## ACKNOWLEDGMENTS

We gratefully acknowledge Dr. Seiichi Mori for providing antibodies for flow cytometry, and Dr. Jeffrey R. Marks for assistance with micrography. This research was supported by the Gail Parkins Ovarian Cancer Awareness Fund and the Ovarian Cancer Research Fund.

## SUPPLEMENTARY INFORMATION

**Figure S1.** Alteration of drug sensitivity in seven ovarian cancer cell lines grown as monolayer cells versus spheroids. UCN-01 (10 nM or 100 nM, left) exhibits higher efficacy against spheroids, in contrast to cisplatin (10 µM, middle) and paclitaxel (100 nM, right). M, monolayer cells; S, spheroids.

**Table S1.** Spheroid-forming characteristics of ovarian cancer cell lines. Shown are the average numbers of spheroids formed (>50 µM in diameter) after 7-10 days in culture from initial plating of 500 dissociated cells/ml under spheroid culture conditions, and the ability of the spheroids to undergo serial passage under spheroid culture conditions.

**Table S2.** Differentially expressed genes that exhibit increased expression in the high spheroid-forming capacity cell lines that discriminate between high and low potential to survive serum-free spheroid culture.

**Table S3.** Differentially expressed genes that exhibit increased expression in the low spheroid-forming capacity cell lines that discriminate between high and low potential to survive serum-free spheroid culture.

**Table S4.** Functional annotations (from the Database for Annotation, Visualization and Integrated Discovery (DAVID) v6.8) for genes with higher expression in cells with high spheroid-forming capacity relative to cells with low spheroid-forming capacity.

**Table S5.** Functional annotations (from DAVID) for genes with higher expression in cells with low spheroid-forming capacity relative to cells with high spheroid-forming capacity.

**Table S6.** Average photon flux measurements from the five experimental arms of the mouse study at each measured time point.

