## Supplementary material for "Targeting dormant ovarian cancer cells *in vitro* and in an *in vivo* model of platinum resistance": Table S1

| Table S1. Spheroid-forming characteristics of ovarian cancer cell lines | | |
|---|---|---|
| **Cell line** | **Spheroid number** | **Maximum serial passages** |
| OVARY1847 | 0 | 0 |
| OV90 | 0 | 1 |
| PEO1 | 1 | 0 |
| OVCAR8 | 1 | 2 |
| HEYA8 | 3 | ≥5 |
| HEY | 4 | ≥5 |
| HEYC2 | 4 | ≥5 |
| OVCAR2 | 9 | ≥5 |
| CAOV2 | 10 | ≥5 |
