## Supplementary material for "Targeting dormant ovarian cancer cells *in vitro* and in an *in vivo* model of platinum resistance": Table S2

### Table S2. Genes with increased expression in high spheroid-forming capacity cell lines

| U133A probe | Gene Symbol | Avg NSF* | Avg SF* | Fold Change | t test p |
|---|---|---|---|---|---|
| 206172_at | IL13RA2 | 4.894 | 9.638 | 22.5 | 0.034 |
| 209160_at | AKR1C3 | 6.252 | 10.594 | 18.9 | 0.008 |
| 212764_at | --- | 6.024 | 10.313 | 18.4 | 0.007 |
| 201645_at | TNC | 5.297 | 9.412 | 16.9 | 0.001 |
| 204955_at | SRPX | 6.811 | 10.891 | 16.6 | 0.002 |
| 209781_s_at | KHDRBS3 | 5.922 | 9.791 | 15.0 | 0.000 |
| 205083_at | AOX1 | 4.918 | 8.600 | 13.6 | 0.044 |
| 204115_at | GNG11 | 6.960 | 10.557 | 12.9 | 0.009 |
| 213338_at | RIS1 | 5.812 | 9.398 | 12.9 | 0.002 |
| 201288_at | ARHGDIB | 6.808 | 10.329 | 12.4 | 0.036 |
| 204823_at | NAV3 | 6.267 | 9.767 | 12.3 | 0.012 |
| 215177_s_at | ITGA6 | 6.672 | 10.123 | 11.9 | 0.003 |
| 213030_s_at | PLXNA2 | 5.143 | 8.541 | 11.5 | 0.002 |
| 209276_s_at | GLRX | 6.890 | 10.232 | 11.2 | 0.000 |
| 212758_s_at | TCF8 | 5.446 | 8.747 | 10.9 | 0.001 |
| 211709_s_at | CLEC11A | 5.197 | 8.404 | 10.3 | 0.039 |
| 213139_at | SNAI2 | 5.376 | 8.574 | 10.2 | 0.014 |
| 201150_s_at | TIMP3 | 4.706 | 7.900 | 10.2 | 0.016 |
| 209101_at | CTGF | 9.073 | 12.234 | 10.0 | 0.012 |
| 205935_at | FOXF1 | 4.650 | 7.809 | 10.0 | 0.000 |
| 204472_at | GEM | 5.805 | 8.900 | 9.6 | 0.001 |
| 213435_at | SATB2 | 6.218 | 9.306 | 9.5 | 0.000 |
| 203939_at | NT5E | 8.290 | 11.280 | 8.9 | 0.003 |
| 207419_s_at | RAC2 | 5.933 | 8.858 | 8.6 | 0.003 |
| 215913_s_at | GULP1 | 5.947 | 8.870 | 8.5 | 0.001 |
| 218211_s_at | MLPH | 8.220 | 11.042 | 8.0 | 0.030 |
| 203058_s_at | PAPSS2 | 8.244 | 11.063 | 7.9 | 0.049 |
| 201280_s_at | DAB2 | 5.864 | 8.666 | 7.8 | 0.029 |
| 210757_x_at | DAB2 | 5.991 | 8.729 | 7.5 | 0.010 |
| 201279_s_at | DAB2 | 5.515 | 8.242 | 7.4 | 0.006 |
| 210299_s_at | FHL1 | 6.150 | 8.871 | 7.4 | 0.034 |
| 210875_s_at | TCF8 | 4.804 | 7.502 | 7.3 | 0.004 |
| 219628_at | WIG1 | 7.038 | 9.730 | 7.3 | 0.037 |
| 221911_at | ETV1 | 6.186 | 8.877 | 7.2 | 0.006 |
| 203300_x_at | AP1S2 | 6.349 | 9.004 | 7.0 | 0.041 |
| 221606_s_at | NSBP1 | 5.790 | 8.395 | 6.8 | 0.019 |
| 211343_s_at | COL13A1 | 5.186 | 7.784 | 6.8 | 0.011 |
| 204319_s_at | RGS10 | 7.940 | 10.523 | 6.7 | 0.047 |
| 202742_s_at | PRKACB | 6.071 | 8.631 | 6.6 | 0.013 |
| 201147_s_at | TIMP3 | 5.219 | 7.755 | 6.4 | 0.033 |

| | | | | | |
|---|---|---|---|---|---|
| 210298_x_at | FHL1 | 5.049 | 7.554 | 6.3 | 0.002 |
| 202202_s_at | LAMA4 | 5.344 | 7.836 | 6.2 | 0.022 |
| 220615_s_at | MLSTD1 | 7.196 | 9.687 | 6.2 | 0.028 |
| 215447_at | --- | 4.736 | 7.178 | 6.0 | 0.000 |
| 209406_at | BAG2 | 8.090 | 10.494 | 5.8 | 0.029 |
| 203504_s_at | ABCA1 | 5.035 | 7.436 | 5.8 | 0.001 |
| 205498_at | GHR | 4.798 | 7.183 | 5.7 | 0.004 |
| 205005_s_at | NMT2 | 6.590 | 8.953 | 5.6 | 0.021 |
| 205006_s_at | NMT2 | 6.016 | 8.337 | 5.4 | 0.024 |
| 214505_s_at | FHL1 | 6.540 | 8.854 | 5.4 | 0.016 |
| 219682_s_at | TBX3 | 4.918 | 7.224 | 5.3 | 0.002 |
| 203409_at | DDB2 | 6.593 | 8.899 | 5.3 | 0.012 |
| 201539_s_at | FHL1 | 5.804 | 8.094 | 5.2 | 0.001 |
| 213421_x_at | PRSS3 | 5.808 | 8.094 | 5.2 | 0.007 |
| 219282_s_at | TRPV2 | 5.816 | 8.091 | 5.2 | 0.035 |
| 213010_at | PRKCDBP | 6.349 | 8.621 | 5.2 | 0.022 |
| 213603_s_at | RAC2 | 8.321 | 10.585 | 5.1 | 0.033 |
| 207463_x_at | PRSS3 | 5.039 | 7.302 | 5.1 | 0.000 |
| 213428_s_at | COL6A1 | 7.506 | 9.747 | 5.0 | 0.030 |
| 213880_at | LGR5 | 4.793 | 7.031 | 5.0 | 0.017 |
| 210317_s_at | YWHAE | 7.927 | 10.163 | 5.0 | 0.004 |
| 220892_s_at | PSAT1 | 8.974 | 11.202 | 5.0 | 0.000 |
| 203320_at | LNK | 7.955 | 10.158 | 4.9 | 0.002 |
| 37965_at | PARVB | 6.203 | 8.401 | 4.8 | 0.003 |
| 208478_s_at | BAX | 6.512 | 8.708 | 4.8 | 0.001 |
| 218000_s_at | PHLDA1 | 6.078 | 8.256 | 4.7 | 0.010 |
| 204220_at | GMFG | 5.496 | 7.650 | 4.6 | 0.047 |
| 214772_at | C11orf41 | 5.009 | 7.151 | 4.6 | 0.041 |
| 205808_at | ASPH | 6.986 | 9.121 | 4.6 | 0.003 |
| 200958_s_at | SDCBP | 10.649 | 12.779 | 4.5 | 0.027 |
| 201278_at | DAB2 | 6.319 | 8.442 | 4.5 | 0.044 |
| 219973_at | ARSJ | 5.091 | 7.200 | 4.4 | 0.019 |
| 203505_at | ABCA1 | 6.254 | 8.363 | 4.4 | 0.013 |
| 203989_x_at | F2R | 5.924 | 8.029 | 4.4 | 0.016 |
| 203059_s_at | PAPSS2 | 6.452 | 8.552 | 4.4 | 0.003 |
| 202143_s_at | COPS8 | 8.536 | 10.626 | 4.4 | 0.002 |
| 217999_s_at | PHLDA1 | 6.514 | 8.599 | 4.3 | 0.031 |
| 205462_s_at | HPCAL1 | 6.933 | 9.017 | 4.3 | 0.017 |
| 217790_s_at | SSR3 | 8.596 | 10.664 | 4.3 | 0.027 |
| 217959_s_at | TRAPPC4 | 9.230 | 11.296 | 4.3 | 0.001 |
| 209629_s_at | NXT2 | 5.789 | 7.852 | 4.3 | 0.012 |
| 217996_at | PHLDA1 | 8.911 | 10.964 | 4.2 | 0.040 |
| 219260_s_at | C17orf81 | 6.078 | 8.131 | 4.2 | 0.009 |
| 203132_at | RB1 | 7.769 | 9.802 | 4.1 | 0.036 |
| 217960_s_at | TOMM22 | 7.740 | 9.770 | 4.1 | 0.001 |

| | | | | | |
|---|---|---|---|---|---|
| 218644_at | PLEK2 | 8.635 | 10.660 | 4.1 | 0.028 |
| 201337_s_at | VAMP3 | 7.890 | 9.900 | 4.0 | 0.040 |
| 219654_at | PTPLA | 7.256 | 9.262 | 4.0 | 0.038 |
| 218398_at | MRPS30 | 8.680 | 10.681 | 4.0 | 0.004 |
| 200776_s_at | BZW1 /// LOC151579 | 9.766 | 11.760 | 4.0 | 0.007 |
| 220606_s_at | C17orf48 | 5.289 | 7.270 | 3.9 | 0.001 |
| 211833_s_at | BAX | 6.634 | 8.603 | 3.9 | 0.006 |
| 201823_s_at | RNF14 | 6.810 | 8.771 | 3.8 | 0.039 |
| 213116_at | NEK3 | 6.895 | 8.852 | 3.8 | 0.030 |
| 221618_s_at | TAF9B /// LOC653068 | 5.621 | 7.576 | 3.8 | 0.009 |
| 201311_s_at | SH3BGRL | 6.958 | 8.911 | 3.8 | 0.029 |
| 209679_s_at | LOC57228 | 7.967 | 9.902 | 3.7 | 0.000 |
| 201559_s_at | CLIC4 | 7.709 | 9.643 | 3.7 | 0.029 |
| 213548_s_at | CDV3 | 6.512 | 8.443 | 3.7 | 0.018 |
| 206113_s_at | RAB5A | 7.379 | 9.305 | 3.7 | 0.046 |
| 202693_s_at | STK17A | 8.632 | 10.555 | 3.7 | 0.006 |
| 203207_s_at | MTFR1 | 7.823 | 9.745 | 3.7 | 0.001 |
| 211681_s_at | PDLIM5 | 5.105 | 7.024 | 3.7 | 0.006 |
| 214835_s_at | SUCLG2 | 5.989 | 7.906 | 3.7 | 0.002 |
| 218718_at | PDGFC | 9.953 | 11.866 | 3.7 | 0.004 |
| 217997_at | PHLDA1 | 8.328 | 10.240 | 3.7 | 0.043 |
| 210732_s_at | LGALS8 | 7.216 | 9.121 | 3.6 | 0.044 |
| 205803_s_at | TRPC1 | 4.999 | 6.903 | 3.6 | 0.000 |
| 210448_s_at | P2RX5 | 6.811 | 8.713 | 3.6 | 0.006 |
| 215997_s_at | CUL4B | 6.117 | 8.019 | 3.6 | 0.000 |
| 206157_at | PTX3 | 7.548 | 9.448 | 3.6 | 0.045 |
| 37966_at | PARVB | 5.731 | 7.627 | 3.6 | 0.011 |
| 205018_s_at | MBNL2 | 5.443 | 7.334 | 3.6 | 0.010 |
| 214446_at | ELL2 | 6.628 | 8.519 | 3.6 | 0.001 |
| 217785_s_at | YKT6 | 5.737 | 7.628 | 3.6 | 0.002 |
| 213309_at | PLCL2 | 5.969 | 7.858 | 3.6 | 0.013 |
| 201554_x_at | GYG1 | 8.947 | 10.829 | 3.5 | 0.015 |
| 216591_s_at | SDHC /// LOC642502 | 7.384 | 9.265 | 3.5 | 0.006 |
| 204629_at | PARVB | 5.946 | 7.819 | 3.5 | 0.014 |
| 215772_x_at | SUCLG2 | 8.259 | 10.129 | 3.5 | 0.003 |
| 202533_s_at | DHFR /// LOC643509 /// LOC653874 | 7.047 | 8.910 | 3.5 | 0.001 |
| 215707_s_at | PRNP | 7.706 | 9.550 | 3.4 | 0.006 |
| 210653_s_at | BCKDHB | 5.231 | 7.074 | 3.4 | 0.000 |
| 209488_s_at | RBPMS | 5.977 | 7.818 | 3.4 | 0.009 |
| 219522_at | FJX1 | 7.665 | 9.495 | 3.3 | 0.049 |
| 217445_s_at | GART | 6.352 | 8.179 | 3.3 | 0.004 |
| 206693_at | IL7 | 4.513 | 6.334 | 3.3 | 0.012 |
| 212843_at | NCAM1 | 4.799 | 6.618 | 3.3 | 0.037 |
| 214007_s_at | PTK9 | 7.864 | 9.683 | 3.3 | 0.031 |

| | | | | | |
|---|---|---|---|---|---|
| 203242_s_at | PDLIM5 | 7.279 | 9.089 | 3.3 | 0.005 |
| 200727_s_at | ACTR2 | 8.603 | 10.409 | 3.3 | 0.048 |
| 213325_at | PVRL3 | 5.767 | 7.572 | 3.3 | 0.009 |
| 206233_at | B4GALT6 | 6.358 | 8.154 | 3.2 | 0.006 |
| 200796_s_at | MCL1 | 5.194 | 6.988 | 3.2 | 0.008 |
| 206544_x_at | SMARCA2 | 5.456 | 7.241 | 3.2 | 0.016 |
| 219553_at | NME7 | 8.240 | 10.025 | 3.2 | 0.016 |
| 213887_s_at | POLR2E | 9.099 | 10.883 | 3.2 | 0.023 |
| 206662_at | GLRX | 8.574 | 10.356 | 3.2 | 0.001 |
| 209208_at | MPDU1 | 7.864 | 9.645 | 3.2 | 0.002 |
| 216574_s_at | RPE /// LOC440001 /// LOC649755 | 6.536 | 8.316 | 3.2 | 0.009 |
| 220334_at | RGS17 | 5.624 | 7.369 | 3.0 | 0.007 |
| 215794_x_at | GLUD2 | 7.624 | 9.358 | 3.0 | 0.000 |
| 220773_s_at | GPHN | 6.537 | 8.272 | 3.0 | 0.014 |
| 208854_s_at | STK24 | 9.378 | 11.109 | 3.0 | 0.018 |
| 204344_s_at | SEC23A | 7.958 | 9.687 | 3.0 | 0.013 |
| 210868_s_at | ELOVL6 | 6.011 | 7.729 | 2.9 | 0.031 |
| 219545_at | KCTD14 | 6.377 | 8.091 | 2.9 | 0.035 |
| 212459_x_at | SUCLG2 | 8.573 | 10.283 | 2.9 | 0.010 |
| 212792_at | DPY19L1 | 8.592 | 10.296 | 2.9 | 0.011 |
| 215617_at | --- | 5.855 | 7.556 | 2.9 | 0.039 |
| 212091_s_at | COL6A1 | 5.368 | 7.066 | 2.9 | 0.004 |
| 211092_s_at | NF2 | 4.764 | 6.455 | 2.9 | 0.038 |
| 221638_s_at | STX16 | 5.989 | 7.680 | 2.9 | 0.009 |
| 202166_s_at | PPP1R2 | 7.995 | 9.686 | 2.9 | 0.033 |
| 203402_at | KCNAB2 | 5.532 | 7.217 | 2.8 | 0.047 |
| 218526_s_at | RANGNRF | 7.034 | 8.707 | 2.8 | 0.009 |
| 211089_s_at | NEK3 | 5.367 | 7.036 | 2.8 | 0.009 |
| 201870_at | TOMM34 | 8.057 | 9.712 | 2.7 | 0.000 |
| 207088_s_at | SLC25A11 | 7.583 | 9.236 | 2.7 | 0.002 |
| 220238_s_at | KLHL7 | 6.250 | 7.899 | 2.7 | 0.025 |
| 210257_x_at | CUL4B | 6.944 | 8.589 | 2.7 | 0.005 |
| 209487_at | RBPMS | 6.694 | 8.335 | 2.7 | 0.035 |
| 210935_s_at | WDR1 | 6.940 | 8.581 | 2.7 | 0.045 |
| 216466_at | NAV3 | 5.066 | 6.706 | 2.7 | 0.041 |
| 201656_at | ITGA6 | 8.023 | 9.662 | 2.7 | 0.047 |
| 219006_at | C6orf66 | 7.768 | 9.402 | 2.7 | 0.011 |
| 213150_at | HOXA10 | 6.757 | 8.390 | 2.7 | 0.049 |
| 202118_s_at | CPNE3 | 9.357 | 10.990 | 2.7 | 0.037 |
| 201316_at | PSMA2 | 7.190 | 8.822 | 2.7 | 0.004 |
| 211954_s_at | RANBP5 | 10.486 | 12.112 | 2.6 | 0.013 |
| 203148_s_at | TRIM14 | 7.212 | 8.837 | 2.6 | 0.020 |
| 221503_s_at | KPNA3 | 9.081 | 10.701 | 2.6 | 0.023 |
| 220477_s_at | C20orf30 | 8.811 | 10.424 | 2.6 | 0.012 |

| | | | | | |
|---|---|---|---|---|---|
| 221727_at | --- | 8.109 | 9.720 | 2.6 | 0.033 |
| 208828_at | POLE3 | 8.323 | 9.932 | 2.6 | 0.018 |
| 219555_s_at | C16orf60 | 9.446 | 11.050 | 2.6 | 0.000 |
| 218852_at | C14orf10 | 8.634 | 10.238 | 2.6 | 0.043 |
| 214483_s_at | ARFIP1 | 5.909 | 7.505 | 2.5 | 0.002 |
| 221078_s_at | KIAA1212 | 5.941 | 7.534 | 2.5 | 0.024 |
| 212838_at | DNMBP | 9.433 | 11.024 | 2.5 | 0.025 |
| 214544_s_at | SNAP23 | 6.359 | 7.948 | 2.5 | 0.003 |
| 211080_s_at | NEK2 | 5.426 | 7.013 | 2.5 | 0.000 |
| 203626_s_at | SKP2 | 5.946 | 7.526 | 2.5 | 0.023 |
| 202731_at | PDCD4 | 5.900 | 7.475 | 2.5 | 0.000 |
| 204969_s_at | RDX | 6.053 | 7.628 | 2.5 | 0.010 |
| 203196_at | ABCC4 | 8.750 | 10.323 | 2.5 | 0.032 |
| 205429_s_at | MPP6 | 7.004 | 8.573 | 2.5 | 0.012 |
| 202142_at | COPS8 | 8.870 | 10.438 | 2.5 | 0.003 |
| 202165_at | PPP1R2 | 7.217 | 8.783 | 2.5 | 0.014 |
| 213117_at | KLHL9 | 6.074 | 7.634 | 2.4 | 0.014 |
| 214512_s_at | SUB1 | 10.106 | 11.665 | 2.4 | 0.013 |
| 219155_at | PITPNC1 | 6.910 | 8.468 | 2.4 | 0.028 |
| 209799_at | PRKAA1 | 6.941 | 8.498 | 2.4 | 0.008 |
| 215780_s_at | SET /// LOC642869 | 9.957 | 11.513 | 2.4 | 0.001 |
| 219390_at | FKBP14 | 6.554 | 8.110 | 2.4 | 0.012 |
| 207686_s_at | CASP8 | 6.043 | 7.598 | 2.4 | 0.002 |
| 203790_s_at | HRSP12 | 8.559 | 10.111 | 2.4 | 0.009 |
| 206308_at | DNMT2 | 5.173 | 6.725 | 2.4 | 0.005 |
| 209131_s_at | SNAP23 | 5.844 | 7.394 | 2.4 | 0.004 |
| 208097_s_at | TXNDC | 8.834 | 10.378 | 2.4 | 0.007 |
| 221551_x_at | ST6GALNAC4 | 6.158 | 7.700 | 2.4 | 0.000 |
| 213133_s_at | GCSH /// LOC653763 | 8.454 | 9.989 | 2.4 | 0.034 |
| 205677_s_at | DLEU1 | 8.851 | 10.385 | 2.4 | 0.035 |
| 204768_s_at | FEN1 | 9.173 | 10.702 | 2.3 | 0.012 |
| 218549_s_at | FAM82B | 8.667 | 10.194 | 2.3 | 0.037 |
| 218013_x_at | DCTN4 | 7.392 | 8.907 | 2.3 | 0.001 |
| 210457_x_at | HMGA1 | 6.467 | 7.981 | 2.3 | 0.013 |
| 218992_at | C9orf46 | 7.221 | 8.734 | 2.3 | 0.025 |
| 202497_x_at | SLC2A3 | 5.894 | 7.406 | 2.3 | 0.038 |
| 201801_s_at | SLC29A1 | 6.721 | 8.230 | 2.3 | 0.007 |
| 210024_s_at | UBE2E3 | 10.405 | 11.912 | 2.3 | 0.016 |
| 220937_s_at | ST6GALNAC4 | 6.024 | 7.531 | 2.3 | 0.004 |
| 201508_at | IGFBP4 | 9.268 | 10.773 | 2.3 | 0.025 |
| 209512_at | HSDL2 | 6.668 | 8.163 | 2.2 | 0.043 |
| 200722_s_at | GPIAP1 | 8.904 | 10.399 | 2.2 | 0.003 |
| 209891_at | SPBC25 | 7.546 | 9.041 | 2.2 | 0.018 |
| 205196_s_at | AP1S1 | 6.621 | 8.105 | 2.2 | 0.038 |
| 211212_s_at | ORC5L | 5.613 | 7.092 | 2.2 | 0.043 |

| | | | | | |
|---|---|---|---|---|---|
| 208741_at | SAP18 | 5.878 | 7.352 | 2.2 | 0.001 |
| 205219_s_at | GALK2 | 6.835 | 8.308 | 2.2 | 0.032 |
| 222192_s_at | FLJ21820 | 7.053 | 8.525 | 2.2 | 0.030 |
| 215195_at | PRKCA | 7.846 | 9.312 | 2.1 | 0.005 |
| 202973_x_at | FAM13A1 | 6.934 | 8.400 | 2.1 | 0.049 |
| 209009_at | ESD | 10.576 | 12.040 | 2.1 | 0.025 |
| 211955_at | RANBP5 | 10.362 | 11.825 | 2.1 | 0.026 |
| 205071_x_at | XRCC4 | 6.925 | 8.387 | 2.1 | 0.016 |
| 216253_s_at | PARVB | 5.342 | 6.803 | 2.1 | 0.006 |
| 205401_at | AGPS | 8.103 | 9.564 | 2.1 | 0.003 |
| 212624_s_at | CHN1 | 7.505 | 8.965 | 2.1 | 0.026 |
| 202593_s_at | MIR16 | 8.487 | 9.945 | 2.1 | 0.009 |
| 219927_at | C14orf111 | 5.295 | 6.753 | 2.1 | 0.005 |
| 218439_s_at | COMMD10 | 7.653 | 9.108 | 2.1 | 0.002 |
| 209363_s_at | SURB7 | 8.205 | 9.660 | 2.1 | 0.001 |
| 221688_s_at | IMP3 | 8.597 | 10.051 | 2.1 | 0.003 |
| 211814_s_at | CCNE2 | 6.131 | 7.581 | 2.1 | 0.031 |
| 210296_s_at | PXMP3 | 8.665 | 10.113 | 2.1 | 0.000 |
| 203340_s_at | SLC25A12 | 8.364 | 9.811 | 2.1 | 0.012 |
| 218597_s_at | C10orf70 | 9.867 | 11.312 | 2.1 | 0.008 |
| 214941_s_at | LOC642455 | 8.227 | 9.662 | 2.1 | 0.045 |
| 203348_s_at | ETV5 | 7.742 | 9.177 | 2.1 | 0.028 |
| 200942_s_at | HSBP1 | 9.893 | 11.327 | 2.1 | 0.026 |
| 207360_s_at | NTSR1 | 5.136 | 6.565 | 2.0 | 0.032 |
| 221423_s_at | YIPF5 | 8.133 | 9.562 | 2.0 | 0.027 |
| 208476_s_at | FRMD4A | 5.367 | 6.794 | 2.0 | 0.033 |
| 218111_s_at | CMAS | 7.764 | 9.189 | 2.0 | 0.027 |
| 216804_s_at | PDLIM5 | 8.782 | 10.201 | 2.0 | 0.035 |
| 219502_at | NEIL3 | 7.007 | 8.425 | 2.0 | 0.038 |
| 202190_at | CSTF1 | 7.389 | 8.806 | 2.0 | 0.001 |
| 202904_s_at | LSM5 | 7.203 | 8.619 | 2.0 | 0.023 |
| 203405_at | DSCR2 | 8.827 | 10.243 | 2.0 | 0.038 |
| 219281_at | MSRA | 6.514 | 7.930 | 2.0 | 0.009 |
| 204944_at | PTPRG | 6.667 | 8.082 | 2.0 | 0.021 |
| 200604_s_at | PRKAR1A | 8.640 | 10.051 | 2.0 | 0.012 |
| 201504_s_at | TSN | 7.723 | 9.134 | 2.0 | 0.002 |
| 219596_at | THAP10 | 6.472 | 7.881 | 2.0 | 0.010 |
| 202213_s_at | CUL4B | 7.675 | 9.082 | 2.0 | 0.000 |
| 210041_s_at | PGM3 | 7.203 | 8.609 | 2.0 | 0.033 |
| 209536_s_at | EHD4 | 9.378 | 10.783 | 2.0 | 0.028 |
| 210154_at | ME2 | 7.265 | 8.661 | 1.9 | 0.007 |
| 203621_at | NDUFB5 | 10.299 | 11.693 | 1.9 | 0.042 |
| 214276_at | KLF12 | 5.559 | 6.953 | 1.9 | 0.005 |
| 201095_at | DAP | 8.604 | 9.998 | 1.9 | 0.033 |
| 204128_s_at | RFC3 | 7.986 | 9.378 | 1.9 | 0.026 |

| | | | | | |
|---|---|---|---|---|---|
| 209628_at | NXT2 | 8.369 | 9.760 | 1.9 | 0.029 |
| 211922_s_at | CAT | 7.641 | 9.032 | 1.9 | 0.026 |
| 200798_x_at | MCL1 | 8.621 | 10.011 | 1.9 | 0.021 |
| 206788_s_at | CBFB | 7.074 | 8.462 | 1.9 | 0.014 |
| 201512_s_at | TOMM70A | 8.091 | 9.479 | 1.9 | 0.031 |
| 200946_x_at | GLUD1 | 9.810 | 11.197 | 1.9 | 0.010 |
| 201633_s_at | CYB5B | 7.976 | 9.362 | 1.9 | 0.026 |
| 206654_s_at | POLR3G | 6.271 | 7.654 | 1.9 | 0.042 |
| 209080_x_at | TXNL2 | 9.709 | 11.090 | 1.9 | 0.013 |
| 213041_s_at | ATP5D | 9.107 | 10.487 | 1.9 | 0.026 |
| 217998_at | PHLDA1 | 5.737 | 7.116 | 1.9 | 0.018 |
| 208074_s_at | AP2S1 | 10.803 | 12.181 | 1.9 | 0.014 |
| 204298_s_at | LOX | 5.441 | 6.818 | 1.9 | 0.035 |
| 201490_s_at | PPIF | 8.454 | 9.831 | 1.9 | 0.011 |
| 203622_s_at | LOC56902 | 8.960 | 10.334 | 1.9 | 0.004 |
| 217299_s_at | NBN | 7.386 | 8.761 | 1.9 | 0.000 |
| 205796_at | TCP11L1 | 5.971 | 7.341 | 1.9 | 0.005 |
| 205802_at | TRPC1 | 6.692 | 8.059 | 1.9 | 0.006 |
| 203198_at | CDK9 | 6.979 | 8.345 | 1.9 | 0.025 |
| 203771_s_at | BLVRA | 6.116 | 7.481 | 1.9 | 0.012 |
| 216375_s_at | ETV5 | 6.059 | 7.423 | 1.9 | 0.014 |
| 217707_x_at | SMARCA2 | 6.056 | 7.418 | 1.9 | 0.026 |
| 201457_x_at | BUB3 | 10.657 | 12.016 | 1.8 | 0.003 |
| 201462_at | SCRN1 | 9.736 | 11.094 | 1.8 | 0.025 |
| 217911_s_at | BAG3 | 8.950 | 10.307 | 1.8 | 0.018 |
| 209445_x_at | FLJ10803 | 9.730 | 11.086 | 1.8 | 0.002 |
| 205701_at | IPO8 | 5.883 | 7.239 | 1.8 | 0.002 |
| 218491_s_at | THYN1 | 8.385 | 9.739 | 1.8 | 0.023 |
| 219099_at | C12orf5 | 8.490 | 9.844 | 1.8 | 0.001 |
| 209592_s_at | WDR68 | 7.141 | 8.493 | 1.8 | 0.021 |
| 206923_at | PRKCA | 5.018 | 6.368 | 1.8 | 0.006 |
| 204950_at | CARD8 | 6.147 | 7.493 | 1.8 | 0.032 |
| 204283_at | FARS2 | 6.184 | 7.528 | 1.8 | 0.047 |
| 219291_at | DTWD1 | 5.492 | 6.833 | 1.8 | 0.009 |
| 203466_at | MPV17 | 8.022 | 9.362 | 1.8 | 0.013 |
| 202654_x_at | 7-Mar | 9.052 | 10.391 | 1.8 | 0.000 |
| 200739_s_at | SUMO3 | 8.926 | 10.265 | 1.8 | 0.032 |
| 211747_s_at | LSM5 | 10.345 | 11.681 | 1.8 | 0.003 |
| 208742_s_at | SAP18 | 10.736 | 12.069 | 1.8 | 0.000 |
| 218449_at | C4orf20 | 7.441 | 8.773 | 1.8 | 0.040 |
| 205053_at | PRIM1 | 8.843 | 10.174 | 1.8 | 0.002 |
| 205579_at | HRH1 | 5.509 | 6.839 | 1.8 | 0.014 |
| 206542_s_at | SMARCA2 | 6.315 | 7.644 | 1.8 | 0.031 |
| 221487_s_at | ENSA | 6.753 | 8.079 | 1.8 | 0.008 |
| 210285_x_at | WTAP | 7.546 | 8.868 | 1.7 | 0.001 |

| | | | | | |
|---|---|---|---|---|---|
| 217958_at | TRAPPC4 | 9.969 | 11.290 | 1.7 | 0.024 |
| 201222_s_at | RAD23B | 11.250 | 12.569 | 1.7 | 0.003 |
| 218001_at | MRPS2 | 8.989 | 10.308 | 1.7 | 0.049 |
| 215037_s_at | BCL2L1 | 6.100 | 7.419 | 1.7 | 0.049 |
| 210009_s_at | GOSR2 | 7.359 | 8.675 | 1.7 | 0.021 |
| 205214_at | STK17B | 5.736 | 7.052 | 1.7 | 0.001 |
| 213262_at | SACS | 9.460 | 10.775 | 1.7 | 0.023 |
| 221502_at | KPNA3 | 9.920 | 11.235 | 1.7 | 0.013 |
| 213461_at | NUDT21 | 7.245 | 8.558 | 1.7 | 0.043 |
| 208764_s_at | ATP5G2 | 11.229 | 12.541 | 1.7 | 0.024 |
| 201308_s_at | 11-Sep | 6.664 | 7.975 | 1.7 | 0.039 |
| 222088_s_at | SLC2A3 | 5.706 | 7.016 | 1.7 | 0.021 |
| 200615_s_at | AP2B1 | 8.819 | 10.130 | 1.7 | 0.001 |
| 218640_s_at | PLEKHF2 | 7.651 | 8.960 | 1.7 | 0.020 |
| 200883_at | UQCRC2 | 9.077 | 10.382 | 1.7 | 0.001 |
| 205217_at | TIMM8A | 7.238 | 8.537 | 1.7 | 0.012 |
| 203333_at | KIFAP3 | 8.075 | 9.372 | 1.7 | 0.033 |
| 212471_at | KIAA0241 | 5.618 | 6.912 | 1.7 | 0.000 |
| 209825_s_at | UCK2 | 8.659 | 9.951 | 1.7 | 0.004 |
| 201120_s_at | PGRMC1 | 8.747 | 10.039 | 1.7 | 0.010 |
| 212552_at | HPCAL1 | 9.155 | 10.445 | 1.7 | 0.017 |
| 221573_at | C7orf25 | 5.487 | 6.776 | 1.7 | 0.002 |
| 210405_x_at | TNFRSF10B | 6.687 | 7.976 | 1.7 | 0.027 |
| 221583_s_at | KCNMA1 | 5.204 | 6.492 | 1.7 | 0.043 |
| 215910_s_at | FNDC3A | 5.936 | 7.222 | 1.7 | 0.015 |
| 219979_s_at | C11orf73 | 6.554 | 7.840 | 1.7 | 0.009 |
| 91684_g_at | EXOSC4 | 7.877 | 9.158 | 1.6 | 0.032 |
| 213465_s_at | PPP1R7 | 9.685 | 10.960 | 1.6 | 0.001 |
| 213617_s_at | C18orf10 | 7.881 | 9.156 | 1.6 | 0.006 |
| 215464_s_at | TAX1BP3 | 9.520 | 10.793 | 1.6 | 0.028 |
| 203339_at | SLC25A12 | 7.207 | 8.480 | 1.6 | 0.010 |
| 202697_at | NUDT21 | 9.309 | 10.580 | 1.6 | 0.005 |
| 219324_at | TRIOBP /// MGC3731 | 7.299 | 8.570 | 1.6 | 0.009 |
| 221770_at | RPE | 8.192 | 9.459 | 1.6 | 0.027 |
| 218479_s_at | XPO4 | 6.140 | 7.404 | 1.6 | 0.043 |
| 219938_s_at | PSTPIP2 | 5.346 | 6.609 | 1.6 | 0.003 |
| 213581_at | PDCD2 | 8.839 | 10.102 | 1.6 | 0.029 |
| 210415_s_at | ODF2 | 5.697 | 6.958 | 1.6 | 0.002 |
| 221082_s_at | NDRG3 | 6.329 | 7.589 | 1.6 | 0.043 |
| 210813_s_at | XRCC4 | 5.942 | 7.202 | 1.6 | 0.038 |
| 215591_at | SATB2 | 5.007 | 6.264 | 1.6 | 0.005 |
| 202141_s_at | COPS8 | 10.216 | 11.472 | 1.6 | 0.019 |
| 201317_s_at | PSMA2 | 10.976 | 12.230 | 1.6 | 0.009 |
| 209287_s_at | CDC42EP3 | 7.387 | 8.635 | 1.6 | 0.037 |
| 216218_s_at | PLCL2 | 5.047 | 6.296 | 1.6 | 0.000 |

| | | | | | |
|---|---|---|---|---|---|
| 211752_s_at | NDUFS7 | 8.680 | 9.927 | 1.6 | 0.050 |
| 213757_at | EIF5A | 9.339 | 10.586 | 1.6 | 0.018 |
| 218748_s_at | EXOC5 | 6.975 | 8.222 | 1.6 | 0.046 |
| 216236_s_at | SLC2A3 | 7.418 | 8.663 | 1.6 | 0.025 |
| 212993_at | --- | 7.769 | 9.014 | 1.6 | 0.029 |
| 206348_s_at | PDK3 | 5.842 | 7.083 | 1.5 | 0.015 |
| 219056_at | DLEU8 | 5.523 | 6.763 | 1.5 | 0.014 |
| 204883_s_at | HUS1 | 7.697 | 8.936 | 1.5 | 0.026 |
| 208733_at | RAB2 | 5.104 | 6.341 | 1.5 | 0.002 |
| 210250_x_at | ADSL | 10.140 | 11.375 | 1.5 | 0.026 |
| 200729_s_at | ACTR2 | 10.149 | 11.382 | 1.5 | 0.012 |
| 202862_at | FAH | 6.666 | 7.899 | 1.5 | 0.047 |
| 212410_at | EFHA1 | 9.309 | 10.541 | 1.5 | 0.032 |
| 201096_s_at | ARF4 | 9.834 | 11.066 | 1.5 | 0.031 |
| 220060_s_at | C12orf48 | 6.070 | 7.300 | 1.5 | 0.015 |
| 218252_at | CKAP2 | 10.140 | 11.369 | 1.5 | 0.043 |
| 217808_s_at | MAPKAP1 | 8.338 | 9.567 | 1.5 | 0.005 |
| 221547_at | PRPF18 | 7.488 | 8.716 | 1.5 | 0.045 |
| 210984_x_at | EGFR | 6.261 | 7.487 | 1.5 | 0.047 |
| 204093_at | CCNH | 10.099 | 11.323 | 1.5 | 0.023 |
| 213655_at | PAFAH1B1 | 11.178 | 12.400 | 1.5 | 0.023 |
| 200712_s_at | MAPRE1 | 9.026 | 10.248 | 1.5 | 0.034 |
| 204479_at | OSTF1 | 7.690 | 8.910 | 1.5 | 0.045 |
| 58780_s_at | FLJ10357 | 6.656 | 7.874 | 1.5 | 0.039 |
| 212246_at | MCFD2 | 8.726 | 9.944 | 1.5 | 0.007 |
| 207334_s_at | TGFBR2 | 5.418 | 6.635 | 1.5 | 0.005 |
| 209233_at | EMG1 | 10.060 | 11.276 | 1.5 | 0.017 |
| 212003_at | C1orf144 | 6.830 | 8.045 | 1.5 | 0.008 |
| 202407_s_at | PRPF31 | 7.284 | 8.499 | 1.5 | 0.014 |
| 200751_s_at | HNRPC | 10.028 | 11.241 | 1.5 | 0.035 |
| 201706_s_at | PEX19 | 6.728 | 7.938 | 1.5 | 0.001 |
| 210667_s_at | C21orf33 | 6.671 | 7.881 | 1.5 | 0.023 |
| 208732_at | RAB2 | 8.342 | 9.548 | 1.5 | 0.004 |
| 219267_at | GLTP | 6.608 | 7.812 | 1.5 | 0.031 |
| 203299_s_at | AP1S2 /// LOC653653 /// LOC654127 | 5.162 | 6.366 | 1.4 | 0.016 |
| 202695_s_at | STK17A | 5.946 | 7.149 | 1.4 | 0.001 |
| 209567_at | RRS1 | 10.192 | 11.394 | 1.4 | 0.000 |
| 218163_at | MCTS1 | 10.194 | 11.395 | 1.4 | 0.013 |
| 203740_at | MPHOSPH6 | 10.031 | 11.232 | 1.4 | 0.008 |
| 203147_s_at | TRIM14 | 6.398 | 7.597 | 1.4 | 0.050 |
| 205091_x_at | RECQL | 7.940 | 9.137 | 1.4 | 0.049 |
| 219715_s_at | TDP1 | 8.256 | 9.453 | 1.4 | 0.001 |
| 217356_s_at | PGK1 | 10.952 | 12.149 | 1.4 | 0.035 |
| 202899_s_at | SFRS3 | 11.412 | 12.608 | 1.4 | 0.005 |

| | | | | | |
|---|---|---|---|---|---|
| 201686_x_at | API5 | 7.138 | 8.333 | 1.4 | 0.025 |
| 212409_s_at | TOR1AIP1 | 6.376 | 7.571 | 1.4 | 0.009 |
| 205446_s_at | ATF2 | 5.728 | 6.922 | 1.4 | 0.001 |
| 220199_s_at | C1orf80 | 8.008 | 9.202 | 1.4 | 0.014 |
| 208296_x_at | TNFAIP8 | 6.153 | 7.346 | 1.4 | 0.003 |
| 219599_at | PRO1843 | 7.351 | 8.544 | 1.4 | 0.016 |
| 221521_s_at | GINS2 | 9.126 | 10.318 | 1.4 | 0.010 |
| 208911_s_at | PDHB | 8.981 | 10.172 | 1.4 | 0.018 |
| 200987_x_at | PSME3 | 9.418 | 10.609 | 1.4 | 0.032 |
| 218122_s_at | SENP2 | 7.143 | 8.334 | 1.4 | 0.043 |
| 203176_s_at | TFAM | 8.573 | 9.762 | 1.4 | 0.022 |
| 219819_s_at | MRPS28 | 10.115 | 11.302 | 1.4 | 0.027 |
| 202019_s_at | LANCL1 | 6.945 | 8.131 | 1.4 | 0.025 |
| 209484_s_at | C1orf48 | 8.628 | 9.814 | 1.4 | 0.021 |
| 221737_at | GNA12 | 6.100 | 7.283 | 1.4 | 0.020 |
| 218104_at | TEX10 | 8.782 | 9.961 | 1.4 | 0.016 |
| 206501_x_at | ETV1 | 5.337 | 6.514 | 1.4 | 0.013 |
| 220122_at | MCTP1 | 5.321 | 6.496 | 1.4 | 0.025 |
| 203439_s_at | STC2 | 6.792 | 7.966 | 1.4 | 0.049 |
| 200612_s_at | AP2B1 | 8.170 | 9.343 | 1.4 | 0.004 |
| 200890_s_at | SSR1 | 8.255 | 9.428 | 1.4 | 0.006 |
| 218993_at | RNMTL1 | 7.417 | 8.589 | 1.4 | 0.035 |
| 209715_at | CBX5 | 8.195 | 9.366 | 1.4 | 0.001 |
| 205501_at | --- | 4.937 | 6.106 | 1.4 | 0.022 |
| 214435_x_at | RALA | 8.702 | 9.871 | 1.4 | 0.019 |
| 212216_at | PREPL | 8.273 | 9.441 | 1.4 | 0.014 |
| 213760_s_at | ZNF330 | 7.097 | 8.262 | 1.4 | 0.018 |
| 201383_s_at | NBR1 /// LOC653347 | 7.760 | 8.921 | 1.3 | 0.044 |
| 218550_s_at | LRRC20 | 6.848 | 8.007 | 1.3 | 0.011 |
| 211382_s_at | TACC2 | 7.206 | 8.360 | 1.3 | 0.009 |
| 203565_s_at | MNAT1 | 8.004 | 9.158 | 1.3 | 0.040 |
| 214336_s_at | COPA | 6.333 | 7.486 | 1.3 | 0.004 |
| 201192_s_at | PITPNA | 8.811 | 9.963 | 1.3 | 0.032 |
| 206686_at | PDK1 | 5.581 | 6.733 | 1.3 | 0.001 |
| 212398_at | RDX | 8.952 | 10.103 | 1.3 | 0.037 |
| 204186_s_at | PPID | 8.210 | 9.358 | 1.3 | 0.042 |
| 219231_at | NCOA6IP | 6.244 | 7.391 | 1.3 | 0.007 |
| 208910_s_at | C1QBP | 11.089 | 12.236 | 1.3 | 0.004 |
| 217821_s_at | WBP11 | 9.003 | 10.150 | 1.3 | 0.036 |
| 216397_s_at | BOP1 /// LOC653119 | 7.449 | 8.594 | 1.3 | 0.021 |
| 212214_at | OPA1 | 8.709 | 9.853 | 1.3 | 0.016 |
| 206920_s_at | GLE1L | 6.662 | 7.807 | 1.3 | 0.002 |
| 209362_at | SURB7 | 8.330 | 9.472 | 1.3 | 0.007 |
| 217883_at | C2orf25 | 11.348 | 12.489 | 1.3 | 0.000 |
| 207845_s_at | ANAPC10 | 7.385 | 8.525 | 1.3 | 0.032 |

| | | | | | |
|---|---|---|---|---|---|
| 202884_s_at | PPP2R1B | 6.634 | 7.772 | 1.3 | 0.027 |
| 201614_s_at | RUVBL1 | 9.307 | 10.444 | 1.3 | 0.024 |
| 214959_s_at | API5 | 7.482 | 8.617 | 1.3 | 0.007 |
| 211318_s_at | RAE1 | 9.267 | 10.402 | 1.3 | 0.002 |
| 208631_s_at | HADHA | 9.975 | 11.106 | 1.3 | 0.005 |
| 202730_s_at | PDCD4 | 7.214 | 8.345 | 1.3 | 0.000 |
| 200737_at | PGK1 | 9.514 | 10.644 | 1.3 | 0.044 |
| 218077_s_at | ZDHHC3 | 5.632 | 6.761 | 1.3 | 0.044 |
| 201742_x_at | SFRS1 | 10.559 | 11.685 | 1.3 | 0.007 |
| 213822_s_at | UBE3B | 7.842 | 8.967 | 1.3 | 0.030 |
| 205397_x_at | SMAD3 | 5.976 | 7.098 | 1.3 | 0.030 |
| 208972_s_at | ATP5G1 | 9.078 | 10.199 | 1.3 | 0.003 |
| 210654_at | TNFRSF10D | 5.065 | 6.187 | 1.3 | 0.025 |
| 221960_s_at | RAB2 | 6.506 | 7.626 | 1.3 | 0.005 |
| 202740_at | ACY1 | 7.817 | 8.936 | 1.3 | 0.036 |
| 204305_at | MIPEP | 6.577 | 7.696 | 1.3 | 0.025 |
| 218160_at | NDUFA8 | 9.646 | 10.764 | 1.3 | 0.045 |
| 209203_s_at | BICD2 | 6.545 | 7.663 | 1.2 | 0.029 |
| 217457_s_at | RAP1GDS1 | 7.197 | 8.314 | 1.2 | 0.010 |
| 200927_s_at | RAB14 | 9.081 | 10.196 | 1.2 | 0.043 |
| 200023_s_at | EIF3S5 | 11.503 | 12.618 | 1.2 | 0.046 |
| 202246_s_at | CDK4 | 10.188 | 11.303 | 1.2 | 0.001 |
| 219960_s_at | UCHL5 | 8.694 | 9.808 | 1.2 | 0.011 |
| 200723_s_at | GPIAP1 | 10.569 | 11.682 | 1.2 | 0.006 |
| 218027_at | MRPL15 | 10.858 | 11.971 | 1.2 | 0.001 |
| 212418_at | ELF1 | 7.772 | 8.883 | 1.2 | 0.002 |
| 205078_at | PIGF | 7.558 | 8.668 | 1.2 | 0.037 |
| 211602_s_at | TRPC1 | 5.523 | 6.633 | 1.2 | 0.002 |
| 208876_s_at | PAK2 | 7.686 | 8.795 | 1.2 | 0.011 |
| 206491_s_at | NAPA | 7.706 | 8.815 | 1.2 | 0.000 |
| 212595_s_at | DAZAP2 | 9.210 | 10.319 | 1.2 | 0.004 |
| 215253_s_at | DSCR1 | 5.240 | 6.346 | 1.2 | 0.001 |
| 208351_s_at | MAPK1 | 6.456 | 7.562 | 1.2 | 0.004 |
| 209078_s_at | TXN2 | 8.000 | 9.104 | 1.2 | 0.025 |
| 203560_at | GGH | 10.124 | 11.227 | 1.2 | 0.023 |
| 221263_s_at | SF3B5 | 10.296 | 11.397 | 1.2 | 0.020 |
| 210260_s_at | TNFAIP8 | 6.517 | 7.617 | 1.2 | 0.005 |
| 207829_s_at | BNIP1 | 6.853 | 7.953 | 1.2 | 0.005 |
| 217717_s_at | YWHAB | 10.540 | 11.640 | 1.2 | 0.001 |
| 218185_s_at | ARMC1 | 9.035 | 10.134 | 1.2 | 0.015 |
| 218653_at | SLC25A15 | 7.763 | 8.860 | 1.2 | 0.043 |
| 209971_x_at | JTV1 | 10.167 | 11.263 | 1.2 | 0.038 |
| 209622_at | STK16 | 5.807 | 6.901 | 1.2 | 0.002 |
| 204386_s_at | MRP63 | 11.148 | 12.243 | 1.2 | 0.017 |
| 216005_at | TNC | 5.013 | 6.107 | 1.2 | 0.011 |

| | | | | | |
|---|---|---|---|---|---|
| 209177_at | C3orf60 | 8.883 | 9.978 | 1.2 | 0.023 |
| 202138_x_at | JTV1 | 9.473 | 10.567 | 1.2 | 0.028 |
| 204899_s_at | SAP30 | 5.461 | 6.554 | 1.2 | 0.047 |
| 212194_s_at | TM9SF4 | 8.548 | 9.639 | 1.2 | 0.039 |
| 221531_at | WDR61 | 7.588 | 8.678 | 1.2 | 0.017 |
| 204207_s_at | RNGTT | 7.203 | 8.293 | 1.2 | 0.038 |
| 203171_s_at | KIAA0409 | 7.023 | 8.111 | 1.2 | 0.031 |
| 202168_at | TAF9 | 10.001 | 11.089 | 1.2 | 0.041 |
| 209520_s_at | NCBP1 | 9.317 | 10.404 | 1.2 | 0.014 |
| 213144_at | GOSR2 | 6.283 | 7.370 | 1.2 | 0.001 |
| 212432_at | GRPEL1 | 8.968 | 10.053 | 1.2 | 0.012 |
| 213047_x_at | SET | 11.056 | 12.141 | 1.2 | 0.000 |
| 209375_at | XPC | 6.928 | 8.009 | 1.2 | 0.041 |
| 210379_s_at | TLK1 | 5.812 | 6.894 | 1.2 | 0.036 |
| 212527_at | D15Wsu75e | 7.311 | 8.390 | 1.2 | 0.020 |
| 206232_s_at | B4GALT6 | 4.835 | 5.913 | 1.2 | 0.025 |
| 206474_at | PCTK2 | 5.348 | 6.424 | 1.2 | 0.001 |
| 221957_at | PDK3 | 5.603 | 6.676 | 1.2 | 0.032 |
| 214205_x_at | TXNL2 | 5.554 | 6.627 | 1.2 | 0.008 |
| 219497_s_at | BCL11A | 5.485 | 6.557 | 1.1 | 0.025 |
| 218119_at | TIMM23 /// LOC653252 | 9.017 | 10.089 | 1.1 | 0.003 |
| 208967_s_at | AK2 | 9.747 | 10.817 | 1.1 | 0.047 |
| 203816_at | DGUOK | 7.302 | 8.365 | 1.1 | 0.042 |
| 202823_at | TCEB1 | 8.021 | 9.084 | 1.1 | 0.004 |
| 36830_at | MIPEP | 6.285 | 7.347 | 1.1 | 0.036 |
| 201872_s_at | ABCE1 | 10.474 | 11.535 | 1.1 | 0.024 |
| 203267_s_at | DRG2 | 7.597 | 8.657 | 1.1 | 0.024 |
| 219176_at | FLJ22555 | 9.523 | 10.580 | 1.1 | 0.048 |
| 201073_s_at | SMARCC1 | 6.397 | 7.453 | 1.1 | 0.045 |
| 202226_s_at | CRK | 8.669 | 9.723 | 1.1 | 0.036 |
| 202691_at | SNRPD1 | 8.606 | 9.660 | 1.1 | 0.008 |
| 201380_at | CRTAP | 7.357 | 8.408 | 1.1 | 0.004 |
| 204699_s_at | C1orf107 | 6.900 | 7.947 | 1.1 | 0.035 |
| 203926_x_at | ATP5D | 6.933 | 7.980 | 1.1 | 0.032 |
| 209853_s_at | PSME3 | 9.264 | 10.307 | 1.1 | 0.028 |
| 212213_x_at | OPA1 | 8.472 | 9.514 | 1.1 | 0.017 |
| 219073_s_at | OSBPL10 | 9.163 | 10.204 | 1.1 | 0.026 |
| 210734_x_at | MAX | 6.728 | 7.767 | 1.1 | 0.032 |
| 209344_at | TPM4 | 9.912 | 10.951 | 1.1 | 0.008 |
| 204833_at | ATG12 | 7.390 | 8.428 | 1.1 | 0.018 |
| 202535_at | FADD | 8.656 | 9.692 | 1.1 | 0.049 |
| 212420_at | ELF1 | 8.215 | 9.250 | 1.1 | 0.045 |
| 219177_at | BXDC2 | 6.894 | 7.927 | 1.1 | 0.029 |
| 202906_s_at | NBN | 8.547 | 9.581 | 1.1 | 0.042 |
| 215058_at | MGC24039 | 5.425 | 6.458 | 1.1 | 0.015 |

| | | | | | |
|---|---|---|---|---|---|
| 202900_s_at | NUP88 | 9.514 | 10.546 | 1.1 | 0.009 |
| 217053_x_at | ETV1 | 5.496 | 6.528 | 1.1 | 0.027 |
| 208848_at | ADH5 | 8.626 | 9.658 | 1.1 | 0.050 |
| 212243_at | GRINL1A /// Gcom1 | 7.377 | 8.408 | 1.1 | 0.029 |
| 201652_at | COPS5 | 10.225 | 11.253 | 1.1 | 0.003 |
| 219575_s_at | PDF /// COG8 | 8.416 | 9.444 | 1.1 | 0.030 |
| 208879_x_at | PRPF6 | 7.215 | 8.242 | 1.1 | 0.020 |
| 203897_at | LOC57149 | 8.335 | 9.360 | 1.1 | 0.050 |
| 205330_at | MN1 | 5.053 | 6.078 | 1.1 | 0.022 |
| 205817_at | SIX1 | 5.701 | 6.726 | 1.0 | 0.041 |
| 202348_s_at | TOR1A | 8.089 | 9.112 | 1.0 | 0.019 |
| 202144_s_at | ADSL | 10.580 | 11.602 | 1.0 | 0.039 |
| 204194_at | BACH1 | 6.562 | 7.583 | 1.0 | 0.003 |
| 214744_s_at | RPL23 | 6.905 | 7.926 | 1.0 | 0.023 |
| 202595_s_at | LEPROTL1 | 8.429 | 9.449 | 1.0 | 0.038 |
| 205631_at | KIAA0586 | 7.324 | 8.343 | 1.0 | 0.005 |
| 209157_at | DNAJA2 | 9.602 | 10.621 | 1.0 | 0.037 |
| 219924_s_at | ZMYM6 | 7.646 | 8.664 | 1.0 | 0.012 |
| 206667_s_at | SCAMP1 | 4.918 | 5.934 | 1.0 | 0.017 |
| 32723_at | CSTF1 | 6.835 | 7.851 | 1.0 | 0.002 |
| 218336_at | PFDN2 | 10.238 | 11.252 | 1.0 | 0.014 |
| 221452_s_at | TMEM14B | 9.827 | 10.841 | 1.0 | 0.033 |
| 202483_s_at | RANBP1 | 10.527 | 11.541 | 1.0 | 0.012 |
| 203105_s_at | DNM1L | 9.742 | 10.755 | 1.0 | 0.006 |
| 221207_s_at | NBEA | 5.978 | 6.990 | 1.0 | 0.042 |
| 200079_s_at | KARS | 10.948 | 11.960 | 1.0 | 0.003 |
| 203150_at | RABEPK | 8.708 | 9.718 | 1.0 | 0.032 |
| 212857_x_at | SUB1 | 11.504 | 12.514 | 1.0 | 0.043 |
| 217915_s_at | C15orf15 | 10.733 | 11.742 | 1.0 | 0.012 |
| 219688_at | BBS7 | 6.290 | 7.296 | 1.0 | 0.019 |
| 206342_x_at | IDS | 7.588 | 8.595 | 1.0 | 0.016 |
| 206558_at | SIM2 | 6.454 | 7.460 | 1.0 | 0.012 |
| 205372_at | PLAG1 | 5.973 | 6.980 | 1.0 | 0.016 |
| 218225_at | SITPEC | 6.468 | 7.474 | 1.0 | 0.006 |
| 203109_at | UBE2M | 8.595 | 9.600 | 1.0 | 0.045 |
| 204212_at | ACOT8 | 6.455 | 7.460 | 1.0 | 0.006 |
| 217540_at | --- | 5.127 | 6.132 | 1.0 | 0.028 |
| 200042_at | HSPC117 | 9.711 | 10.716 | 1.0 | 0.032 |
| 212481_s_at | TPM4 | 9.045 | 10.050 | 1.0 | 0.005 |
| 204460_s_at | RAD1 | 8.318 | 9.322 | 1.0 | 0.048 |
| 201521_s_at | NCBP2 | 8.760 | 9.763 | 1.0 | 0.043 |
| 218894_s_at | FLJ10292 | 7.688 | 8.690 | 1.0 | 0.002 |
| 203243_s_at | PDLIM5 | 8.984 | 9.985 | 1.0 | 0.032 |
| 201921_at | GNG10 /// LOC552891 /// LOC653503 | 10.439 | 11.441 | 1.0 | 0.026 |

| | | | | | |
|---|---|---|---|---|---|
| 211150_s_at | DLAT | 9.069 | 10.069 | 1.0 | 0.004 |
| 209464_at | AURKB | 9.317 | 10.317 | 1.0 | 0.008 |
| 209049_s_at | PRKCBP1 | 9.777 | 10.776 | 1.0 | 0.030 |
| 201821_s_at | TIMM17A | 9.097 | 10.095 | 1.0 | 0.041 |
| 207827_x_at | SNCA | 6.559 | 7.557 | 1.0 | 0.039 |
| 215952_s_at | OAZ1 | 11.831 | 12.829 | 1.0 | 0.037 |
| 201234_at | ILK | 9.541 | 10.538 | 1.0 | 0.022 |
| 218258_at | POLR1D | 10.344 | 11.341 | 1.0 | 0.006 |
| 220183_s_at | NUDT6 | 5.824 | 6.819 | 1.0 | 0.009 |
| 221570_s_at | METTL5 | 10.825 | 11.820 | 1.0 | 0.023 |
| 203265_s_at | MAP2K4 | 6.142 | 7.136 | 1.0 | 0.012 |
| 205566_at | ABHD2 | 6.123 | 7.117 | 1.0 | 0.012 |
| 202623_at | C14orf11 | 8.789 | 9.781 | 1.0 | 0.042 |
| 211684_s_at | DYNC1I2 | 11.116 | 12.108 | 1.0 | 0.007 |
| 211121_s_at | DOK1 | 4.935 | 5.927 | 1.0 | 0.001 |
| 218450_at | HEBP1 | 10.019 | 11.010 | 1.0 | 0.038 |
| 218046_s_at | MRPS16 | 9.182 | 10.170 | 1.0 | 0.029 |
| 205361_s_at | PFDN4 | 10.055 | 11.042 | 1.0 | 0.007 |
| 216593_s_at | PIGC | 7.042 | 8.029 | 1.0 | 0.013 |
| 217140_s_at | VDAC1 | 10.379 | 11.365 | 1.0 | 0.022 |
| 210216_x_at | RAD1 | 8.799 | 9.783 | 1.0 | 0.028 |
| 208899_x_at | ATP6V1D | 9.718 | 10.700 | 1.0 | 0.045 |
| 208969_at | NDUFA9 | 10.723 | 11.704 | 1.0 | 0.015 |
| 221036_s_at | APH1B | 5.827 | 6.807 | 1.0 | 0.008 |
| 219622_at | RAB20 | 6.747 | 7.728 | 1.0 | 0.017 |
| 220285_at | C9orf77 | 6.346 | 7.325 | 1.0 | 0.022 |
| 202903_at | LSM5 | 5.543 | 6.519 | 1.0 | 0.041 |
| 202457_s_at | PPP3CA | 9.332 | 10.304 | 0.9 | 0.028 |
| 218156_s_at | TSR1 | 9.071 | 10.043 | 0.9 | 0.024 |
| 212191_x_at | RPL13 | 12.672 | 13.644 | 0.9 | 0.031 |
| 203515_s_at | PMVK | 7.586 | 8.556 | 0.9 | 0.000 |
| 217061_s_at | ETV1 | 5.678 | 6.646 | 0.9 | 0.037 |
| 218817_at | SPCS3 | 7.070 | 8.035 | 0.9 | 0.007 |
| 206809_s_at | HNRPA3P1 /// HNRPA3 | 9.313 | 10.276 | 0.9 | 0.007 |
| 221620_s_at | MGC4825 | 8.404 | 9.367 | 0.9 | 0.040 |
| 221803_s_at | NRBF2 | 7.141 | 8.102 | 0.9 | 0.012 |
| 220789_s_at | TBRG4 | 8.834 | 9.794 | 0.9 | 0.048 |
| 207980_s_at | CITED2 | 7.354 | 8.313 | 0.9 | 0.018 |
| 220588_at | BCAS4 | 5.308 | 6.266 | 0.9 | 0.004 |
| 208734_x_at | RAB2 | 9.994 | 10.952 | 0.9 | 0.005 |
| 203328_x_at | IDE | 8.132 | 9.090 | 0.9 | 0.037 |
| 201214_s_at | PPP1R7 | 9.032 | 9.988 | 0.9 | 0.044 |
| 213788_s_at | FLJ35348 | 6.129 | 7.084 | 0.9 | 0.001 |
| 218220_at | C12orf10 | 7.977 | 8.932 | 0.9 | 0.045 |
| 202162_s_at | CNOT8 | 8.152 | 9.106 | 0.9 | 0.016 |

| | | | | | |
|---|---|---|---|---|---|
| 220631_at | OSGEPL1 | 7.468 | 8.420 | 0.9 | 0.005 |
| 208758_at | ATIC | 10.545 | 11.497 | 0.9 | 0.015 |
| 200005_at | EIF3S7 | 11.651 | 12.600 | 0.9 | 0.039 |
| 217741_s_at | ZA20D2 | 10.020 | 10.967 | 0.9 | 0.048 |
| 200715_x_at | RPL13A | 12.310 | 13.257 | 0.9 | 0.002 |
| 202690_s_at | SNRPD1 | 10.911 | 11.851 | 0.9 | 0.001 |
| 218738_s_at | RNF138 | 9.740 | 10.679 | 0.9 | 0.017 |
| 216457_s_at | SF3A1 | 9.162 | 10.101 | 0.9 | 0.000 |
| 218716_x_at | MTO1 | 7.807 | 8.745 | 0.9 | 0.038 |
| 214214_s_at | C1QBP | 11.126 | 12.064 | 0.9 | 0.011 |
| 217854_s_at | POLR2E | 10.178 | 11.114 | 0.9 | 0.037 |
| 202753_at | PSMD6 | 11.161 | 12.097 | 0.9 | 0.034 |
| 214437_s_at | SHMT2 | 8.488 | 9.423 | 0.9 | 0.000 |
| 203676_at | GNS | 6.661 | 7.596 | 0.9 | 0.018 |
| 218651_s_at | LARP6 | 6.845 | 7.779 | 0.9 | 0.037 |
| 209239_at | NFKB1 | 7.927 | 8.857 | 0.9 | 0.025 |
| 202461_at | EIF2B2 | 9.031 | 9.961 | 0.9 | 0.031 |
| 204729_s_at | STX1A | 5.671 | 6.599 | 0.9 | 0.015 |
| 201167_x_at | ARHGDIA | 5.334 | 6.262 | 0.9 | 0.020 |
| 213897_s_at | MRPL23 | 9.781 | 10.708 | 0.9 | 0.019 |
| 212385_at | --- | 5.590 | 6.517 | 0.9 | 0.034 |
| 206445_s_at | PRMT1 | 11.370 | 12.295 | 0.9 | 0.019 |
| 216532_x_at | --- | 6.777 | 7.702 | 0.9 | 0.009 |
| 201066_at | CYC1 | 10.761 | 11.686 | 0.9 | 0.021 |
| 212563_at | BOP1 /// LOC653119 | 9.192 | 10.116 | 0.9 | 0.010 |
| 212973_at | RPIA | 8.902 | 9.825 | 0.9 | 0.019 |
| 206670_s_at | GAD1 | 5.530 | 6.452 | 0.9 | 0.017 |
| 216088_s_at | PSMA7 | 9.314 | 10.237 | 0.9 | 0.001 |
| 215380_s_at | C7orf24 | 11.025 | 11.946 | 0.8 | 0.034 |
| 201784_s_at | C11orf58 | 9.585 | 10.505 | 0.8 | 0.020 |
| 216392_s_at | SEC23IP | 7.344 | 8.264 | 0.8 | 0.005 |
| 214668_at | C13orf1 | 5.005 | 5.925 | 0.8 | 0.017 |
| 76897_s_at | KIAA0674 | 5.172 | 6.091 | 0.8 | 0.026 |
| 218890_x_at | MRPL35 | 9.883 | 10.802 | 0.8 | 0.018 |
| 219193_at | WDR70 | 8.784 | 9.703 | 0.8 | 0.015 |
| 218513_at | FLJ11184 | 4.943 | 5.861 | 0.8 | 0.005 |
| 200038_s_at | RPL17 | 13.021 | 13.939 | 0.8 | 0.013 |
| 201695_s_at | NP | 9.934 | 10.851 | 0.8 | 0.031 |
| 212475_at | KIAA0241 | 5.694 | 6.611 | 0.8 | 0.013 |
| 210534_s_at | EPPB9 | 7.537 | 8.453 | 0.8 | 0.018 |
| 213687_s_at | RPL35A | 12.451 | 13.366 | 0.8 | 0.000 |
| 213287_s_at | KRT10 | 8.867 | 9.781 | 0.8 | 0.018 |
| 213223_at | RPL28 | 7.673 | 8.585 | 0.8 | 0.024 |
| 200889_s_at | SSR1 | 8.946 | 9.858 | 0.8 | 0.035 |
| 204559_s_at | LSM7 | 10.889 | 11.801 | 0.8 | 0.043 |

| | | | | | |
|---|---|---|---|---|---|
| 204209_at | PCYT1A | 6.134 | 7.046 | 0.8 | 0.020 |
| 218253_s_at | LGTN | 8.907 | 9.818 | 0.8 | 0.028 |
| 217955_at | BCL2L13 | 6.764 | 7.674 | 0.8 | 0.006 |
| 213907_at | EEF1E1 | 5.739 | 6.648 | 0.8 | 0.045 |
| 201300_s_at | PRNP | 10.620 | 11.529 | 0.8 | 0.039 |
| 218481_at | EXOSC5 | 7.960 | 8.869 | 0.8 | 0.026 |
| 204593_s_at | RP5-1104E15.5 | 8.110 | 9.015 | 0.8 | 0.002 |
| 219449_s_at | TMEM70 | 10.446 | 11.350 | 0.8 | 0.003 |
| 201138_s_at | SSB | 10.109 | 11.013 | 0.8 | 0.018 |
| 209549_s_at | DGUOK | 9.900 | 10.803 | 0.8 | 0.023 |
| 202472_at | MPI | 6.115 | 7.017 | 0.8 | 0.003 |
| 210188_at | GABPA /// GABPAP | 5.770 | 6.672 | 0.8 | 0.008 |
| 208796_s_at | CCNG1 | 11.546 | 12.447 | 0.8 | 0.023 |
| 203208_s_at | MTFR1 | 10.098 | 10.996 | 0.8 | 0.010 |
| 217851_s_at | C20orf45 | 8.800 | 9.698 | 0.8 | 0.012 |
| 222045_s_at | C20orf67 | 5.392 | 6.289 | 0.8 | 0.003 |
| 202858_at | U2AF1 | 11.348 | 12.243 | 0.8 | 0.039 |
| 203138_at | HAT1 | 10.736 | 11.628 | 0.8 | 0.002 |
| 210458_s_at | TANK | 5.256 | 6.148 | 0.8 | 0.003 |
| 219109_at | SPAG16 | 6.796 | 7.687 | 0.8 | 0.032 |
| 214579_at | NPAL3 | 6.090 | 6.981 | 0.8 | 0.010 |
| 216508_x_at | HMGB1 /// HMG1L1 /// LOC644380 | 9.005 | 9.894 | 0.8 | 0.005 |
| 203941_at | RC74 | 6.748 | 7.637 | 0.8 | 0.008 |
| 212476_at | CENTB2 | 8.832 | 9.721 | 0.8 | 0.037 |
| 208369_s_at | GCDH | 6.426 | 7.314 | 0.8 | 0.039 |
| 200777_s_at | BZW1 /// LOC151579 | 11.465 | 12.353 | 0.8 | 0.042 |
| 221987_s_at | TSR1 | 8.152 | 9.040 | 0.8 | 0.017 |
| 219767_s_at | CRYZL1 | 7.053 | 7.938 | 0.8 | 0.023 |
| 208101_s_at | C9orf74 | 7.001 | 7.886 | 0.8 | 0.027 |
| 217725_x_at | SERBP1 | 10.142 | 11.027 | 0.8 | 0.013 |
| 211255_x_at | DEDD | 5.374 | 6.258 | 0.8 | 0.024 |
| 204560_at | FKBP5 | 5.606 | 6.489 | 0.8 | 0.031 |
| 203401_at | PRPS2 | 8.997 | 9.880 | 0.8 | 0.029 |
| 203177_x_at | TFAM | 9.610 | 10.492 | 0.8 | 0.008 |
| 206236_at | GPR4 | 4.909 | 5.790 | 0.8 | 0.043 |
| 201435_s_at | EIF4E | 8.916 | 9.797 | 0.8 | 0.003 |
| 211622_s_at | ARF3 | 8.825 | 9.706 | 0.8 | 0.008 |
| 201179_s_at | GNAI3 | 8.709 | 9.590 | 0.8 | 0.004 |
| 222118_at | C16orf60 | 5.478 | 6.358 | 0.8 | 0.019 |
| 209565_at | RNF113A | 8.165 | 9.042 | 0.8 | 0.005 |
| 206668_s_at | SCAMP1 | 5.881 | 6.757 | 0.8 | 0.024 |
| 208103_s_at | ANP32E | 8.396 | 9.270 | 0.8 | 0.024 |
| 217737_x_at | C20orf43 | 8.362 | 9.236 | 0.8 | 0.011 |
| 211932_at | HNRPA3 | 9.299 | 10.173 | 0.8 | 0.028 |

| | | | | | |
|---|---|---|---|---|---|
| 205017_s_at | MBNL2 | 5.643 | 6.516 | 0.8 | 0.025 |
| 209048_s_at | PRKCBP1 | 9.951 | 10.823 | 0.8 | 0.046 |
| 208945_s_at | BECN1 | 7.932 | 8.804 | 0.8 | 0.050 |
| 204507_s_at | PPP3R1 | 6.253 | 7.125 | 0.8 | 0.003 |
| 203360_s_at | MYCBP | 8.823 | 9.694 | 0.8 | 0.036 |
| 209714_s_at | CDKN3 | 10.468 | 11.339 | 0.8 | 0.037 |
| 220688_s_at | C1orf33 | 9.786 | 10.656 | 0.8 | 0.046 |
| 200868_s_at | ZNF313 | 9.238 | 10.107 | 0.8 | 0.039 |
| 203163_at | KATNB1 | 6.145 | 7.009 | 0.7 | 0.023 |
| 203270_at | DTYMK /// LOC653208 | 9.154 | 10.018 | 0.7 | 0.037 |
| 207290_at | PLXNA2 | 5.012 | 5.875 | 0.7 | 0.042 |
| 214351_x_at | RPL13 /// LOC388344 | 12.069 | 12.931 | 0.7 | 0.038 |
| 203800_s_at | MRPS14 | 9.291 | 10.150 | 0.7 | 0.010 |
| 218286_s_at | RNF7 | 8.411 | 9.270 | 0.7 | 0.018 |
| 211824_x_at | NALP1 | 5.146 | 6.005 | 0.7 | 0.024 |
| 222159_at | PLXNA2 | 5.524 | 6.379 | 0.7 | 0.020 |
| 215113_s_at | SENP3 | 7.413 | 8.268 | 0.7 | 0.049 |
| 210338_s_at | HSPA8 | 12.399 | 13.252 | 0.7 | 0.005 |
| 219357_at | GTPBP1 | 7.688 | 8.541 | 0.7 | 0.021 |
| 205133_s_at | HSPE1 | 11.191 | 12.041 | 0.7 | 0.004 |
| 201226_at | NDUFB8 | 9.966 | 10.815 | 0.7 | 0.038 |
| 208781_x_at | SNX3 | 10.064 | 10.913 | 0.7 | 0.043 |
| 203363_s_at | KIAA0652 | 6.677 | 7.526 | 0.7 | 0.018 |
| 218049_s_at | MRPL13 | 9.600 | 10.448 | 0.7 | 0.031 |
| 40189_at | SET | 11.085 | 11.933 | 0.7 | 0.004 |
| 203141_s_at | AP3B1 | 6.704 | 7.549 | 0.7 | 0.035 |
| 207099_s_at | CHM | 5.141 | 5.986 | 0.7 | 0.045 |
| 201239_s_at | SPCS2 /// LOC653566 | 10.881 | 11.724 | 0.7 | 0.039 |
| 200892_s_at | SFRS10 | 10.010 | 10.853 | 0.7 | 0.025 |
| 221726_at | RPL22 | 10.429 | 11.271 | 0.7 | 0.036 |
| 213373_s_at | CASP8 | 8.775 | 9.617 | 0.7 | 0.002 |
| 217874_at | SUCLG1 | 10.915 | 11.757 | 0.7 | 0.019 |
| 205315_s_at | SNTB2 | 7.977 | 8.819 | 0.7 | 0.032 |
| 201902_s_at | YY1 | 5.994 | 6.834 | 0.7 | 0.017 |
| 201054_at | HNRPA0 | 10.484 | 11.323 | 0.7 | 0.018 |
| 209141_at | UBE2G1 | 9.589 | 10.428 | 0.7 | 0.039 |
| 200086_s_at | COX4I1 | 11.320 | 12.159 | 0.7 | 0.025 |
| 221381_s_at | MORF4L1 /// MORF4 | 10.225 | 11.063 | 0.7 | 0.021 |
| 210904_s_at | IL13RA1 | 6.829 | 7.667 | 0.7 | 0.033 |
| 210681_s_at | USP15 | 9.083 | 9.920 | 0.7 | 0.035 |
| 210691_s_at | CACYBP | 9.904 | 10.740 | 0.7 | 0.034 |
| 204300_at | PET112L | 7.094 | 7.929 | 0.7 | 0.016 |
| 200978_at | MDH1 | 11.522 | 12.357 | 0.7 | 0.000 |
| 213698_at | ZMYM6 | 7.998 | 8.831 | 0.7 | 0.045 |
| 200056_s_at | C1D | 9.012 | 9.845 | 0.7 | 0.033 |

| | | | | | |
|---|---|---|---|---|---|
| 201091_s_at | CBX3 /// LOC653972 | 10.273 | 11.106 | 0.7 | 0.046 |
| 218919_at | ZFAND1 | 9.210 | 10.040 | 0.7 | 0.027 |
| 217919_s_at | MRPL42 | 10.200 | 11.030 | 0.7 | 0.022 |
| 200812_at | CCT7 | 11.309 | 12.137 | 0.7 | 0.009 |
| 201988_s_at | CREBL2 | 6.700 | 7.528 | 0.7 | 0.027 |
| 203103_s_at | PRPF19 | 9.269 | 10.095 | 0.7 | 0.014 |
| 210131_x_at | SDHC | 10.046 | 10.871 | 0.7 | 0.013 |
| 210153_s_at | ME2 | 8.542 | 9.367 | 0.7 | 0.034 |
| 207040_s_at | ST13 | 10.750 | 11.575 | 0.7 | 0.042 |
| 201459_at | RUVBL2 | 11.011 | 11.833 | 0.7 | 0.002 |
| 212145_at | MRPS27 | 9.415 | 10.237 | 0.7 | 0.020 |
| 214167_s_at | RPLP0 /// RPLP0-like | 12.242 | 13.063 | 0.7 | 0.017 |
| 200679_x_at | HMGB1 | 12.043 | 12.864 | 0.7 | 0.011 |
| 202681_at | USP4 | 7.573 | 8.394 | 0.7 | 0.047 |
| 200886_s_at | PGAM1 /// LOC642969 /// LOC643576 | 12.275 | 13.096 | 0.7 | 0.032 |
| 205398_s_at | SMAD3 | 7.645 | 8.465 | 0.7 | 0.048 |
| 220258_s_at | WDR79 | 6.930 | 7.750 | 0.7 | 0.004 |
| 207791_s_at | RAB1A | 10.095 | 10.915 | 0.7 | 0.047 |
| 200634_at | PFN1 | 12.683 | 13.502 | 0.7 | 0.015 |
| 201573_s_at | ETF1 | 9.292 | 10.108 | 0.7 | 0.041 |
| 200997_at | RBM4 /// LOC650029 | 9.277 | 10.092 | 0.7 | 0.010 |
| 207629_s_at | ARHGEF2 | 5.905 | 6.720 | 0.7 | 0.004 |
| 59705_at | SCLY | 5.471 | 6.286 | 0.7 | 0.022 |
| 204336_s_at | RGS19 | 7.797 | 8.611 | 0.7 | 0.015 |
| 203137_at | WTAP | 9.410 | 10.224 | 0.7 | 0.021 |
| 203613_s_at | NDUFB6 | 9.770 | 10.583 | 0.7 | 0.019 |
| 208693_s_at | GARS | 11.698 | 12.510 | 0.7 | 0.035 |
| 211797_s_at | NFYC | 7.029 | 7.841 | 0.7 | 0.019 |
| 214590_s_at | UBE2D1 | 5.752 | 6.563 | 0.7 | 0.000 |
| 202810_at | DRG1 | 9.996 | 10.807 | 0.7 | 0.015 |
| 208726_s_at | EIF2S2 | 12.139 | 12.948 | 0.7 | 0.000 |
| 209649_at | STAM2 | 7.376 | 8.185 | 0.7 | 0.027 |
| 203214_x_at | CDC2 | 10.088 | 10.896 | 0.7 | 0.032 |
| 211547_s_at | PAFAH1B1 | 6.608 | 7.415 | 0.7 | 0.033 |
| 209615_s_at | PAK1 | 5.476 | 6.282 | 0.6 | 0.027 |
| 217726_at | COPZ1 | 9.074 | 9.880 | 0.6 | 0.027 |
| 205394_at | CHEK1 | 8.515 | 9.318 | 0.6 | 0.021 |
| 216100_s_at | TOR1AIP1 | 5.295 | 6.099 | 0.6 | 0.026 |
| 212217_at | PREPL | 8.517 | 9.319 | 0.6 | 0.024 |
| 218535_s_at | RIOK2 | 8.366 | 9.168 | 0.6 | 0.019 |
| 202467_s_at | COPS2 | 10.136 | 10.938 | 0.6 | 0.027 |
| 207319_s_at | CDC2L5 | 5.251 | 6.052 | 0.6 | 0.021 |
| 204335_at | CCDC94 | 5.888 | 6.688 | 0.6 | 0.006 |
| 218982_s_at | MRPS17 | 9.440 | 10.239 | 0.6 | 0.021 |

| | | | | | |
|---|---|---|---|---|---|
| 200922_at | KDELR1 | 8.051 | 8.848 | 0.6 | 0.030 |
| 219769_at | INCENP | 5.823 | 6.620 | 0.6 | 0.017 |
| 219487_at | BBS10 | 6.220 | 7.016 | 0.6 | 0.011 |
| 209096_at | UBE2V2 | 10.558 | 11.353 | 0.6 | 0.044 |
| 207023_x_at | KRT10 | 9.432 | 10.227 | 0.6 | 0.026 |
| 208909_at | UQCRFS1 | 11.499 | 12.294 | 0.6 | 0.032 |
| 210759_s_at | PSMA1 | 11.238 | 12.030 | 0.6 | 0.050 |
| 208630_at | HADHA | 9.817 | 10.608 | 0.6 | 0.016 |
| 202910_s_at | CD97 | 8.185 | 8.976 | 0.6 | 0.016 |
| 207812_s_at | GORASP2 | 9.723 | 10.514 | 0.6 | 0.025 |
| 202004_x_at | SDHC /// LOC642502 | 9.646 | 10.436 | 0.6 | 0.018 |
| 212724_at | RND3 | 10.143 | 10.933 | 0.6 | 0.036 |
| 212333_at | FAM98A | 10.202 | 10.989 | 0.6 | 0.001 |
| 208878_s_at | PAK2 | 8.016 | 8.802 | 0.6 | 0.022 |
| 216295_s_at | CLTA | 10.948 | 11.731 | 0.6 | 0.047 |
| 203538_at | CAMLG | 9.581 | 10.365 | 0.6 | 0.033 |
| 218150_at | ARL5A | 10.060 | 10.842 | 0.6 | 0.029 |
| 218616_at | INTS12 | 8.468 | 9.251 | 0.6 | 0.037 |
| 210932_s_at | RNF6 | 4.981 | 5.763 | 0.6 | 0.003 |
| 209316_s_at | HBS1L | 9.474 | 10.256 | 0.6 | 0.046 |
| 218512_at | WDR12 | 10.258 | 11.040 | 0.6 | 0.014 |
| 214378_at | TFPI | 5.188 | 5.969 | 0.6 | 0.000 |
| 217448_s_at | C14orf92 | 6.636 | 7.416 | 0.6 | 0.041 |
| 203142_s_at | AP3B1 | 8.602 | 9.382 | 0.6 | 0.027 |
| 202698_x_at | COX4I1 | 12.403 | 13.183 | 0.6 | 0.028 |
| 212417_at | SCAMP1 | 7.707 | 8.487 | 0.6 | 0.015 |
| 211077_s_at | TLK1 | 5.174 | 5.951 | 0.6 | 0.013 |
| 201263_at | TARS | 11.364 | 12.141 | 0.6 | 0.020 |
| 211774_s_at | MMACHC | 5.771 | 6.547 | 0.6 | 0.001 |
| 218585_s_at | DTL | 9.370 | 10.145 | 0.6 | 0.021 |
| 201357_s_at | SF3A1 | 7.551 | 8.325 | 0.6 | 0.002 |
| 212018_s_at | RSL1D1 | 10.488 | 11.261 | 0.6 | 0.010 |
| 200631_s_at | SET | 11.907 | 12.679 | 0.6 | 0.010 |
| 218238_at | GTPBP4 | 9.541 | 10.313 | 0.6 | 0.010 |
| 201433_s_at | PTDSS1 | 11.161 | 11.932 | 0.6 | 0.017 |
| 218408_at | TIMM10 | 9.510 | 10.281 | 0.6 | 0.023 |
| 218207_s_at | STMN3 | 4.853 | 5.623 | 0.6 | 0.001 |
| 216383_at | LOC347544 | 6.167 | 6.936 | 0.6 | 0.025 |
| 200828_s_at | ZNF207 | 11.033 | 11.800 | 0.6 | 0.026 |
| 205628_at | PRIM2A | 6.123 | 6.890 | 0.6 | 0.040 |
| 205188_s_at | SMAD5 | 7.305 | 8.070 | 0.6 | 0.039 |
| 215044_s_at | STAM2 | 7.003 | 7.768 | 0.6 | 0.032 |
| 208679_s_at | ARPC2 | 12.121 | 12.885 | 0.6 | 0.023 |
| 215424_s_at | SNW1 | 9.652 | 10.416 | 0.6 | 0.045 |
| 203213_at | CDC2 | 11.227 | 11.990 | 0.6 | 0.033 |

| | | | | | |
|---|---|---|---|---|---|
| 211933_s_at | HNRPA3P1 /// HNRPA3 | 10.740 | 11.502 | 0.6 | 0.009 |
| 200669_s_at | UBE2D3 | 10.090 | 10.853 | 0.6 | 0.005 |
| 202706_s_at | UMPS | 8.796 | 9.559 | 0.6 | 0.025 |
| 216348_at | RPS17 /// LOC402057 | 8.497 | 9.259 | 0.6 | 0.025 |
| 221025_x_at | PUS7L | 6.721 | 7.482 | 0.6 | 0.009 |
| 205580_s_at | HRH1 | 5.537 | 6.298 | 0.6 | 0.031 |
| 208546_x_at | HIST1H2BH | 6.714 | 7.474 | 0.6 | 0.045 |
| 207350_s_at | VAMP4 | 5.399 | 6.156 | 0.6 | 0.011 |
| 202703_at | DUSP11 | 8.751 | 9.508 | 0.6 | 0.025 |
| 209044_x_at | SF3B4 | 9.286 | 10.042 | 0.6 | 0.018 |
| 205393_s_at | CHEK1 | 7.440 | 8.195 | 0.6 | 0.028 |
| 210180_s_at | SFRS10 | 8.604 | 9.358 | 0.6 | 0.022 |
| 218832_x_at | ARRB1 | 5.097 | 5.850 | 0.6 | 0.007 |
| 219408_at | PRMT7 | 6.060 | 6.813 | 0.6 | 0.045 |
| 219931_s_at | KLHL12 | 6.147 | 6.899 | 0.6 | 0.015 |
| 217772_s_at | MTCH2 | 11.093 | 11.845 | 0.6 | 0.017 |
| 208748_s_at | FLOT1 | 5.824 | 6.575 | 0.6 | 0.039 |
| 214729_at | TWISTNB | 4.846 | 5.595 | 0.6 | 0.009 |
| 202060_at | CTR9 | 9.158 | 9.906 | 0.6 | 0.040 |
| 219498_s_at | BCL11A | 5.299 | 6.045 | 0.6 | 0.021 |
| 218489_s_at | ALAD | 6.044 | 6.790 | 0.6 | 0.004 |
| 204226_at | STAU2 | 7.789 | 8.534 | 0.6 | 0.005 |
| 200809_x_at | RPL12 | 13.571 | 14.315 | 0.6 | 0.023 |
| 221158_at | C21orf66 | 7.285 | 8.028 | 0.6 | 0.047 |
| 200093_s_at | HINT1 | 12.008 | 12.749 | 0.5 | 0.015 |
| 202824_s_at | TCEB1 | 11.700 | 12.441 | 0.5 | 0.034 |
| 214095_at | SHMT2 | 8.031 | 8.771 | 0.5 | 0.045 |
| 201863_at | FAM32A | 9.329 | 10.065 | 0.5 | 0.017 |
| 220397_at | MDM1 | 5.241 | 5.976 | 0.5 | 0.035 |
| 209511_at | POLR2F | 9.247 | 9.982 | 0.5 | 0.029 |
| 209974_s_at | BUB3 | 11.209 | 11.943 | 0.5 | 0.043 |
| 218658_s_at | ACTR8 | 6.325 | 7.059 | 0.5 | 0.012 |
| 207722_s_at | BTBD2 | 5.819 | 6.552 | 0.5 | 0.017 |
| 218823_s_at | KCTD9 | 7.759 | 8.492 | 0.5 | 0.013 |
| 217909_s_at | MLX | 7.855 | 8.588 | 0.5 | 0.002 |
| 218564_at | RFWD3 | 7.837 | 8.570 | 0.5 | 0.021 |
| 201379_s_at | TPD52L2 | 9.865 | 10.597 | 0.5 | 0.019 |
| 200744_s_at | GNB1 | 10.070 | 10.803 | 0.5 | 0.042 |
| 217747_s_at | RPS9 | 12.864 | 13.596 | 0.5 | 0.001 |
| 204742_s_at | APRIN | 6.511 | 7.241 | 0.5 | 0.025 |
| 214008_at | PTK9 | 5.969 | 6.699 | 0.5 | 0.009 |
| 220086_at | ZNFN1A5 | 6.600 | 7.329 | 0.5 | 0.050 |
| 212537_x_at | RPL17 | 13.597 | 14.326 | 0.5 | 0.008 |
| 216962_at | RPAIN | 5.702 | 6.430 | 0.5 | 0.004 |
| 209330_s_at | HNRPD | 10.144 | 10.872 | 0.5 | 0.019 |

| | | | | | |
|---|---|---|---|---|---|
| 208541_x_at | TFAM | 7.087 | 7.814 | 0.5 | 0.017 |
| 218774_at | DCPS | 8.330 | 9.057 | 0.5 | 0.016 |
| 214040_s_at | GSN | 5.674 | 6.401 | 0.5 | 0.018 |
| 221604_s_at | PEX16 | 6.627 | 7.353 | 0.5 | 0.001 |
| 211251_x_at | NFYC | 7.460 | 8.185 | 0.5 | 0.036 |
| 221749_at | YTHDF3 | 9.903 | 10.628 | 0.5 | 0.039 |
| 215357_s_at | POLDIP3 | 6.214 | 6.938 | 0.5 | 0.001 |
| 200947_s_at | GLUD1 | 10.273 | 10.997 | 0.5 | 0.004 |
| 207618_s_at | BCS1L | 8.373 | 9.097 | 0.5 | 0.024 |
| 209786_at | HMGN4 | 8.250 | 8.973 | 0.5 | 0.045 |
| 207180_s_at | HTATIP2 | 8.253 | 8.975 | 0.5 | 0.047 |
| 200880_at | DNAJA1 | 10.373 | 11.094 | 0.5 | 0.041 |
| 209690_s_at | DOK4 | 5.409 | 6.130 | 0.5 | 0.025 |
| 218862_at | ASB13 | 7.051 | 7.771 | 0.5 | 0.032 |
| 203764_at | DLG7 | 10.344 | 11.064 | 0.5 | 0.010 |
| 218089_at | C20orf4 | 7.880 | 8.597 | 0.5 | 0.023 |
| 219539_at | GEMIN6 | 8.642 | 9.359 | 0.5 | 0.000 |
| 218057_x_at | COX4NB | 9.408 | 10.125 | 0.5 | 0.034 |
| 212270_x_at | RPL17 | 13.627 | 14.343 | 0.5 | 0.005 |
| 209304_x_at | GADD45B | 7.448 | 8.165 | 0.5 | 0.026 |
| 208047_s_at | NAB1 | 5.962 | 6.675 | 0.5 | 0.034 |
| 219133_at | OXSM | 8.339 | 9.051 | 0.5 | 0.013 |
| 201953_at | CIB1 | 9.452 | 10.163 | 0.5 | 0.023 |
| 219861_at | DNAJC17 | 6.381 | 7.091 | 0.5 | 0.010 |
| 203741_s_at | ADCY7 | 7.515 | 8.225 | 0.5 | 0.049 |
| 201897_s_at | CKS1B | 11.215 | 11.923 | 0.5 | 0.002 |
| 200833_s_at | RAP1B /// LOC643752 | 11.270 | 11.974 | 0.5 | 0.018 |
| 209584_x_at | APOBEC3C | 7.277 | 7.982 | 0.5 | 0.032 |
| 201877_s_at | PPP2R5C | 8.803 | 9.508 | 0.5 | 0.044 |
| 210470_x_at | NONO | 10.817 | 11.521 | 0.5 | 0.040 |
| 218485_s_at | SLC35C1 | 6.134 | 6.837 | 0.5 | 0.000 |
| 206220_s_at | RASA3 | 6.151 | 6.852 | 0.5 | 0.042 |
| 203664_s_at | POLR2D | 8.425 | 9.125 | 0.5 | 0.042 |
| 208719_s_at | DDX17 | 6.177 | 6.877 | 0.5 | 0.039 |
| 213321_at | BCKDHB | 5.690 | 6.391 | 0.5 | 0.036 |
| 218569_s_at | KBTBD4 | 6.628 | 7.328 | 0.5 | 0.008 |
| 212499_s_at | C14orf111 /// C14orf32 | 8.026 | 8.723 | 0.5 | 0.034 |
| 215948_x_at | ZMYM5 | 6.905 | 7.602 | 0.5 | 0.018 |
| 209448_at | HTATIP2 | 9.511 | 10.207 | 0.5 | 0.014 |
| 200767_s_at | C9orf10 | 7.896 | 8.590 | 0.5 | 0.007 |
| 219262_at | SUV39H2 | 4.713 | 5.406 | 0.5 | 0.032 |
| 201707_at | PEX19 | 7.568 | 8.259 | 0.5 | 0.012 |
| 205708_s_at | TRPM2 | 6.509 | 7.199 | 0.5 | 0.029 |
| 212055_at | C18orf10 | 9.255 | 9.945 | 0.5 | 0.043 |
| 202214_s_at | CUL4B | 9.345 | 10.034 | 0.5 | 0.031 |

| | | | | | |
|---|---|---|---|---|---|
| 207721_x_at | HINT1 | 12.329 | 13.016 | 0.5 | 0.045 |
| 217595_at | --- | 4.929 | 5.614 | 0.5 | 0.012 |
| 206141_at | MOCS3 | 5.695 | 6.381 | 0.5 | 0.033 |
| 201503_at | G3BP | 10.917 | 11.601 | 0.5 | 0.028 |
| 204459_at | CSTF2 | 8.401 | 9.084 | 0.5 | 0.045 |
| 212002_at | --- | 7.187 | 7.870 | 0.5 | 0.038 |
| 213606_s_at | ARHGDIA | 5.263 | 5.945 | 0.5 | 0.009 |
| 212600_s_at | UQCRC2 | 11.251 | 11.933 | 0.5 | 0.016 |
| 219104_at | RNF141 | 6.345 | 7.027 | 0.5 | 0.004 |
| 220329_s_at | C6orf96 | 7.474 | 8.153 | 0.5 | 0.030 |
| 211406_at | IER3IP1 | 6.502 | 7.179 | 0.5 | 0.030 |
| 201676_x_at | PSMA1 | 11.835 | 12.512 | 0.5 | 0.040 |
| 221094_s_at | ELP3 | 6.909 | 7.586 | 0.5 | 0.023 |
| 212497_at | C14orf32 | 5.973 | 6.649 | 0.5 | 0.011 |
| 211937_at | EIF4B | 10.680 | 11.355 | 0.5 | 0.003 |
| 200840_at | KARS | 12.448 | 13.118 | 0.5 | 0.037 |
| 219974_x_at | ECHDC1 | 9.256 | 9.926 | 0.4 | 0.008 |
| 202026_at | SDHD | 9.614 | 10.283 | 0.4 | 0.032 |
| 213861_s_at | FAM119B | 5.769 | 6.438 | 0.4 | 0.029 |
| 222229_x_at | RPL26 /// LOC400055 | 11.644 | 12.313 | 0.4 | 0.015 |
| 87100_at | ABHD2 | 5.144 | 5.812 | 0.4 | 0.029 |
| 207071_s_at | ACO1 | 8.388 | 9.055 | 0.4 | 0.034 |
| 215165_x_at | UMPS | 8.372 | 9.038 | 0.4 | 0.027 |
| 218405_at | ABT1 | 6.844 | 7.510 | 0.4 | 0.047 |
| 209958_s_at | PTHB1 | 5.693 | 6.360 | 0.4 | 0.002 |
| 221291_at | ULBP2 | 6.164 | 6.828 | 0.4 | 0.017 |
| 212937_s_at | COL6A1 | 5.668 | 6.332 | 0.4 | 0.008 |
| 204992_s_at | PFN2 | 11.697 | 12.360 | 0.4 | 0.019 |
| 211505_s_at | STAU1 | 9.988 | 10.650 | 0.4 | 0.019 |
| 218624_s_at | MGC2752 | 7.619 | 8.280 | 0.4 | 0.034 |
| 203013_at | ECD | 9.273 | 9.933 | 0.4 | 0.034 |
| 208696_at | CCT5 | 12.131 | 12.792 | 0.4 | 0.007 |
| 208743_s_at | YWHAB | 10.903 | 11.564 | 0.4 | 0.043 |
| 201532_at | PSMA3 | 11.661 | 12.322 | 0.4 | 0.026 |
| 209134_s_at | RPS6 | 12.926 | 13.586 | 0.4 | 0.034 |
| 200705_s_at | EEF1B2 | 12.524 | 13.184 | 0.4 | 0.004 |
| 212962_at | SYDE1 | 5.597 | 6.256 | 0.4 | 0.026 |
| 204641_at | NEK2 | 8.741 | 9.398 | 0.4 | 0.017 |
| 209249_s_at | GHITM | 11.414 | 12.071 | 0.4 | 0.028 |
| 218230_at | ARFIP1 | 8.732 | 9.386 | 0.4 | 0.021 |
| 211098_x_at | TMCO1 | 8.568 | 9.220 | 0.4 | 0.003 |
| 201256_at | COX7A2L | 9.819 | 10.470 | 0.4 | 0.045 |
| 221871_s_at | TFG | 6.013 | 6.664 | 0.4 | 0.012 |
| 213048_s_at | --- | 12.151 | 12.802 | 0.4 | 0.018 |
| 217852_s_at | ARL8B | 10.287 | 10.937 | 0.4 | 0.009 |

| | | | | | |
|---|---|---|---|---|---|
| 217839_at | TFG | 10.465 | 11.114 | 0.4 | 0.037 |
| 208643_s_at | XRCC5 | 10.667 | 11.317 | 0.4 | 0.044 |
| 201593_s_at | LEREPO4 | 10.966 | 11.614 | 0.4 | 0.023 |
| 200088_x_at | RPL12 | 13.729 | 14.375 | 0.4 | 0.029 |
| 208640_at | RAC1 | 12.404 | 13.049 | 0.4 | 0.036 |
| 218850_s_at | LIMD1 | 5.779 | 6.424 | 0.4 | 0.008 |
| 218420_s_at | C13orf23 | 7.540 | 8.183 | 0.4 | 0.038 |
| 209139_s_at | PRKRA | 9.249 | 9.892 | 0.4 | 0.024 |
| 211849_s_at | RNGTT | 5.119 | 5.760 | 0.4 | 0.013 |
| 210633_x_at | KRT10 | 9.475 | 10.116 | 0.4 | 0.044 |
| 215891_s_at | GM2A | 5.112 | 5.752 | 0.4 | 0.012 |
| 212004_at | C1orf144 | 7.313 | 7.953 | 0.4 | 0.044 |
| 200949_x_at | RPS20 | 13.453 | 14.091 | 0.4 | 0.041 |
| 201726_at | ELAVL1 | 10.818 | 11.456 | 0.4 | 0.014 |
| 211297_s_at | CDK7 | 9.535 | 10.172 | 0.4 | 0.028 |
| 209151_x_at | TCF3 | 4.766 | 5.403 | 0.4 | 0.044 |
| 200877_at | CCT4 | 12.497 | 13.134 | 0.4 | 0.036 |
| 206584_at | LY96 | 4.746 | 5.383 | 0.4 | 0.018 |
| 214738_s_at | NEK9 | 5.553 | 6.189 | 0.4 | 0.014 |
| 218453_s_at | C6orf35 /// LOC653041 | 5.425 | 6.061 | 0.4 | 0.031 |
| 203437_at | TMEM11 | 8.213 | 8.848 | 0.4 | 0.013 |
| 213756_s_at | HSF1 | 4.886 | 5.521 | 0.4 | 0.001 |
| 212824_at | FUBP3 | 8.472 | 9.106 | 0.4 | 0.017 |
| 202031_s_at | WIPI2 | 9.123 | 9.756 | 0.4 | 0.034 |
| 203403_s_at | RNF6 | 9.614 | 10.244 | 0.4 | 0.047 |
| 218388_at | PGLS | 9.053 | 9.683 | 0.4 | 0.042 |
| 200881_s_at | DNAJA1 | 11.643 | 12.271 | 0.4 | 0.013 |
| 209773_s_at | RRM2 | 11.946 | 12.574 | 0.4 | 0.001 |
| 208842_s_at | GORASP2 | 9.586 | 10.214 | 0.4 | 0.012 |
| 216283_s_at | PVR | 5.057 | 5.684 | 0.4 | 0.007 |
| 218170_at | ISOC1 | 8.607 | 9.233 | 0.4 | 0.032 |
| 214960_at | API5 | 6.592 | 7.219 | 0.4 | 0.048 |
| 201198_s_at | PSMD1 | 9.731 | 10.355 | 0.4 | 0.016 |
| 37232_at | KIAA0586 | 6.009 | 6.633 | 0.4 | 0.029 |
| 218361_at | GOLPH3L | 8.197 | 8.818 | 0.4 | 0.041 |
| 212257_s_at | SMARCA2 | 5.556 | 6.176 | 0.4 | 0.017 |
| 202232_s_at | PCID1 | 11.159 | 11.779 | 0.4 | 0.040 |
| 200036_s_at | RPL10A | 12.993 | 13.611 | 0.4 | 0.007 |
| 210349_at | CAMK4 | 5.362 | 5.978 | 0.4 | 0.030 |
| 222335_at | --- | 5.952 | 6.566 | 0.4 | 0.023 |
| 219004_s_at | C21orf45 | 8.706 | 9.319 | 0.4 | 0.039 |
| 200813_s_at | PAFAH1B1 | 8.585 | 9.196 | 0.4 | 0.035 |
| 216247_at | LOC440992 | 4.863 | 5.473 | 0.4 | 0.008 |
| 209927_s_at | C1orf77 | 6.423 | 7.030 | 0.4 | 0.017 |
| 209727_at | GM2A | 5.604 | 6.211 | 0.4 | 0.013 |

| | | | | | |
|---|---|---|---|---|---|
| 208752_x_at | NAP1L1 | 12.014 | 12.620 | 0.4 | 0.013 |
| 201937_s_at | DNPEP | 7.915 | 8.521 | 0.4 | 0.011 |
| 205015_s_at | TGFA | 5.216 | 5.821 | 0.4 | 0.039 |
| 203261_at | DCTN6 | 9.754 | 10.356 | 0.4 | 0.010 |
| 202126_at | PRPF4B | 9.509 | 10.110 | 0.4 | 0.043 |
| 202634_at | POLR2K | 8.528 | 9.128 | 0.4 | 0.012 |
| 211938_at | EIF4B | 11.402 | 12.002 | 0.4 | 0.025 |
| 219810_at | VCPIP1 | 5.695 | 6.294 | 0.4 | 0.030 |
| 203272_s_at | TUSC2 | 7.881 | 8.480 | 0.4 | 0.020 |
| 213688_at | CALM1 | 5.659 | 6.258 | 0.4 | 0.019 |
| 201558_at | RAE1 | 10.815 | 11.412 | 0.4 | 0.018 |
| 202972_s_at | FAM13A1 | 5.805 | 6.402 | 0.4 | 0.043 |
| 210378_s_at | SSNA1 | 7.891 | 8.488 | 0.4 | 0.034 |
| 212711_at | CAMSAP1 | 8.335 | 8.931 | 0.4 | 0.014 |
| 37226_at | BNIP1 | 5.825 | 6.419 | 0.4 | 0.031 |
| 215185_at | --- | 5.166 | 5.756 | 0.3 | 0.004 |
| 207396_s_at | ALG3 | 8.511 | 9.102 | 0.3 | 0.048 |
| 201799_s_at | OSBP | 7.486 | 8.075 | 0.3 | 0.046 |
| 206917_at | GNA13 | 5.319 | 5.908 | 0.3 | 0.049 |
| 221279_at | GDAP1 | 5.523 | 6.112 | 0.3 | 0.011 |
| 200658_s_at | PHB | 10.389 | 10.977 | 0.3 | 0.030 |
| 206942_s_at | PMCH | 4.799 | 5.387 | 0.3 | 0.036 |
| 218118_s_at | TIMM23 | 9.811 | 10.399 | 0.3 | 0.003 |
| 208826_x_at | HINT1 | 12.423 | 13.010 | 0.3 | 0.046 |
| 217859_s_at | SLC39A9 | 5.867 | 6.453 | 0.3 | 0.015 |
| 217309_s_at | DSCR3 | 5.963 | 6.548 | 0.3 | 0.011 |
| 65591_at | WDR48 | 6.700 | 7.285 | 0.3 | 0.017 |
| 216855_s_at | HNRPU | 5.586 | 6.170 | 0.3 | 0.019 |
| 202829_s_at | SYBL1 | 9.786 | 10.370 | 0.3 | 0.016 |
| 214271_x_at | RPL12 | 13.652 | 14.235 | 0.3 | 0.023 |
| 201241_at | DDX1 | 11.615 | 12.197 | 0.3 | 0.003 |
| 200873_s_at | CCT8 | 12.223 | 12.802 | 0.3 | 0.043 |
| 201218_at | CTBP2 /// LOC645291 /// LOC645508 /// LOC650999 | 10.257 | 10.835 | 0.3 | 0.020 |
| 203458_at | SPR | 8.368 | 8.944 | 0.3 | 0.028 |
| 210658_s_at | GGA2 | 6.564 | 7.139 | 0.3 | 0.035 |
| 201805_at | PRKAG1 | 8.856 | 9.428 | 0.3 | 0.031 |
| 204033_at | TRIP13 | 10.181 | 10.753 | 0.3 | 0.020 |
| 200793_s_at | ACO2 | 10.060 | 10.630 | 0.3 | 0.012 |
| 211499_s_at | MAPK11 | 5.752 | 6.321 | 0.3 | 0.016 |
| 201323_at | EBNA1BP2 | 10.035 | 10.601 | 0.3 | 0.041 |
| 64432_at | C12orf47 | 7.539 | 8.106 | 0.3 | 0.009 |
| 212113_at | LOC552889 | 5.786 | 6.352 | 0.3 | 0.010 |
| 209868_s_at | RBMS1 /// LOC648293 | 10.111 | 10.676 | 0.3 | 0.026 |

| | | | | | |
|---|---|---|---|---|---|
| 219904_at | ZSCAN5 | 5.652 | 6.216 | 0.3 | 0.045 |
| 201595_s_at | LEREPO4 | 10.534 | 11.097 | 0.3 | 0.023 |
| 202352_s_at | PSMD12 | 10.835 | 11.395 | 0.3 | 0.022 |
| 213762_x_at | RBMX | 11.000 | 11.559 | 0.3 | 0.009 |
| 205362_s_at | PFDN4 | 6.239 | 6.797 | 0.3 | 0.025 |
| 200084_at | C11orf58 | 11.040 | 11.597 | 0.3 | 0.047 |
| 213468_at | ERCC2 | 7.227 | 7.782 | 0.3 | 0.018 |
| 200937_s_at | RPL5 | 12.664 | 13.219 | 0.3 | 0.020 |
| 221550_at | COX15 | 7.062 | 7.616 | 0.3 | 0.025 |
| 210466_s_at | SERBP1 | 12.390 | 12.943 | 0.3 | 0.017 |
| 200013_at | RPL24 | 13.098 | 13.650 | 0.3 | 0.007 |
| 200014_s_at | HNRPC | 11.239 | 11.791 | 0.3 | 0.026 |
| 215230_x_at | EIF3S8 /// LOC653352 | 11.949 | 12.500 | 0.3 | 0.018 |
| 213851_at | TMEM110 | 5.959 | 6.509 | 0.3 | 0.017 |
| 221679_s_at | ABHD6 | 5.386 | 5.935 | 0.3 | 0.019 |
| 214849_at | C6orf69 | 5.400 | 5.945 | 0.3 | 0.047 |
| 218949_s_at | QRSL1 | 7.905 | 8.449 | 0.3 | 0.050 |
| 208778_s_at | TCP1 | 11.714 | 12.257 | 0.3 | 0.020 |
| 209476_at | TXNDC | 10.746 | 11.289 | 0.3 | 0.029 |
| 205264_at | CD3EAP | 5.940 | 6.479 | 0.3 | 0.026 |
| 213696_s_at | MED8 | 8.074 | 8.613 | 0.3 | 0.028 |
| 220953_s_at | MTMR12 | 5.935 | 6.473 | 0.3 | 0.012 |
| 53202_at | C7orf25 | 5.117 | 5.655 | 0.3 | 0.025 |
| 220408_x_at | FAM48A | 8.786 | 9.323 | 0.3 | 0.020 |
| 204716_at | CCDC6 | 7.127 | 7.661 | 0.3 | 0.041 |
| 215481_s_at | PEX5 | 5.244 | 5.778 | 0.3 | 0.012 |
| 202266_at | TTRAP | 9.788 | 10.320 | 0.3 | 0.006 |
| 218202_x_at | MRPL44 | 5.183 | 5.714 | 0.3 | 0.002 |
| 201653_at | CNIH | 10.586 | 11.115 | 0.3 | 0.023 |
| 203068_at | KLHL21 | 6.699 | 7.226 | 0.3 | 0.025 |
| 213253_at | SMC2L1 | 7.670 | 8.197 | 0.3 | 0.023 |
| 212264_s_at | WAPAL | 8.983 | 9.509 | 0.3 | 0.021 |
| 212967_x_at | NAP1L1 | 11.760 | 12.283 | 0.3 | 0.016 |
| 217822_at | WBP11 | 10.001 | 10.524 | 0.3 | 0.017 |
| 210076_x_at | SERBP1 | 7.603 | 8.124 | 0.3 | 0.028 |
| 206351_s_at | PEX10 | 5.292 | 5.814 | 0.3 | 0.003 |
| 213521_at | PTPN18 | 5.677 | 6.197 | 0.3 | 0.011 |
| 202589_at | TYMS | 12.174 | 12.694 | 0.3 | 0.048 |
| 221910_at | --- | 4.882 | 5.402 | 0.3 | 0.001 |
| 211942_x_at | RPL13A /// LOC283340 /// RP11-365K22.1 /// LOC642209 /// LOC644511 | 13.316 | 13.835 | 0.3 | 0.042 |
| 209313_at | XAB1 | 9.720 | 10.238 | 0.3 | 0.026 |
| 218438_s_at | MED28 | 9.317 | 9.835 | 0.3 | 0.037 |

| | | | | | |
|---|---|---|---|---|---|
| 207219_at | ZNF643 | 5.039 | 5.552 | 0.3 | 0.012 |
| 219361_s_at | ISG20L1 | 7.443 | 7.956 | 0.3 | 0.041 |
| 208540_x_at | S100A11 | 9.835 | 10.348 | 0.3 | 0.027 |
| 205920_at | SLC6A6 | 5.270 | 5.783 | 0.3 | 0.033 |
| 208194_s_at | STAM2 | 5.456 | 5.967 | 0.3 | 0.005 |
| 210818_s_at | BACH1 | 5.490 | 6.001 | 0.3 | 0.005 |
| 201577_at | NME1 | 12.620 | 13.131 | 0.3 | 0.025 |
| 209818_s_at | HABP4 | 5.537 | 6.046 | 0.3 | 0.048 |
| 207304_at | ZNF45 | 5.568 | 6.076 | 0.3 | 0.012 |
| 214698_at | ROD1 | 5.903 | 6.411 | 0.3 | 0.009 |
| 201745_at | PTK9 | 10.421 | 10.928 | 0.3 | 0.041 |
| 208697_s_at | EIF3S6 | 12.562 | 13.069 | 0.3 | 0.017 |
| 213864_s_at | NAP1L1 | 12.039 | 12.545 | 0.3 | 0.010 |
| 200012_x_at | RPL21 /// LOC653156 /// LOC653713 /// LOC653730 /// LOC653737 | 13.712 | 14.216 | 0.3 | 0.031 |
| 212844_at | KIAA0179 | 6.203 | 6.705 | 0.3 | 0.049 |
| 211761_s_at | CACYBP | 11.077 | 11.578 | 0.3 | 0.042 |
| 212525_s_at | H2AFX | 5.953 | 6.453 | 0.2 | 0.033 |
| 203818_s_at | SF3A3 | 9.847 | 10.345 | 0.2 | 0.021 |
| 216421_at | --- | 5.137 | 5.634 | 0.2 | 0.003 |
| 213314_at | C6orf162 | 5.552 | 6.048 | 0.2 | 0.021 |
| 208642_s_at | XRCC5 | 12.093 | 12.588 | 0.2 | 0.012 |
| 201461_s_at | MAPKAPK2 | 4.865 | 5.359 | 0.2 | 0.008 |
| 200630_x_at | SET | 12.434 | 12.928 | 0.2 | 0.001 |
| 200647_x_at | EIF3S8 /// LOC653352 | 11.950 | 12.443 | 0.2 | 0.023 |
| 208403_x_at | MAX | 5.519 | 6.011 | 0.2 | 0.037 |
| 211051_s_at | EXTL3 | 5.556 | 6.046 | 0.2 | 0.024 |
| 201400_at | PSMB3 | 11.572 | 12.061 | 0.2 | 0.031 |
| 219131_at | UBIAD1 | 6.817 | 7.305 | 0.2 | 0.038 |
| 211931_s_at | HNRPA3P1 /// HNRPA3 /// LOC643689 /// LOC647474 | 11.423 | 11.911 | 0.2 | 0.006 |
| 217559_at | RPL10L | 4.935 | 5.423 | 0.2 | 0.001 |
| 215150_at | YOD1 | 4.888 | 5.376 | 0.2 | 0.015 |
| 218235_s_at | UTP11L | 10.040 | 10.527 | 0.2 | 0.029 |
| 206248_at | PRKCE | 5.847 | 6.331 | 0.2 | 0.001 |
| 203856_at | VRK1 | 10.289 | 10.771 | 0.2 | 0.038 |
| 216180_s_at | SYNJ2 | 6.240 | 6.721 | 0.2 | 0.036 |
| 206016_at | CCDC22 | 7.322 | 7.803 | 0.2 | 0.028 |
| 208361_s_at | POLR3D | 5.508 | 5.989 | 0.2 | 0.011 |
| 205411_at | STK4 | 5.162 | 5.641 | 0.2 | 0.041 |
| 203415_at | PDCD6 | 9.847 | 10.324 | 0.2 | 0.033 |
| 219273_at | CCNK | 5.170 | 5.647 | 0.2 | 0.002 |

| | | | | | |
|---|---|---|---|---|---|
| 221486_at | ENSA | 9.003 | 9.479 | 0.2 | 0.029 |
| 214993_at | ASPHD1 | 6.234 | 6.709 | 0.2 | 0.038 |
| 207754_at | RASSF8 | 5.073 | 5.546 | 0.2 | 0.024 |
| 219234_x_at | SCRN3 | 4.866 | 5.339 | 0.2 | 0.009 |
| 200735_x_at | NACA | 13.551 | 14.023 | 0.2 | 0.028 |
| 206911_at | TRIM25 | 5.632 | 6.104 | 0.2 | 0.009 |
| 32699_s_at | PVR | 5.278 | 5.748 | 0.2 | 0.007 |
| 210027_s_at | APEX1 | 12.063 | 12.530 | 0.2 | 0.043 |
| 207143_at | CDK6 | 5.648 | 6.115 | 0.2 | 0.031 |
| 201322_at | ATP5B | 11.756 | 12.222 | 0.2 | 0.031 |
| 212215_at | PREPL | 9.712 | 10.178 | 0.2 | 0.019 |
| 203075_at | SMAD2 | 8.075 | 8.540 | 0.2 | 0.010 |
| 219094_at | ARMC8 | 6.367 | 6.832 | 0.2 | 0.003 |
| 211787_s_at | EIF4A1 | 13.190 | 13.654 | 0.2 | 0.021 |
| 214190_x_at | GGA2 | 4.904 | 5.367 | 0.2 | 0.028 |
| 217719_at | EIF3S6IP | 12.814 | 13.276 | 0.2 | 0.015 |
| 218383_at | C14orf94 | 8.351 | 8.811 | 0.2 | 0.022 |
| 200957_s_at | SSRP1 | 10.597 | 11.056 | 0.2 | 0.039 |
| 211091_s_at | NF2 | 5.436 | 5.893 | 0.2 | 0.046 |
| 216856_s_at | --- | 5.500 | 5.955 | 0.2 | 0.039 |
| 220797_at | METT10D | 5.513 | 5.964 | 0.2 | 0.030 |
| 214687_x_at | ALDOA | 12.268 | 12.717 | 0.2 | 0.015 |
| 218708_at | NXT1 | 9.331 | 9.780 | 0.2 | 0.048 |
| 203101_s_at | MGAT2 | 5.602 | 6.049 | 0.2 | 0.021 |
| 200966_x_at | ALDOA | 12.292 | 12.736 | 0.2 | 0.010 |
| 204841_s_at | EEA1 | 5.271 | 5.716 | 0.2 | 0.034 |
| 209776_s_at | SLC19A1 | 5.248 | 5.690 | 0.2 | 0.020 |
| 217313_at | --- | 6.158 | 6.597 | 0.2 | 0.049 |
| 33646_g_at | GM2A | 5.493 | 5.931 | 0.2 | 0.019 |
| 221938_x_at | THRAP5 | 5.874 | 6.312 | 0.2 | 0.015 |
| 208635_x_at | NACA | 13.656 | 14.092 | 0.2 | 0.023 |
| 206473_at | MBTPS2 | 5.365 | 5.800 | 0.2 | 0.049 |
| 201281_at | ADRM1 | 10.206 | 10.640 | 0.2 | 0.028 |
| 215820_x_at | SNX13 | 5.075 | 5.507 | 0.2 | 0.033 |
| 212102_s_at | KPNA6 | 5.397 | 5.829 | 0.2 | 0.009 |
| 210812_at | XRCC4 | 5.478 | 5.909 | 0.2 | 0.023 |
| 217724_at | SERBP1 | 11.629 | 12.059 | 0.2 | 0.011 |
| 200991_s_at | SNX17 | 9.167 | 9.596 | 0.2 | 0.015 |
| 217461_x_at | BTF3 | 5.198 | 5.627 | 0.2 | 0.019 |
| 209984_at | JMJD2C | 6.122 | 6.549 | 0.2 | 0.011 |
| 213810_s_at | C6orf166 | 5.070 | 5.494 | 0.2 | 0.025 |
| 201114_x_at | PSMA7 | 12.106 | 12.530 | 0.2 | 0.021 |
| 218669_at | RAP2C | 8.722 | 9.144 | 0.2 | 0.007 |
| 209830_s_at | SLC9A3R2 | 4.632 | 5.052 | 0.2 | 0.008 |
| 222011_s_at | TCP1 | 8.515 | 8.935 | 0.2 | 0.024 |

| | | | | | |
|---|---|---|---|---|---|
| 202243_s_at | PSMB4 | 11.768 | 12.187 | 0.2 | 0.029 |
| 201623_s_at | DARS | 12.010 | 12.425 | 0.2 | 0.002 |
| 218442_at | TTC4 | 7.346 | 7.760 | 0.2 | 0.037 |
| 205604_at | HOXD9 | 6.009 | 6.421 | 0.2 | 0.032 |
| 213372_at | PAQR3 | 7.631 | 8.038 | 0.2 | 0.029 |
| 210846_x_at | TRIM14 | 5.117 | 5.523 | 0.2 | 0.005 |
| 216880_at | RAD51L1 | 5.278 | 5.683 | 0.2 | 0.010 |
| 202298_at | NDUFA1 | 11.735 | 12.139 | 0.2 | 0.040 |
| 200829_x_at | ZNF207 | 10.939 | 11.343 | 0.2 | 0.033 |
| 217019_at | --- | 5.069 | 5.472 | 0.2 | 0.013 |
| 201213_at | --- | 5.967 | 6.369 | 0.2 | 0.029 |
| 200066_at | IK | 10.397 | 10.797 | 0.2 | 0.005 |
| 208692_at | RPS3 | 13.510 | 13.910 | 0.2 | 0.037 |
| 201270_x_at | NUDCD3 | 8.711 | 9.108 | 0.2 | 0.001 |
| 210455_at | C10orf28 | 5.261 | 5.655 | 0.2 | 0.027 |
| 208836_at | ATP1B3 | 12.084 | 12.475 | 0.2 | 0.027 |
| 200099_s_at | RPS3A /// LOC439992 | 13.696 | 14.088 | 0.2 | 0.037 |
| 203898_at | RCP9 | 7.484 | 7.874 | 0.2 | 0.029 |
| 218301_at | RNPEPL1 | 5.594 | 5.985 | 0.2 | 0.041 |
| 201007_at | HADHB | 10.806 | 11.195 | 0.2 | 0.038 |
| 217503_at | --- | 5.021 | 5.408 | 0.1 | 0.005 |
| 200077_s_at | OAZ1 | 13.304 | 13.690 | 0.1 | 0.023 |
| 89948_at | C20orf67 | 5.549 | 5.932 | 0.1 | 0.045 |
| 217092_x_at | RPL7 /// LOC389305 /// LOC641750 /// LOC643906 /// LOC645737 /// LOC648000 /// LOC649312 /// LOC653702 /// LOC653949 | 10.454 | 10.834 | 0.1 | 0.039 |
| 208619_at | DDB1 | 9.708 | 10.088 | 0.1 | 0.036 |
| 220762_s_at | GNB1L | 6.133 | 6.511 | 0.1 | 0.006 |
| 213323_s_at | ZC3H7B | 5.741 | 6.119 | 0.1 | 0.020 |
| 206744_s_at | ZMYM5 | 4.899 | 5.276 | 0.1 | 0.048 |
| 201665_x_at | RPS17 | 13.825 | 14.201 | 0.1 | 0.024 |
| 212084_at | TEX261 | 6.511 | 6.885 | 0.1 | 0.049 |
| 203721_s_at | UTP18 | 10.984 | 11.356 | 0.1 | 0.047 |
| 211141_s_at | CNOT3 | 5.152 | 5.524 | 0.1 | 0.026 |
| 221098_x_at | UTP14A | 4.775 | 5.143 | 0.1 | 0.020 |
| 205461_at | RAB35 | 5.723 | 6.090 | 0.1 | 0.025 |
| 209519_at | --- | 5.202 | 5.568 | 0.1 | 0.047 |
| 201699_at | PSMC6 | 10.960 | 11.325 | 0.1 | 0.034 |
| 216650_at | --- | 5.909 | 6.272 | 0.1 | 0.038 |
| 221038_at | --- | 4.941 | 5.304 | 0.1 | 0.031 |

| | | | | | |
|---|---|---|---|---|---|
| 217416_x_at | --- | 5.152 | 5.503 | 0.1 | 0.037 |
| 204528_s_at | NAP1L1 | 11.792 | 12.143 | 0.1 | 0.044 |
| 41657_at | STK11 | 6.541 | 6.891 | 0.1 | 0.038 |
| 219153_s_at | THSD4 | 4.901 | 5.250 | 0.1 | 0.026 |
| 206536_s_at | BIRC4 | 5.157 | 5.503 | 0.1 | 0.031 |
| 208843_s_at | GORASP2 | 10.099 | 10.444 | 0.1 | 0.049 |
| 220473_s_at | ZCCHC4 | 4.794 | 5.137 | 0.1 | 0.023 |
| 202550_s_at | VAPB | 10.133 | 10.475 | 0.1 | 0.013 |
| 202657_s_at | SERTAD2 | 9.065 | 9.407 | 0.1 | 0.034 |
| 215519_x_at | RUTBC3 | 5.409 | 5.750 | 0.1 | 0.026 |
| 214737_x_at | HNRPC | 12.031 | 12.369 | 0.1 | 0.030 |
| 208115_x_at | C10orf137 | 5.200 | 5.534 | 0.1 | 0.042 |
| 212042_x_at | RPL7 /// LOC653702 /// LOC653949 | 13.653 | 13.979 | 0.1 | 0.043 |
| 216271_x_at | SYDE1 | 4.890 | 5.215 | 0.1 | 0.014 |
| 210061_at | ZNF589 | 5.358 | 5.682 | 0.1 | 0.031 |
| 211378_x_at | PPIA | 13.934 | 14.257 | 0.1 | 0.035 |
| 209346_s_at | PI4KII | 6.198 | 6.520 | 0.1 | 0.012 |
| 200858_s_at | RPS8 | 12.363 | 12.683 | 0.1 | 0.038 |
| 218278_at | WDR74 | 7.732 | 8.048 | 0.1 | 0.041 |
| 220330_s_at | SAMSN1 | 4.669 | 4.982 | 0.1 | 0.043 |
| 65521_at | UBE2D4 | 5.578 | 5.888 | 0.1 | 0.011 |
| 200888_s_at | RPL23 | 13.191 | 13.500 | 0.1 | 0.037 |
| 200017_at | RPS27A | 13.778 | 14.087 | 0.1 | 0.046 |
| 217924_at | C6orf106 | 5.460 | 5.766 | 0.1 | 0.003 |
| 212425_at | SCAMP1 | 4.978 | 5.282 | 0.1 | 0.003 |
| 208229_at | FGFR2 | 5.171 | 5.475 | 0.1 | 0.037 |
| 211765_x_at | PPIA | 13.930 | 14.233 | 0.1 | 0.041 |
| 215343_at | KIAA1509 | 4.914 | 5.215 | 0.1 | 0.036 |
| 204525_at | PHF14 | 4.844 | 5.143 | 0.1 | 0.026 |
| 214143_x_at | RPL24 /// SLC36A2 | 13.864 | 14.158 | 0.1 | 0.044 |
| 200034_s_at | RPL6 | 13.456 | 13.745 | 0.1 | 0.028 |
| 207594_s_at | SYNJ1 | 4.700 | 4.978 | 0.1 | 0.001 |
| 220103_s_at | MRPS18C | 4.716 | 4.993 | 0.1 | 0.041 |
| 216315_x_at | UBE2V1 /// Kua-UEV | 5.139 | 5.414 | 0.1 | 0.017 |
| 211750_x_at | TUBA6 | 13.789 | 14.063 | 0.1 | 0.039 |
| 215282_at | ANAPC13 | 4.661 | 4.934 | 0.1 | 0.003 |
| 212661_x_at | PPIA | 13.969 | 14.237 | 0.1 | 0.042 |
| 208007_at | --- | 4.612 | 4.876 | 0.1 | 0.008 |
| 214443_at | PVR | 5.462 | 5.719 | 0.1 | 0.047 |
| 208496_x_at | HIST1H3G | 5.488 | 5.744 | 0.1 | 0.020 |
| 208687_x_at | HSPA8 | 13.543 | 13.795 | 0.1 | 0.038 |
| 220472_at | ZCCHC4 | 4.856 | 5.106 | 0.1 | 0.035 |
| 215724_at | PLD1 | 5.167 | 5.416 | 0.1 | 0.038 |
| 222261_at | KIAA1609 | 5.557 | 5.805 | 0.1 | 0.042 |

| | | | | | |
|---|---|---|---|---|---|
| 204884_s_at | HUS1 | 4.757 | 5.004 | 0.1 | 0.019 |
| 208275_x_at | UTF1 | 4.822 | 5.069 | 0.1 | 0.005 |
| 207455_at | P2RY1 | 4.760 | 5.003 | 0.1 | 0.017 |
| 214999_s_at | RAB11FIP3 | 5.505 | 5.747 | 0.1 | 0.008 |
| 210689_at | CLDN14 | 4.884 | 5.124 | 0.1 | 0.048 |
| 220671_at | CCRN4L | 5.226 | 5.465 | 0.1 | 0.038 |
| 200680_x_at | HMGB1 | 13.792 | 14.030 | 0.1 | 0.039 |
| 222142_at | CYLD | 4.818 | 5.051 | 0.1 | 0.022 |
| 207495_at | RAB28 | 5.534 | 5.767 | 0.1 | 0.007 |
| 215927_at | MUC1 | 5.643 | 5.875 | 0.1 | 0.024 |
| 202480_s_at | DEDD | 6.363 | 6.593 | 0.1 | 0.001 |
| 213640_s_at | LOX | 4.554 | 4.783 | 0.1 | 0.014 |
| 219834_at | ALS2CR8 | 4.911 | 5.139 | 0.1 | 0.015 |
| 216354_at | --- | 6.208 | 6.432 | 0.0 | 0.004 |
| 200029_at | RPL19 | 13.718 | 13.939 | 0.0 | 0.041 |
| 211455_at | IFP38 | 4.526 | 4.747 | 0.0 | 0.010 |
| 209223_at | NDUFA2 | 4.980 | 5.200 | 0.0 | 0.004 |
| 208724_s_at | RAB1A | 12.193 | 12.410 | 0.0 | 0.033 |
| 216279_at | ZNF272 | 4.856 | 5.072 | 0.0 | 0.016 |
| 220315_at | PARP11 | 4.817 | 5.033 | 0.0 | 0.019 |
| 220236_at | PDPR | 4.990 | 5.204 | 0.0 | 0.021 |
| 220457_at | SAMD4B | 4.900 | 5.114 | 0.0 | 0.037 |
| 208049_s_at | TACR1 | 4.940 | 5.152 | 0.0 | 0.031 |
| 220229_s_at | AP4E1 | 5.002 | 5.213 | 0.0 | 0.022 |
| 210972_x_at | TRA@ /// TRDV2 /// TRAV20 /// TRAC | 4.844 | 5.054 | 0.0 | 0.041 |
| 212819_at | ASB1 | 5.169 | 5.372 | 0.0 | 0.033 |
| 206243_at | TIMP4 | 5.084 | 5.286 | 0.0 | 0.020 |
| 213829_x_at | RTEL1 | 6.110 | 6.311 | 0.0 | 0.032 |
| 220931_at | MGC5590 | 4.836 | 5.037 | 0.0 | 0.030 |
| 206743_s_at | ASGR1 | 5.210 | 5.409 | 0.0 | 0.030 |
| 222224_at | NACAL | 4.788 | 4.987 | 0.0 | 0.013 |
| 204270_at | --- | 5.132 | 5.329 | 0.0 | 0.011 |
| 217714_x_at | STMN1 | 6.620 | 6.814 | 0.0 | 0.043 |
| 211782_at | IDS | 4.749 | 4.940 | 0.0 | 0.048 |
| 210000_s_at | SOCS1 | 4.706 | 4.892 | 0.0 | 0.007 |
| 214994_at | APOBEC3F | 4.943 | 5.128 | 0.0 | 0.034 |
| 220360_at | THAP9 | 4.483 | 4.668 | 0.0 | 0.033 |
| 210067_at | AQP4 | 4.857 | 5.042 | 0.0 | 0.026 |
| 202526_at | SMAD4 | 4.869 | 5.054 | 0.0 | 0.041 |
| 220437_at | LOC55908 | 4.867 | 5.052 | 0.0 | 0.010 |
| 215324_at | SEMA3D | 4.830 | 5.008 | 0.0 | 0.036 |
| 210997_at | HGF | 4.631 | 4.807 | 0.0 | 0.048 |
| 207451_at | NKX2-8 | 5.289 | 5.464 | 0.0 | 0.049 |
| 207797_s_at | LRP2BP | 4.893 | 5.067 | 0.0 | 0.021 |

| | | | | | |
|---|---|---|---|---|---|
| 201187_s_at | ITPR3 | 5.141 | 5.307 | 0.0 | 0.029 |
| 211174_s_at | CCKAR | 4.692 | 4.858 | 0.0 | 0.007 |
| 216093_at | NCAM1 | 4.824 | 4.987 | 0.0 | 0.037 |
| 216737_at | --- | 4.733 | 4.894 | 0.0 | 0.036 |
| 210193_at | MOBP | 5.458 | 5.618 | 0.0 | 0.042 |
| 208166_at | MMP16 | 4.859 | 5.019 | 0.0 | 0.018 |
| 214938_x_at | HMGB1 | 13.749 | 13.904 | 0.0 | 0.039 |
| 215345_x_at | TRGV7 | 5.008 | 5.163 | 0.0 | 0.014 |
| 221405_at | LOC51190 | 4.815 | 4.965 | 0.0 | 0.044 |
| 216369_at | --- | 4.649 | 4.799 | 0.0 | 0.023 |
| 211445_x_at | NACAP1 /// LOC389240 | 11.658 | 11.804 | 0.0 | 0.046 |
| 204150_at | STAB1 | 4.929 | 5.075 | 0.0 | 0.044 |
| 216490_x_at | --- | 5.282 | 5.425 | 0.0 | 0.029 |
| 213265_at | PGA5 /// LOC643834 /// LOC643847 | 5.146 | 5.289 | 0.0 | 0.019 |
| 207457_s_at | LY6G6D | 5.113 | 5.255 | 0.0 | 0.016 |
| 210626_at | AKAP1 | 5.143 | 5.283 | 0.0 | 0.023 |
| 220679_s_at | CDH7 | 4.820 | 4.954 | 0.0 | 0.026 |
| 211663_x_at | PTGDS | 4.743 | 4.872 | 0.0 | 0.016 |
| 207552_at | ATP5G2 | 4.762 | 4.886 | 0.0 | 0.042 |
| 207158_at | APOBEC1 | 4.782 | 4.905 | 0.0 | 0.045 |
| 220167_s_at | TP53TG3 /// LOC653535 /// LOC653550 /// LOC653551 | 4.929 | 5.052 | 0.0 | 0.050 |
| 207978_s_at | NR4A3 | 4.957 | 5.079 | 0.0 | 0.048 |
| 216981_x_at | SPN | 5.034 | 5.155 | 0.0 | 0.036 |
| 51226_at | RBM12B | 4.776 | 4.895 | 0.0 | 0.003 |
| 209987_s_at | ASCL1 | 4.716 | 4.835 | 0.0 | 0.043 |
| 208569_at | HIST1H2AB | 4.704 | 4.818 | 0.0 | 0.021 |
| 209671_x_at | TRA@ /// TRAC | 4.837 | 4.948 | 0.0 | 0.037 |
| 216440_at | RAB6IP2 | 4.761 | 4.868 | 0.0 | 0.044 |
| 204938_s_at | PLN | 4.475 | 4.580 | 0.0 | 0.026 |
| 207938_at | PI15 | 4.963 | 5.069 | 0.0 | 0.048 |
| 219436_s_at | EMCN | 4.785 | 4.890 | 0.0 | 0.041 |
| 216410_at | --- | 4.798 | 4.902 | 0.0 | 0.034 |
| 207161_at | KIAA0087 | 4.638 | 4.742 | 0.0 | 0.024 |
| 215666_at | HLA-DRB4 | 4.792 | 4.893 | 0.0 | 0.044 |
| 215524_x_at | TRA@ /// TRDV2 /// TRAV20 /// TRAC | 4.640 | 4.736 | 0.0 | 0.025 |
| 217474_at | --- | 5.064 | 5.159 | 0.0 | 0.012 |
| 210411_s_at | GRIN2B | 4.825 | 4.919 | 0.0 | 0.048 |
| 217568_at | FAM12A | 4.592 | 4.675 | 0.0 | 0.039 |
| *** log2 transformed RMA values** | | | | | |
