## Supplementary material for "Targeting dormant ovarian cancer cells *in vitro* and in an *in vivo* model of platinum resistance": Table S3

| Table S3. Genes with increased expression in low spheroid-forming capacity cell lines | | | | | |
|---|---|---|---|---|---|
| **U133A probe** | **Gene Symbol** | **Avg NSF*** | **Avg SF*** | **Fold Change** | **t test p** |
| 209771_x_at | CD24 | 12.557 | 7.213 | 28.556 | 0.001 |
| 216379_x_at | CD24 | 12.570 | 7.386 | 26.873 | 0.000 |
| 201839_s_at | TACSTD1 | 12.104 | 6.997 | 26.080 | 0.007 |
| 266_s_at | CD24 | 10.982 | 6.014 | 24.681 | 0.000 |
| 208651_x_at | CD24 | 10.288 | 5.762 | 20.480 | 0.000 |
| 202826_at | SPINT1 | 9.370 | 4.951 | 19.527 | 0.040 |
| 208650_s_at | CD24 /// LOC647456 | 11.094 | 6.740 | 18.962 | 0.000 |
| 212314_at | KIAA0746 | 10.416 | 6.383 | 16.263 | 0.009 |
| 202149_at | NEDD9 | 9.361 | 5.331 | 16.243 | 0.010 |
| 202454_s_at | ERBB3 | 9.385 | 5.490 | 15.166 | 0.003 |
| 203066_at | GALNAC4S-6ST | 8.714 | 4.951 | 14.163 | 0.002 |
| 203397_s_at | GALNT3 | 9.601 | 5.861 | 13.987 | 0.019 |
| 214355_x_at | LOC441296 | 10.763 | 7.061 | 13.708 | 0.006 |
| 208190_s_at | LSR | 9.482 | 5.871 | 13.034 | 0.007 |
| 209772_s_at | CD24 | 10.194 | 6.586 | 13.016 | 0.002 |
| 202838_at | FUCA1 | 10.234 | 6.649 | 12.853 | 0.006 |
| 215549_x_at | LOC643854 | 9.650 | 6.137 | 12.337 | 0.009 |
| 212311_at | KIAA0746 | 9.341 | 5.895 | 11.877 | 0.018 |
| 200606_at | DSP | 12.010 | 8.586 | 11.723 | 0.023 |
| 204114_at | NID2 | 7.827 | 4.486 | 11.158 | 0.007 |
| 202948_at | IL1R1 | 8.878 | 5.609 | 10.684 | 0.013 |
| 32137_at | JAG2 | 8.911 | 5.651 | 10.626 | 0.028 |
| 218706_s_at | GRAMD3 | 10.515 | 7.276 | 10.489 | 0.005 |
| 212816_s_at | CBS | 9.605 | 6.416 | 10.170 | 0.008 |
| 65517_at | AP1M2 | 9.747 | 6.619 | 9.785 | 0.018 |
| 210715_s_at | SPINT2 | 11.742 | 8.648 | 9.570 | 0.031 |
| 204990_s_at | ITGB4 | 9.428 | 6.336 | 9.557 | 0.003 |
| 37117_at | ARHGAP8 /// LOC553158 | 8.446 | 5.376 | 9.420 | 0.032 |
| 218803_at | CHFR | 9.125 | 6.097 | 9.171 | 0.007 |
| 218261_at | AP1M2 | 9.960 | 6.994 | 8.796 | 0.036 |
| 209784_s_at | JAG2 | 7.938 | 5.005 | 8.601 | 0.044 |
| 210749_x_at | DDR1 | 9.512 | 6.624 | 8.339 | 0.013 |
| 202295_s_at | CTSH | 9.702 | 6.821 | 8.297 | 0.010 |
| 204029_at | CELSR2 | 8.555 | 5.705 | 8.121 | 0.043 |
| 207169_x_at | DDR1 | 9.658 | 6.818 | 8.060 | 0.011 |
| 201015_s_at | JUP | 9.444 | 6.631 | 7.910 | 0.020 |
| 206683_at | ZNF165 | 7.419 | 4.611 | 7.887 | 0.022 |
| 36499_at | CELSR2 | 8.353 | 5.576 | 7.716 | 0.049 |
| 204765_at | ARHGEF5 | 8.774 | 6.106 | 7.115 | 0.002 |
| 201694_s_at | EGR1 | 9.220 | 6.581 | 6.965 | 0.045 |
| 1007_s_at | DDR1 | 9.841 | 7.223 | 6.859 | 0.016 |
| 208779_x_at | DDR1 | 8.765 | 6.156 | 6.807 | 0.025 |

| | | | | | |
|---|---|---|---|---|---|
| 41660_at | CELSR1 | 9.088 | 6.527 | 6.560 | 0.020 |
| 219228_at | ZNF331 | 9.258 | 6.751 | 6.287 | 0.020 |
| 219976_at | HOOK1 | 8.097 | 5.619 | 6.141 | 0.013 |
| 221645_s_at | ZNF83 | 9.596 | 7.169 | 5.893 | 0.016 |
| 215299_x_at | SULT1A1 | 9.688 | 7.263 | 5.882 | 0.017 |
| 220668_s_at | DNMT3B | 9.375 | 6.955 | 5.853 | 0.024 |
| 202421_at | IGSF3 | 9.805 | 7.390 | 5.833 | 0.023 |
| 220609_at | LOC202181 | 7.969 | 5.583 | 5.693 | 0.000 |
| 218312_s_at | ZNF447 | 8.507 | 6.125 | 5.672 | 0.022 |
| 213359_at | HNRPD | 10.818 | 8.463 | 5.546 | 0.002 |
| 206295_at | IL18 | 8.852 | 6.503 | 5.517 | 0.041 |
| 203139_at | DAPK1 | 9.848 | 7.502 | 5.505 | 0.027 |
| 202761_s_at | SYNE2 | 9.910 | 7.588 | 5.389 | 0.049 |
| 218066_at | SLC12A7 | 8.741 | 6.421 | 5.382 | 0.004 |
| 218780_at | HOOK2 | 8.804 | 6.530 | 5.172 | 0.007 |
| 213929_at | --- | 8.835 | 6.608 | 4.957 | 0.028 |
| 210598_at | --- | 8.142 | 5.926 | 4.909 | 0.012 |
| 203713_s_at | LLGL2 | 7.693 | 5.568 | 4.516 | 0.003 |
| 203569_s_at | OFD1 | 10.266 | 8.142 | 4.508 | 0.023 |
| 219681_s_at | RAB11FIP1 | 8.086 | 5.995 | 4.370 | 0.031 |
| 209873_s_at | PKP3 | 8.280 | 6.197 | 4.338 | 0.001 |
| 203071_at | SEMA3B | 7.808 | 5.740 | 4.276 | 0.042 |
| 213792_s_at | --- | 8.380 | 6.322 | 4.236 | 0.019 |
| 216836_s_at | ERBB2 | 9.321 | 7.270 | 4.205 | 0.001 |
| 206247_at | MICB | 9.679 | 7.632 | 4.191 | 0.027 |
| 203615_x_at | SULT1A1 | 9.138 | 7.093 | 4.185 | 0.050 |
| 206687_s_at | PTPN6 | 8.582 | 6.536 | 4.185 | 0.003 |
| 206261_at | ZNF239 | 8.465 | 6.424 | 4.166 | 0.027 |
| 202178_at | PRKCZ | 8.745 | 6.714 | 4.123 | 0.010 |
| 213737_x_at | GOLGA8G /// GOLGA8D /// LOC388189 /// GOLGA8E /// GOLGA8C /// GOLGA8F | 10.638 | 8.619 | 4.076 | 0.000 |
| 40837_at | TLE2 | 7.888 | 5.877 | 4.043 | 0.018 |
| 204717_s_at | SLC29A2 | 7.642 | 5.646 | 3.984 | 0.022 |
| 217820_s_at | ENAH | 10.323 | 8.329 | 3.974 | 0.046 |
| 206723_s_at | EDG4 | 7.914 | 5.925 | 3.954 | 0.010 |
| 209607_x_at | SULT1A3 /// SULT1A4 /// LOC648394 | 9.281 | 7.324 | 3.828 | 0.025 |
| 201613_s_at | AP1G2 | 9.696 | 7.745 | 3.805 | 0.025 |
| 217520_x_at | LOC283683 /// LOC646278 | 7.966 | 6.022 | 3.780 | 0.032 |
| 212807_s_at | SORT1 | 9.045 | 7.105 | 3.764 | 0.004 |
| 213340_s_at | KIAA0495 | 7.824 | 5.912 | 3.655 | 0.007 |
| 221860_at | HNRPL | 10.391 | 8.493 | 3.603 | 0.030 |

| | | | | | |
|---|---|---|---|---|---|
| 212727_at | DLG3 | 8.342 | 6.456 | 3.556 | 0.013 |
| 203765_at | GCA | 7.350 | 5.475 | 3.515 | 0.002 |
| 204321_at | NEO1 | 7.404 | 5.533 | 3.498 | 0.000 |
| 205525_at | CALD1 | 8.321 | 6.451 | 3.497 | 0.041 |
| 210266_s_at | TRIM33 | 11.407 | 9.548 | 3.455 | 0.050 |
| 220728_at | --- | 8.237 | 6.392 | 3.401 | 0.018 |
| 202908_at | WFS1 | 9.015 | 7.187 | 3.340 | 0.003 |
| 204497_at | ADCY9 | 9.252 | 7.426 | 3.335 | 0.000 |
| 215006_at | EZH2 | 9.028 | 7.218 | 3.275 | 0.013 |
| 209163_at | CYB561 | 9.119 | 7.312 | 3.269 | 0.023 |
| 207394_at | ZNF137 | 6.933 | 5.129 | 3.253 | 0.006 |
| 209845_at | MKRN1 | 9.672 | 7.870 | 3.247 | 0.007 |
| 216211_at | C10orf18 | 8.568 | 6.770 | 3.235 | 0.010 |
| 212321_at | --- | 8.640 | 6.844 | 3.226 | 0.000 |
| 204671_s_at | ANKRD6 | 6.693 | 4.905 | 3.200 | 0.028 |
| 212441_at | KIAA0232 | 10.411 | 8.628 | 3.179 | 0.022 |
| 216563_at | ANKRD12 | 8.324 | 6.544 | 3.169 | 0.027 |
| 215807_s_at | PLXNB1 | 7.288 | 5.512 | 3.155 | 0.027 |
| 213939_s_at | RUFY3 | 9.877 | 8.109 | 3.127 | 0.000 |
| 214114_x_at | FASTK | 9.169 | 7.418 | 3.066 | 0.012 |
| 214703_s_at | MAN2B2 | 8.987 | 7.243 | 3.043 | 0.019 |
| 205514_at | ZNF415 | 7.139 | 5.397 | 3.034 | 0.023 |
| 210074_at | CTSL2 | 8.937 | 7.201 | 3.015 | 0.001 |
| 222062_at | IL27RA | 7.721 | 5.985 | 3.014 | 0.012 |
| 222316_at | VDP | 7.803 | 6.073 | 2.994 | 0.000 |
| 209403_at | TBC1D3 /// TBC1D3C /// TBC1D3D /// LOC653380 /// LOC653498 /// TBC1D3H /// TBC1D3F /// TBC1D3E | 9.042 | 7.326 | 2.944 | 0.004 |
| 209369_at | ANXA3 | 11.191 | 9.477 | 2.940 | 0.006 |
| 213085_s_at | WWC1 | 10.280 | 8.593 | 2.845 | 0.031 |
| 210580_x_at | SULT1A3 /// SULT1A4 | 10.107 | 8.423 | 2.836 | 0.049 |
| 203608_at | ALDH5A1 | 8.074 | 6.393 | 2.827 | 0.016 |
| 220954_s_at | PILRB | 9.207 | 7.544 | 2.764 | 0.001 |
| 214948_s_at | TMF1 | 11.149 | 9.491 | 2.747 | 0.020 |
| 204088_at | P2RX4 | 7.885 | 6.234 | 2.729 | 0.013 |
| 202871_at | TRAF4 | 7.972 | 6.337 | 2.672 | 0.011 |
| 216115_at | NF1 | 8.728 | 7.097 | 2.660 | 0.001 |
| 216060_s_at | DAAM1 | 9.170 | 7.552 | 2.619 | 0.020 |
| 200704_at | LITAF | 9.745 | 8.132 | 2.602 | 0.024 |
| 207746_at | POLQ | 8.900 | 7.295 | 2.579 | 0.037 |
| 209211_at | KLF5 | 9.058 | 7.459 | 2.559 | 0.038 |
| 215191_at | --- | 8.610 | 7.011 | 2.555 | 0.046 |
| 221088_s_at | PPP1R9A | 8.088 | 6.491 | 2.549 | 0.045 |

| | | | | | |
|---|---|---|---|---|---|
| 213230_at | CDR2L | 8.387 | 6.791 | 2.546 | 0.004 |
| 203973_s_at | CEBPD | 8.591 | 6.997 | 2.541 | 0.030 |
| 216550_x_at | ANKRD12 | 7.580 | 5.989 | 2.533 | 0.025 |
| 215206_at | EXT1 | 10.501 | 8.914 | 2.518 | 0.026 |
| 212978_at | LRRC8B | 9.296 | 7.713 | 2.506 | 0.006 |
| 204328_at | TMC6 | 7.316 | 5.734 | 2.501 | 0.017 |
| 211996_s_at | LOC23117 /// DKFZp547E087 /// LOC440345 /// LOC440353 /// LOC613037 /// LOC647048 | 10.901 | 9.320 | 2.499 | 0.019 |
| 202813_at | TARBP1 | 9.558 | 7.983 | 2.482 | 0.001 |
| 204730_at | RIMS3 | 6.998 | 5.424 | 2.478 | 0.026 |
| 222258_s_at | SH3BP4 | 11.155 | 9.588 | 2.457 | 0.021 |
| 212285_s_at | AGRN | 9.662 | 8.095 | 2.454 | 0.014 |
| 209925_at | OCLN /// LOC647859 /// LOC653400 | 6.610 | 5.056 | 2.415 | 0.028 |
| 206023_at | NMU | 10.366 | 8.813 | 2.412 | 0.007 |
| 207011_s_at | PTK7 | 8.688 | 7.139 | 2.397 | 0.018 |
| 218942_at | PIP5K2C | 9.958 | 8.412 | 2.388 | 0.019 |
| 213266_at | 76P | 8.603 | 7.067 | 2.360 | 0.008 |
| 217419_x_at | AGRN | 9.138 | 7.609 | 2.339 | 0.006 |
| 208812_x_at | HLA-C | 11.797 | 10.269 | 2.334 | 0.018 |
| 215474_at | C8orf36 | 7.341 | 5.814 | 2.333 | 0.032 |
| 214715_x_at | ZNF160 | 10.718 | 9.197 | 2.312 | 0.017 |
| 219573_at | LRRC16 | 7.481 | 5.961 | 2.311 | 0.020 |
| 202106_at | GOLGA3 | 9.380 | 7.865 | 2.294 | 0.001 |
| 203016_s_at | SSX2IP | 9.906 | 8.393 | 2.289 | 0.037 |
| 214048_at | MBD4 | 8.263 | 6.752 | 2.281 | 0.014 |
| 220777_at | KIF13A | 6.622 | 5.120 | 2.254 | 0.009 |
| 218429_s_at | FLJ11286 | 8.453 | 6.962 | 2.223 | 0.011 |
| 204980_at | CLOCK | 10.373 | 8.884 | 2.219 | 0.032 |
| 215066_at | PTPRF | 8.359 | 6.870 | 2.216 | 0.011 |
| 207120_at | ZNF667 | 6.032 | 4.544 | 2.214 | 0.048 |
| 217679_x_at | --- | 11.084 | 9.597 | 2.211 | 0.007 |
| 212322_at | SGPL1 | 10.024 | 8.539 | 2.204 | 0.006 |
| 214060_at | SSBP1 | 7.313 | 5.829 | 2.201 | 0.006 |
| 212517_at | ATRN | 8.699 | 7.216 | 2.199 | 0.009 |
| 53720_at | FLJ11286 | 7.120 | 5.638 | 2.194 | 0.006 |
| 213351_s_at | TMCC1 | 7.731 | 6.251 | 2.191 | 0.027 |
| 214046_at | --- | 5.999 | 4.522 | 2.181 | 0.046 |
| 214594_x_at | ATP8B1 | 9.355 | 7.884 | 2.165 | 0.036 |
| 212359_s_at | KIAA0913 | 8.619 | 7.148 | 2.163 | 0.009 |
| 212959_s_at | GNPTAB | 9.900 | 8.430 | 2.162 | 0.009 |

| | | | | | |
|---|---|---|---|---|---|
| 202894_at | EPHB4 | 9.352 | 7.883 | 2.159 | 0.003 |
| 204148_s_at | ZP3 /// POMZP3 | 7.859 | 6.401 | 2.126 | 0.016 |
| 213089_at | LOC153561 /// LOC643367 /// LOC643373 /// LOC653412 | 9.795 | 8.340 | 2.116 | 0.002 |
| 220711_at | --- | 6.841 | 5.397 | 2.087 | 0.015 |
| 215866_at | EEF1G | 7.117 | 5.682 | 2.058 | 0.024 |
| 218652_s_at | PIGG | 9.983 | 8.549 | 2.055 | 0.009 |
| 207365_x_at | USP34 | 9.233 | 7.802 | 2.045 | 0.009 |
| 213212_x_at | LOC161527 /// LOC642346 /// LOC643696 | 10.034 | 8.605 | 2.041 | 0.003 |
| 211034_s_at | FLJ30092 | 9.083 | 7.657 | 2.033 | 0.004 |
| 204927_at | RASSF7 | 8.138 | 6.715 | 2.025 | 0.035 |
| 213700_s_at | PKM2 | 7.801 | 6.379 | 2.021 | 0.022 |
| 202704_at | TOB1 | 11.125 | 9.704 | 2.018 | 0.001 |
| 204447_at | ProSAPiP1 | 7.986 | 6.567 | 2.014 | 0.005 |
| 222186_at | ZA20D3 | 8.372 | 6.955 | 2.007 | 0.036 |
| 202812_at | GAA | 8.176 | 6.762 | 1.998 | 0.013 |
| 221768_at | SFPQ | 11.070 | 9.657 | 1.997 | 0.013 |
| 212876_at | B4GALT4 | 8.430 | 7.018 | 1.992 | 0.018 |
| 221249_s_at | FAM117A | 8.620 | 7.209 | 1.991 | 0.047 |
| 209212_s_at | KLF5 | 9.394 | 7.984 | 1.989 | 0.032 |
| 215588_x_at | RIOK3 | 9.183 | 7.774 | 1.986 | 0.020 |
| 218319_at | PELI1 | 8.976 | 7.571 | 1.974 | 0.036 |
| 215470_at | DKFZP686M0199 | 10.026 | 8.627 | 1.955 | 0.011 |
| 212757_s_at | CAMK2G | 8.667 | 7.270 | 1.952 | 0.015 |
| 201056_at | GOLGB1 | 8.720 | 7.330 | 1.931 | 0.013 |
| 202951_at | STK38 | 10.085 | 8.696 | 1.928 | 0.003 |
| 222030_at | SIVA | 7.410 | 6.023 | 1.923 | 0.003 |
| 213046_at | PABPN1 | 8.134 | 6.747 | 1.923 | 0.000 |
| 215067_x_at | PRDX2 | 7.820 | 6.437 | 1.913 | 0.018 |
| 218686_s_at | RHBDF1 | 8.872 | 7.492 | 1.905 | 0.037 |
| 211048_s_at | PDIA4 | 11.446 | 10.067 | 1.899 | 0.038 |
| 201817_at | UBE3C | 9.997 | 8.620 | 1.897 | 0.026 |
| 222371_at | PIAS1 | 8.828 | 7.458 | 1.877 | 0.003 |
| 202435_s_at | CYP1B1 | 6.516 | 5.150 | 1.867 | 0.042 |
| 219500_at | CLCF1 | 7.985 | 6.623 | 1.857 | 0.014 |
| 213605_s_at | LOC643373 /// LOC653080 /// LOC653188 /// LOC653412 | 11.300 | 9.943 | 1.840 | 0.006 |
| 216960_s_at | ZNF133 | 7.711 | 6.361 | 1.822 | 0.018 |
| 212080_at | MLL | 9.802 | 8.452 | 1.821 | 0.030 |
| 216109_at | THRAP2 | 6.773 | 5.423 | 1.821 | 0.042 |

| | | | | | |
|---|---|---|---|---|---|
| 207730_x_at | HDGF2 | 8.346 | 7.002 | 1.807 | 0.013 |
| 212975_at | DENND3 | 7.026 | 5.683 | 1.806 | 0.016 |
| 204453_at | ZNF84 | 9.098 | 7.758 | 1.797 | 0.001 |
| 217586_x_at | --- | 10.179 | 8.840 | 1.794 | 0.000 |
| 214322_at | CAMK2G | 7.418 | 6.079 | 1.792 | 0.003 |
| 201285_at | MKRN1 | 9.637 | 8.300 | 1.787 | 0.000 |
| 218476_at | POMT1 | 7.909 | 6.573 | 1.786 | 0.033 |
| 213727_x_at | MPPE1 | 8.265 | 6.929 | 1.783 | 0.008 |
| 202185_at | PLOD3 | 10.693 | 9.358 | 1.783 | 0.035 |
| 212753_at | PCGF3 | 9.935 | 8.603 | 1.775 | 0.004 |
| 201057_s_at | GOLGB1 | 10.461 | 9.131 | 1.769 | 0.028 |
| 215235_at | SPTAN1 | 11.075 | 9.746 | 1.765 | 0.029 |
| 201368_at | ZFP36L2 | 11.028 | 9.703 | 1.756 | 0.004 |
| 40093_at | BCAM | 7.713 | 6.395 | 1.737 | 0.039 |
| 218683_at | PTBP2 | 9.015 | 7.698 | 1.734 | 0.001 |
| 217653_x_at | LOC653471 /// LOC654000 | 10.360 | 9.048 | 1.722 | 0.016 |
| 214459_x_at | HLA-C | 11.273 | 9.963 | 1.715 | 0.003 |
| 210425_x_at | GOLGA8B | 10.219 | 8.915 | 1.701 | 0.027 |
| 222282_at | PAPD4 | 9.600 | 8.300 | 1.691 | 0.004 |
| 210540_s_at | B4GALT4 | 8.482 | 7.186 | 1.680 | 0.003 |
| 215599_at | SMA4 /// LOC643367 /// LOC643373 /// LOC652924 /// LOC653869 | 9.686 | 8.391 | 1.676 | 0.025 |
| 206548_at | FLJ23556 | 8.382 | 7.089 | 1.673 | 0.048 |
| 220917_s_at | WDR19 | 8.283 | 6.991 | 1.670 | 0.010 |
| 202307_s_at | TAP1 | 7.599 | 6.307 | 1.668 | 0.020 |
| 217200_x_at | CYB561 | 8.821 | 7.531 | 1.665 | 0.014 |
| 209858_x_at | MPPE1 | 7.847 | 6.560 | 1.654 | 0.007 |
| 218920_at | FLJ10404 | 8.946 | 7.661 | 1.652 | 0.008 |
| 212611_at | DTX4 | 7.203 | 5.929 | 1.621 | 0.027 |
| 206848_at | HOXA7 | 8.478 | 7.208 | 1.612 | 0.037 |
| 215029_at | --- | 7.487 | 6.217 | 1.612 | 0.024 |
| 218684_at | LRRC8D | 10.986 | 9.718 | 1.608 | 0.020 |
| 212980_at | AHSA2 | 8.842 | 7.575 | 1.605 | 0.011 |
| 215387_x_at | GPC6 | 6.987 | 5.722 | 1.601 | 0.047 |
| 201204_s_at | RRBP1 | 10.685 | 9.420 | 1.601 | 0.043 |
| 202423_at | MYST3 | 10.052 | 8.788 | 1.600 | 0.040 |
| 203566_s_at | AGL | 10.020 | 8.759 | 1.591 | 0.020 |
| 220796_x_at | SLC35E1 | 10.986 | 9.728 | 1.583 | 0.012 |
| 213742_at | SFRS11 | 9.253 | 7.996 | 1.580 | 0.044 |
| 216187_x_at | KNS2 | 11.061 | 9.805 | 1.579 | 0.006 |
| 209256_s_at | KIAA0265 | 8.876 | 7.624 | 1.568 | 0.036 |
| 215385_at | FTO | 8.672 | 7.420 | 1.566 | 0.002 |

| | | | | | |
|---|---|---|---|---|---|
| 48106_at | FLJ20489 | 8.035 | 6.788 | 1.556 | 0.019 |
| 202676_x_at | FASTK | 8.651 | 7.406 | 1.551 | 0.020 |
| 218639_s_at | ZXDC | 8.352 | 7.108 | 1.547 | 0.008 |
| 208009_s_at | ARHGEF16 | 6.593 | 5.350 | 1.547 | 0.023 |
| 219595_at | ZNF26 | 9.010 | 7.768 | 1.542 | 0.023 |
| 215604_x_at | UBE2D2 | 9.235 | 7.995 | 1.538 | 0.014 |
| 210694_s_at | --- | 8.053 | 6.813 | 1.536 | 0.005 |
| 209367_at | STXBP2 | 8.547 | 7.308 | 1.535 | 0.007 |
| 201369_s_at | ZFP36L2 | 7.070 | 5.833 | 1.530 | 0.007 |
| 216170_at | EEF1G | 6.631 | 5.397 | 1.522 | 0.006 |
| 201906_s_at | CTDSPL | 8.475 | 7.243 | 1.516 | 0.017 |
| 205417_s_at | DAG1 | 11.047 | 9.818 | 1.511 | 0.001 |
| 209989_at | ZNF268 | 8.104 | 6.877 | 1.506 | 0.008 |
| 218017_s_at | TMEM76 /// LOC643642 | 9.248 | 8.025 | 1.495 | 0.043 |
| 203623_at | PLXNA3 | 6.464 | 5.243 | 1.490 | 0.031 |
| 218159_at | C20orf116 | 9.796 | 8.577 | 1.485 | 0.001 |
| 214686_at | ZNF266 | 9.887 | 8.668 | 1.485 | 0.045 |
| 212053_at | KIAA0251 | 9.851 | 8.649 | 1.446 | 0.001 |
| 206764_x_at | MPPE1 | 7.518 | 6.318 | 1.441 | 0.011 |
| 212339_at | EPB41L1 | 7.065 | 5.864 | 1.441 | 0.033 |
| 215200_x_at | VIL2 | 8.618 | 7.422 | 1.431 | 0.001 |
| 217704_x_at | SUZ12P | 9.576 | 8.380 | 1.428 | 0.006 |
| 215750_at | KIAA1659 | 6.984 | 5.791 | 1.423 | 0.033 |
| 40446_at | PHF1 | 10.113 | 8.922 | 1.421 | 0.043 |
| 213670_x_at | NSUN5B | 9.296 | 8.104 | 1.420 | 0.034 |
| 215269_at | TMEM1 | 8.147 | 6.955 | 1.420 | 0.012 |
| 205059_s_at | IDUA | 6.384 | 5.193 | 1.418 | 0.033 |
| 217703_x_at | SPON1 | 7.337 | 6.147 | 1.416 | 0.044 |
| 203301_s_at | DMTF1 | 10.768 | 9.579 | 1.415 | 0.003 |
| 203221_at | TLE1 | 10.574 | 9.386 | 1.411 | 0.036 |
| 213460_x_at | NSUN5C | 9.798 | 8.611 | 1.411 | 0.027 |
| 215978_x_at | LOC152719 | 11.393 | 10.206 | 1.410 | 0.015 |
| 212860_at | ZDHHC18 | 7.243 | 6.058 | 1.405 | 0.040 |
| 213424_at | KIAA0895 | 6.921 | 5.736 | 1.404 | 0.004 |
| 203518_at | LYST | 6.766 | 5.581 | 1.404 | 0.041 |
| 214707_x_at | ALMS1 | 8.015 | 6.832 | 1.398 | 0.019 |
| 212987_at | FBXO9 | 10.920 | 9.743 | 1.383 | 0.048 |
| 214100_x_at | NSUN5B | 10.049 | 8.875 | 1.378 | 0.046 |
| 209467_s_at | MKNK1 | 9.685 | 8.516 | 1.367 | 0.016 |
| 218417_s_at | FLJ20489 | 6.996 | 5.827 | 1.367 | 0.037 |
| 208246_x_at | --- | 11.167 | 9.998 | 1.367 | 0.004 |
| 216026_s_at | POLE | 8.942 | 7.775 | 1.361 | 0.001 |
| 209446_s_at | --- | 7.701 | 6.534 | 1.360 | 0.015 |
| 78495_at | DKFZp762P2111 | 7.679 | 6.513 | 1.360 | 0.011 |
| 218259_at | MKL2 | 9.075 | 7.909 | 1.359 | 0.001 |

| | | | | | |
|---|---|---|---|---|---|
| 215545_at | --- | 8.346 | 7.181 | 1.357 | 0.033 |
| 208798_x_at | GOLGA8A | 11.819 | 10.654 | 1.357 | 0.029 |
| 216858_x_at | --- | 9.101 | 7.937 | 1.355 | 0.004 |
| 213650_at | GOLGA8A /// GOLGA8B | 8.576 | 7.413 | 1.354 | 0.001 |
| 204137_at | GPR137B | 7.444 | 6.281 | 1.351 | 0.010 |
| 204161_s_at | ENPP4 | 6.052 | 4.891 | 1.348 | 0.048 |
| 222214_at | SUZ12P | 8.690 | 7.529 | 1.348 | 0.024 |
| 209315_at | HBS1L | 7.165 | 6.006 | 1.345 | 0.011 |
| 203600_s_at | C4orf8 | 8.944 | 7.786 | 1.340 | 0.000 |
| 215786_at | HBXAP | 7.579 | 6.424 | 1.335 | 0.017 |
| 212747_at | ANKS1A | 8.571 | 7.416 | 1.334 | 0.034 |
| 215201_at | REPS1 | 6.903 | 5.749 | 1.331 | 0.014 |
| 216526_x_at | HLA-C | 11.592 | 10.439 | 1.330 | 0.009 |
| 222266_at | C19orf2 | 10.699 | 9.548 | 1.325 | 0.013 |
| 202281_at | GAK | 9.448 | 8.298 | 1.322 | 0.018 |
| 218844_at | FLJ20920 | 9.101 | 7.955 | 1.315 | 0.028 |
| 214016_s_at | SFPQ | 11.130 | 9.988 | 1.304 | 0.031 |
| 219632_s_at | TRPV1 | 6.320 | 5.179 | 1.303 | 0.046 |
| 215898_at | TTLL5 | 7.640 | 6.499 | 1.302 | 0.034 |
| 213136_at | PTPN2 | 11.179 | 10.039 | 1.299 | 0.015 |
| 209561_at | THBS3 | 7.926 | 6.788 | 1.296 | 0.002 |
| 218900_at | CNNM4 | 7.079 | 5.943 | 1.291 | 0.015 |
| 220720_x_at | FLJ14346 | 8.284 | 7.148 | 1.290 | 0.004 |
| 222111_at | --- | 8.769 | 7.634 | 1.290 | 0.043 |
| 218536_at | MRS2L | 8.475 | 7.339 | 1.289 | 0.037 |
| 205255_x_at | TCF7 | 9.265 | 8.130 | 1.288 | 0.027 |
| 208137_x_at | ZNF611 | 8.172 | 7.041 | 1.279 | 0.007 |
| 213672_at | MARS | 7.923 | 6.795 | 1.274 | 0.039 |
| 201099_at | USP9X | 10.206 | 9.077 | 1.273 | 0.004 |
| 210679_x_at | --- | 9.350 | 8.223 | 1.271 | 0.008 |
| 214902_x_at | LPP | 9.096 | 7.970 | 1.268 | 0.001 |
| 213269_at | ZNF248 | 8.469 | 7.343 | 1.267 | 0.013 |
| 202032_s_at | MAN2A2 | 8.642 | 7.518 | 1.263 | 0.041 |
| 221971_x_at | CTGLF1 /// RP11-144G6.7 /// LOC399761 /// LOC653468 | 10.334 | 9.211 | 1.263 | 0.034 |
| 200871_s_at | PSAP | 12.889 | 11.767 | 1.259 | 0.012 |
| 209831_x_at | DNASE2 | 9.443 | 8.321 | 1.258 | 0.040 |
| 213204_at | PARC | 7.290 | 6.168 | 1.257 | 0.005 |
| 218543_s_at | PARP12 | 8.852 | 7.731 | 1.257 | 0.000 |
| 204861_s_at | BIRC1 /// LOC653371 | 7.004 | 5.884 | 1.254 | 0.029 |
| 213927_at | MAP3K9 | 7.988 | 6.869 | 1.252 | 0.023 |
| 37254_at | ZNF133 | 7.424 | 6.305 | 1.250 | 0.042 |
| 206169_x_at | ZC3H7B | 8.308 | 7.190 | 1.250 | 0.045 |
| 204873_at | PEX1 | 7.212 | 6.095 | 1.249 | 0.048 |

| | | | | | |
|---|---|---|---|---|---|
| 210210_at | MPZL1 | 7.505 | 6.389 | 1.246 | 0.021 |
| 215600_x_at | FBXW12 | 8.809 | 7.695 | 1.241 | 0.009 |
| 208165_s_at | PRSS16 | 6.073 | 4.961 | 1.237 | 0.000 |
| 210686_x_at | SLC25A16 | 10.695 | 9.584 | 1.234 | 0.009 |
| 209215_at | TETRAN | 9.066 | 7.955 | 1.234 | 0.005 |
| 202743_at | PIK3R3 | 6.987 | 5.877 | 1.231 | 0.037 |
| 202455_at | HDAC5 | 7.848 | 6.742 | 1.223 | 0.031 |
| 209454_s_at | TEAD3 | 6.583 | 5.478 | 1.221 | 0.004 |
| 221876_at | DKFZp762P2111 | 7.087 | 5.986 | 1.213 | 0.036 |
| 205370_x_at | DBT | 11.729 | 10.628 | 1.212 | 0.002 |
| 219351_at | TRAPPC2 | 7.634 | 6.534 | 1.210 | 0.049 |
| 220948_s_at | ATP1A1 | 11.713 | 10.613 | 1.209 | 0.037 |
| 205926_at | IL27RA | 7.266 | 6.168 | 1.205 | 0.049 |
| 204523_at | ZNF140 | 7.744 | 6.646 | 1.205 | 0.003 |
| 202127_at | PRPF4B | 10.067 | 8.969 | 1.204 | 0.007 |
| 213279_at | DHRS1 | 9.188 | 8.092 | 1.202 | 0.035 |
| 212991_at | FBXO9 | 7.468 | 6.373 | 1.199 | 0.016 |
| 214035_x_at | LOC399491 | 11.465 | 10.383 | 1.171 | 0.001 |
| 208238_x_at | --- | 7.221 | 6.141 | 1.166 | 0.004 |
| 203752_s_at | JUND | 10.942 | 9.865 | 1.159 | 0.038 |
| 207700_s_at | NCOA3 | 9.404 | 8.328 | 1.158 | 0.011 |
| 220030_at | STYK1 | 6.346 | 5.271 | 1.156 | 0.016 |
| 204573_at | CROT | 8.017 | 6.942 | 1.155 | 0.004 |
| 201206_s_at | RRBP1 | 9.423 | 8.350 | 1.152 | 0.038 |
| 207809_s_at | ATP6AP1 | 10.843 | 9.770 | 1.150 | 0.040 |
| 219481_at | TTC13 | 9.320 | 8.249 | 1.148 | 0.024 |
| 203882_at | ISGF3G | 8.041 | 6.971 | 1.146 | 0.050 |
| 209916_at | DHTKD1 | 8.541 | 7.472 | 1.142 | 0.009 |
| 200636_s_at | PTPRF | 11.644 | 10.576 | 1.141 | 0.043 |
| 209061_at | NCOA3 | 8.448 | 7.383 | 1.134 | 0.003 |
| 210910_s_at | ZP3 /// POMZP3 | 6.598 | 5.533 | 1.134 | 0.019 |
| 202057_at | KPNA1 | 7.478 | 6.413 | 1.134 | 0.017 |
| 212640_at | PTPLB | 10.973 | 9.911 | 1.128 | 0.023 |
| 222310_at | SFRS15 | 7.347 | 6.285 | 1.128 | 0.009 |
| 207186_s_at | FALZ | 10.576 | 9.515 | 1.126 | 0.001 |
| 40225_at | GAK | 10.419 | 9.358 | 1.125 | 0.023 |
| 222358_x_at | GLT28D1 | 8.994 | 7.935 | 1.123 | 0.015 |
| 216350_s_at | ZNF10 | 6.546 | 5.487 | 1.121 | 0.010 |
| 55583_at | DOCK6 | 7.642 | 6.584 | 1.120 | 0.013 |
| 215828_at | ATBF1 | 7.219 | 6.165 | 1.110 | 0.006 |
| 201536_at | DUSP3 | 9.738 | 8.685 | 1.108 | 0.024 |
| 206551_x_at | KLHL24 | 8.769 | 7.720 | 1.101 | 0.006 |
| 213485_s_at | ABCC10 | 8.805 | 7.757 | 1.099 | 0.035 |
| 207598_x_at | XRCC2 | 8.645 | 7.598 | 1.095 | 0.035 |
| 220661_s_at | ZNF692 | 9.382 | 8.338 | 1.089 | 0.022 |

| | | | | | |
|---|---|---|---|---|---|
| 210943_s_at | LYST | 6.770 | 5.727 | 1.087 | 0.012 |
| 206792_x_at | PDE4C | 10.306 | 9.264 | 1.086 | 0.025 |
| 214198_s_at | DGCR2 | 7.374 | 6.334 | 1.083 | 0.016 |
| 208082_x_at | --- | 9.479 | 8.439 | 1.080 | 0.003 |
| 217593_at | ZNF447 | 6.523 | 5.484 | 1.079 | 0.006 |
| 202642_s_at | TRRAP | 9.822 | 8.784 | 1.079 | 0.038 |
| 215179_x_at | PGF | 9.957 | 8.921 | 1.073 | 0.014 |
| 206109_at | FUT1 | 6.868 | 5.832 | 1.073 | 0.013 |
| 213502_x_at | LOC91316 | 6.837 | 5.804 | 1.067 | 0.017 |
| 213242_x_at | KIAA0284 | 7.898 | 6.870 | 1.057 | 0.030 |
| 201079_at | SYNGR2 | 9.636 | 8.609 | 1.054 | 0.039 |
| 214870_x_at | NPIP /// LOC23117 /// LOC339047 /// LOC440341 /// LOC642778 /// LOC642799 | 11.485 | 10.459 | 1.053 | 0.000 |
| 211105_s_at | NFATC1 | 6.303 | 5.277 | 1.053 | 0.018 |
| 210975_x_at | FASTK | 8.640 | 7.615 | 1.050 | 0.030 |
| 204267_x_at | PKMYT1 | 8.335 | 7.310 | 1.050 | 0.030 |
| 209366_x_at | CYB5A | 9.940 | 8.918 | 1.046 | 0.009 |
| 210336_x_at | ZNF42 | 8.865 | 7.843 | 1.044 | 0.042 |
| 220243_at | BTBD15 | 6.950 | 5.929 | 1.043 | 0.022 |
| 204040_at | RNF144 | 6.890 | 5.869 | 1.041 | 0.009 |
| 222200_s_at | BSDC1 | 8.458 | 7.438 | 1.040 | 0.037 |
| 213313_at | RABGAP1 | 10.323 | 9.303 | 1.039 | 0.013 |
| 213077_at | YTHDC2 | 10.009 | 8.990 | 1.037 | 0.019 |
| 213694_at | RSBN1 | 8.572 | 7.555 | 1.034 | 0.042 |
| 217141_at | BTBD7 | 7.090 | 6.074 | 1.034 | 0.015 |
| 215577_at | UBE2E1 | 7.883 | 6.866 | 1.033 | 0.048 |
| 201987_at | THRAP1 | 10.659 | 9.648 | 1.022 | 0.031 |
| 207843_x_at | CYB5A | 9.443 | 8.432 | 1.022 | 0.013 |
| 212704_at | ZCCHC11 | 9.424 | 8.416 | 1.017 | 0.002 |
| 203669_s_at | DGAT1 /// LOC642255 | 7.941 | 6.933 | 1.017 | 0.001 |
| 217526_at | NFATC2IP | 9.643 | 8.635 | 1.016 | 0.020 |
| 219504_s_at | C1orf82 | 7.034 | 6.026 | 1.016 | 0.004 |
| 215628_x_at | PPP2CA | 7.558 | 6.550 | 1.016 | 0.029 |
| 214109_at | LRBA | 8.459 | 7.452 | 1.015 | 0.005 |
| 212686_at | PPM1H | 6.875 | 5.867 | 1.015 | 0.038 |
| 55616_at | PERLD1 | 6.739 | 5.733 | 1.013 | 0.046 |
| 215398_at | CDC73 | 7.569 | 6.564 | 1.011 | 0.015 |
| 221230_s_at | ARID4B | 9.758 | 8.754 | 1.008 | 0.000 |
| 205442_at | MFAP3L | 6.089 | 5.090 | 0.998 | 0.038 |
| 201100_s_at | USP9X | 11.693 | 10.694 | 0.997 | 0.012 |
| 202193_at | LIMK2 | 7.771 | 6.774 | 0.994 | 0.017 |
| 203739_at | ZNF217 | 10.170 | 9.175 | 0.991 | 0.034 |

| | | | | | |
|---|---|---|---|---|---|
| 201904_s_at | CTDSPL | 8.494 | 7.500 | 0.988 | 0.040 |
| 201957_at | PPP1R12B | 6.754 | 5.762 | 0.983 | 0.044 |
| 215726_s_at | CYB5A | 9.287 | 8.297 | 0.980 | 0.013 |
| 215528_at | MGAT5 | 8.451 | 7.461 | 0.980 | 0.020 |
| 221501_x_at | LOC339047 | 11.648 | 10.660 | 0.977 | 0.000 |
| 220113_x_at | POLR1B | 8.343 | 7.358 | 0.970 | 0.000 |
| 48531_at | TNIP2 | 9.184 | 8.200 | 0.968 | 0.014 |
| 202804_at | ABCC1 | 11.100 | 10.117 | 0.966 | 0.028 |
| 215529_x_at | DIP2A | 9.388 | 8.405 | 0.966 | 0.027 |
| 215595_x_at | GCNT2 | 6.925 | 5.944 | 0.964 | 0.027 |
| 32259_at | EZH1 | 8.321 | 7.339 | 0.964 | 0.030 |
| 221875_x_at | HLA-F | 8.996 | 8.015 | 0.963 | 0.032 |
| 209195_s_at | ADCY6 | 7.796 | 6.815 | 0.961 | 0.042 |
| 203487_s_at | ARMC8 | 8.347 | 7.367 | 0.961 | 0.028 |
| 214989_x_at | PLEKHA5 | 7.913 | 6.933 | 0.960 | 0.035 |
| 204028_s_at | RABGAP1 | 10.334 | 9.355 | 0.958 | 0.008 |
| 209705_at | --- | 9.171 | 8.193 | 0.956 | 0.014 |
| 207115_x_at | MBTD1 | 7.220 | 6.245 | 0.950 | 0.012 |
| 215281_x_at | POGZ | 8.778 | 7.804 | 0.950 | 0.012 |
| 201825_s_at | SCCPDH | 10.261 | 9.288 | 0.947 | 0.037 |
| 221850_x_at | CTGLF1 | 10.057 | 9.084 | 0.947 | 0.043 |
| 213185_at | KIAA0556 | 7.631 | 6.659 | 0.944 | 0.012 |
| 207986_x_at | CYB561 | 8.672 | 7.700 | 0.944 | 0.005 |
| 212847_at | FUBP1 | 8.718 | 7.748 | 0.941 | 0.008 |
| 202390_s_at | HD | 7.140 | 6.170 | 0.940 | 0.001 |
| 219510_at | POLQ | 9.241 | 8.272 | 0.939 | 0.016 |
| 219980_at | FLJ21106 | 7.113 | 6.146 | 0.935 | 0.002 |
| 203651_at | ZFYVE16 | 9.256 | 8.290 | 0.933 | 0.001 |
| 203577_at | GTF2H4 | 7.696 | 6.731 | 0.931 | 0.027 |
| 217588_at | CATSPER2 /// LOC440278 | 6.797 | 5.836 | 0.924 | 0.030 |
| 221802_s_at | KIAA1598 | 10.109 | 9.148 | 0.923 | 0.015 |
| 211799_x_at | HLA-C | 8.327 | 7.368 | 0.919 | 0.009 |
| 222006_at | LETM1 | 8.220 | 7.264 | 0.913 | 0.007 |
| 204819_at | FGD1 | 7.565 | 6.614 | 0.905 | 0.049 |
| 212356_at | KIAA0323 | 8.491 | 7.540 | 0.904 | 0.050 |
| 216147_at | 11-Sep | 6.896 | 5.947 | 0.900 | 0.015 |
| 204538_x_at | NPIP | 11.505 | 10.559 | 0.894 | 0.000 |
| 212176_at | C6orf111 | 10.409 | 9.466 | 0.889 | 0.040 |
| 47608_at | TJAP1 | 9.083 | 8.142 | 0.887 | 0.022 |
| 211386_at | MGC12488 | 6.458 | 5.517 | 0.886 | 0.025 |
| 215587_x_at | BTBD14B | 8.729 | 7.789 | 0.884 | 0.044 |
| 215791_at | ITSN1 | 7.307 | 6.369 | 0.879 | 0.011 |
| 208723_at | USP11 | 10.115 | 9.178 | 0.877 | 0.019 |
| 39650_s_at | PCNXL2 | 7.405 | 6.470 | 0.876 | 0.045 |

| | | | | | |
|---|---|---|---|---|---|
| 218524_at | E4F1 | 8.475 | 7.540 | 0.873 | 0.001 |
| 31837_at | TMEM153 | 9.459 | 8.525 | 0.872 | 0.011 |
| 220712_at | C8orf60 | 6.702 | 5.768 | 0.872 | 0.038 |
| 213367_at | LOC155060 | 6.672 | 5.739 | 0.870 | 0.020 |
| 215032_at | RREB1 | 7.303 | 6.370 | 0.870 | 0.013 |
| 218155_x_at | TSR1 | 8.476 | 7.545 | 0.867 | 0.005 |
| 209275_s_at | CLN3 | 8.705 | 7.777 | 0.861 | 0.001 |
| 218079_s_at | ZNF403 | 10.175 | 9.249 | 0.859 | 0.004 |
| 216310_at | TAOK1 | 7.016 | 6.094 | 0.850 | 0.040 |
| 209352_s_at | SIN3B | 7.805 | 6.887 | 0.843 | 0.027 |
| 221794_at | DOCK6 | 7.626 | 6.713 | 0.834 | 0.018 |
| 222366_at | ADNP | 7.749 | 6.837 | 0.830 | 0.017 |
| 216229_x_at | HCG2P7 | 6.721 | 5.811 | 0.829 | 0.001 |
| 220659_s_at | FLJ10925 | 7.246 | 6.338 | 0.823 | 0.008 |
| 219722_s_at | GDPD3 | 6.553 | 5.647 | 0.820 | 0.046 |
| 220221_at | VPS13D /// LOC654256 | 6.486 | 5.580 | 0.819 | 0.017 |
| 46256_at | SPSB3 | 9.433 | 8.528 | 0.819 | 0.012 |
| 210582_s_at | LIMK2 | 7.481 | 6.577 | 0.817 | 0.009 |
| 212027_at | RBM25 | 10.486 | 9.583 | 0.817 | 0.011 |
| 214911_s_at | BRD2 | 11.150 | 10.246 | 0.817 | 0.021 |
| 220352_x_at | --- | 7.504 | 6.602 | 0.813 | 0.008 |
| 222361_at | LOC643224 | 6.110 | 5.213 | 0.806 | 0.040 |
| 220071_x_at | CEP27 | 9.240 | 8.348 | 0.797 | 0.000 |
| 200678_x_at | GRN | 10.837 | 9.948 | 0.789 | 0.038 |
| 215553_x_at | WDR45 | 6.583 | 5.696 | 0.788 | 0.022 |
| 220370_s_at | USP36 | 7.797 | 6.910 | 0.788 | 0.012 |
| 221696_s_at | STYK1 | 6.261 | 5.375 | 0.785 | 0.008 |
| 218099_at | TEX2 | 8.307 | 7.425 | 0.779 | 0.031 |
| 216123_x_at | ST7L | 6.617 | 5.735 | 0.778 | 0.002 |
| 211284_s_at | GRN | 10.076 | 9.195 | 0.776 | 0.027 |
| 219906_at | FLJ10213 | 7.457 | 6.577 | 0.774 | 0.023 |
| 207090_x_at | ZFP30 | 6.887 | 6.008 | 0.773 | 0.013 |
| 204202_at | IQCE | 6.373 | 5.494 | 0.772 | 0.023 |
| 208829_at | TAPBP | 10.078 | 9.203 | 0.766 | 0.010 |
| 217527_s_at | NFATC2IP | 9.604 | 8.729 | 0.766 | 0.013 |
| 37652_at | CABIN1 | 7.874 | 7.001 | 0.762 | 0.003 |
| 208611_s_at | SPTAN1 | 11.139 | 10.267 | 0.761 | 0.010 |
| 215754_at | SCARB2 | 6.645 | 5.774 | 0.759 | 0.001 |
| 219757_s_at | C14orf101 | 8.737 | 7.866 | 0.759 | 0.041 |
| 214055_x_at | BAT2D1 | 11.493 | 10.622 | 0.757 | 0.016 |
| 202560_s_at | C1orf77 | 10.389 | 9.524 | 0.750 | 0.025 |
| 218874_s_at | C6orf134 | 6.431 | 5.565 | 0.749 | 0.032 |
| 214093_s_at | FUBP1 | 10.365 | 9.500 | 0.748 | 0.012 |
| 210539_at | TTLL5 | 6.646 | 5.781 | 0.748 | 0.016 |
| 214742_at | AZI1 | 7.205 | 6.341 | 0.746 | 0.004 |

| | | | | | |
|---|---|---|---|---|---|
| 211454_x_at | FKSG49 | 10.666 | 9.802 | 0.745 | 0.033 |
| 216041_x_at | GRN | 10.420 | 9.557 | 0.744 | 0.043 |
| 203727_at | SKIV2L | 8.515 | 7.655 | 0.739 | 0.028 |
| 46665_at | SEMA4C | 9.666 | 8.806 | 0.738 | 0.002 |
| 212760_at | UBR2 | 9.614 | 8.755 | 0.738 | 0.000 |
| 220744_s_at | IFT122 | 7.944 | 7.087 | 0.735 | 0.001 |
| 215383_x_at | SPG21 | 8.113 | 7.258 | 0.732 | 0.011 |
| 212375_at | EP400 | 9.898 | 9.042 | 0.732 | 0.000 |
| 207078_at | MED6 | 7.202 | 6.347 | 0.730 | 0.028 |
| 217939_s_at | AFTIPHILIN | 10.005 | 9.156 | 0.721 | 0.036 |
| 212252_at | CAMKK2 | 7.975 | 7.129 | 0.717 | 0.004 |
| 216524_x_at | ROBO2 | 9.721 | 8.875 | 0.716 | 0.012 |
| 219639_x_at | PARP6 | 9.572 | 8.726 | 0.716 | 0.019 |
| 220335_x_at | FLJ21736 | 6.489 | 5.644 | 0.713 | 0.002 |
| 215373_x_at | FLJ12151 | 8.460 | 7.616 | 0.712 | 0.043 |
| 202500_at | DNAJB2 | 6.342 | 5.501 | 0.707 | 0.044 |
| 209250_at | DEGS1 | 11.023 | 10.182 | 0.706 | 0.050 |
| 214339_s_at | MAP4K1 | 6.034 | 5.194 | 0.705 | 0.047 |
| 216527_at | HCG18 | 6.139 | 5.302 | 0.701 | 0.001 |
| 214340_at | ALOX12P2 | 6.060 | 5.223 | 0.701 | 0.047 |
| 222152_at | PDCD6 | 7.976 | 7.140 | 0.699 | 0.045 |
| 201449_at | TIA1 | 8.886 | 8.050 | 0.698 | 0.031 |
| 210859_x_at | CLN3 | 7.441 | 6.606 | 0.698 | 0.007 |
| 216745_x_at | --- | 7.450 | 6.615 | 0.698 | 0.002 |
| 200931_s_at | VCL | 13.011 | 12.177 | 0.696 | 0.000 |
| 212485_at | KIAA0553 | 9.115 | 8.280 | 0.696 | 0.000 |
| 213743_at | CCNT2 | 9.037 | 8.204 | 0.694 | 0.005 |
| 208120_x_at | FKSG49 | 10.415 | 9.582 | 0.694 | 0.048 |
| 208610_s_at | SRRM2 | 11.270 | 10.439 | 0.690 | 0.000 |
| 204558_at | RAD54L | 8.993 | 8.163 | 0.688 | 0.021 |
| 202616_s_at | MECP2 | 8.625 | 7.801 | 0.680 | 0.038 |
| 203693_s_at | E2F3 | 9.857 | 9.033 | 0.680 | 0.010 |
| 219290_x_at | DAPP1 | 5.748 | 4.924 | 0.679 | 0.015 |
| 207625_s_at | CBFA2T2 | 8.257 | 7.433 | 0.678 | 0.003 |
| 209871_s_at | APBA2 | 6.034 | 5.210 | 0.678 | 0.021 |
| 217849_s_at | CDC42BPB | 8.665 | 7.842 | 0.677 | 0.043 |
| 218335_x_at | TNIP2 | 9.076 | 8.254 | 0.676 | 0.001 |
| 221139_s_at | CSAD | 7.763 | 6.941 | 0.676 | 0.046 |
| 212968_at | RFNG | 7.895 | 7.078 | 0.669 | 0.042 |
| 217164_at | --- | 8.408 | 7.592 | 0.666 | 0.025 |
| 208099_x_at | TTLL5 | 6.938 | 6.123 | 0.663 | 0.001 |
| 217922_at | --- | 9.947 | 9.133 | 0.662 | 0.024 |
| 212403_at | UBE3B | 8.403 | 7.593 | 0.656 | 0.001 |
| 215213_at | NUP54 | 6.056 | 5.246 | 0.655 | 0.019 |
| 213639_s_at | ZNF500 | 7.552 | 6.743 | 0.654 | 0.035 |

| | | | | | |
|---|---|---|---|---|---|
| 202071_at | SDC4 | 10.822 | 10.015 | 0.652 | 0.040 |
| 202611_s_at | CRSP2 | 8.367 | 7.560 | 0.651 | 0.006 |
| 205323_s_at | MTF1 | 6.814 | 6.008 | 0.650 | 0.042 |
| 203756_at | ARHGEF17 | 8.343 | 7.538 | 0.648 | 0.024 |
| 212571_at | CHD8 | 9.527 | 8.722 | 0.647 | 0.000 |
| 211342_x_at | MED12 | 8.676 | 7.872 | 0.647 | 0.043 |
| 216071_x_at | MED12 | 8.314 | 7.511 | 0.644 | 0.017 |
| 203704_s_at | --- | 10.118 | 9.316 | 0.644 | 0.004 |
| 206374_at | DUSP8 | 5.921 | 5.119 | 0.644 | 0.012 |
| 215404_x_at | FGFR1 | 7.244 | 6.442 | 0.643 | 0.011 |
| 203255_at | FBXO11 | 10.609 | 9.808 | 0.642 | 0.005 |
| 221812_at | FBXO42 | 7.338 | 6.538 | 0.641 | 0.002 |
| 220910_at | FRAS1 | 6.224 | 5.424 | 0.640 | 0.039 |
| 213403_at | --- | 5.997 | 5.197 | 0.640 | 0.012 |
| 204739_at | CENPC1 | 7.816 | 7.018 | 0.637 | 0.047 |
| 212179_at | C6orf111 | 10.120 | 9.323 | 0.636 | 0.019 |
| 222207_x_at | LOC441258 /// LOC442580 /// LOC641771 /// LOC641776 /// LOC643862 /// LOC643920 | 10.034 | 9.238 | 0.633 | 0.015 |
| 220079_s_at | USP48 | 10.494 | 9.699 | 0.632 | 0.043 |
| 209290_s_at | NFIB | 10.554 | 9.760 | 0.630 | 0.041 |
| 204568_at | KIAA0831 | 8.038 | 7.246 | 0.628 | 0.004 |
| 201703_s_at | PPP1R10 | 8.807 | 8.020 | 0.619 | 0.042 |
| 216859_x_at | --- | 6.234 | 5.451 | 0.614 | 0.001 |
| 203944_x_at | BTN2A1 | 9.087 | 8.304 | 0.613 | 0.016 |
| 216161_at | SBNO1 | 5.791 | 5.014 | 0.604 | 0.001 |
| 200866_s_at | PSAP | 10.731 | 9.954 | 0.603 | 0.006 |
| 212487_at | KIAA0553 | 9.016 | 8.240 | 0.603 | 0.000 |
| 209054_s_at | WHSC1 | 10.206 | 9.432 | 0.599 | 0.034 |
| 200748_s_at | FTH1 | 13.412 | 12.639 | 0.598 | 0.028 |
| 207435_s_at | SRRM2 | 9.619 | 8.848 | 0.595 | 0.009 |
| 212139_at | GCN1L1 | 10.431 | 9.661 | 0.593 | 0.009 |
| 208663_s_at | TTC3 | 10.437 | 9.668 | 0.591 | 0.026 |
| 222149_x_at | GOLGA8G /// GOLGA8D /// GOLGA8E /// GOLGA8C /// GOLGA8F | 6.872 | 6.105 | 0.589 | 0.000 |
| 202860_at | DENND4B | 9.478 | 8.711 | 0.588 | 0.027 |
| 206431_x_at | KIAA0676 | 7.531 | 6.765 | 0.586 | 0.002 |
| 212323_s_at | VPS13D | 8.320 | 7.555 | 0.586 | 0.023 |
| 215017_s_at | FNBP1L | 9.316 | 8.551 | 0.585 | 0.033 |
| 215583_at | TMEM63A | 6.810 | 6.046 | 0.584 | 0.023 |
| 200058_s_at | ASCC3L1 | 12.310 | 11.546 | 0.584 | 0.027 |
| 222311_s_at | SFRS15 | 8.192 | 7.430 | 0.580 | 0.002 |

| | | | | | |
|---|---|---|---|---|---|
| 217713_x_at | DKFZP566N034 | 6.898 | 6.136 | 0.580 | 0.002 |
| 208823_s_at | PCTK1 | 7.099 | 6.341 | 0.575 | 0.032 |
| 209246_at | ABCF2 | 7.869 | 7.115 | 0.569 | 0.045 |
| 200757_s_at | CALU | 10.917 | 10.163 | 0.568 | 0.026 |
| 34260_at | KIAA0683 | 7.494 | 6.741 | 0.567 | 0.013 |
| 210094_s_at | PARD3 | 10.011 | 9.258 | 0.567 | 0.049 |
| 212383_at | ATP6V0A1 | 8.812 | 8.063 | 0.562 | 0.035 |
| 212293_at | HIPK1 | 9.472 | 8.722 | 0.562 | 0.022 |
| 220739_s_at | CNNM3 | 8.442 | 7.692 | 0.561 | 0.018 |
| 200862_at | DHCR24 | 11.622 | 10.874 | 0.560 | 0.020 |
| 215208_x_at | RPL35A | 8.569 | 7.821 | 0.559 | 0.046 |
| 205486_at | TESK2 | 6.143 | 5.396 | 0.558 | 0.000 |
| 210981_s_at | GRK6 | 7.408 | 6.661 | 0.558 | 0.023 |
| 210251_s_at | RUFY3 | 6.763 | 6.016 | 0.558 | 0.009 |
| 222252_x_at | UBQLN4 /// UBQLN4P | 8.204 | 7.457 | 0.558 | 0.002 |
| 218419_s_at | MGC3123 | 7.116 | 6.369 | 0.558 | 0.023 |
| 219854_at | ZNF14 | 6.533 | 5.788 | 0.555 | 0.032 |
| 41113_at | ZNF500 | 6.421 | 5.676 | 0.554 | 0.012 |
| 202197_at | MTMR3 | 7.248 | 6.505 | 0.552 | 0.005 |
| 218478_s_at | ZCCHC8 | 10.662 | 9.919 | 0.551 | 0.001 |
| 207006_s_at | CCDC106 | 6.744 | 6.003 | 0.549 | 0.020 |
| 208685_x_at | BRD2 | 11.272 | 10.531 | 0.549 | 0.021 |
| 208021_s_at | RFC1 | 9.380 | 8.640 | 0.548 | 0.031 |
| 220734_s_at | MGC10334 | 7.918 | 7.179 | 0.545 | 0.047 |
| 211464_x_at | CASP6 | 8.155 | 7.418 | 0.543 | 0.034 |
| 202256_at | CD2BP2 | 7.896 | 7.159 | 0.543 | 0.007 |
| 215351_at | RTCD1 | 5.960 | 5.223 | 0.542 | 0.021 |
| 208835_s_at | CROP | 11.726 | 10.993 | 0.537 | 0.013 |
| 202250_s_at | WDR42A | 8.331 | 7.600 | 0.534 | 0.033 |
| 215994_x_at | KIAA0676 | 8.321 | 7.591 | 0.533 | 0.001 |
| 216284_at | --- | 6.871 | 6.142 | 0.532 | 0.002 |
| 38069_at | CLCN7 | 9.144 | 8.417 | 0.529 | 0.010 |
| 215900_at | C1orf121 | 5.726 | 5.002 | 0.525 | 0.009 |
| 214241_at | NDUFB8 | 7.918 | 7.195 | 0.524 | 0.011 |
| 218308_at | TACC3 | 10.872 | 10.149 | 0.522 | 0.024 |
| 221260_s_at | C12orf22 | 8.024 | 7.302 | 0.521 | 0.024 |
| 204074_s_at | KIAA0562 | 8.065 | 7.344 | 0.521 | 0.047 |
| 202610_s_at | CRSP2 | 9.209 | 8.489 | 0.518 | 0.011 |
| 47069_at | PRR5 | 6.273 | 5.556 | 0.513 | 0.017 |
| 212779_at | KIAA1109 | 9.525 | 8.809 | 0.512 | 0.013 |
| 217665_at | --- | 6.167 | 5.453 | 0.511 | 0.032 |
| 205178_s_at | RBBP6 | 10.194 | 9.482 | 0.507 | 0.044 |
| 201549_x_at | JARID1B | 8.843 | 8.132 | 0.506 | 0.036 |
| 205181_at | ZNF193 | 7.729 | 7.018 | 0.505 | 0.005 |
| 215873_x_at | ABCC10 | 7.631 | 6.920 | 0.505 | 0.029 |

| | | | | | |
|---|---|---|---|---|---|
| 221882_s_at | TMEM8 | 8.171 | 7.460 | 0.505 | 0.012 |
| 212842_x_at | RGPD5 /// RGPD4 /// LOC653086 /// LOC653596 | 10.499 | 9.789 | 0.503 | 0.029 |
| 204947_at | E2F1 | 8.586 | 7.877 | 0.503 | 0.002 |
| 218834_s_at | TMEM132A | 7.734 | 7.026 | 0.502 | 0.026 |
| 203229_s_at | CLK2 | 10.097 | 9.389 | 0.501 | 0.009 |
| 205933_at | SETBP1 | 5.930 | 5.222 | 0.501 | 0.017 |
| 209826_at | EGFL8 /// LOC653870 | 6.958 | 6.253 | 0.497 | 0.013 |
| 215699_x_at | SFI1 | 6.483 | 5.779 | 0.496 | 0.005 |
| 204649_at | TROAP | 8.380 | 7.676 | 0.496 | 0.020 |
| 213647_at | DNA2L | 7.636 | 6.932 | 0.496 | 0.048 |
| 204403_x_at | KIAA0738 | 8.823 | 8.119 | 0.495 | 0.014 |
| 212054_x_at | KIAA0676 | 8.389 | 7.686 | 0.495 | 0.004 |
| 212052_s_at | KIAA0676 | 9.506 | 8.803 | 0.494 | 0.010 |
| 211202_s_at | JARID1B | 9.085 | 8.383 | 0.493 | 0.047 |
| 206056_x_at | SPN | 6.880 | 6.178 | 0.493 | 0.033 |
| 212518_at | PIP5K1C | 7.752 | 7.051 | 0.491 | 0.007 |
| 213350_at | RPS11 | 9.126 | 8.426 | 0.489 | 0.006 |
| 218539_at | FBXO34 | 9.442 | 8.743 | 0.489 | 0.033 |
| 211374_x_at | --- | 7.145 | 6.446 | 0.488 | 0.001 |
| 220215_at | ZNF669 | 6.520 | 5.826 | 0.483 | 0.002 |
| 204060_s_at | PRKX /// PRKY | 7.244 | 6.549 | 0.482 | 0.028 |
| 202152_x_at | USF2 | 8.172 | 7.478 | 0.482 | 0.018 |
| 221939_at | YIPF2 | 8.338 | 7.645 | 0.480 | 0.020 |
| 219815_at | GAL3ST4 | 6.353 | 5.665 | 0.474 | 0.020 |
| 214047_s_at | MBD4 | 9.101 | 8.414 | 0.473 | 0.006 |
| 215848_at | ZNF291 | 6.162 | 5.475 | 0.472 | 0.040 |
| 213908_at | WHDC1L1 /// WHDC1L2 /// LOC643458 /// LOC646456 | 6.020 | 5.333 | 0.472 | 0.031 |
| 221155_x_at | --- | 6.251 | 5.564 | 0.471 | 0.009 |
| 203558_at | CUL7 | 7.700 | 7.014 | 0.470 | 0.028 |
| 203379_at | RPS6KA1 | 7.350 | 6.667 | 0.467 | 0.038 |
| 222104_x_at | GTF2H3 | 10.870 | 10.189 | 0.463 | 0.013 |
| 211628_x_at | FTHP1 | 11.904 | 11.224 | 0.463 | 0.009 |
| 212031_at | RBM25 | 10.659 | 9.980 | 0.461 | 0.027 |
| 38157_at | DOM3Z | 8.037 | 7.358 | 0.461 | 0.013 |
| 219951_s_at | C20orf12 | 5.864 | 5.186 | 0.460 | 0.043 |
| 212153_at | POGZ | 10.033 | 9.355 | 0.459 | 0.007 |
| 200686_s_at | SFRS11 | 12.324 | 11.648 | 0.457 | 0.009 |
| 212778_at | PACS2 | 7.926 | 7.250 | 0.457 | 0.010 |
| 202809_s_at | INTS3 | 10.108 | 9.432 | 0.456 | 0.004 |
| 216113_at | ABI2 | 5.676 | 5.001 | 0.456 | 0.038 |
| 216129_at | ATP9A | 6.659 | 5.986 | 0.454 | 0.050 |
| 217877_s_at | GPBP1L1 | 9.615 | 8.943 | 0.452 | 0.032 |

| | | | | | |
|---|---|---|---|---|---|
| 220905_at | --- | 6.720 | 6.048 | 0.452 | 0.042 |
| 207436_x_at | KIAA0894 | 8.408 | 7.737 | 0.450 | 0.043 |
| 215070_x_at | RABGAP1 | 7.742 | 7.072 | 0.449 | 0.004 |
| 221827_at | C20orf18 | 9.915 | 9.246 | 0.447 | 0.034 |
| 202322_s_at | GGPS1 | 9.644 | 8.977 | 0.445 | 0.009 |
| 203709_at | PHKG2 | 8.213 | 7.547 | 0.444 | 0.006 |
| 209205_s_at | LMO4 | 9.759 | 9.092 | 0.444 | 0.044 |
| 202866_at | DNAJB12 | 8.899 | 8.233 | 0.444 | 0.020 |
| 201975_at | RSN | 10.158 | 9.492 | 0.443 | 0.016 |
| 202700_s_at | TMEM63A | 6.698 | 6.033 | 0.442 | 0.045 |
| 208216_at | DLX4 | 5.960 | 5.296 | 0.441 | 0.040 |
| 217715_x_at | ZNF354A | 5.943 | 5.280 | 0.440 | 0.024 |
| 203907_s_at | IQSEC1 | 7.819 | 7.156 | 0.440 | 0.041 |
| 219096_at | ARMC7 | 7.912 | 7.255 | 0.431 | 0.003 |
| 207133_x_at | ALPK1 | 5.619 | 4.965 | 0.428 | 0.008 |
| 204587_at | SLC25A14 | 8.391 | 7.737 | 0.427 | 0.025 |
| 210714_at | R3HDM1 | 6.147 | 5.494 | 0.426 | 0.022 |
| 202328_s_at | PKD1 | 6.080 | 5.428 | 0.426 | 0.028 |
| 215296_at | CDC42BPA | 6.572 | 5.920 | 0.425 | 0.011 |
| 219676_at | ZNF435 | 6.678 | 6.028 | 0.423 | 0.011 |
| 214002_at | MYL6 | 6.704 | 6.055 | 0.420 | 0.041 |
| 212101_at | KPNA6 | 9.845 | 9.198 | 0.419 | 0.050 |
| 203127_s_at | SPTLC2 | 9.840 | 9.193 | 0.418 | 0.033 |
| 207188_at | CDK3 | 6.868 | 6.222 | 0.418 | 0.046 |
| 203118_at | PCSK7 | 7.895 | 7.249 | 0.417 | 0.018 |
| 221526_x_at | PARD3 | 9.697 | 9.051 | 0.417 | 0.028 |
| 219086_at | C14orf131 | 6.375 | 5.732 | 0.413 | 0.023 |
| 213980_s_at | CTBP1 | 10.442 | 9.799 | 0.413 | 0.045 |
| 203985_at | ZNF212 | 7.949 | 7.307 | 0.412 | 0.047 |
| 207855_s_at | CLCC1 | 8.366 | 7.724 | 0.412 | 0.011 |
| 218490_s_at | ZNF302 | 9.071 | 8.431 | 0.410 | 0.028 |
| 221811_at | PERLD1 | 6.486 | 5.847 | 0.408 | 0.037 |
| 211256_x_at | BTN2A1 | 7.363 | 6.725 | 0.407 | 0.043 |
| 220341_s_at | LOC51149 | 7.298 | 6.662 | 0.405 | 0.002 |
| 215063_x_at | LRRC40 | 7.639 | 7.003 | 0.404 | 0.002 |
| 218151_x_at | GPR172A | 9.080 | 8.446 | 0.402 | 0.025 |
| 205107_s_at | EFNA4 | 7.494 | 6.860 | 0.401 | 0.042 |
| 213526_s_at | F25965 | 8.657 | 8.026 | 0.399 | 0.022 |
| 209934_s_at | ATP2C1 | 9.250 | 8.621 | 0.397 | 0.021 |
| 209497_s_at | RBM4B | 8.904 | 8.275 | 0.396 | 0.033 |
| 211143_x_at | NR4A1 | 6.640 | 6.012 | 0.394 | 0.011 |
| 202519_at | MLXIP | 9.024 | 8.398 | 0.392 | 0.007 |
| 214736_s_at | ADD1 | 9.635 | 9.012 | 0.388 | 0.027 |
| 213190_at | COG7 | 8.022 | 7.400 | 0.387 | 0.037 |
| 216112_at | PKN2 | 6.702 | 6.080 | 0.387 | 0.037 |

| | | | | | |
|---|---|---|---|---|---|
| 218742_at | NARFL | 7.469 | 6.847 | 0.387 | 0.019 |
| 212756_s_at | UBR2 | 8.660 | 8.040 | 0.385 | 0.012 |
| 203804_s_at | CROP | 11.597 | 10.979 | 0.382 | 0.048 |
| 208797_s_at | GOLGA8B | 5.954 | 5.337 | 0.381 | 0.031 |
| 218977_s_at | TRSPAP1 | 8.442 | 7.825 | 0.381 | 0.030 |
| 35671_at | GTF3C1 | 9.510 | 8.894 | 0.380 | 0.028 |
| 205887_x_at | MSH3 | 6.621 | 6.005 | 0.379 | 0.009 |
| 209586_s_at | PRUNE | 8.140 | 7.527 | 0.376 | 0.020 |
| 217579_x_at | ARL6IP2 | 7.539 | 6.926 | 0.375 | 0.010 |
| 202326_at | EHMT2 | 8.847 | 8.237 | 0.372 | 0.044 |
| 214751_at | ZNF468 | 7.815 | 7.206 | 0.371 | 0.043 |
| 203009_at | BCAM | 6.331 | 5.722 | 0.371 | 0.005 |
| 212856_at | DIP | 6.055 | 5.446 | 0.371 | 0.049 |
| 216000_at | KIAA0484 | 5.725 | 5.117 | 0.369 | 0.035 |
| 206471_s_at | PLXNC1 | 5.743 | 5.137 | 0.368 | 0.000 |
| 217828_at | SLTM | 10.436 | 9.830 | 0.368 | 0.034 |
| 218410_s_at | MGC4692 | 6.211 | 5.605 | 0.367 | 0.021 |
| 221250_s_at | MXD3 | 7.396 | 6.792 | 0.364 | 0.001 |
| 222375_at | PPIG | 8.158 | 7.556 | 0.363 | 0.026 |
| 205090_s_at | NAGPA | 8.145 | 7.546 | 0.359 | 0.040 |
| 200599_s_at | HSP90B1 | 12.900 | 12.302 | 0.357 | 0.038 |
| 212786_at | KIAA0350 | 6.928 | 6.331 | 0.356 | 0.039 |
| 212752_at | CLASP1 | 8.852 | 8.256 | 0.355 | 0.020 |
| 201598_s_at | INPPL1 | 7.464 | 6.870 | 0.353 | 0.034 |
| 216756_at | --- | 5.828 | 5.235 | 0.351 | 0.032 |
| 205812_s_at | TMED9 | 12.101 | 11.509 | 0.350 | 0.037 |
| 212542_s_at | PHIP | 10.800 | 10.210 | 0.349 | 0.009 |
| 215467_x_at | LOC647070 | 5.932 | 5.341 | 0.349 | 0.003 |
| 220584_at | FLJ22184 | 5.703 | 5.116 | 0.344 | 0.000 |
| 222169_x_at | SH2D3A | 5.841 | 5.256 | 0.342 | 0.048 |
| 202913_at | ARHGEF11 | 7.100 | 6.516 | 0.342 | 0.041 |
| 201997_s_at | SPEN | 10.018 | 9.434 | 0.340 | 0.020 |
| 209155_s_at | NT5C2 | 9.374 | 8.791 | 0.340 | 0.009 |
| 203368_at | CRELD1 | 7.101 | 6.517 | 0.340 | 0.021 |
| 212207_at | THRAP2 | 6.532 | 5.951 | 0.338 | 0.026 |
| 52975_at | C9orf28 | 6.631 | 6.050 | 0.338 | 0.026 |
| 212380_at | KIAA0082 | 9.136 | 8.556 | 0.337 | 0.023 |
| 202849_x_at | GRK6 | 7.184 | 6.605 | 0.336 | 0.018 |
| 212455_at | YTHDC1 | 11.263 | 10.684 | 0.335 | 0.039 |
| 201696_at | SFRS4 | 10.871 | 10.293 | 0.335 | 0.013 |
| 210132_at | EFNA3 | 6.987 | 6.410 | 0.333 | 0.001 |
| 47773_at | FBXO42 | 7.064 | 6.488 | 0.332 | 0.028 |
| 219392_x_at | PRR11 | 11.108 | 10.534 | 0.330 | 0.034 |
| 215586_at | PPP3CB | 5.787 | 5.213 | 0.329 | 0.038 |
| 221879_at | CALML4 | 7.793 | 7.220 | 0.328 | 0.022 |

| | | | | | |
|---|---|---|---|---|---|
| 213396_s_at | AKAP10 | 8.109 | 7.538 | 0.326 | 0.011 |
| 208845_at | VDAC3 | 12.624 | 12.056 | 0.323 | 0.040 |
| 218231_at | NAGK | 9.257 | 8.691 | 0.321 | 0.047 |
| 210977_s_at | HSF4 | 6.514 | 5.948 | 0.321 | 0.040 |
| 36545_s_at | SFI1 | 6.941 | 6.376 | 0.319 | 0.024 |
| 208809_s_at | C6orf62 | 10.762 | 10.197 | 0.319 | 0.036 |
| 221857_s_at | TJAP1 | 6.593 | 6.031 | 0.315 | 0.016 |
| 202702_at | TRIM26 | 8.037 | 7.476 | 0.314 | 0.038 |
| 206721_at | C1orf114 | 5.561 | 5.000 | 0.314 | 0.023 |
| 220632_s_at | POMT2 | 6.651 | 6.091 | 0.314 | 0.005 |
| 217619_x_at | --- | 6.544 | 5.984 | 0.313 | 0.041 |
| 222047_s_at | ARS2 | 11.853 | 11.295 | 0.312 | 0.023 |
| 218441_s_at | RPAP1 | 7.748 | 7.190 | 0.311 | 0.017 |
| 213936_x_at | SFTPB | 5.921 | 5.364 | 0.310 | 0.012 |
| 215013_s_at | USP34 | 7.467 | 6.911 | 0.308 | 0.020 |
| 208030_s_at | ADD1 | 9.317 | 8.763 | 0.308 | 0.040 |
| 205271_s_at | CCRK | 5.897 | 5.342 | 0.307 | 0.023 |
| 221064_s_at | C16orf28 | 6.734 | 6.181 | 0.306 | 0.024 |
| 210157_at | C19orf2 | 7.679 | 7.128 | 0.304 | 0.030 |
| 219379_x_at | ZNF358 | 6.373 | 5.823 | 0.303 | 0.017 |
| 211040_x_at | GTSE1 | 9.758 | 9.208 | 0.302 | 0.025 |
| 217643_x_at | FLJ22795 | 6.881 | 6.333 | 0.300 | 0.028 |
| 201011_at | RPN1 | 11.353 | 10.805 | 0.300 | 0.022 |
| 217501_at | WDR39 | 9.086 | 8.540 | 0.298 | 0.036 |
| 219786_at | MTL5 | 5.509 | 4.964 | 0.297 | 0.010 |
| 203160_s_at | RNF8 | 8.483 | 7.939 | 0.296 | 0.043 |
| 214354_x_at | SFTPB | 6.178 | 5.634 | 0.296 | 0.023 |
| 212152_x_at | ARID1A | 10.494 | 9.951 | 0.295 | 0.027 |
| 201024_x_at | EIF5B | 12.253 | 11.711 | 0.294 | 0.007 |
| 213081_at | ZBTB22 | 6.159 | 5.617 | 0.294 | 0.003 |
| 208709_s_at | NRD1 | 11.373 | 10.831 | 0.294 | 0.031 |
| 220242_x_at | ZNF701 | 6.099 | 5.562 | 0.288 | 0.018 |
| 212431_at | KIAA0194 | 7.261 | 6.729 | 0.283 | 0.019 |
| 212424_at | PDCD11 | 8.728 | 8.198 | 0.281 | 0.011 |
| 206254_at | EGF | 5.363 | 4.832 | 0.281 | 0.025 |
| 212376_s_at | EP400 | 8.245 | 7.716 | 0.280 | 0.043 |
| 212443_at | NBEAL2 | 6.838 | 6.312 | 0.277 | 0.004 |
| 212863_x_at | CTBP1 | 10.843 | 10.318 | 0.275 | 0.010 |
| 221806_s_at | SETD5 | 10.608 | 10.084 | 0.274 | 0.041 |
| 208469_s_at | EGFL8 /// LOC653870 | 6.163 | 5.640 | 0.273 | 0.015 |
| 215284_at | SNX9 | 5.614 | 5.093 | 0.272 | 0.029 |
| 217594_at | ZCCHC11 | 5.821 | 5.300 | 0.272 | 0.035 |
| 202340_x_at | NR4A1 | 6.195 | 5.675 | 0.270 | 0.000 |
| 219460_s_at | TMEM127 | 8.458 | 7.938 | 0.270 | 0.045 |
| 218509_at | LPPR2 | 6.490 | 5.971 | 0.270 | 0.044 |

| | | | | | |
|---|---|---|---|---|---|
| 213177_at | MAPK8IP3 | 6.126 | 5.609 | 0.267 | 0.013 |
| 213235_at | LOC400506 | 7.893 | 7.377 | 0.266 | 0.008 |
| 220600_at | TMEM103 | 6.125 | 5.610 | 0.265 | 0.006 |
| 213660_s_at | TOP3B | 5.727 | 5.212 | 0.265 | 0.040 |
| 203956_at | MORC2 | 9.641 | 9.128 | 0.263 | 0.016 |
| 218373_at | FTS | 8.540 | 8.029 | 0.261 | 0.034 |
| 203714_s_at | TBCE | 9.855 | 9.346 | 0.260 | 0.036 |
| 206317_s_at | ABCB8 | 6.445 | 5.937 | 0.259 | 0.000 |
| 220111_s_at | TMEM16B | 5.373 | 4.868 | 0.254 | 0.030 |
| 213119_at | SLC36A1 | 7.202 | 6.700 | 0.252 | 0.020 |
| 222046_at | ARS2 | 6.113 | 5.611 | 0.252 | 0.016 |
| 209167_at | GPM6B | 5.433 | 4.933 | 0.251 | 0.027 |
| 213612_x_at | NBPF15 /// NBPF10 /// NBPF16 | 12.223 | 11.724 | 0.249 | 0.013 |
| 209935_at | ATP2C1 | 9.158 | 8.659 | 0.248 | 0.030 |
| 207064_s_at | AOC2 | 5.465 | 4.967 | 0.248 | 0.001 |
| 202360_at | MAML1 | 9.998 | 9.504 | 0.245 | 0.033 |
| 213570_at | EIF4E2 | 5.831 | 5.336 | 0.245 | 0.046 |
| 218765_at | SIDT2 | 6.330 | 5.836 | 0.244 | 0.009 |
| 214760_at | ZNF337 | 8.314 | 7.820 | 0.244 | 0.027 |
| 204524_at | PDPK1 | 9.685 | 9.196 | 0.239 | 0.016 |
| 202775_s_at | SFRS8 | 8.896 | 8.410 | 0.236 | 0.049 |
| 213512_at | C14orf79 | 6.697 | 6.216 | 0.232 | 0.002 |
| 215620_at | RREB1 | 5.394 | 4.913 | 0.231 | 0.005 |
| 219760_at | LIN7B | 6.355 | 5.876 | 0.229 | 0.025 |
| 216642_at | SEC14L1 | 5.634 | 5.156 | 0.229 | 0.032 |
| 221769_at | SPSB3 | 6.469 | 5.991 | 0.229 | 0.016 |
| 218173_s_at | WHSC1L1 | 5.310 | 4.837 | 0.224 | 0.015 |
| 222276_at | --- | 5.959 | 5.487 | 0.223 | 0.025 |
| 210499_s_at | PQBP1 | 6.241 | 5.770 | 0.222 | 0.024 |
| 222304_x_at | OR7E47P | 6.400 | 5.932 | 0.219 | 0.022 |
| 201103_x_at | NBPF11 /// NBPF15 /// NBPF9 /// NBPF10 /// NBPF16 | 12.156 | 11.689 | 0.218 | 0.022 |
| 215597_x_at | MYST4 | 5.777 | 5.310 | 0.218 | 0.015 |
| 91617_at | DGCR8 | 8.111 | 7.645 | 0.217 | 0.046 |
| 210718_s_at | ARL17P1 | 6.165 | 5.700 | 0.217 | 0.035 |
| 220012_at | ERO1LB | 5.806 | 5.341 | 0.216 | 0.022 |
| 215525_at | --- | 5.828 | 5.366 | 0.214 | 0.009 |
| 202692_s_at | UBTF | 7.448 | 6.988 | 0.212 | 0.025 |
| 209170_s_at | GPM6B | 5.065 | 4.607 | 0.210 | 0.011 |
| 211970_x_at | ACTG1 | 14.152 | 13.694 | 0.210 | 0.043 |
| 218506_x_at | N-PAC | 8.700 | 8.244 | 0.208 | 0.003 |
| 201806_s_at | ATXN2L | 7.052 | 6.596 | 0.208 | 0.022 |
| 31861_at | IGHMBP2 | 7.663 | 7.208 | 0.207 | 0.004 |

| | | | | | |
|---|---|---|---|---|---|
| 203506_s_at | MED12 | 8.020 | 7.565 | 0.207 | 0.034 |
| 217446_x_at | --- | 6.353 | 5.901 | 0.205 | 0.049 |
| 214192_at | NUP88 | 5.591 | 5.140 | 0.203 | 0.001 |
| 202051_s_at | ZMYM4 | 9.697 | 9.248 | 0.202 | 0.020 |
| 215636_at | ZUBR1 | 5.978 | 5.529 | 0.202 | 0.043 |
| 202474_s_at | HCFC1 | 9.740 | 9.295 | 0.198 | 0.004 |
| 206813_at | CTF1 | 5.995 | 5.553 | 0.195 | 0.011 |
| 216459_x_at | --- | 5.296 | 4.856 | 0.194 | 0.010 |
| 219039_at | SEMA4C | 6.743 | 6.306 | 0.191 | 0.015 |
| 214404_x_at | SPDEF | 5.890 | 5.457 | 0.188 | 0.046 |
| 214409_at | RFPL3S | 4.837 | 4.410 | 0.182 | 0.006 |
| 204792_s_at | IFT140 | 5.664 | 5.238 | 0.182 | 0.020 |
| 212324_s_at | VPS13D | 6.629 | 6.205 | 0.180 | 0.046 |
| 203082_at | BMS1L | 10.009 | 9.585 | 0.180 | 0.006 |
| 202486_at | AFG3L2 | 8.666 | 8.244 | 0.178 | 0.031 |
| 203694_s_at | DHX16 | 9.644 | 9.226 | 0.175 | 0.027 |
| 220605_s_at | SIRT2 | 8.250 | 7.835 | 0.172 | 0.048 |
| 57082_at | LDLRAP1 | 7.072 | 6.657 | 0.172 | 0.026 |
| 203671_at | TPMT | 6.801 | 6.388 | 0.171 | 0.013 |
| 212854_x_at | NBPF10 | 11.467 | 11.054 | 0.170 | 0.012 |
| 200925_at | COX6A1 | 13.638 | 13.226 | 0.169 | 0.023 |
| 218836_at | RPP21 | 9.883 | 9.471 | 0.169 | 0.024 |
| 219102_at | RCN3 | 5.918 | 5.507 | 0.169 | 0.016 |
| 216768_x_at | FLJ20699 | 6.355 | 5.946 | 0.167 | 0.037 |
| 222090_at | --- | 6.007 | 5.599 | 0.167 | 0.010 |
| 209913_x_at | --- | 6.751 | 6.344 | 0.166 | 0.032 |
| 201216_at | ERP29 | 11.442 | 11.038 | 0.164 | 0.043 |
| 216750_at | APBB2 | 5.685 | 5.282 | 0.163 | 0.015 |
| 214338_at | DNAJB12 | 6.568 | 6.165 | 0.162 | 0.026 |
| 212982_at | ZDHHC17 | 9.286 | 8.886 | 0.160 | 0.007 |
| 216121_at | VPS35 | 5.416 | 5.019 | 0.158 | 0.046 |
| 216765_at | MAP2K5 | 5.527 | 5.134 | 0.154 | 0.047 |
| 211610_at | KLF6 | 5.914 | 5.523 | 0.153 | 0.023 |
| 211459_at | --- | 5.080 | 4.690 | 0.152 | 0.001 |
| 32091_at | KIAA0446 | 9.612 | 9.222 | 0.152 | 0.022 |
| 212683_at | KIAA0446 | 7.101 | 6.712 | 0.152 | 0.015 |
| 219232_s_at | EGLN3 | 5.654 | 5.266 | 0.150 | 0.050 |
| 215326_at | PAK4 | 5.503 | 5.116 | 0.149 | 0.032 |
| 222073_at | COL4A3 | 5.126 | 4.740 | 0.149 | 0.034 |
| 219897_at | RNF122 | 5.892 | 5.507 | 0.148 | 0.033 |
| 212369_at | ZNF384 | 9.205 | 8.822 | 0.147 | 0.020 |
| 212231_at | FBXO21 | 9.593 | 9.210 | 0.147 | 0.025 |
| 208979_at | NCOA6 | 9.947 | 9.565 | 0.145 | 0.023 |
| 214253_s_at | DTNB | 6.006 | 5.627 | 0.144 | 0.008 |
| 206068_s_at | ACADL | 5.116 | 4.737 | 0.144 | 0.022 |

| | | | | | |
|---|---|---|---|---|---|
| 77508_r_at | RABEP2 /// LOC652743 | 6.234 | 5.856 | 0.143 | 0.012 |
| 215402_at | APPBP2 | 5.848 | 5.472 | 0.142 | 0.039 |
| 219316_s_at | C14orf58 | 5.506 | 5.130 | 0.142 | 0.029 |
| 210835_s_at | CTBP2 | 11.369 | 10.993 | 0.142 | 0.022 |
| 216246_at | --- | 8.987 | 8.611 | 0.141 | 0.032 |
| 220779_at | PADI3 | 5.606 | 5.232 | 0.140 | 0.046 |
| 205245_at | PARD6A | 6.222 | 5.849 | 0.140 | 0.040 |
| 211531_x_at | PRB1 /// PRB2 | 5.900 | 5.528 | 0.138 | 0.030 |
| 216735_x_at | HRH1 | 4.983 | 4.612 | 0.138 | 0.008 |
| 220689_at | --- | 5.511 | 5.140 | 0.138 | 0.008 |
| 214861_at | JMJD2C | 6.278 | 5.907 | 0.138 | 0.050 |
| 217734_s_at | WDR6 | 9.984 | 9.614 | 0.137 | 0.028 |
| 205432_at | OVGP1 | 5.951 | 5.582 | 0.136 | 0.005 |
| 212108_at | UBXD8 | 10.702 | 10.335 | 0.135 | 0.014 |
| 204611_s_at | PPP2R5B | 6.989 | 6.623 | 0.133 | 0.031 |
| 203796_s_at | BCL7A | 6.300 | 5.936 | 0.133 | 0.044 |
| 203398_s_at | GALNT3 | 5.051 | 4.689 | 0.132 | 0.033 |
| 220618_s_at | ZCWPW1 | 5.546 | 5.184 | 0.131 | 0.010 |
| 213339_at | KIAA0495 | 5.339 | 4.977 | 0.131 | 0.010 |
| 208820_at | PTK2 | 11.142 | 10.780 | 0.131 | 0.050 |
| 216537_s_at | SIGLEC7 | 5.908 | 5.546 | 0.130 | 0.035 |
| 214982_at | ASCC3L1 /// LOC652147 | 5.083 | 4.722 | 0.130 | 0.001 |
| 219941_at | TMEM19 | 5.610 | 5.250 | 0.130 | 0.017 |
| 219286_s_at | RBM15 | 10.079 | 9.721 | 0.128 | 0.045 |
| 209910_at | SLC25A16 | 5.802 | 5.445 | 0.128 | 0.048 |
| 208558_at | OR10H1 | 5.732 | 5.375 | 0.128 | 0.028 |
| 209169_at | GPM6B | 4.851 | 4.496 | 0.126 | 0.006 |
| 206264_at | GPLD1 | 4.974 | 4.620 | 0.125 | 0.022 |
| 202726_at | LIG1 | 8.859 | 8.507 | 0.124 | 0.032 |
| 213172_at | TTC9 | 5.344 | 4.993 | 0.123 | 0.003 |
| 205156_s_at | ACCN2 | 5.900 | 5.551 | 0.122 | 0.036 |
| 210737_at | TUB | 5.817 | 5.469 | 0.121 | 0.023 |
| 221887_s_at | DFNB31 | 6.856 | 6.509 | 0.120 | 0.027 |
| 213270_at | MPP2 | 6.329 | 5.984 | 0.119 | 0.050 |
| 215845_x_at | --- | 5.577 | 5.234 | 0.118 | 0.003 |
| 215589_at | SMURF1 | 5.314 | 4.971 | 0.118 | 0.027 |
| 206398_s_at | CD19 | 5.276 | 4.933 | 0.117 | 0.020 |
| 215895_x_at | ADFP | 5.137 | 4.795 | 0.117 | 0.031 |
| 219309_at | CTA-216E10.6 | 5.394 | 5.052 | 0.117 | 0.009 |
| 220524_at | LOC653326 | 5.423 | 5.082 | 0.116 | 0.048 |
| 211002_s_at | TRIM29 | 5.504 | 5.164 | 0.116 | 0.046 |
| 211451_s_at | KCNJ4 | 5.327 | 4.987 | 0.116 | 0.001 |
| 209976_s_at | CYP2E1 | 5.389 | 5.050 | 0.115 | 0.036 |
| 217280_x_at | GABRA5 /// LOC653222 | 5.028 | 4.689 | 0.115 | 0.001 |
| 220107_s_at | C14orf140 | 5.408 | 5.071 | 0.114 | 0.009 |

| | | | | | |
|---|---|---|---|---|---|
| 220989_s_at | AMN | 6.173 | 5.838 | 0.113 | 0.032 |
| 216204_at | COMT | 5.766 | 5.431 | 0.112 | 0.005 |
| 215907_at | BACH2 | 6.558 | 6.225 | 0.111 | 0.018 |
| 215554_at | GPLD1 | 5.237 | 4.905 | 0.110 | 0.002 |
| 206709_x_at | GPT | 5.554 | 5.223 | 0.109 | 0.018 |
| 221991_at | NXPH3 | 5.854 | 5.524 | 0.109 | 0.038 |
| 204129_at | BCL9 | 5.855 | 5.526 | 0.108 | 0.036 |
| 50221_at | TFEB | 5.319 | 4.990 | 0.108 | 0.004 |
| 216051_x_at | KIAA1217 | 5.944 | 5.615 | 0.108 | 0.006 |
| 220469_at | COPE | 5.584 | 5.255 | 0.108 | 0.017 |
| 206119_at | BHMT | 5.182 | 4.854 | 0.108 | 0.010 |
| 207252_at | INE1 | 5.469 | 5.142 | 0.107 | 0.041 |
| 205988_at | CD84 | 5.068 | 4.741 | 0.107 | 0.030 |
| 222341_x_at | CCDC14 | 5.299 | 4.972 | 0.107 | 0.016 |
| 32029_at | PDPK1 | 8.073 | 7.747 | 0.106 | 0.003 |
| 217342_x_at | FLJ11292 | 5.400 | 5.076 | 0.105 | 0.008 |
| 207468_s_at | SFRP5 | 5.348 | 5.025 | 0.104 | 0.001 |
| 212137_at | LARP1 | 12.072 | 11.751 | 0.103 | 0.021 |
| 220858_at | SORBS2 | 5.104 | 4.783 | 0.103 | 0.023 |
| 219916_s_at | RNF39 | 5.685 | 5.364 | 0.103 | 0.020 |
| 201085_s_at | SON | 9.731 | 9.412 | 0.102 | 0.023 |
| 213202_at | SETD1A | 6.271 | 5.952 | 0.102 | 0.014 |
| 205998_x_at | CYP3A4 | 5.901 | 5.583 | 0.101 | 0.017 |
| 221022_s_at | PMFBP1 | 5.347 | 5.030 | 0.101 | 0.032 |
| 210365_at | RUNX1 | 5.599 | 5.286 | 0.098 | 0.029 |
| 211424_x_at | METTL7A | 4.879 | 4.570 | 0.096 | 0.028 |
| 220891_at | C4orf23 | 5.040 | 4.732 | 0.095 | 0.006 |
| 213958_at | CD6 | 6.133 | 5.825 | 0.095 | 0.010 |
| 205332_at | RCE1 | 5.590 | 5.283 | 0.094 | 0.024 |
| 217041_at | NPTXR | 5.164 | 4.858 | 0.094 | 0.020 |
| 209171_at | ITPA | 9.445 | 9.141 | 0.093 | 0.029 |
| 216223_at | CPN2 | 4.863 | 4.559 | 0.093 | 0.026 |
| 220377_at | FAM30A | 5.583 | 5.279 | 0.093 | 0.029 |
| 206718_at | LMO1 | 5.172 | 4.871 | 0.091 | 0.038 |
| 203326_x_at | --- | 5.739 | 5.438 | 0.090 | 0.026 |
| 202881_x_at | --- | 5.615 | 5.314 | 0.090 | 0.016 |
| 207312_at | PHKG1 | 5.148 | 4.847 | 0.090 | 0.003 |
| 207370_at | IBSP | 5.154 | 4.854 | 0.090 | 0.003 |
| 207767_s_at | EGR4 | 5.170 | 4.870 | 0.090 | 0.021 |
| 203969_at | LOC153914 | 5.344 | 5.045 | 0.089 | 0.013 |
| 216924_s_at | DRD2 | 5.396 | 5.098 | 0.089 | 0.009 |
| 49327_at | SIRT3 | 7.128 | 6.829 | 0.089 | 0.033 |
| 207185_at | SLC10A1 | 5.315 | 5.018 | 0.088 | 0.003 |
| 221240_s_at | B3GNT4 | 5.685 | 5.388 | 0.088 | 0.027 |
| 210304_at | PDE6B | 5.498 | 5.201 | 0.088 | 0.050 |

| | | | | | |
|---|---|---|---|---|---|
| 214408_s_at | RFPL3S /// RFPL1S | 5.213 | 4.916 | 0.088 | 0.001 |
| 221164_x_at | CHST5 | 5.141 | 4.847 | 0.087 | 0.022 |
| 210459_at | PSMD4 | 5.196 | 4.902 | 0.086 | 0.011 |
| 216362_at | --- | 5.381 | 5.088 | 0.086 | 0.006 |
| 221789_x_at | RHOT2 | 7.468 | 7.175 | 0.086 | 0.020 |
| 217384_x_at | LOC388255 /// LOC440361 /// LOC652159 | 4.886 | 4.594 | 0.085 | 0.009 |
| 216326_s_at | HDAC3 | 8.442 | 8.151 | 0.085 | 0.029 |
| 216290_x_at | DPP6 | 4.942 | 4.651 | 0.085 | 0.021 |
| 208040_s_at | MYBPC3 | 6.014 | 5.723 | 0.085 | 0.021 |
| 207187_at | JAK3 | 5.130 | 4.840 | 0.084 | 0.011 |
| 214987_at | --- | 5.572 | 5.282 | 0.084 | 0.039 |
| 216243_s_at | IL1RN | 5.384 | 5.096 | 0.083 | 0.005 |
| 212700_x_at | PLEKHM1 /// LOC440456 | 5.766 | 5.478 | 0.083 | 0.045 |
| 206490_at | DLGAP1 | 5.028 | 4.740 | 0.083 | 0.039 |
| 204577_s_at | CLUAP1 | 5.496 | 5.214 | 0.080 | 0.026 |
| 208071_s_at | LAIR1 | 5.546 | 5.264 | 0.079 | 0.034 |
| 213433_at | ARL3 | 5.543 | 5.263 | 0.078 | 0.028 |
| 207060_at | EN2 | 5.130 | 4.851 | 0.078 | 0.044 |
| 216910_at | XPNPEP2 | 5.287 | 5.008 | 0.078 | 0.025 |
| 213925_at | C1orf95 | 6.295 | 6.018 | 0.077 | 0.002 |
| 210370_s_at | LY9 | 5.097 | 4.819 | 0.077 | 0.049 |
| 215654_at | BCAT2 | 5.438 | 5.161 | 0.077 | 0.033 |
| 220290_at | AIM1L | 5.479 | 5.202 | 0.077 | 0.020 |
| 221171_at | RP4-692D3.1 | 5.250 | 4.974 | 0.077 | 0.041 |
| 216702_x_at | ATP8A2 | 5.810 | 5.535 | 0.076 | 0.028 |
| 220283_at | KIAA1822L | 5.101 | 4.826 | 0.075 | 0.040 |
| 220364_at | FLJ11235 | 5.164 | 4.890 | 0.075 | 0.023 |
| 206604_at | OVOL1 | 5.256 | 4.982 | 0.075 | 0.024 |
| 216185_at | FUT9 | 5.008 | 4.734 | 0.075 | 0.024 |
| 216844_at | ZC3H7B | 4.921 | 4.648 | 0.075 | 0.002 |
| 222337_at | OSBPL9 | 5.780 | 5.507 | 0.075 | 0.037 |
| 210072_at | CCL19 | 5.083 | 4.810 | 0.074 | 0.032 |
| 207267_s_at | DSCR6 | 5.116 | 4.845 | 0.074 | 0.027 |
| 221541_at | CRISPLD2 | 5.873 | 5.603 | 0.073 | 0.024 |
| 207684_at | TBX6 | 5.323 | 5.054 | 0.073 | 0.005 |
| 205230_at | RPH3A | 5.707 | 5.438 | 0.072 | 0.040 |
| 214357_at | C1orf105 | 5.431 | 5.163 | 0.072 | 0.047 |
| 216670_at | KLK13 | 5.119 | 4.851 | 0.072 | 0.044 |
| 215202_at | LOC91316 | 5.789 | 5.522 | 0.071 | 0.001 |
| 213920_at | CUTL2 | 5.434 | 5.169 | 0.070 | 0.031 |
| 206223_at | LMTK2 | 5.479 | 5.215 | 0.070 | 0.032 |
| 206189_at | UNC5C | 5.036 | 4.773 | 0.069 | 0.007 |
| 222115_x_at | N-PAC | 6.599 | 6.338 | 0.068 | 0.008 |

| | | | | | |
|---|---|---|---|---|---|
| 209983_s_at | NRXN2 | 5.295 | 5.034 | 0.068 | 0.018 |
| 210944_s_at | CAPN3 | 5.199 | 4.938 | 0.068 | 0.015 |
| 220224_at | HAO1 | 5.116 | 4.856 | 0.068 | 0.048 |
| 206950_at | SCN9A | 4.724 | 4.463 | 0.068 | 0.032 |
| 217079_at | LOC646452 | 5.717 | 5.456 | 0.068 | 0.035 |
| 213539_at | CD3D | 5.297 | 5.037 | 0.068 | 0.011 |
| 37953_s_at | ACCN2 | 5.522 | 5.262 | 0.068 | 0.024 |
| 205039_s_at | ZNFN1A1 | 6.001 | 5.743 | 0.067 | 0.023 |
| 214402_s_at | SFI1 | 5.821 | 5.562 | 0.067 | 0.026 |
| 210330_at | SGCD | 4.872 | 4.615 | 0.066 | 0.036 |
| 207998_s_at | CACNA1D | 5.354 | 5.097 | 0.066 | 0.030 |
| 217242_at | ZNF154 | 5.004 | 4.747 | 0.066 | 0.022 |
| 221225_at | DCAKD | 5.528 | 5.272 | 0.066 | 0.013 |
| 204829_s_at | FOLR2 | 5.518 | 5.262 | 0.066 | 0.029 |
| 207609_s_at | CYP1A2 | 5.638 | 5.383 | 0.065 | 0.027 |
| 205119_s_at | FPR1 | 5.453 | 5.199 | 0.065 | 0.013 |
| 204684_at | NPTX1 | 5.466 | 5.212 | 0.064 | 0.015 |
| 211811_s_at | PCDHA6 | 5.162 | 4.909 | 0.064 | 0.019 |
| 222218_s_at | PILRA | 5.268 | 5.015 | 0.064 | 0.020 |
| 216851_at | IGL@ | 5.295 | 5.042 | 0.064 | 0.007 |
| 202312_s_at | COL1A1 | 5.650 | 5.398 | 0.064 | 0.034 |
| 207757_at | ZFP2 | 5.265 | 5.013 | 0.063 | 0.045 |
| 220667_at | --- | 5.314 | 5.062 | 0.063 | 0.020 |
| 210736_x_at | DTNA | 5.684 | 5.434 | 0.062 | 0.021 |
| 207073_at | CDKL2 | 5.179 | 4.930 | 0.062 | 0.028 |
| 215552_s_at | ESR1 | 5.052 | 4.804 | 0.062 | 0.047 |
| 205338_s_at | DCT | 5.031 | 4.783 | 0.062 | 0.009 |
| 222091_at | --- | 5.647 | 5.399 | 0.061 | 0.040 |
| 208279_s_at | CDRT1 | 5.128 | 4.881 | 0.061 | 0.045 |
| 205484_at | SIT1 | 5.498 | 5.251 | 0.061 | 0.015 |
| 208520_at | OR10H3 | 4.910 | 4.664 | 0.061 | 0.003 |
| 210660_at | LILRA1 | 5.347 | 5.101 | 0.061 | 0.004 |
| 205987_at | CD1C | 5.113 | 4.868 | 0.060 | 0.030 |
| 38447_at | ADRBK1 | 5.749 | 5.505 | 0.060 | 0.017 |
| 210800_at | --- | 4.800 | 4.556 | 0.060 | 0.009 |
| 207144_s_at | CITED1 | 5.517 | 5.273 | 0.059 | 0.009 |
| 207084_at | POU3F2 | 5.782 | 5.538 | 0.059 | 0.045 |
| 211135_x_at | LILRB2 /// LILRB3 | 5.485 | 5.241 | 0.059 | 0.023 |
| 209842_at | SOX10 | 5.288 | 5.044 | 0.059 | 0.007 |
| 207561_s_at | ACCN3 | 5.258 | 5.015 | 0.059 | 0.008 |
| 216769_x_at | C9orf150 | 5.190 | 4.947 | 0.059 | 0.012 |
| 205146_x_at | APBA3 | 5.999 | 5.756 | 0.059 | 0.007 |
| 217348_x_at | ARHGEF15 | 5.086 | 4.844 | 0.059 | 0.018 |
| 202015_x_at | METAP2 | 4.742 | 4.501 | 0.058 | 0.000 |
| 207771_at | SLC5A2 | 5.632 | 5.391 | 0.058 | 0.017 |

| | | | | | |
|---|---|---|---|---|---|
| 207313_x_at | KIR3DL2 | 5.311 | 5.071 | 0.057 | 0.037 |
| 206705_at | TULP1 | 5.471 | 5.232 | 0.057 | 0.002 |
| 210197_at | ITPK1 | 5.290 | 5.051 | 0.057 | 0.019 |
| 211835_at | IGH@ /// IGHA1 /// IGHA2 /// IGHD /// IGHG1 /// IGHG3 /// IGHG4 /// IGHM /// LOC642131 /// LOC652065 /// LOC652106 /// LOC652848 | 5.205 | 4.966 | 0.057 | 0.047 |
| 219824_at | SLC13A4 | 5.557 | 5.318 | 0.057 | 0.020 |
| 217161_x_at | AGC1 | 5.432 | 5.194 | 0.057 | 0.032 |
| 208173_at | IFNB1 | 5.209 | 4.971 | 0.057 | 0.024 |
| 211364_at | MTAP | 4.955 | 4.717 | 0.057 | 0.002 |
| 215971_at | C14orf135 | 5.515 | 5.277 | 0.057 | 0.005 |
| 207864_at | SCN7A | 5.024 | 4.788 | 0.056 | 0.023 |
| 208599_at | HUWE1 | 5.296 | 5.061 | 0.055 | 0.039 |
| 210782_x_at | GRIN1 | 6.002 | 5.766 | 0.055 | 0.025 |
| 215618_at | RSU1 | 5.697 | 5.461 | 0.055 | 0.018 |
| 215943_at | KIAA1661 | 5.015 | 4.781 | 0.055 | 0.001 |
| 209638_x_at | RGS12 | 4.857 | 4.623 | 0.055 | 0.007 |
| 210181_s_at | CABP1 | 5.528 | 5.294 | 0.055 | 0.004 |
| 208105_at | GIPR | 5.257 | 5.023 | 0.055 | 0.008 |
| 219684_at | RTP4 | 4.960 | 4.726 | 0.055 | 0.049 |
| 217459_at | --- | 5.117 | 4.884 | 0.054 | 0.014 |
| 222123_s_at | HIF3A | 5.452 | 5.221 | 0.053 | 0.027 |
| 209959_at | NR4A3 | 5.333 | 5.102 | 0.053 | 0.014 |
| 203666_at | CXCL12 | 4.789 | 4.558 | 0.053 | 0.001 |
| 204685_s_at | ATP2B2 | 5.397 | 5.166 | 0.053 | 0.023 |
| 206353_at | COX6A2 | 5.405 | 5.174 | 0.053 | 0.036 |
| 220412_x_at | KCNK7 | 5.171 | 4.940 | 0.053 | 0.008 |
| 214405_at | CUGBP2 | 4.774 | 4.544 | 0.053 | 0.025 |
| 215863_at | TFR2 | 5.726 | 5.498 | 0.052 | 0.007 |
| 211310_at | EZH1 | 4.988 | 4.761 | 0.052 | 0.005 |
| 214839_at | LOC157627 | 5.686 | 5.459 | 0.052 | 0.030 |
| 214081_at | PLXDC1 | 4.950 | 4.723 | 0.052 | 0.019 |
| 215121_x_at | IGL@ /// IGLC1 /// IGLC2 /// IGLV4-3 /// IGLV3-25 /// IGLV2-14 | 5.279 | 5.052 | 0.051 | 0.030 |
| 210723_x_at | MGC4771 | 5.078 | 4.851 | 0.051 | 0.040 |
| 207202_s_at | NR1I2 | 5.079 | 4.853 | 0.051 | 0.028 |
| 206796_at | WISP1 | 4.802 | 4.575 | 0.051 | 0.006 |
| 207834_at | FBLN1 | 5.168 | 4.942 | 0.051 | 0.016 |
| 204890_s_at | LCK | 5.223 | 4.998 | 0.051 | 0.032 |
| 214181_x_at | LST1 | 5.137 | 4.912 | 0.051 | 0.029 |
| 214416_at | --- | 4.806 | 4.580 | 0.051 | 0.020 |

| | | | | | |
|---|---|---|---|---|---|
| 220879_at | --- | 5.194 | 4.969 | 0.051 | 0.037 |
| 211583_x_at | NCR3 | 5.117 | 4.892 | 0.051 | 0.043 |
| 206697_s_at | HP | 5.734 | 5.510 | 0.050 | 0.025 |
| 211545_at | GHRHR | 5.208 | 4.984 | 0.050 | 0.015 |
| 214514_at | MCM3AP | 5.127 | 4.905 | 0.050 | 0.045 |
| 203470_s_at | PLEK | 5.361 | 5.138 | 0.049 | 0.034 |
| 205938_at | PPM1E | 4.928 | 4.706 | 0.049 | 0.010 |
| 211199_s_at | ICOSLG | 5.290 | 5.068 | 0.049 | 0.004 |
| 220348_at | KBTBD9 | 5.192 | 4.970 | 0.049 | 0.026 |
| 215810_x_at | DST | 4.815 | 4.594 | 0.049 | 0.041 |
| 207969_x_at | ACRV1 | 5.050 | 4.828 | 0.049 | 0.001 |
| 221174_at | --- | 5.022 | 4.800 | 0.049 | 0.020 |
| 211127_x_at | EDA | 4.955 | 4.734 | 0.049 | 0.039 |
| 207955_at | CCL27 | 5.189 | 4.968 | 0.049 | 0.013 |
| 205624_at | CPA3 | 5.429 | 5.208 | 0.049 | 0.021 |
| 220813_at | CYSLTR2 | 5.493 | 5.272 | 0.049 | 0.036 |
| 215638_at | ERBB3 | 5.045 | 4.825 | 0.048 | 0.019 |
| 208609_s_at | TNXA /// TNXB | 5.074 | 4.855 | 0.048 | 0.030 |
| 206836_at | SLC6A3 | 5.514 | 5.296 | 0.048 | 0.007 |
| 208253_at | SIGLEC8 | 4.833 | 4.616 | 0.047 | 0.023 |
| 208028_s_at | GPX5 | 4.971 | 4.754 | 0.047 | 0.028 |
| 207251_at | MEP1B | 5.007 | 4.790 | 0.047 | 0.001 |
| 207129_at | CA5B | 5.573 | 5.356 | 0.047 | 0.024 |
| 221118_at | PKD2L2 | 4.949 | 4.733 | 0.047 | 0.007 |
| 213096_at | TMCC2 | 5.670 | 5.454 | 0.047 | 0.036 |
| 215861_at | RP4-724E16.2 | 5.047 | 4.831 | 0.047 | 0.004 |
| 210810_s_at | SLC6A5 | 4.891 | 4.675 | 0.047 | 0.049 |
| 220844_at | TCEB3B | 4.897 | 4.681 | 0.046 | 0.000 |
| 220376_at | LRRC19 | 4.837 | 4.622 | 0.046 | 0.022 |
| 215805_at | --- | 5.131 | 4.916 | 0.046 | 0.001 |
| 215449_at | BZRPL1 | 5.102 | 4.888 | 0.046 | 0.015 |
| 222225_at | FLJ45055 | 5.906 | 5.691 | 0.046 | 0.031 |
| 215799_at | --- | 5.298 | 5.085 | 0.046 | 0.021 |
| 40640_at | hCAP-H2 | 5.946 | 5.733 | 0.045 | 0.039 |
| 208263_at | --- | 4.832 | 4.619 | 0.045 | 0.012 |
| 206974_at | CXCR6 | 5.216 | 5.004 | 0.045 | 0.017 |
| 217424_at | --- | 5.242 | 5.031 | 0.044 | 0.021 |
| 202222_s_at | DES | 5.558 | 5.348 | 0.044 | 0.049 |
| 206455_s_at | RHO | 5.085 | 4.876 | 0.044 | 0.002 |
| 207217_s_at | NOX1 | 5.331 | 5.122 | 0.044 | 0.023 |
| 206502_s_at | INSM1 | 5.011 | 4.802 | 0.044 | 0.032 |
| 211577_s_at | IGF1 | 4.842 | 4.634 | 0.043 | 0.045 |
| 210341_at | MYT1 | 5.162 | 4.954 | 0.043 | 0.037 |
| 221303_at | PCDHB1 | 4.954 | 4.746 | 0.043 | 0.009 |
| 205524_s_at | HAPLN1 | 5.373 | 5.166 | 0.043 | 0.038 |

| | | | | | |
|---|---|---|---|---|---|
| 215278_at | NNT | 5.422 | 5.216 | 0.043 | 0.029 |
| 222058_at | --- | 5.117 | 4.911 | 0.042 | 0.013 |
| 221072_at | C9orf31 | 5.171 | 4.966 | 0.042 | 0.036 |
| 216984_x_at | IGL@ | 5.280 | 5.075 | 0.042 | 0.016 |
| 206773_at | LY6H | 4.663 | 4.459 | 0.042 | 0.034 |
| 217469_at | IGHG1 | 5.319 | 5.115 | 0.042 | 0.030 |
| 205683_x_at | TPSAB1 | 5.013 | 4.808 | 0.042 | 0.011 |
| 217674_at | --- | 4.841 | 4.637 | 0.042 | 0.017 |
| 217323_at | HLA-DRB6 | 5.099 | 4.895 | 0.042 | 0.021 |
| 219337_at | C1orf159 | 5.983 | 5.779 | 0.042 | 0.043 |
| 219991_at | SLC2A9 | 5.257 | 5.053 | 0.041 | 0.047 |
| 211820_x_at | GYPA | 4.904 | 4.700 | 0.041 | 0.003 |
| 206776_x_at | ACRV1 | 4.946 | 4.743 | 0.041 | 0.011 |
| 215881_x_at | SSX2 /// SSX4 /// SSX3 /// SSX7 /// SSX9 /// LOC648415 /// LOC652163 /// LOC652630 /// LOC653088 | 5.253 | 5.050 | 0.041 | 0.037 |
| 210659_at | CMKLR1 | 5.049 | 4.847 | 0.041 | 0.028 |
| 220476_s_at | C1orf183 | 5.497 | 5.294 | 0.041 | 0.008 |
| 208224_at | HOXB1 | 5.295 | 5.093 | 0.041 | 0.028 |
| 208128_x_at | KIF25 | 5.500 | 5.298 | 0.041 | 0.043 |
| 208516_at | MTNR1B | 5.157 | 4.954 | 0.041 | 0.011 |
| 206815_at | SPAG8 | 5.166 | 4.964 | 0.041 | 0.046 |
| 215328_at | KIAA0953 | 4.980 | 4.779 | 0.041 | 0.000 |
| 214207_s_at | CARD10 | 5.295 | 5.095 | 0.040 | 0.030 |
| 214685_at | C4orf10 | 5.115 | 4.915 | 0.040 | 0.010 |
| 218621_at | HEMK1 | 5.969 | 5.770 | 0.040 | 0.045 |
| 214669_x_at | IGKC | 5.190 | 4.990 | 0.040 | 0.024 |
| 216761_at | RAB33A | 5.304 | 5.105 | 0.040 | 0.030 |
| 213439_x_at | RPIP8 | 5.328 | 5.129 | 0.040 | 0.022 |
| 220737_at | RPS6KA6 | 4.868 | 4.671 | 0.039 | 0.007 |
| 210031_at | CD247 | 4.978 | 4.780 | 0.039 | 0.023 |
| 206301_at | TEC | 5.129 | 4.932 | 0.039 | 0.022 |
| 208357_x_at | CSH1 | 5.074 | 4.877 | 0.039 | 0.030 |
| 217123_x_at | PMCHL1 | 5.269 | 5.073 | 0.039 | 0.049 |
| 220055_at | ZNF287 | 4.920 | 4.724 | 0.039 | 0.028 |
| 203747_at | AQP3 | 5.950 | 5.754 | 0.038 | 0.041 |
| 214165_s_at | HS6ST1 | 4.731 | 4.536 | 0.038 | 0.043 |
| 206646_at | GLI1 | 5.208 | 5.014 | 0.038 | 0.033 |
| 220257_x_at | NXF2 /// LOC650686 /// LOC653244 | 5.133 | 4.938 | 0.038 | 0.021 |
| 221347_at | CHRM5 | 4.813 | 4.618 | 0.038 | 0.029 |
| 214128_at | C11orf11 | 5.684 | 5.490 | 0.038 | 0.018 |

| | | | | | |
|---|---|---|---|---|---|
| 208314_at | RRH | 5.030 | 4.836 | 0.038 | 0.019 |
| 222295_x_at | --- | 5.429 | 5.235 | 0.038 | 0.011 |
| 220920_at | ATP10B | 5.292 | 5.098 | 0.038 | 0.010 |
| 213845_at | GRIK2 | 5.130 | 4.937 | 0.037 | 0.004 |
| 219977_at | AIPL1 | 5.020 | 4.827 | 0.037 | 0.030 |
| 215865_at | SYT12 | 5.091 | 4.898 | 0.037 | 0.005 |
| 219365_s_at | CAMKV | 5.427 | 5.234 | 0.037 | 0.019 |
| 217250_s_at | CHD5 | 5.091 | 4.898 | 0.037 | 0.017 |
| 221298_s_at | SLC22A8 | 4.980 | 4.788 | 0.037 | 0.008 |
| 64064_at | GIMAP5 | 5.536 | 5.344 | 0.037 | 0.004 |
| 220902_at | FLJ12616 | 5.434 | 5.243 | 0.036 | 0.046 |
| 219735_s_at | TFCP2L1 | 5.334 | 5.144 | 0.036 | 0.030 |
| 222174_at | MAX | 4.908 | 4.718 | 0.036 | 0.002 |
| 210166_at | TLR5 | 5.285 | 5.096 | 0.036 | 0.029 |
| 211248_s_at | CHRD | 5.141 | 4.952 | 0.036 | 0.035 |
| 214328_s_at | HSP90AA1 | 14.088 | 13.898 | 0.036 | 0.024 |
| 216665_s_at | TTTY2 | 4.700 | 4.512 | 0.036 | 0.007 |
| 214646_at | HIST1H3J | 5.053 | 4.865 | 0.035 | 0.030 |
| 217648_at | RWDD3 | 4.902 | 4.714 | 0.035 | 0.032 |
| 220958_at | ULK4 | 4.968 | 4.781 | 0.035 | 0.001 |
| 221200_at | LOC646541 | 4.873 | 4.686 | 0.035 | 0.030 |
| 220804_s_at | TP73 | 5.241 | 5.055 | 0.035 | 0.023 |
| 38964_r_at | WAS | 6.902 | 6.715 | 0.035 | 0.028 |
| 211369_at | --- | 4.984 | 4.799 | 0.034 | 0.022 |
| 216651_s_at | GAD2 | 4.942 | 4.756 | 0.034 | 0.021 |
| 216062_at | CD44 | 4.922 | 4.737 | 0.034 | 0.015 |
| 220791_x_at | SCN11A | 5.011 | 4.828 | 0.034 | 0.001 |
| 216214_at | --- | 4.939 | 4.756 | 0.033 | 0.005 |
| 211390_at | CG018 | 4.767 | 4.585 | 0.033 | 0.035 |
| 215755_at | --- | 5.093 | 4.910 | 0.033 | 0.036 |
| 214636_at | CALCB | 4.829 | 4.646 | 0.033 | 0.024 |
| 205577_at | PYGM | 4.986 | 4.803 | 0.033 | 0.039 |
| 215461_at | ZNRF4 | 5.301 | 5.118 | 0.033 | 0.015 |
| 213830_at | TRA@ | 4.839 | 4.657 | 0.033 | 0.048 |
| 207572_at | --- | 5.198 | 5.015 | 0.033 | 0.010 |
| 215911_x_at | ATP2B3 | 5.146 | 4.964 | 0.033 | 0.047 |
| 208573_s_at | OR2H2 | 5.329 | 5.147 | 0.033 | 0.012 |
| 220006_at | CCDC48 | 5.018 | 4.836 | 0.033 | 0.023 |
| 221373_x_at | PSPN | 5.582 | 5.401 | 0.033 | 0.035 |
| 220834_at | MS4A12 | 4.948 | 4.768 | 0.033 | 0.033 |
| 221150_at | MEPE | 5.640 | 5.460 | 0.032 | 0.009 |
| 205834_s_at | PART1 | 5.131 | 4.951 | 0.032 | 0.019 |
| 218623_at | HMP19 | 5.078 | 4.899 | 0.032 | 0.042 |
| 208574_at | SOX14 | 5.144 | 4.966 | 0.032 | 0.009 |
| 210259_s_at | DLX4 | 4.761 | 4.583 | 0.032 | 0.038 |

| | | | | | |
|---|---|---|---|---|---|
| 204851_s_at | DCX | 4.900 | 4.723 | 0.032 | 0.040 |
| 205456_at | CD3E | 4.921 | 4.743 | 0.031 | 0.021 |
| 211215_x_at | DIO2 | 5.359 | 5.182 | 0.031 | 0.011 |
| 217187_at | MUC5AC | 5.234 | 5.057 | 0.031 | 0.034 |
| 222300_at | CAPZA1 | 5.083 | 4.907 | 0.031 | 0.031 |
| 215846_at | CDC42SE2 | 5.131 | 4.955 | 0.031 | 0.048 |
| 221162_at | HHLA1 | 5.162 | 4.986 | 0.031 | 0.000 |
| 206844_at | FBP2 | 5.334 | 5.158 | 0.031 | 0.008 |
| 206458_s_at | WNT2B | 4.804 | 4.629 | 0.031 | 0.028 |
| 217483_at | FOLH1 | 5.014 | 4.839 | 0.031 | 0.019 |
| 207363_at | RS1 | 4.917 | 4.742 | 0.031 | 0.005 |
| 205114_s_at | CCL3 /// CCL3L1 /// CCL3L3 /// LOC643930 | 4.990 | 4.815 | 0.031 | 0.039 |
| 217303_s_at | ADRB3 | 5.524 | 5.349 | 0.030 | 0.045 |
| 207944_at | OCM | 5.074 | 4.900 | 0.030 | 0.001 |
| 213525_at | --- | 5.804 | 5.630 | 0.030 | 0.013 |
| 211287_x_at | CSF2RA | 5.208 | 5.035 | 0.030 | 0.039 |
| 219167_at | RASL12 | 5.040 | 4.867 | 0.030 | 0.003 |
| 215234_at | LOC653542 | 4.651 | 4.477 | 0.030 | 0.007 |
| 218369_s_at | EXOSC1 | 4.926 | 4.754 | 0.030 | 0.044 |
| 214400_at | INSL3 | 5.072 | 4.900 | 0.030 | 0.028 |
| 211314_at | CACNA1G | 5.118 | 4.946 | 0.030 | 0.005 |
| 214275_at | MED12 | 4.940 | 4.768 | 0.030 | 0.046 |
| 215902_at | 6-Mar | 4.921 | 4.750 | 0.030 | 0.021 |
| 220604_x_at | FTCD | 5.295 | 5.124 | 0.029 | 0.029 |
| 221159_at | --- | 5.008 | 4.837 | 0.029 | 0.027 |
| 216451_at | STK38 | 5.218 | 5.047 | 0.029 | 0.037 |
| 212868_x_at | C12orf47 | 5.205 | 5.035 | 0.029 | 0.002 |
| 217706_at | LRRC51 | 5.148 | 4.977 | 0.029 | 0.046 |
| 200865_at | --- | 4.889 | 4.719 | 0.029 | 0.001 |
| 210391_at | NR6A1 | 5.194 | 5.024 | 0.029 | 0.044 |
| 208239_at | FOXE1 | 5.053 | 4.883 | 0.029 | 0.022 |
| 221378_at | CER1 | 5.384 | 5.214 | 0.029 | 0.011 |
| 207093_s_at | OMG | 4.907 | 4.737 | 0.029 | 0.045 |
| 208172_s_at | KCNB2 | 4.810 | 4.640 | 0.029 | 0.018 |
| 215598_at | TTC12 | 5.273 | 5.103 | 0.029 | 0.030 |
| 221157_s_at | FBXO24 | 5.167 | 4.997 | 0.029 | 0.036 |
| 219873_at | COLEC11 | 5.074 | 4.904 | 0.029 | 0.016 |
| 205890_s_at | GABBR1 /// UBD | 4.953 | 4.784 | 0.028 | 0.038 |
| 206360_s_at | SOCS3 | 4.933 | 4.764 | 0.028 | 0.025 |
| 220747_at | HSPC072 | 5.544 | 5.376 | 0.028 | 0.050 |
| 206643_at | HAL | 5.277 | 5.109 | 0.028 | 0.023 |
| 220583_at | FLJ22596 | 5.444 | 5.278 | 0.028 | 0.026 |
| 216612_x_at | --- | 4.963 | 4.797 | 0.028 | 0.026 |
| 208222_at | ACVR1B | 5.102 | 4.935 | 0.028 | 0.007 |

| | | | | | |
|---|---|---|---|---|---|
| 206166_s_at | CLCA2 | 5.089 | 4.923 | 0.028 | 0.002 |
| 206916_x_at | TAT | 4.990 | 4.825 | 0.027 | 0.022 |
| 215972_at | PART1 | 5.032 | 4.867 | 0.027 | 0.049 |
| 217637_at | --- | 5.216 | 5.051 | 0.027 | 0.048 |
| 205270_s_at | LCP2 | 4.828 | 4.664 | 0.027 | 0.024 |
| 209791_at | PADI2 | 5.396 | 5.232 | 0.027 | 0.017 |
| 221244_s_at | --- | 5.338 | 5.175 | 0.027 | 0.049 |
| 214082_at | CA5B | 5.409 | 5.246 | 0.027 | 0.001 |
| 211142_x_at | HLA-DOA | 5.111 | 4.949 | 0.027 | 0.027 |
| 211830_s_at | CACNA1I | 5.813 | 5.651 | 0.026 | 0.026 |
| 206489_s_at | DLGAP1 | 4.893 | 4.730 | 0.026 | 0.025 |
| 219845_at | BARX1 | 5.261 | 5.099 | 0.026 | 0.039 |
| 216596_at | DKFZP434L187 | 5.271 | 5.109 | 0.026 | 0.044 |
| 210921_at | --- | 5.494 | 5.333 | 0.026 | 0.034 |
| 213841_at | --- | 5.250 | 5.089 | 0.026 | 0.049 |
| 211648_at | IGHG1 | 4.899 | 4.738 | 0.026 | 0.015 |
| 208006_at | FOXI1 | 5.084 | 4.924 | 0.026 | 0.021 |
| 206459_s_at | WNT2B | 5.215 | 5.055 | 0.026 | 0.024 |
| 217233_at | --- | 4.918 | 4.758 | 0.025 | 0.025 |
| 211856_x_at | CD28 | 5.203 | 5.044 | 0.025 | 0.020 |
| 217115_at | --- | 5.316 | 5.157 | 0.025 | 0.039 |
| 217361_at | --- | 5.383 | 5.225 | 0.025 | 0.036 |
| 221001_at | LOC647835 | 5.041 | 4.882 | 0.025 | 0.039 |
| 216162_at | SBNO1 | 4.857 | 4.699 | 0.025 | 0.028 |
| 220492_s_at | OTOF | 4.790 | 4.632 | 0.025 | 0.023 |
| 216704_at | TBCA | 5.155 | 4.998 | 0.025 | 0.045 |
| 216053_x_at | C20orf91 /// RP13-401N8.2 | 5.140 | 4.984 | 0.024 | 0.023 |
| 216168_at | RHOH | 4.898 | 4.742 | 0.024 | 0.018 |
| 217676_at | --- | 5.109 | 4.954 | 0.024 | 0.038 |
| 216122_at | NAV1 | 4.954 | 4.799 | 0.024 | 0.024 |
| 206735_at | CHRNA4 | 5.426 | 5.270 | 0.024 | 0.003 |
| 207248_at | KCNA4 | 5.497 | 5.343 | 0.024 | 0.031 |
| 211192_s_at | CD84 | 4.841 | 4.686 | 0.024 | 0.041 |
| 217587_at | HNRPF | 4.997 | 4.842 | 0.024 | 0.047 |
| 210019_at | --- | 5.009 | 4.855 | 0.024 | 0.006 |
| 211129_x_at | EDA | 5.073 | 4.920 | 0.023 | 0.029 |
| 207672_at | --- | 4.980 | 4.827 | 0.023 | 0.022 |
| 206390_x_at | PF4 | 5.958 | 5.807 | 0.023 | 0.023 |
| 215608_at | --- | 5.132 | 4.980 | 0.023 | 0.037 |
| 216024_at | DNM2 | 4.765 | 4.614 | 0.023 | 0.024 |
| 217085_at | SLC14A2 | 4.770 | 4.619 | 0.023 | 0.042 |
| 216437_at | EPC1 | 4.705 | 4.554 | 0.023 | 0.035 |
| 211207_s_at | ACSL6 | 4.599 | 4.448 | 0.023 | 0.050 |
| 221594_at | DKFZP564O0523 | 4.998 | 4.847 | 0.023 | 0.026 |

| | | | | | |
|---|---|---|---|---|---|
| 216578_at | --- | 5.216 | 5.065 | 0.023 | 0.015 |
| 206605_at | P11 | 4.994 | 4.844 | 0.022 | 0.031 |
| 217219_at | DKFZP434A062 | 5.644 | 5.494 | 0.022 | 0.045 |
| 210262_at | CRISP2 | 4.860 | 4.710 | 0.022 | 0.044 |
| 209863_s_at | TP73L | 5.043 | 4.893 | 0.022 | 0.014 |
| 206250_x_at | AVPR1A | 5.079 | 4.930 | 0.022 | 0.033 |
| 215662_at | --- | 4.928 | 4.780 | 0.022 | 0.022 |
| 217096_at | PCLO | 4.946 | 4.798 | 0.022 | 0.026 |
| 216638_s_at | PRLR | 4.759 | 4.610 | 0.022 | 0.006 |
| 214530_x_at | EPB41 | 4.704 | 4.556 | 0.022 | 0.014 |
| 210133_at | CCL11 | 5.005 | 4.858 | 0.022 | 0.008 |
| 216400_at | GBA /// GBAP | 4.959 | 4.812 | 0.021 | 0.030 |
| 215840_at | DNHD3 | 5.226 | 5.080 | 0.021 | 0.031 |
| 205912_at | PNLIP | 4.823 | 4.677 | 0.021 | 0.018 |
| 37145_at | GNLY | 4.822 | 4.676 | 0.021 | 0.031 |
| 207224_s_at | SIGLEC7 | 4.966 | 4.821 | 0.021 | 0.033 |
| 215822_x_at | MYT1 | 5.435 | 5.289 | 0.021 | 0.036 |
| 207381_at | ALOX12B | 5.037 | 4.892 | 0.021 | 0.034 |
| 221239_s_at | FCRL2 | 4.809 | 4.664 | 0.021 | 0.012 |
| 220359_s_at | ARPP-21 | 4.783 | 4.638 | 0.021 | 0.014 |
| 207461_at | --- | 4.639 | 4.494 | 0.021 | 0.010 |
| 215777_at | --- | 4.870 | 4.726 | 0.021 | 0.038 |
| 214621_at | GYS2 | 4.687 | 4.543 | 0.020 | 0.023 |
| 222086_s_at | WNT6 | 4.947 | 4.804 | 0.020 | 0.019 |
| 210669_at | TFAP2A | 5.266 | 5.123 | 0.020 | 0.023 |
| 206952_at | G6PC | 4.645 | 4.502 | 0.020 | 0.046 |
| 204723_at | SCN3B | 4.862 | 4.720 | 0.020 | 0.043 |
| 210356_x_at | MS4A1 | 4.996 | 4.854 | 0.020 | 0.037 |
| 214928_at | OBSL1 | 4.931 | 4.790 | 0.020 | 0.044 |
| 202485_s_at | MBD2 | 5.125 | 4.983 | 0.020 | 0.021 |
| 214973_x_at | IGHD | 5.302 | 5.161 | 0.020 | 0.029 |
| 1255_g_at | GUCA1A | 4.689 | 4.548 | 0.020 | 0.029 |
| 219827_at | UCP3 | 5.428 | 5.288 | 0.020 | 0.042 |
| 206426_at | MLANA | 4.868 | 4.727 | 0.020 | 0.021 |
| 215056_at | --- | 5.118 | 4.978 | 0.020 | 0.022 |
| 210579_s_at | TRIM10 | 4.890 | 4.751 | 0.019 | 0.048 |
| 208548_at | IFNA6 | 4.741 | 4.602 | 0.019 | 0.012 |
| 221287_at | RNASEL | 5.188 | 5.049 | 0.019 | 0.018 |
| 203808_at | --- | 4.732 | 4.594 | 0.019 | 0.030 |
| 221160_s_at | CABP3 /// CABP5 | 4.923 | 4.786 | 0.019 | 0.050 |
| 214072_x_at | NENF | 5.509 | 5.372 | 0.019 | 0.046 |
| 209793_at | GRIA1 | 4.951 | 4.815 | 0.019 | 0.007 |
| 208382_s_at | DMC1 | 5.045 | 4.909 | 0.018 | 0.047 |
| 206069_s_at | ACADL | 4.595 | 4.459 | 0.018 | 0.008 |
| 221092_at | ZNFN1A3 | 4.789 | 4.653 | 0.018 | 0.047 |

| | | | | | |
|---|---|---|---|---|---|
| 205831_at | CD2 | 5.278 | 5.143 | 0.018 | 0.039 |
| 207885_at | S100G | 4.991 | 4.857 | 0.018 | 0.004 |
| 220871_at | --- | 4.788 | 4.654 | 0.018 | 0.034 |
| 215979_s_at | SLC7A1 | 5.418 | 5.284 | 0.018 | 0.029 |
| 216877_at | DKFZp686O1327 | 4.917 | 4.785 | 0.017 | 0.047 |
| 220540_at | KCNK15 | 4.786 | 4.654 | 0.017 | 0.013 |
| 203877_at | --- | 4.927 | 4.795 | 0.017 | 0.041 |
| 222220_s_at | TSNAXIP1 | 5.360 | 5.229 | 0.017 | 0.011 |
| 215332_s_at | CD8B | 4.901 | 4.770 | 0.017 | 0.045 |
| 216276_s_at | ADAM3A | 4.688 | 4.557 | 0.017 | 0.025 |
| 216409_at | ACSL6 | 4.963 | 4.832 | 0.017 | 0.043 |
| 211617_at | ALDOAP2 | 4.834 | 4.704 | 0.017 | 0.040 |
| 216412_x_at | --- | 5.228 | 5.099 | 0.017 | 0.018 |
| 31835_at | HRG | 4.795 | 4.667 | 0.016 | 0.031 |
| 220505_at | C9orf53 | 4.909 | 4.781 | 0.016 | 0.011 |
| 210833_at | PTGER3 | 4.821 | 4.695 | 0.016 | 0.024 |
| 210548_at | CCL23 | 4.876 | 4.751 | 0.016 | 0.034 |
| 207316_at | HAS1 | 4.926 | 4.800 | 0.016 | 0.046 |
| 216898_s_at | COL4A3 | 4.824 | 4.699 | 0.016 | 0.009 |
| 207058_s_at | PARK2 | 4.857 | 4.732 | 0.016 | 0.033 |
| 216780_at | --- | 5.135 | 5.010 | 0.015 | 0.050 |
| 210159_s_at | TRIM31 | 5.026 | 4.902 | 0.015 | 0.040 |
| 222271_at | --- | 5.015 | 4.894 | 0.015 | 0.033 |
| 203948_s_at | MPO | 4.811 | 4.690 | 0.015 | 0.002 |
| 206608_s_at | RPGRIP1 | 4.948 | 4.827 | 0.015 | 0.033 |
| 217271_at | GNA11 | 4.819 | 4.699 | 0.015 | 0.019 |
| 216018_at | RNF5 | 5.059 | 4.938 | 0.014 | 0.037 |
| 215340_at | ADCY1 | 5.119 | 4.999 | 0.014 | 0.037 |
| 213832_at | --- | 4.857 | 4.738 | 0.014 | 0.039 |
| 209915_s_at | NRXN1 | 4.927 | 4.808 | 0.014 | 0.050 |
| 205686_s_at | CD86 | 4.692 | 4.574 | 0.014 | 0.024 |
| 217428_s_at | COL10A1 | 4.843 | 4.725 | 0.014 | 0.048 |
| 217090_at | ADAM3A | 4.736 | 4.618 | 0.014 | 0.048 |
| 206078_at | KALRN | 4.883 | 4.765 | 0.014 | 0.043 |
| 222383_s_at | ALOXE3 | 5.015 | 4.898 | 0.014 | 0.029 |
| 221121_at | CXorf48 | 4.991 | 4.874 | 0.014 | 0.045 |
| 214503_x_at | GPR135 | 4.816 | 4.700 | 0.013 | 0.041 |
| 213831_at | HLA-DQA1 | 4.881 | 4.768 | 0.013 | 0.036 |
| 216159_s_at | --- | 5.043 | 4.930 | 0.013 | 0.046 |
| 214981_at | POSTN | 4.759 | 4.647 | 0.012 | 0.027 |
| 217464_at | RPS2 | 4.902 | 4.792 | 0.012 | 0.038 |
| 217563_at | CLOCK | 5.102 | 4.992 | 0.012 | 0.048 |
| 221424_s_at | OR51E2 | 4.785 | 4.676 | 0.012 | 0.029 |
| 221217_s_at | A2BP1 | 5.467 | 5.359 | 0.012 | 0.038 |
| 37201_at | ITIH4 | 5.155 | 5.047 | 0.012 | 0.047 |

| | | | | | |
|---|---|---|---|---|---|
| 221271_at | IL21 | 5.042 | 4.934 | 0.012 | 0.020 |
| 206577_at | VIP | 4.832 | 4.725 | 0.011 | 0.031 |
| 216807_at | KIAA1751 /// LOC642155 | 4.925 | 4.819 | 0.011 | 0.006 |
| 34210_at | CD52 | 4.768 | 4.664 | 0.011 | 0.019 |
| 211917_s_at | PRLR | 4.837 | 4.735 | 0.010 | 0.018 |
| 217272_s_at | SERPINB13 | 4.988 | 4.886 | 0.010 | 0.024 |
| 216575_at | --- | 4.746 | 4.647 | 0.010 | 0.036 |
| 206759_at | FCER2 | 4.978 | 4.880 | 0.010 | 0.045 |
| 206841_at | PDE6H | 4.812 | 4.715 | 0.009 | 0.048 |
| 216127_at | PDIA2 | 5.089 | 4.992 | 0.009 | 0.048 |
| 210375_at | PTGER3 | 4.674 | 4.580 | 0.009 | 0.013 |
| 216897_s_at | FAM76A | 5.043 | 4.951 | 0.008 | 0.036 |
| 215272_at | --- | 4.943 | 4.851 | 0.008 | 0.004 |
| 221682_s_at | PCDHGB6 | 4.892 | 4.804 | 0.008 | 0.009 |
| 209474_s_at | ENTPD1 | 4.633 | 4.546 | 0.008 | 0.040 |
| 91580_at | LRTM1 | 4.660 | 4.574 | 0.007 | 0.016 |
| 214208_at | FLJ33790 | 5.315 | 5.232 | 0.007 | 0.042 |
| 216255_s_at | GRM8 | 4.948 | 4.866 | 0.007 | 0.022 |
| 216441_at | --- | 4.735 | 4.660 | 0.006 | 0.022 |
| 207906_at | IL3 | 4.480 | 4.411 | 0.005 | 0.038 |
| 206535_at | SLC2A2 | 4.661 | 4.593 | 0.005 | 0.031 |
| **2 transformed RMA values** | | | | | |
