## Supplementary material for "Targeting dormant ovarian cancer cells *in vitro* and in an *in vivo* model of platinum resistance": Table S4

| **Table S4. Functional annotations for genes with higher expression in cells with high spheroid-forming capacity** | | | | | | | | | | |
|---|---|---|---|---|---|---|---|---|---|---|
| Annotation Cluster 1 | Enrichment Score: 11.568772130117555 | | | | | | | | | |
| Category | Term | Count | % | PValue | Genes | List Total | Pop Hits | Pop Total | Fold Enrich | FDR |
| UP_KEYWORDS | Mitochondrion | 137 | 12.85178 | 1.59E-22 | 220103_S_AT, 204283_AT, 221550_AT, 201256_AT, 200086_S_AT, 201490_S_AT, 216591_S_AT, 207829_S_AT, 209549_S_AT, 218046_S_AT, 200946_X_AT, 201821_S_AT, 205217_AT, 201226_AT, 201300_S_AT, 217919_S_AT, 204386_S_AT, 202004_X_AT, 217772_S_AT, 221620_S_AT, 203213_AT, 205361_S_AT, 200796_S_AT, 208369_S_AT, 208910_S_AT, 203613_S_AT, 203437_AT, 203926_X_AT, 200798_X_AT, 218202_X_AT, 209346_S_AT, 217874_AT, 220631_AT, 208541_X_AT, 200658_S_AT, 215195_AT, 215524_X_AT, 218027_AT, 206686_AT, 217140_S_AT, 201559_S_AT, 210626_AT, 218890_X_AT, 215707_S_AT, 214437_S_AT, 37226_AT, 218118_S_AT, 209249_S_AT, 219006_AT, 209445_X_AT, 36830_AT, 212600_S_AT, 202026_AT, 218119_AT, 218993_AT, 204305_AT, 210154_AT, 215037_S_AT, 218597_S_AT, 204300_AT, 218220_AT, | 1035 | 1119 | 20581 | 2.434538 | 2.25E-19 |
| UP_KEYWORDS | Transit peptide | 68 | 6.378987 | 5.27E-12 | 220103_S_AT, 208969_AT, 204283_AT, 203177_X_AT, 208972_S_AT, 201256_AT, 200086_S_AT, 201490_S_AT, 216591_S_AT, 209549_S_AT, 209078_S_AT, 200947_S_AT, 219281_AT, 200946_X_AT, 218046_S_AT, 203207_S_AT, 201226_AT, 217919_S_AT, 202004_X_AT, 208369_S_AT, 208910_S_AT, 220329_S_AT, 203926_X_AT, 208764_S_AT, 218202_X_AT, 214214_S_AT, 208909_AT, 217874_AT, 220631_AT, 208541_X_AT, 219575_S_AT, 201066_AT, 218027_AT, 211752_S_AT, 208911_S_AT, 206686_AT, 214095_AT, 218890_X_AT, 210626_AT, 210153_S_AT, 214437_S_AT, 211150_S_AT, 207552_AT, 209249_S_AT, 205401_AT, 206348_S_AT, 36830_AT, 210131_X_AT, 212600_S_AT, 202026_AT, 201007_AT, 218993_AT, 200793_S_AT, 210154_AT, 204305_AT, 203176_S_AT, 212410_AT, 204300_AT, 218220_AT, 218982_S_AT, 203621_AT, 203208_S_AT, | 1035 | 536 | 20581 | 2.522727 | 7.48E-09 |
| UP_SEQ_FEATURE | transit peptide:Mitochond | 60 | 5.628518 | 4.79E-10 | 220103_S_AT, 208969_AT, 204283_AT, 203177_X_AT, 208972_S_AT, 201256_AT, 200086_S_AT, 201490_S_AT, 216591_S_AT, 209549_S_AT, 209078_S_AT, 200947_S_AT, 200946_X_AT, 218046_S_AT, 201226_AT, 217919_S_AT, 202004_X_AT, 208369_S_AT, 208910_S_AT, 203926_X_AT, 208764_S_AT, 218202_X_AT, 214214_S_AT, 208909_AT, 217874_AT, 208541_X_AT, 219575_S_AT, 201066_AT, 218027_AT, 211752_S_AT, 208911_S_AT, 206686_AT, 214095_AT, 218890_X_AT, 210626_AT, 210153_S_AT, 214437_S_AT, 211150_S_AT, 207552_AT, 206348_S_AT, 36830_AT, 210131_X_AT, 212600_S_AT, 202026_AT, 201007_AT, 200793_S_AT, 204305_AT, 210154_AT, 203176_S_AT, 204300_AT, 218220_AT, 218982_S_AT, 203621_AT, 215794_X_AT, 215772_X_AT, 214835_S_AT, 210667_S_AT, 219449_S_AT, 212432_AT, 217883_AT, 201322_AT, | 1029 | 482 | 20063 | 2.427083 | 8.39E-07 |

| | | | | | | | | | | |
|---|---|---|---|---|---|---|---|---|---|---|
| GOTERM_CC_DIRECT | GO:0005759~mitochond | 36 | 3.377111 | 1.33E-04 | 204283_AT, 200840_AT, 208969_AT, 203177_X_AT, 201490_S_AT, 209549_S_AT, 203270_AT, 200947_S_AT, 209078_S_AT, 200946_X_AT, 200079_S_AT, 200796_S_AT, 208369_S_AT, 208910_S_AT, 203926_X_AT, 200798_X_AT, 214214_S_AT, 201872_S_AT, 205133_S_AT, 217874_AT, 208541_X_AT, 211752_S_AT, 208911_S_AT, 206686_AT, 214095_AT, 210153_S_AT, 214437_S_AT, 211150_S_AT, 206348_S_AT, 36830_AT, 200793_S_AT, 204305_AT, 210154_AT, 215037_S_AT, 203176_S_AT, 215772_X_AT, 214835_S_AT, 208693_S_AT, 212432_AT, 212459_X_AT, 201322_AT, 208692_AT, 213133_S_AT, 213321_AT, 220236_AT, 213041_S_AT, 202589_AT, 210653_S_AT, 219176_AT, 203816_AT, 221957_AT | 1004 | 327 | 18224 | 1.998319 | 0.198946 |
| Annotation Cluster 2 | Enrichment Score: 5.913648776789434 | | | | | | | | | |
| Category | Term | Count | % | PValue | Genes | List Total | Pop Hits | Pop Total | Fold Enrich | FDR |
| UP_KEYWORDS | DNA damage | 46 | 4.315197 | 8.68E-09 | 203103_S_AT, 213829_X_AT, 213468_AT, 212525_S_AT, 210379_S_AT, 219960_S_AT, 200957_S_AT, 210813_S_AT, 218564_AT, 205393_S_AT, 200679_X_AT, 200669_S_AT, 208642_S_AT, 210257_X_AT, 218163_AT, 219361_S_AT, 201902_S_AT, 203198_AT, 204883_S_AT, 218738_S_AT, 41657_AT, 219502_AT, 210216_X_AT, 200680_X_AT, 218585_S_AT, 201459_AT, 217299_S_AT, 211297_S_AT, 202906_S_AT, 211077_S_AT, 219715_S_AT, 204460_S_AT, 220060_S_AT, 218658_S_AT, 210812_AT, 205446_S_AT, 209375_AT, 208619_AT, 208643_S_AT, 204884_S_AT, 214938_X_AT, 205394_AT, 210470_X_AT, 221381_S_AT, 201222_S_AT, 205071_X_AT, 215997_S_AT, 201461_S_AT, 210458_S_AT, 201614_S_AT, 216880_AT, 204768_S_AT, 208692_AT, 210027_S_AT, 202266_AT, 219628_AT, | 1035 | 352 | 20581 | 2.598611 | 1.23E-05 |
| UP_KEYWORDS | DNA repair | 36 | 3.377111 | 1.91E-06 | 203103_S_AT, 201459_AT, 217299_S_AT, 213829_X_AT, 212525_S_AT, 213468_AT, 211297_S_AT, 219960_S_AT, 202906_S_AT, 219715_S_AT, 204460_S_AT, 220060_S_AT, 200957_S_AT, 210813_S_AT, 218564_AT, 210812_AT, 218658_S_AT, 205393_S_AT, 200679_X_AT, 200669_S_AT, 209375_AT, 208619_AT, 208643_S_AT, 208642_S_AT, 214938_X_AT, 205394_AT, 210257_X_AT, 210470_X_AT, 221381_S_AT, 201222_S_AT, 205071_X_AT, 215997_S_AT, 201614_S_AT, 204768_S_AT, 216880_AT, 203198_AT, 201902_S_AT, 218738_S_AT, 208692_AT, 210027_S_AT, 202266_AT, 219502_AT, 210216_X_AT, 200680_X_AT, 203409_AT, 202214_S_AT, 202213_S_AT | 1035 | 293 | 20581 | 2.443211 | 0.002708 |

| | | | | | | | | | | |
|---|---|---|---|---|---|---|---|---|---|---|
| GOTERM_BP_DIRECT | GO:0006281~DNA repai | 30 | 2.814259 | 1.10E-04 | 201459_AT, 217299_S_AT, 213829_X_AT, 203565_S_AT, 216962_AT, 202906_S_AT, 219960_S_AT, 219715_S_AT, 204460_S_AT, 220060_S_AT, 200957_S_AT, 218564_AT, 218658_S_AT, 205393_S_AT, 200669_S_AT, 209375_AT, 203213_AT, 204194_AT, 208619_AT, 205091_X_AT, 204884_S_AT, 205394_AT, 210470_X_AT, 204742_S_AT, 201614_S_AT, 203214_X_AT, 216880_AT, 204768_S_AT, 204883_S_AT, 203198_AT, 208692_AT, 200997_AT, 210027_S_AT, 210216_X_AT, 203409_AT | 980 | 235 | 16792 | 2.187408 | 0.200438 |
| Annotation Cluster 3 | Enrichment Score: 5.597281746589656 | | | | | | | | | |
| Category | Term | Count | % | PValue | Genes | List Total | Pop Hits | Pop Total | Fold Enrich | FDR |
| GOTERM_BP_DIRECT | GO:0006412~translation | 44 | 4.12758 | 2.06E-10 | 209134_S_AT, 220103_S_AT, 201665_X_AT, 218398_AT, 200099_S_AT, 210984_X_AT, 218046_S_AT, 212018_S_AT, 200036_S_AT, 204386_S_AT, 217919_S_AT, 200715_X_AT, 220329_S_AT, 201652_AT, 211942_X_AT, 217915_S_AT, 221726_AT, 214143_X_AT, 212042_X_AT, 219575_S_AT, 218027_AT, 200017_AT, 218001_AT, 203340_S_AT, 218890_X_AT, 200088_X_AT, 200809_X_AT, 200029_AT, 209445_X_AT, 218653_AT, 201263_AT, 217747_S_AT, 204300_AT, 218982_S_AT, 200735_X_AT, 208635_X_AT, 209316_S_AT, 213897_S_AT, 201623_S_AT, 218049_S_AT, 200013_AT, 207088_S_AT, 214271_X_AT, 200034_S_AT, 213687_S_AT, 208692_AT, 217559_AT, 203800_S_AT, 203339_AT, 214167_S_AT | 980 | 253 | 16792 | 2.979947 | 3.78E-07 |
| GOTERM_CC_DIRECT | GO:0005840~ribosome | 32 | 3.001876 | 1.84E-09 | 220103_S_AT, 209134_S_AT, 218513_AT, 200017_AT, 218001_AT, 218890_X_AT, 201665_X_AT, 218398_AT, 200099_S_AT, 200029_AT, 218046_S_AT, 200036_S_AT, 217747_S_AT, 218982_S_AT, 217919_S_AT, 200715_X_AT, 213897_S_AT, 212145_AT, 218049_S_AT, 200013_AT, 207827_X_AT, 211942_X_AT, 217915_S_AT, 200034_S_AT, 213687_S_AT, 218202_X_AT, 208692_AT, 221726_AT, 210027_S_AT, 214143_X_AT, 203800_S_AT, 212042_X_AT, 214167_S_AT, 219819_S_AT | 1004 | 166 | 18224 | 3.499064 | 2.76E-06 |

| | | | | | | | | | | |
|---|---|---|---|---|---|---|---|---|---|---|
| UP_KEYWORDS | Ribosomal protein | 32 | 3.001876 | 3.30E-09 | 220103_S_AT, 209134_S_AT, 218027_AT, 200017_AT, 218001_AT, 218890_X_AT, 201665_X_AT, 200088_X_AT, 218398_AT, 200099_S_AT, 200809_X_AT, 200029_AT, 218046_S_AT, 200036_S_AT, 217747_S_AT, 204386_S_AT, 218982_S_AT, 217919_S_AT, 200715_X_AT, 213897_S_AT, 212145_AT, 218049_S_AT, 200013_AT, 214271_X_AT, 211942_X_AT, 200034_S_AT, 213687_S_AT, 218202_X_AT, 208692_AT, 217559_AT, 221726_AT, 214143_X_AT, 203800_S_AT, 212042_X_AT, 214167_S_AT, 219819_S_AT | 1035 | 185 | 20581 | 3.439572 | 4.69E-06 |
| UP_KEYWORDS | Ribonucleoprotein | 42 | 3.939962 | 3.85E-09 | 209134_S_AT, 220103_S_AT, 201665_X_AT, 218398_AT, 200099_S_AT, 218046_S_AT, 202690_S_AT, 200036_S_AT, 214737_X_AT, 204386_S_AT, 217919_S_AT, 200715_X_AT, 221688_S_AT, 202903_AT, 202691_AT, 211942_X_AT, 218202_X_AT, 221726_AT, 209330_S_AT, 214143_X_AT, 212042_X_AT, 218027_AT, 200017_AT, 202407_S_AT, 218001_AT, 218890_X_AT, 211747_S_AT, 200088_X_AT, 200809_X_AT, 202904_S_AT, 200751_S_AT, 200029_AT, 217747_S_AT, 218982_S_AT, 200014_S_AT, 201054_AT, 211932_AT, 213897_S_AT, 212145_AT, 218049_S_AT, 200013_AT, 214271_X_AT, 216855_S_AT, 200034_S_AT, 204559_S_AT, 213687_S_AT, 208692_AT, 217559_AT, 203800_S_AT, 214167_S_AT, 219819_S_AT | 1035 | 296 | 20581 | 2.821524 | 5.47E-06 |
| GOTERM_BP_DIRECT | GO:0006364~rRNA proc | 37 | 3.470919 | 8.14E-09 | 209134_S_AT, 212499_S_AT, 203721_S_AT, 200017_AT, 201665_X_AT, 200088_X_AT, 221098_X_AT, 200099_S_AT, 200809_X_AT, 203740_AT, 200056_S_AT, 200029_AT, 217822_AT, 220688_S_AT, 218481_AT, 218535_S_AT, 91684_G_AT, 203622_S_AT, 221987_S_AT, 218993_AT, 200036_S_AT, 217747_S_AT, 219927_AT, 218104_AT, 212844_AT, 200715_X_AT, 209233_AT, 218156_S_AT, 221688_S_AT, 200013_AT, 218512_AT, 214271_X_AT, 211942_X_AT, 204699_S_AT, 200034_S_AT, 213687_S_AT, 208692_AT, 221726_AT, 203171_S_AT, 214143_X_AT, 217821_S_AT, 212042_X_AT, 214167_S_AT, 218235_S_AT | 980 | 214 | 16792 | 2.962541 | 1.49E-05 |

| | | | | | | | | | | |
|---|---|---|---|---|---|---|---|---|---|---|
| GOTERM_BP_DIRECT | GO:0006413~translation | 28 | 2.626642 | 2.00E-08 | 209134_S_AT, 200017_AT, 200005_AT, 201665_X_AT, 200088_X_AT, 200023_S_AT, 200099_S_AT, 200809_X_AT, 200029_AT, 211937_AT, 201435_S_AT, 200036_S_AT, 217747_S_AT, 200647_X_AT, 211938_AT, 208726_S_AT, 200715_X_AT, 200013_AT, 219599_AT, 214271_X_AT, 201652_AT, 202232_S_AT, 211942_X_AT, 200034_S_AT, 215230_X_AT, 213687_S_AT, 217719_AT, 208692_AT, 221726_AT, 214143_X_AT, 201872_S_AT, 202461_AT, 212042_X_AT, 208697_S_AT, 214167_S_AT | 980 | 137 | 16792 | 3.501981 | 3.66E-05 |
| GOTERM_MF_DIRECT | GO:0003735~structural c | 36 | 3.377111 | 4.97E-08 | 220103_S_AT, 209134_S_AT, 218027_AT, 200017_AT, 218890_X_AT, 218001_AT, 203340_S_AT, 201665_X_AT, 200088_X_AT, 218398_AT, 200099_S_AT, 200809_X_AT, 200029_AT, 220688_S_AT, 218653_AT, 209445_X_AT, 218046_S_AT, 200036_S_AT, 217747_S_AT, 204386_S_AT, 218982_S_AT, 217919_S_AT, 200715_X_AT, 213897_S_AT, 221688_S_AT, 218049_S_AT, 200013_AT, 207088_S_AT, 214271_X_AT, 211942_X_AT, 217915_S_AT, 200034_S_AT, 213687_S_AT, 217559_AT, 208692_AT, 221726_AT, 214143_X_AT, 203800_S_AT, 203339_AT, 212042_X_AT, 214167_S_AT | 971 | 222 | 16881 | 2.819217 | 7.97E-05 |
| KEGG_PATHWAY | hsa03010:Ribosome | 27 | 2.532833 | 4.52E-06 | 220103_S_AT, 209134_S_AT, 218027_AT, 200017_AT, 218001_AT, 218890_X_AT, 201665_X_AT, 200088_X_AT, 200099_S_AT, 200809_X_AT, 200029_AT, 218046_S_AT, 200036_S_AT, 217747_S_AT, 218982_S_AT, 200715_X_AT, 213897_S_AT, 218049_S_AT, 200013_AT, 214271_X_AT, 211942_X_AT, 200034_S_AT, 217915_S_AT, 213687_S_AT, 208692_AT, 217559_AT, 221726_AT, 214143_X_AT, 203800_S_AT, 212042_X_AT, 214167_S_AT | 505 | 136 | 6879 | 2.704324 | 0.005927 |

| | | | | | | | | | | |
|---|---|---|---|---|---|---|---|---|---|---|
| GOTERM_BP_DIRECT | GO:0000184~nuclear-tra | 22 | 2.06379 | 4.77E-06 | 209134_S_AT, 200017_AT, 201665_X_AT, 200088_X_AT, 200099_S_AT, 201573_S_AT, 200809_X_AT, 209519_AT, 200029_AT, 200036_S_AT, 217747_S_AT, 200715_X_AT, 209520_S_AT, 218894_S_AT, 217595_AT, 201521_S_AT, 200013_AT, 214271_X_AT, 211942_X_AT, 200034_S_AT, 213687_S_AT, 208692_AT, 221726_AT, 214143_X_AT, 212042_X_AT, 208697_S_AT, 214167_S_AT | 980 | 119 | 16792 | 3.167759 | 0.008719 |
| GOTERM_CC_DIRECT | GO:0022625~cytosolic la | 15 | 1.407129 | 1.62E-05 | 200013_AT, 200088_X_AT, 214271_X_AT, 200809_X_AT, 211942_X_AT, 200029_AT, 217915_S_AT, 200034_S_AT, 220688_S_AT, 209445_X_AT, 213687_S_AT, 217559_AT, 221726_AT, 212018_S_AT, 214143_X_AT, 200036_S_AT, 212042_X_AT, 214167_S_AT, 200715_X_AT | 1004 | 68 | 18224 | 4.003984 | 0.024333 |
| GOTERM_BP_DIRECT | GO:0006614~SRP-depe | 17 | 1.594747 | 9.86E-05 | 209134_S_AT, 200017_AT, 201665_X_AT, 200013_AT, 200088_X_AT, 200099_S_AT, 214271_X_AT, 200809_X_AT, 211942_X_AT, 200029_AT, 200034_S_AT, 213687_S_AT, 217790_S_AT, 208692_AT, 221726_AT, 214143_X_AT, 217747_S_AT, 200036_S_AT, 212042_X_AT, 214167_S_AT, 200715_X_AT | 980 | 94 | 16792 | 3.098828 | 0.180274 |

| | | | | | | | | | | |
|---|---|---|---|---|---|---|---|---|---|---|
| GOTERM_BP_DIRECT | GO:0019083~viral transc | 18 | 1.688555 | 2.57E-04 | 209134_S_AT, 200017_AT, 201558_AT, 201665_X_AT, 200013_AT, 200088_X_AT, 200099_S_AT, 214271_X_AT, 202900_S_AT, 200809_X_AT, 211942_X_AT, 200029_AT, 200034_S_AT, 213687_S_AT, 208692_AT, 221726_AT, 214143_X_AT, 217747_S_AT, 200036_S_AT, 211318_S_AT, 212042_X_AT, 214167_S_AT, 200715_X_AT | 980 | 112 | 16792 | 2.75379 | 0.469146 |
| GOTERM_CC_DIRECT | GO:0005763~mitochond | 7 | 0.65666 | 0.001965 | 220103_S_AT, 218046_S_AT, 218001_AT, 203800_S_AT, 218982_S_AT, 217919_S_AT, 219819_S_AT | 1004 | 25 | 18224 | 5.08239 | 2.9103 |
| GOTERM_BP_DIRECT | GO:0070125~mitochond | 13 | 1.219512 | 0.003705 | 213897_S_AT, 220103_S_AT, 218027_AT, 212145_AT, 218049_S_AT, 218890_X_AT, 218398_AT, 218202_X_AT, 218046_S_AT, 203800_S_AT, 217919_S_AT, 218982_S_AT, 219819_S_AT | 980 | 85 | 16792 | 2.6206 | 6.566023 |
| GOTERM_BP_DIRECT | GO:0070126~mitochond | 13 | 1.219512 | 0.004083 | 213897_S_AT, 220103_S_AT, 218027_AT, 212145_AT, 218049_S_AT, 218890_X_AT, 218398_AT, 218202_X_AT, 218046_S_AT, 203800_S_AT, 217919_S_AT, 218982_S_AT, 219819_S_AT | 980 | 86 | 16792 | 2.590128 | 7.212874 |

| | | | | | | | | | | |
|---|---|---|---|---|---|---|---|---|---|---|
| GOTERM_CC_DIRECT | GO:0005762~mitochond | 5 | 0.469043 | 0.270679 | 213897_S_AT, 218027_AT, 218890_X_AT, 218049_S_AT, 217919_S_AT | 1004 | 48 | 18224 | 1.89077 | 99.12589 |
| Annotation Cluster 4 | Enrichment Score: 5.199792855918165 | | | | | | | | | |
| Category | Term | Count | % | PValue | Genes | List Total | Pop Hits | Pop Total | Fold Enrich | FDR |
| UP_KEYWORDS | Cell cycle | 68 | 6.378987 | 1.91E-08 | 203163_AT, 214999_S_AT, 204641_AT, 203132_AT, 212525_S_AT, 201863_AT, 210379_S_AT, 203565_S_AT, 203068_AT, 203764_AT, 211089_S_AT, 201300_S_AT, 212264_S_AT, 216295_S_AT, 219262_AT, 203856_AT, 205393_S_AT, 201308_S_AT, 215519_X_AT, 214738_S_AT, 202246_S_AT, 203213_AT, 208351_S_AT, 201457_X_AT, 207845_S_AT, 210257_X_AT, 218163_AT, 213116_AT, 204742_S_AT, 221487_S_AT, 203272_S_AT, 207143_AT, 41657_AT, 210285_X_AT, 209714_S_AT, 203137_AT, 200987_X_AT, 213117_AT, 207319_S_AT, 215707_S_AT, 217299_S_AT, 211080_S_AT, 219769_AT, 219273_AT, 211297_S_AT, 200813_S_AT, 202906_S_AT, 207629_S_AT, 211077_S_AT, 209974_S_AT, 213253_AT, 214435_X_AT, 209891_AT, 209464_AT, 218658_S_AT, 221486_AT, 218252_AT, 217852_S_AT, 215282_AT, 200829_X_AT, 214941_S_AT, | 1035 | 650 | 20581 | 2.080279 | 2.71E-05 |
| UP_KEYWORDS | Cell division | 45 | 4.221388 | 4.10E-07 | 214999_S_AT, 203163_AT, 204641_AT, 203068_AT, 211089_S_AT, 212264_S_AT, 216295_S_AT, 203856_AT, 201308_S_AT, 214738_S_AT, 202246_S_AT, 203213_AT, 201457_X_AT, 207845_S_AT, 204742_S_AT, 213116_AT, 221487_S_AT, 207143_AT, 213117_AT, 207319_S_AT, 211080_S_AT, 219273_AT, 211297_S_AT, 219769_AT, 200813_S_AT, 207629_S_AT, 209974_S_AT, 213253_AT, 214435_X_AT, 209464_AT, 209891_AT, 218658_S_AT, 221486_AT, 217852_S_AT, 215282_AT, 200829_X_AT, 214941_S_AT, 200828_S_AT, 211814_S_AT, 219004_S_AT, 201897_S_AT, 201614_S_AT, 211547_S_AT, 203214_X_AT, 201953_AT, 208692_AT, 200712_S_AT, 209484_S_AT, 208945_S_AT, 208796_S_AT, 201179_S_AT, 206248_AT | 1035 | 388 | 20581 | 2.306253 | 5.82E-04 |
| UP_KEYWORDS | Mitosis | 30 | 2.814259 | 5.86E-05 | 213117_AT, 203163_AT, 204641_AT, 211080_S_AT, 219273_AT, 219769_AT, 200813_S_AT, 207629_S_AT, 203068_AT, 209974_S_AT, 213253_AT, 211089_S_AT, 212264_S_AT, 216295_S_AT, 209891_AT, 209464_AT, 203856_AT, 218658_S_AT, 221486_AT, 215282_AT, 214738_S_AT, 203213_AT, 200829_X_AT, 200828_S_AT, 207845_S_AT, 201457_X_AT, 219004_S_AT, 204742_S_AT, 213116_AT, 201614_S_AT, 211547_S_AT, 203214_X_AT, 221487_S_AT, 200712_S_AT, 208692_AT, 209484_S_AT, 208796_S_AT | 1035 | 262 | 20581 | 2.276911 | 0.083156 |

| | | | | | | | | | | |
|---|---|---|---|---|---|---|---|---|---|---|
| GOTERM_BP_DIRECT | GO:0051301~cell divisio | 40 | 3.752345 | 7.85E-05 | 203163_AT, 207319_S_AT, 204641_AT, 203132_AT, 211080_S_AT, 211297_S_AT, 219273_AT, 207629_S_AT, 203068_AT, 209974_S_AT, 213253_AT, 211089_S_AT, 212264_S_AT, 216295_S_AT, 209891_AT, 203856_AT, 218658_S_AT, 201308_S_AT, 221486_AT, 217852_S_AT, 215282_AT, 202246_S_AT, 214738_S_AT, 203213_AT, 211750_X_AT, 214941_S_AT, 200829_X_AT, 200828_S_AT, 207845_S_AT, 201457_X_AT, 219004_S_AT, 211814_S_AT, 213116_AT, 204742_S_AT, 201897_S_AT, 201614_S_AT, 203214_X_AT, 221487_S_AT, 207143_AT, 200712_S_AT, 208692_AT, 201953_AT, 209484_S_AT, 208796_S_AT, 201179_S_AT, 206248_AT | 980 | 350 | 16792 | 1.958251 | 0.143452 |
| GOTERM_BP_DIRECT | GO:0007067~mitotic nuc | 30 | 2.814259 | 2.79E-04 | 209134_S_AT, 213117_AT, 204641_AT, 222118_AT, 211080_S_AT, 219273_AT, 219769_AT, 200813_S_AT, 207629_S_AT, 203068_AT, 211089_S_AT, 216295_S_AT, 209891_AT, 209464_AT, 203856_AT, 218658_S_AT, 221486_AT, 215282_AT, 214738_S_AT, 203213_AT, 219555_S_AT, 207845_S_AT, 210997_AT, 219004_S_AT, 202900_S_AT, 213116_AT, 201614_S_AT, 211547_S_AT, 203214_X_AT, 221487_S_AT, 200712_S_AT, 208692_AT, 219810_AT, 204472_AT, 209484_S_AT, 208796_S_AT | 980 | 248 | 16792 | 2.072745 | 0.509601 |
| Annotation Cluster 5 | Enrichment Score: 4.847881370420967 | | | | | | | | | |
| Category | Term | Count | % | PValue | Genes | List Total | Pop Hits | Pop Total | Fold Enrich | FDR |
| UP_KEYWORDS | Chaperone | 29 | 2.72045 | 9.93E-07 | 200877_AT, 219861_AT, 207040_S_AT, 205362_S_AT, 204560_AT, 202348_S_AT, 205217_AT, 200881_S_AT, 219487_AT, 217911_S_AT, 203405_AT, 208635_X_AT, 200735_X_AT, 200880_AT, 213262_AT, 200873_S_AT, 218408_AT, 40189_AT, 205361_S_AT, 204186_S_AT, 200812_AT, 212432_AT, 208696_AT, 209406_AT, 213047_X_AT, 207618_S_AT, 200630_X_AT, 201870_AT, 200631_S_AT, 210338_S_AT, 201872_S_AT, 208687_X_AT, 205133_S_AT, 215780_S_AT, 209157_AT, 208103_S_AT, 218336_AT | 1035 | 201 | 20581 | 2.868984 | 0.00141 |

| | | | | | | | | | | |
|---|---|---|---|---|---|---|---|---|---|---|
| GOTERM_BP_DIRECT | GO:0006457~protein fol | 29 | 2.72045 | 1.86E-06 | 200877_AT, 208097_S_AT, 201459_AT, 207040_S_AT, 205362_S_AT, 204560_AT, 201490_S_AT, 209078_S_AT, 211797_S_AT, 200881_S_AT, 211765_X_AT, 217911_S_AT, 209208_AT, 200880_AT, 219390_AT, 213262_AT, 209476_AT, 200873_S_AT, 205361_S_AT, 204186_S_AT, 200812_AT, 212432_AT, 76897_S_AT, 211251_X_AT, 208696_AT, 209406_AT, 201270_X_AT, 210338_S_AT, 208687_X_AT, 205133_S_AT, 209157_AT, 200744_S_AT, 201179_S_AT, 218336_AT | 980 | 180 | 16792 | 2.76059 | 0.003409 |
| GOTERM_MF_DIRECT | GO:0051082~unfolded p | 16 | 1.500938 | 0.001546 | 200873_S_AT, 205361_S_AT, 200877_AT, 201459_AT, 207040_S_AT, 212432_AT, 200812_AT, 205362_S_AT, 208696_AT, 202348_S_AT, 201270_X_AT, 210338_S_AT, 211765_X_AT, 200881_S_AT, 208687_X_AT, 205133_S_AT, 209157_AT, 200880_AT, 218336_AT | 971 | 110 | 16881 | 2.528752 | 2.451839 |
| Annotation Cluster 6 | Enrichment Score: 4.396274427968596 | | | | | | | | | |
| Category | Term | Count | % | PValue | Genes | List Total | Pop Hits | Pop Total | Fold Enrich | FDR |
| UP_KEYWORDS | mRNA processing | 44 | 4.12758 | 1.25E-08 | 203103_S_AT, 202190_AT, 205018_S_AT, 209781_S_AT, 219539_AT, 203818_S_AT, 221263_S_AT, 202690_S_AT, 214737_X_AT, 215424_S_AT, 201742_X_AT, 209520_S_AT, 208742_S_AT, 202858_AT, 201521_S_AT, 208910_S_AT, 205017_S_AT, 32723_AT, 202903_AT, 202691_AT, 210338_S_AT, 214214_S_AT, 204207_S_AT, 208687_X_AT, 210285_X_AT, 220671_AT, 216457_S_AT, 203137_AT, 200892_S_AT, 202407_S_AT, 211747_S_AT, 207319_S_AT, 203013_AT, 221547_AT, 202904_S_AT, 200751_S_AT, 209519_AT, 211849_S_AT, 217822_AT, 202126_AT, 201241_AT, 210180_S_AT, 208741_AT, 200014_S_AT, 218894_S_AT, 202899_S_AT, 211932_AT, 201357_S_AT, 214941_S_AT, 210470_X_AT, 204459_AT, 214698_AT, 202697_AT, 216855_S_AT, 204559_S_AT, 213461_AT, 218774_AT, 217821_S_AT, 207158_AT, | 1035 | 332 | 20581 | 2.635365 | 1.77E-05 |

| | | | | | | | | | | |
|---|---|---|---|---|---|---|---|---|---|---|
| GOTERM_BP_DIRECT | GO:0000398~mRNA spl | 36 | 3.377111 | 7.08E-08 | 203103_S_AT, 202190_AT, 209487_AT, 202407_S_AT, 200892_S_AT, 211747_S_AT, 219539_AT, 203818_S_AT, 200751_S_AT, 202904_S_AT, 209519_AT, 202634_AT, 221263_S_AT, 202690_S_AT, 202126_AT, 201726_AT, 214737_X_AT, 210180_S_AT, 215424_S_AT, 201742_X_AT, 200014_S_AT, 209520_S_AT, 201054_AT, 203664_S_AT, 202899_S_AT, 201357_S_AT, 211932_AT, 214941_S_AT, 202858_AT, 201521_S_AT, 209488_S_AT, 210470_X_AT, 204459_AT, 202903_AT, 32723_AT, 202691_AT, 202697_AT, 216855_S_AT, 204559_S_AT, 213887_S_AT, 213461_AT, 209330_S_AT, 210338_S_AT, 208687_X_AT, 209511_AT, 209044_X_AT, 216457_S_AT, 217854_S_AT, 208879_X_AT | 980 | 222 | 16792 | 2.778599 | 1.30E-04 |
| UP_KEYWORDS | mRNA splicing | 36 | 3.377111 | 1.08E-07 | 203103_S_AT, 205018_S_AT, 219539_AT, 203818_S_AT, 221263_S_AT, 202690_S_AT, 214737_X_AT, 215424_S_AT, 201742_X_AT, 209520_S_AT, 208742_S_AT, 202858_AT, 201521_S_AT, 208910_S_AT, 205017_S_AT, 202903_AT, 202691_AT, 210338_S_AT, 214214_S_AT, 208687_X_AT, 210285_X_AT, 216457_S_AT, 203137_AT, 202407_S_AT, 200892_S_AT, 211747_S_AT, 203013_AT, 207319_S_AT, 221547_AT, 202904_S_AT, 200751_S_AT, 209519_AT, 217822_AT, 202126_AT, 210180_S_AT, 208741_AT, 200014_S_AT, 218894_S_AT, 202899_S_AT, 201357_S_AT, 211932_AT, 214941_S_AT, 210470_X_AT, 214698_AT, 216855_S_AT, 204559_S_AT, 218774_AT, 217821_S_AT, 209044_X_AT, 208879_X_AT | 1035 | 260 | 20581 | 2.753311 | 1.54E-04 |
| KEGG_PATHWAY | hsa03040:Spliceosome | 25 | 2.345216 | 2.89E-05 | 203103_S_AT, 200892_S_AT, 202407_S_AT, 211747_S_AT, 221547_AT, 203818_S_AT, 200751_S_AT, 202904_S_AT, 209519_AT, 217822_AT, 221263_S_AT, 202690_S_AT, 214737_X_AT, 210180_S_AT, 215424_S_AT, 201742_X_AT, 200014_S_AT, 209520_S_AT, 218894_S_AT, 202899_S_AT, 211932_AT, 201357_S_AT, 214941_S_AT, 202858_AT, 201521_S_AT, 202903_AT, 202691_AT, 216855_S_AT, 204559_S_AT, 210338_S_AT, 217821_S_AT, 208687_X_AT, 209044_X_AT, 216457_S_AT, 208879_X_AT | 505 | 133 | 6879 | 2.560485 | 0.037866 |

| | | | | | | | | | | |
|---|---|---|---|---|---|---|---|---|---|---|
| UP_KEYWORDS | Spliceosome | 19 | 1.782364 | 6.55E-05 | 203103_S_AT, 202407_S_AT, 211747_S_AT, 221547_AT, 200751_S_AT, 202904_S_AT, 203818_S_AT, 221263_S_AT, 202690_S_AT, 202126_AT, 214737_X_AT, 215424_S_AT, 201742_X_AT, 200014_S_AT, 211932_AT, 201357_S_AT, 202858_AT, 202903_AT, 202691_AT, 216855_S_AT, 204559_S_AT, 210338_S_AT, 208687_X_AT, 209044_X_AT, 216457_S_AT, 208879_X_AT | 1035 | 127 | 20581 | 2.974925 | 0.09292 |
| GOTERM_CC_DIRECT | GO:0071013~catalytic st | 14 | 1.313321 | 0.00154 | 203103_S_AT, 202858_AT, 203818_S_AT, 200751_S_AT, 202691_AT, 216855_S_AT, 204559_S_AT, 202690_S_AT, 202126_AT, 214737_X_AT, 215424_S_AT, 201742_X_AT, 216457_S_AT, 200014_S_AT, 218894_S_AT, 208879_X_AT, 201357_S_AT, 211932_AT | 1004 | 92 | 18224 | 2.762169 | 2.288368 |
| GOTERM_BP_DIRECT | GO:0008380~RNA splici | 19 | 1.782364 | 0.008216 | 205018_S_AT, 203013_AT, 221547_AT, 200751_S_AT, 203818_S_AT, 209519_AT, 217822_AT, 202690_S_AT, 202126_AT, 214737_X_AT, 208741_AT, 200014_S_AT, 209520_S_AT, 218894_S_AT, 208742_S_AT, 202858_AT, 201521_S_AT, 208910_S_AT, 210470_X_AT, 205017_S_AT, 214698_AT, 202691_AT, 214214_S_AT, 217821_S_AT, 210285_X_AT, 209044_X_AT, 203137_AT, 208879_X_AT | 980 | 166 | 16792 | 1.9612 | 14.01098 |

| | | | | | | | | | | |
|---|---|---|---|---|---|---|---|---|---|---|
| GOTERM_BP_DIRECT | GO:0006397~mRNA pro | 20 | 1.876173 | 0.008417 | 209781_S_AT, 205018_S_AT, 211747_S_AT, 203013_AT, 221547_AT, 202904_S_AT, 203818_S_AT, 208741_AT, 201742_X_AT, 218894_S_AT, 201054_AT, 208742_S_AT, 201357_S_AT, 202858_AT, 208910_S_AT, 210470_X_AT, 205017_S_AT, 202903_AT, 214698_AT, 202697_AT, 213461_AT, 214214_S_AT, 210285_X_AT, 207158_AT, 209044_X_AT, 216457_S_AT, 220671_AT, 203137_AT | 980 | 179 | 16792 | 1.914491 | 14.32919 |
| GOTERM_CC_DIRECT | GO:0005681~spliceosor | 12 | 1.125704 | 0.014082 | 201686_X_AT, 202858_AT, 211747_S_AT, 221547_AT, 202903_AT, 203818_S_AT, 200751_S_AT, 202904_S_AT, 217822_AT, 214960_AT, 210338_S_AT, 217821_S_AT, 208687_X_AT, 214959_S_AT, 214737_X_AT, 215424_S_AT, 209044_X_AT, 216457_S_AT, 200014_S_AT, 208879_X_AT, 201357_S_AT | 1004 | 94 | 18224 | 2.317199 | 19.18112 |
| Annotation Cluster 7 | Enrichment Score: 4.34099497275795 | | | | | | | | | |
| Category | Term | Count | % | PValue | Genes | List Total | Pop Hits | Pop Total | Fold Enrich | FDR |
| KEGG_PATHWAY | hsa01200:Carbon metab | 25 | 2.345216 | 1.52E-06 | 217356_S_AT, 208911_S_AT, 214095_AT, 203401_AT, 210153_S_AT, 214437_S_AT, 216591_S_AT, 214687_X_AT, 211150_S_AT, 200947_S_AT, 210131_X_AT, 200946_X_AT, 202026_AT, 216574_S_AT, 200793_S_AT, 210154_AT, 208848_AT, 202004_X_AT, 212973_AT, 218388_AT, 215794_X_AT, 209009_AT, 207071_S_AT, 215772_X_AT, 200978_AT, 200886_S_AT, 214835_S_AT, 212459_X_AT, 220892_S_AT, 200737_AT, 208631_S_AT, 217874_AT, 208630_AT, 200966_X_AT, 221770_AT, 211922_S_AT | 505 | 113 | 6879 | 3.013669 | 0.001994 |

| | | | | | | | | | | |
|---|---|---|---|---|---|---|---|---|---|---|
| UP_KEYWORDS | Tricarboxylic acid cycle | 9 | 0.844278 | 1.41E-05 | 210131_X_AT, 207071_S_AT, 215772_X_AT, 202026_AT, 208911_S_AT, 200978_AT, 214835_S_AT, 200793_S_AT, 216591_S_AT, 202004_X_AT, 217874_AT, 211150_S_AT, 212459_X_AT | 1035 | 24 | 20581 | 7.456884 | 0.02006 |
| GOTERM_BP_DIRECT | GO:0006099~tricarboxyl | 9 | 0.844278 | 1.85E-04 | 210131_X_AT, 207071_S_AT, 215772_X_AT, 202026_AT, 208911_S_AT, 200978_AT, 214835_S_AT, 200793_S_AT, 216591_S_AT, 202004_X_AT, 217874_AT, 211150_S_AT, 212459_X_AT | 980 | 29 | 16792 | 5.317664 | 0.338733 |
| KEGG_PATHWAY | hsa00020:Citrate cycle (T | 9 | 0.844278 | 0.001086 | 210131_X_AT, 207071_S_AT, 215772_X_AT, 202026_AT, 208911_S_AT, 200978_AT, 214835_S_AT, 200793_S_AT, 216591_S_AT, 202004_X_AT, 217874_AT, 211150_S_AT, 212459_X_AT | 505 | 30 | 6879 | 4.086535 | 1.415737 |
| Annotation Cluster 8 | Enrichment Score: 4.047213636078432 | | | | | | | | | |
| Category | Term | Count | % | PValue | Genes | List Total | Pop Hits | Pop Total | Fold Enrich | FDR |

| | | | | | | | | | | |
|---|---|---|---|---|---|---|---|---|---|---|
| GOTERM_CC_DIRECT | GO:0005913~cell-cell ac | 38 | 3.564728 | 2.00E-05 | 208496_X_AT, 200958_S_AT, 202483_S_AT, 201595_S_AT, 210984_X_AT, 208876_S_AT, 208724_S_AT, 212018_S_AT, 201656_AT, 218230_AT, 217911_S_AT, 208758_AT, 201745_AT, 216804_S_AT, 200634_AT, 210317_S_AT, 209679_S_AT, 214007_S_AT, 217717_S_AT, 202550_S_AT, 32699_S_AT, 208878_S_AT, 204969_S_AT, 216283_S_AT, 210338_S_AT, 208743_S_AT, 214143_X_AT, 208687_X_AT, 210466_S_AT, 210076_X_AT, 207791_S_AT, 200966_X_AT, 208697_S_AT, 212398_AT, 208748_S_AT, 206911_AT, 214687_X_AT, 214008_AT, 217724_AT, 217785_S_AT, 213325_AT, 214443_AT, 200873_S_AT, 201593_S_AT, 200013_AT, 200776_S_AT, 203242_S_AT, 217725_X_AT, 200777_S_AT, 201614_S_AT, 200034_S_AT, 208854_S_AT, 200712_S_AT, 209536_S_AT, 209830_S_AT, 215177_S_AT, | 1004 | 323 | 18224 | 2.135458 | 0.030018 |
| GOTERM_MF_DIRECT | GO:0098641~cadherin b | 34 | 3.189493 | 1.39E-04 | 208496_X_AT, 200958_S_AT, 202483_S_AT, 201595_S_AT, 210984_X_AT, 208876_S_AT, 208724_S_AT, 212018_S_AT, 201656_AT, 218230_AT, 217911_S_AT, 208758_AT, 201745_AT, 216804_S_AT, 200634_AT, 210317_S_AT, 214007_S_AT, 217717_S_AT, 202550_S_AT, 208878_S_AT, 204969_S_AT, 210338_S_AT, 208743_S_AT, 214143_X_AT, 208687_X_AT, 210466_S_AT, 210076_X_AT, 207791_S_AT, 200966_X_AT, 208697_S_AT, 212398_AT, 206911_AT, 214687_X_AT, 214008_AT, 217724_AT, 217785_S_AT, 200873_S_AT, 201593_S_AT, 200013_AT, 200776_S_AT, 203242_S_AT, 217725_X_AT, 200777_S_AT, 201614_S_AT, 200034_S_AT, 208854_S_AT, 200712_S_AT, 209536_S_AT, 209830_S_AT, 215177_S_AT, 214483_S_AT, 211681_S_AT, 203243_S_AT | 971 | 290 | 16881 | 2.038261 | 0.223364 |
| GOTERM_BP_DIRECT | GO:0098609~cell-cell ac | 32 | 3.001876 | 2.59E-04 | 208496_X_AT, 206911_AT, 200958_S_AT, 214687_X_AT, 201595_S_AT, 202483_S_AT, 214008_AT, 208876_S_AT, 217724_AT, 208724_S_AT, 212018_S_AT, 217785_S_AT, 218230_AT, 217911_S_AT, 208758_AT, 201745_AT, 216804_S_AT, 201593_S_AT, 200873_S_AT, 210317_S_AT, 200634_AT, 200013_AT, 214007_S_AT, 217717_S_AT, 203242_S_AT, 200776_S_AT, 202550_S_AT, 217725_X_AT, 200777_S_AT, 200034_S_AT, 201614_S_AT, 208854_S_AT, 208878_S_AT, 200712_S_AT, 204969_S_AT, 210338_S_AT, 209536_S_AT, 208743_S_AT, 214143_X_AT, 209830_S_AT, 208687_X_AT, 210466_S_AT, 210076_X_AT, 207791_S_AT, 214483_S_AT, 200966_X_AT, 211681_S_AT, 208697_S_AT, 212398_AT, 203243_S_AT | 980 | 271 | 16792 | 2.023285 | 0.472792 |
| Annotation Cluster 9 | Enrichment Score: 3.992984322400946 | | | | | | | | | |
| Category | Term | Count | % | PValue | Genes | List Total | Pop Hits | Pop Total | Fold Enrich | FDR |

| | | | | | | | | | | |
|---|---|---|---|---|---|---|---|---|---|---|
| UP_KEYWORDS | Protein biosynthesis | 29 | 2.72045 | 2.13E-09 | 204283_AT, 200840_AT, 200005_AT, 200023_S_AT, 201573_S_AT, 218253_S_AT, 211937_AT, 201435_S_AT, 201263_AT, 204300_AT, 200079_S_AT, 200647_X_AT, 211938_AT, 209316_S_AT, 208726_S_AT, 218949_S_AT, 217595_AT, 213907_AT, 201623_S_AT, 202138_X_AT, 208693_S_AT, 219599_AT, 220329_S_AT, 209971_X_AT, 218163_AT, 202232_S_AT, 202823_AT, 200705_S_AT, 215230_X_AT, 217719_AT, 202461_AT, 202824_S_AT, 208697_S_AT, 219575_S_AT, 214446_AT | 1035 | 152 | 20581 | 3.793853 | 3.03E-06 |
| GOTERM_BP_DIRECT | GO:0006446~regulation | 12 | 1.125704 | 3.90E-06 | 206920_S_AT, 200005_AT, 219599_AT, 201521_S_AT, 200023_S_AT, 202232_S_AT, 209519_AT, 211937_AT, 215230_X_AT, 217719_AT, 201241_AT, 202461_AT, 200647_X_AT, 211938_AT, 208697_S_AT, 209520_S_AT | 980 | 36 | 16792 | 5.711565 | 0.007133 |
| GOTERM_CC_DIRECT | GO:0005852~eukaryotic | 8 | 0.750469 | 1.79E-05 | 217719_AT, 200005_AT, 201872_S_AT, 200023_S_AT, 201652_AT, 200647_X_AT, 202232_S_AT, 208697_S_AT, 215230_X_AT | 1004 | 17 | 18224 | 8.541833 | 0.026867 |
| GOTERM_BP_DIRECT | GO:0001731~formation o | 9 | 0.844278 | 2.90E-05 | 217719_AT, 200005_AT, 219599_AT, 200023_S_AT, 218163_AT, 200647_X_AT, 202232_S_AT, 211938_AT, 218253_S_AT, 208697_S_AT, 211937_AT, 215230_X_AT | 980 | 23 | 16792 | 6.70488 | 0.052984 |

| | | | | | | | | | | |
|---|---|---|---|---|---|---|---|---|---|---|
| UP_KEYWORDS | Initiation factor | 12 | 1.125704 | 1.26E-04 | 200005_AT, 219599_AT, 200023_S_AT, 218163_AT, 202232_S_AT, 218253_S_AT, 211937_AT, 215230_X_AT, 201435_S_AT, 217719_AT, 202461_AT, 200647_X_AT, 211938_AT, 208697_S_AT, 208726_S_AT | 1035 | 58 | 20581 | 4.114143 | 0.179357 |
| GOTERM_MF_DIRECT | GO:0003743~translation | 13 | 1.219512 | 1.54E-04 | 200005_AT, 219599_AT, 200023_S_AT, 201652_AT, 218163_AT, 202232_S_AT, 218253_S_AT, 211937_AT, 215230_X_AT, 201435_S_AT, 217719_AT, 202461_AT, 200647_X_AT, 211938_AT, 208726_S_AT, 208697_S_AT | 971 | 61 | 16881 | 3.705036 | 0.247375 |
| GOTERM_CC_DIRECT | GO:0016282~eukaryotic | 6 | 0.562852 | 9.44E-04 | 217719_AT, 200005_AT, 200023_S_AT, 200647_X_AT, 202232_S_AT, 208697_S_AT, 215230_X_AT | 1004 | 15 | 18224 | 7.260558 | 1.408581 |
| GOTERM_CC_DIRECT | GO:0033290~eukaryotic | 6 | 0.562852 | 9.44E-04 | 217719_AT, 200005_AT, 200023_S_AT, 200647_X_AT, 202232_S_AT, 208697_S_AT, 215230_X_AT | 1004 | 15 | 18224 | 7.260558 | 1.408581 |
| GOTERM_MF_DIRECT | GO:0031369~translation | 5 | 0.469043 | 0.029634 | 206920_S_AT, 200023_S_AT, 200647_X_AT, 202232_S_AT, 203664_S_AT, 215230_X_AT | 971 | 21 | 16881 | 4.139326 | 38.28139 |

| | | | | | | | | | | |
|---|---|---|---|---|---|---|---|---|---|---|
| GOTERM_CC_DIRECT | GO:0071541~eukaryotic | 3 | 0.281426 | 0.052831 | 200005_AT, 200023_S_AT, 202232_S_AT | 1004 | 7 | 18224 | 7.779169 | 55.73809 |
| Annotation Cluster 10 | Enrichment Score: 3.385942979887091 | | | | | | | | | |
| Category | Term | Count | % | PValue | Genes | List Total | Pop Hits | Pop Total | Fold Enrich | FDR |
| KEGG_PATHWAY | hsa05016:Huntington's c | 34 | 3.189493 | 2.86E-06 | 213373_S_AT, 211752_S_AT, 208969_AT, 211833_S_AT, 203177_X_AT, 217140_S_AT, 201256_AT, 208972_S_AT, 200086_S_AT, 219545_AT, 216591_S_AT, 201490_S_AT, 207552_AT, 218013_X_AT, 202634_AT, 210131_X_AT, 212600_S_AT, 202026_AT, 203176_S_AT, 201226_AT, 216295_S_AT, 202004_X_AT, 202298_AT, 209223_AT, 203664_S_AT, 203621_AT, 218160_AT, 210411_S_AT, 208074_S_AT, 208478_S_AT, 200615_S_AT, 203613_S_AT, 201322_AT, 203926_X_AT, 208764_S_AT, 213887_S_AT, 213041_S_AT, 208909_AT, 200612_S_AT, 202698_X_AT, 209511_AT, 207686_S_AT, 200883_AT, 208541_X_AT, 201066_AT, 217854_S_AT | 505 | 192 | 6879 | 2.412191 | 0.003748 |
| KEGG_PATHWAY | hsa05010:Alzheimer's di | 31 | 2.908068 | 3.65E-06 | 213373_S_AT, 211752_S_AT, 208969_AT, 202535_AT, 201256_AT, 208972_S_AT, 200086_S_AT, 213688_AT, 219545_AT, 216591_S_AT, 207552_AT, 210131_X_AT, 212600_S_AT, 202026_AT, 204507_S_AT, 201226_AT, 202298_AT, 202004_X_AT, 209223_AT, 203621_AT, 208351_S_AT, 218160_AT, 210411_S_AT, 202457_S_AT, 207827_X_AT, 201322_AT, 203613_S_AT, 203328_X_AT, 208764_S_AT, 203926_X_AT, 213041_S_AT, 208909_AT, 202698_X_AT, 221036_S_AT, 207686_S_AT, 200883_AT, 201187_S_AT, 201066_AT | 505 | 168 | 6879 | 2.513543 | 0.004793 |
| UP_KEYWORDS | Electron transport | 18 | 1.688555 | 2.66E-05 | 211752_S_AT, 208969_AT, 218160_AT, 208097_S_AT, 209276_S_AT, 219545_AT, 201633_S_AT, 216591_S_AT, 203613_S_AT, 209078_S_AT, 210131_X_AT, 212600_S_AT, 202026_AT, 206662_AT, 208909_AT, 201226_AT, 202298_AT, 202004_X_AT, 200883_AT, 209223_AT, 201066_AT, 209476_AT, 203621_AT | 1035 | 108 | 20581 | 3.314171 | 0.037724 |

| | | | | | | | | | | |
|---|---|---|---|---|---|---|---|---|---|---|
| KEGG_PATHWAY | hsa05012:Parkinson's di | 26 | 2.439024 | 3.05E-05 | 211752_S_AT, 208969_AT, 209141_AT, 217140_S_AT, 201256_AT, 208972_S_AT, 200086_S_AT, 219545_AT, 216591_S_AT, 201490_S_AT, 207552_AT, 210131_X_AT, 212600_S_AT, 202026_AT, 202742_S_AT, 201226_AT, 202298_AT, 202004_X_AT, 209223_AT, 203621_AT, 218160_AT, 207827_X_AT, 201322_AT, 203613_S_AT, 208764_S_AT, 203926_X_AT, 213041_S_AT, 208909_AT, 202698_X_AT, 200883_AT, 201066_AT, 201179_S_AT | 505 | 142 | 6879 | 2.494129 | 0.039996 |
| GOTERM_BP_DIRECT | GO:0032981~mitochond | 13 | 1.219512 | 2.43E-04 | 218225_AT, 209445_X_AT, 208969_AT, 211752_S_AT, 218160_AT, 201226_AT, 202298_AT, 203613_S_AT, 209223_AT, 219006_AT, 207618_S_AT, 209177_AT, 203621_AT | 980 | 63 | 16792 | 3.53573 | 0.444135 |
| UP_KEYWORDS | Respiratory chain | 12 | 1.125704 | 2.72E-04 | 211752_S_AT, 208969_AT, 218160_AT, 219545_AT, 203613_S_AT, 212600_S_AT, 208909_AT, 201226_AT, 202298_AT, 200883_AT, 209223_AT, 201066_AT, 203621_AT | 1035 | 63 | 20581 | 3.787624 | 0.385329 |

| | | | | | | | | | | |
|---|---|---|---|---|---|---|---|---|---|---|
| KEGG_PATHWAY | hsa00190:Oxidative pho | 22 | 2.06379 | 6.12E-04 | 208969_AT, 211752_S_AT, 221550_AT, 208972_S_AT, 201256_AT, 200086_S_AT, 219545_AT, 216591_S_AT, 207552_AT, 210131_X_AT, 212600_S_AT, 202026_AT, 201226_AT, 202004_X_AT, 202298_AT, 209223_AT, 203621_AT, 218160_AT, 201322_AT, 203613_S_AT, 203926_X_AT, 208764_S_AT, 213041_S_AT, 208909_AT, 202698_X_AT, 208899_X_AT, 200883_AT, 201066_AT | 505 | 133 | 6879 | 2.253227 | 0.800153 |
| KEGG_PATHWAY | hsa04932:Non-alcoholic | 23 | 2.157598 | 0.001383 | 213373_S_AT, 217909_S_AT, 211833_S_AT, 208969_AT, 211752_S_AT, 201256_AT, 208640_AT, 200086_S_AT, 219545_AT, 216591_S_AT, 201805_AT, 210131_X_AT, 212600_S_AT, 202026_AT, 201226_AT, 209799_AT, 202004_X_AT, 202298_AT, 209239_AT, 209223_AT, 203621_AT, 218160_AT, 208478_S_AT, 203613_S_AT, 208909_AT, 202698_X_AT, 207686_S_AT, 200883_AT, 201066_AT | 505 | 151 | 6879 | 2.074841 | 1.799034 |
| GOTERM_CC_DIRECT | GO:0005747~mitochond | 9 | 0.844278 | 0.00494 | 208969_AT, 211752_S_AT, 218160_AT, 201226_AT, 202298_AT, 207827_X_AT, 203613_S_AT, 209223_AT, 203621_AT | 1004 | 49 | 18224 | 3.33393 | 7.167036 |
| GOTERM_MF_DIRECT | GO:0008137~NADH deh | 9 | 0.844278 | 0.00561 | 208969_AT, 211752_S_AT, 218160_AT, 201226_AT, 219545_AT, 202298_AT, 203613_S_AT, 209223_AT, 203621_AT | 971 | 48 | 16881 | 3.259719 | 8.629687 |

| | | | | | | | | | | |
|---|---|---|---|---|---|---|---|---|---|---|
| GOTERM_BP_DIRECT | GO:0006120~mitochond | 9 | 0.844278 | 0.006942 | 208969_AT, 211752_S_AT, 218160_AT, 201226_AT, 219545_AT, 202298_AT, 203613_S_AT, 209223_AT, 203621_AT | 980 | 49 | 16792 | 3.147189 | 11.96778 |
| UP_KEYWORDS | Ubiquinone | 4 | 0.375235 | 0.256456 | 211752_S_AT, 218160_AT, 202298_AT, 203621_AT | 1035 | 35 | 20581 | 2.272574 | 98.51236 |
| Annotation Cluster 11 | Enrichment Score: 3.1625954587923193 | | | | | | | | | |
| Category | Term | Count | % | PValue | Genes | List Total | Pop Hits | Pop Total | Fold Enrich | FDR |
| GOTERM_BP_DIRECT | GO:0043488~regulation | 24 | 2.251407 | 1.93E-08 | 200017_AT, 201532_AT, 201198_S_AT, 201114_X_AT, 221573_AT, 202243_S_AT, 216088_S_AT, 217724_AT, 201316_AT, 218481_AT, 91684_G_AT, 210759_S_AT, 206923_AT, 201726_AT, 201400_AT, 201676_X_AT, 40189_AT, 201317_S_AT, 202753_AT, 217717_S_AT, 217725_X_AT, 201461_S_AT, 202352_S_AT, 213047_X_AT, 200630_X_AT, 200631_S_AT, 210027_S_AT, 210338_S_AT, 201699_AT, 209330_S_AT, 208743_S_AT, 209853_S_AT, 208687_X_AT, 210466_S_AT, 210076_X_AT, 215780_S_AT, 53202_AT, 215195_AT, 200987_X_AT, 220457_AT | 980 | 103 | 16792 | 3.99255 | 3.53E-05 |
| UP_KEYWORDS | Proteasome | 14 | 1.313321 | 1.55E-06 | 201676_X_AT, 201317_S_AT, 202753_AT, 201532_AT, 201198_S_AT, 201114_X_AT, 221573_AT, 202243_S_AT, 201222_S_AT, 202352_S_AT, 216088_S_AT, 219960_S_AT, 201316_AT, 201699_AT, 210759_S_AT, 209853_S_AT, 201281_AT, 53202_AT, 200987_X_AT, 201400_AT | 1035 | 53 | 20581 | 5.252648 | 0.002206 |

| | | | | | | | | | | |
|---|---|---|---|---|---|---|---|---|---|---|
| GOTERM_BP_DIRECT | GO:0031145~anaphase- | 18 | 1.688555 | 2.34E-06 | 200017_AT, 201532_AT, 201198_S_AT, 201114_X_AT, 202243_S_AT, 221573_AT, 216088_S_AT, 201316_AT, 209974_S_AT, 210759_S_AT, 214590_S_AT, 209464_AT, 203626_S_AT, 201400_AT, 203213_AT, 201676_X_AT, 201317_S_AT, 202753_AT, 207845_S_AT, 201457_X_AT, 202352_S_AT, 203214_X_AT, 201699_AT, 209853_S_AT, 53202_AT, 200987_X_AT | 980 | 79 | 16792 | 3.904107 | 0.004284 |
| GOTERM_BP_DIRECT | GO:0051436~negative re | 16 | 1.500938 | 1.16E-05 | 201676_X_AT, 203213_AT, 201317_S_AT, 202753_AT, 200017_AT, 201457_X_AT, 201532_AT, 207845_S_AT, 201114_X_AT, 201198_S_AT, 221573_AT, 202243_S_AT, 202352_S_AT, 216088_S_AT, 201316_AT, 203214_X_AT, 209974_S_AT, 201699_AT, 210759_S_AT, 209853_S_AT, 214590_S_AT, 53202_AT, 200987_X_AT, 201400_AT | 980 | 71 | 16792 | 3.861339 | 0.021157 |
| GOTERM_BP_DIRECT | GO:0051437~positive re | 16 | 1.500938 | 2.73E-05 | 201676_X_AT, 203213_AT, 201317_S_AT, 202753_AT, 200017_AT, 201457_X_AT, 201532_AT, 207845_S_AT, 201114_X_AT, 201198_S_AT, 221573_AT, 202243_S_AT, 202352_S_AT, 216088_S_AT, 201316_AT, 203214_X_AT, 209974_S_AT, 201699_AT, 210759_S_AT, 209853_S_AT, 214590_S_AT, 53202_AT, 200987_X_AT, 201400_AT | 980 | 76 | 16792 | 3.607304 | 0.049981 |

| | | | | | | | | | | |
|---|---|---|---|---|---|---|---|---|---|---|
| GOTERM_CC_DIRECT | GO:0000502~proteasom | 13 | 1.219512 | 8.71E-05 | 201676_X_AT, 201317_S_AT, 202753_AT, 201532_AT, 201198_S_AT, 201114_X_AT, 221573_AT, 202243_S_AT, 201222_S_AT, 202352_S_AT, 216088_S_AT, 201316_AT, 201699_AT, 210759_S_AT, 209853_S_AT, 201281_AT, 53202_AT, 200987_X_AT, 201400_AT | 1004 | 60 | 18224 | 3.932802 | 0.130757 |
| GOTERM_BP_DIRECT | GO:0038061~NIK/NF-ka | 14 | 1.313321 | 9.47E-05 | 201676_X_AT, 201317_S_AT, 202753_AT, 200017_AT, 201532_AT, 201198_S_AT, 201114_X_AT, 221573_AT, 202243_S_AT, 202352_S_AT, 216088_S_AT, 201316_AT, 203109_AT, 201699_AT, 210759_S_AT, 209853_S_AT, 53202_AT, 209239_AT, 200987_X_AT, 201400_AT | 980 | 66 | 16792 | 3.634632 | 0.173121 |
| GOTERM_BP_DIRECT | GO:0006521~regulation | 12 | 1.125704 | 1.38E-04 | 201676_X_AT, 201317_S_AT, 202753_AT, 215952_S_AT, 201532_AT, 200077_S_AT, 201198_S_AT, 201114_X_AT, 221573_AT, 202243_S_AT, 202352_S_AT, 216088_S_AT, 201316_AT, 201699_AT, 210759_S_AT, 209853_S_AT, 53202_AT, 200987_X_AT, 201400_AT | 980 | 51 | 16792 | 4.031693 | 0.252628 |

| | | | | | | | | | | |
|---|---|---|---|---|---|---|---|---|---|---|
| GOTERM_BP_DIRECT | GO:0060071~Wnt signal | 16 | 1.500938 | 2.63E-04 | 201676_X_AT, 201317_S_AT, 202753_AT, 200017_AT, 200634_AT, 208074_S_AT, 201532_AT, 208640_AT, 201114_X_AT, 201198_S_AT, 221573_AT, 200615_S_AT, 202243_S_AT, 202352_S_AT, 216088_S_AT, 201316_AT, 201699_AT, 210759_S_AT, 209853_S_AT, 200612_S_AT, 53202_AT, 200987_X_AT, 201400_AT | 980 | 92 | 16792 | 2.979947 | 0.480984 |
| GOTERM_BP_DIRECT | GO:0002223~stimulatory | 17 | 1.594747 | 3.69E-04 | 201676_X_AT, 201317_S_AT, 202753_AT, 200017_AT, 201532_AT, 201198_S_AT, 201114_X_AT, 221573_AT, 209615_S_AT, 202243_S_AT, 202352_S_AT, 208876_S_AT, 216088_S_AT, 201316_AT, 208878_S_AT, 201699_AT, 210759_S_AT, 202742_S_AT, 209853_S_AT, 214590_S_AT, 53202_AT, 209239_AT, 200987_X_AT, 201400_AT | 980 | 105 | 16792 | 2.774189 | 0.673116 |
| GOTERM_BP_DIRECT | GO:0050852~T cell rece | 20 | 1.876173 | 9.91E-04 | 209671_X_AT, 200017_AT, 201532_AT, 201198_S_AT, 201114_X_AT, 221573_AT, 202243_S_AT, 209615_S_AT, 216088_S_AT, 208876_S_AT, 201316_AT, 210759_S_AT, 215666_AT, 214590_S_AT, 209239_AT, 201400_AT, 201676_X_AT, 208351_S_AT, 201317_S_AT, 202753_AT, 202352_S_AT, 208878_S_AT, 201699_AT, 41657_AT, 209853_S_AT, 53202_AT, 200987_X_AT | 980 | 148 | 16792 | 2.315499 | 1.797483 |

| | | | | | | | | | | |
|---|---|---|---|---|---|---|---|---|---|---|
| KEGG_PATHWAY | hsa03050:Proteasome | 11 | 1.031895 | 0.001042 | 201676_X_AT, 201317_S_AT, 202753_AT, 201532_AT, 201198_S_AT, 201114_X_AT, 221573_AT, 202243_S_AT, 202352_S_AT, 216088_S_AT, 201316_AT, 201699_AT, 210759_S_AT, 209853_S_AT, 53202_AT, 200987_X_AT, 201400_AT | 505 | 44 | 6879 | 3.405446 | 1.358742 |
| GOTERM_BP_DIRECT | GO:0000209~protein po | 23 | 2.157598 | 0.001084 | 203103_S_AT, 200017_AT, 201532_AT, 201198_S_AT, 201114_X_AT, 221573_AT, 202243_S_AT, 216088_S_AT, 200868_S_AT, 201316_AT, 210759_S_AT, 214590_S_AT, 203626_S_AT, 206788_S_AT, 200669_S_AT, 201400_AT, 201676_X_AT, 201317_S_AT, 201823_S_AT, 202753_AT, 202352_S_AT, 218738_S_AT, 201699_AT, 209853_S_AT, 53202_AT, 203409_AT, 209096_AT, 200987_X_AT, 218585_S_AT | 980 | 184 | 16792 | 2.141837 | 1.96423 |
| GOTERM_BP_DIRECT | GO:0090090~negative re | 21 | 1.969981 | 0.001303 | 200017_AT, 201532_AT, 201198_S_AT, 201114_X_AT, 221573_AT, 202243_S_AT, 216088_S_AT, 201316_AT, 201278_AT, 210759_S_AT, 213139_AT, 201400_AT, 201676_X_AT, 201317_S_AT, 222142_AT, 202753_AT, 201279_S_AT, 219931_S_AT, 201508_AT, 204336_S_AT, 202352_S_AT, 205411_AT, 201699_AT, 210757_X_AT, 209853_S_AT, 201503_AT, 218850_S_AT, 53202_AT, 201280_S_AT, 200987_X_AT | 980 | 163 | 16792 | 2.207537 | 2.35808 |

| | | | | | | | | | | |
|---|---|---|---|---|---|---|---|---|---|---|
| GOTERM_BP_DIRECT | GO:0090263~positive re | 17 | 1.594747 | 0.001619 | 201676_X_AT, 201317_S_AT, 208229_AT, 202753_AT, 200017_AT, 213880_AT, 201532_AT, 201198_S_AT, 201114_X_AT, 221573_AT, 202243_S_AT, 202352_S_AT, 216088_S_AT, 201316_AT, 206536_S_AT, 201234_AT, 201699_AT, 210759_S_AT, 209853_S_AT, 53202_AT, 209239_AT, 200987_X_AT, 201400_AT | 980 | 120 | 16792 | 2.427415 | 2.921128 |
| GOTERM_BP_DIRECT | GO:0038095~Fc-epsilon | 22 | 2.06379 | 0.001646 | 200017_AT, 201532_AT, 208640_AT, 213688_AT, 201198_S_AT, 201114_X_AT, 221573_AT, 202243_S_AT, 209615_S_AT, 216088_S_AT, 208876_S_AT, 201316_AT, 210759_S_AT, 204507_S_AT, 214590_S_AT, 209239_AT, 201400_AT, 201676_X_AT, 201317_S_AT, 208351_S_AT, 202753_AT, 203265_S_AT, 202457_S_AT, 202352_S_AT, 208878_S_AT, 201699_AT, 209853_S_AT, 53202_AT, 200987_X_AT | 980 | 178 | 16792 | 2.117771 | 2.96893 |
| GOTERM_BP_DIRECT | GO:0043161~proteasom | 24 | 2.251407 | 0.001745 | 218569_S_AT, 200017_AT, 201532_AT, 210932_S_AT, 201198_S_AT, 207722_S_AT, 201114_X_AT, 221573_AT, 202243_S_AT, 216088_S_AT, 201316_AT, 209974_S_AT, 210759_S_AT, 214590_S_AT, 200669_S_AT, 201400_AT, 201676_X_AT, 203213_AT, 201317_S_AT, 202753_AT, 218832_X_AT, 208619_AT, 201457_X_AT, 207845_S_AT, 201877_S_AT, 201222_S_AT, 202352_S_AT, 203214_X_AT, 201699_AT, 209853_S_AT, 53202_AT, 200987_X_AT, 203403_S_AT | 980 | 203 | 16792 | 2.025777 | 3.145333 |

| | | | | | | | | | | |
|---|---|---|---|---|---|---|---|---|---|---|
| UP_KEYWORDS | Threonine protease | 6 | 0.562852 | 0.002612 | 201676_X_AT, 201317_S_AT, 210759_S_AT, 201532_AT, 53202_AT, 201114_X_AT, 221573_AT, 202243_S_AT, 216088_S_AT, 201316_AT, 201400_AT | 1035 | 20 | 20581 | 5.965507 | 3.645199 |
| INTERPRO | IPR001353:Proteasome, | 6 | 0.562852 | 0.00286 | 201676_X_AT, 201317_S_AT, 210759_S_AT, 201532_AT, 53202_AT, 201114_X_AT, 221573_AT, 202243_S_AT, 216088_S_AT, 201316_AT, 201400_AT | 1008 | 19 | 18559 | 5.814223 | 4.747015 |
| GOTERM_BP_DIRECT | GO:0002479~antigen pr | 11 | 1.031895 | 0.003305 | 201676_X_AT, 201317_S_AT, 202753_AT, 201532_AT, 201198_S_AT, 201114_X_AT, 221573_AT, 202243_S_AT, 202352_S_AT, 216088_S_AT, 201316_AT, 201699_AT, 210759_S_AT, 209853_S_AT, 53202_AT, 200987_X_AT, 201400_AT | 980 | 63 | 16792 | 2.991772 | 5.877919 |
| GOTERM_CC_DIRECT | GO:0005839~proteasom | 6 | 0.562852 | 0.004859 | 201676_X_AT, 201317_S_AT, 210759_S_AT, 201532_AT, 53202_AT, 201114_X_AT, 221573_AT, 202243_S_AT, 216088_S_AT, 201316_AT, 201400_AT | 1004 | 21 | 18224 | 5.186113 | 7.052526 |

| | | | | | | | | | | |
|---|---|---|---|---|---|---|---|---|---|---|
| SMART | SM00948:SM00948 | 4 | 0.375235 | 0.005766 | 201676_X_AT, 201317_S_AT, 210759_S_AT, 201532_AT, 53202_AT, 201114_X_AT, 221573_AT, 216088_S_AT, 201316_AT | 504 | 8 | 10057 | 9.977183 | 7.5732 |
| GOTERM_MF_DIRECT | GO:0004298~threonine- | 6 | 0.562852 | 0.00583 | 201676_X_AT, 201317_S_AT, 210759_S_AT, 201532_AT, 53202_AT, 201114_X_AT, 221573_AT, 202243_S_AT, 216088_S_AT, 201316_AT, 201400_AT | 971 | 21 | 16881 | 4.967191 | 8.953996 |
| INTERPRO | IPR023332:Proteasome | 4 | 0.375235 | 0.00726 | 201676_X_AT, 201317_S_AT, 210759_S_AT, 201532_AT, 53202_AT, 201114_X_AT, 221573_AT, 216088_S_AT, 201316_AT | 1008 | 8 | 18559 | 9.205853 | 11.639 |
| INTERPRO | IPR000426:Proteasome, | 4 | 0.375235 | 0.00726 | 201676_X_AT, 201317_S_AT, 210759_S_AT, 201532_AT, 53202_AT, 201114_X_AT, 221573_AT, 216088_S_AT, 201316_AT | 1008 | 8 | 18559 | 9.205853 | 11.639 |

| | | | | | | | | | | |
|---|---|---|---|---|---|---|---|---|---|---|
| GOTERM_CC_DIRECT | GO:0019773~proteasom | 4 | 0.375235 | 0.007554 | 201676_X_AT, 201317_S_AT, 210759_S_AT, 201532_AT, 53202_AT, 201114_X_AT, 221573_AT, 216088_S_AT, 201316_AT | 1004 | 8 | 18224 | 9.075697 | 10.76223 |
| GOTERM_BP_DIRECT | GO:0000165~MAPK cas | 25 | 2.345216 | 0.019128 | 208229_AT, 200017_AT, 201532_AT, 213688_AT, 206220_S_AT, 201198_S_AT, 201114_X_AT, 221573_AT, 210984_X_AT, 209615_S_AT, 202243_S_AT, 216088_S_AT, 201316_AT, 210759_S_AT, 213372_AT, 209690_S_AT, 212843_AT, 201400_AT, 201676_X_AT, 201317_S_AT, 208351_S_AT, 202753_AT, 210411_S_AT, 217717_S_AT, 201461_S_AT, 211499_S_AT, 202352_S_AT, 201699_AT, 208743_S_AT, 209853_S_AT, 53202_AT, 200987_X_AT | 980 | 262 | 16792 | 1.63499 | 29.76871 |
| GOTERM_BP_DIRECT | GO:0033209~tumor necr | 14 | 1.313321 | 0.019991 | 201676_X_AT, 201317_S_AT, 202753_AT, 200017_AT, 210654_AT, 201532_AT, 201198_S_AT, 201114_X_AT, 221573_AT, 202243_S_AT, 202352_S_AT, 216088_S_AT, 210405_X_AT, 201316_AT, 201699_AT, 210759_S_AT, 209853_S_AT, 53202_AT, 200987_X_AT, 201400_AT | 980 | 118 | 16792 | 2.03293 | 30.8906 |
| GOTERM_BP_DIRECT | GO:0051603~proteolysis | 8 | 0.750469 | 0.020387 | 213373_S_AT, 201676_X_AT, 201317_S_AT, 201532_AT, 201114_X_AT, 221573_AT, 202243_S_AT, 216088_S_AT, 203328_X_AT, 201316_AT, 210759_S_AT, 53202_AT, 207686_S_AT, 201400_AT | 980 | 48 | 16792 | 2.855782 | 31.39969 |

| | | | | | | | | | | |
|---|---|---|---|---|---|---|---|---|---|---|
| GOTERM_CC_DIRECT | GO:0022624~proteasom | 4 | 0.375235 | 0.063482 | 201699_AT, 202753_AT, 201198_S_AT, 202352_S_AT | 1004 | 17 | 18224 | 4.270916 | 62.651 |
| Annotation Cluster 12 | Enrichment Score: 3.1052966921105227 | | | | | | | | | |
| Category | Term | Count | % | PValue | Genes | List Total | Pop Hits | Pop Total | Fold Enrich | FDR |
| GOTERM_BP_DIRECT | GO:0006368~transcriptio | 18 | 1.688555 | 7.93E-06 | 201521_S_AT, 204093_AT, 202823_AT, 213468_AT, 209519_AT, 203565_S_AT, 211297_S_AT, 219273_AT, 202634_AT, 203198_AT, 213887_S_AT, 200957_S_AT, 221618_S_AT, 201281_AT, 209511_AT, 202824_S_AT, 209520_S_AT, 217854_S_AT, 214446_AT, 203664_S_AT, 221094_S_AT | 980 | 86 | 16792 | 3.586331 | 0.014508 |
| GOTERM_BP_DIRECT | GO:0006370~7-methylg | 11 | 1.031895 | 1.17E-05 | 201521_S_AT, 204093_AT, 213468_AT, 209519_AT, 203565_S_AT, 211297_S_AT, 211849_S_AT, 202634_AT, 213887_S_AT, 204207_S_AT, 209511_AT, 209520_S_AT, 217854_S_AT, 203664_S_AT | 980 | 33 | 16792 | 5.711565 | 0.021417 |
| GOTERM_BP_DIRECT | GO:0010467~gene expr | 12 | 1.125704 | 7.70E-05 | 201521_S_AT, 200751_S_AT, 209519_AT, 216855_S_AT, 202634_AT, 213887_S_AT, 209330_S_AT, 214737_X_AT, 209511_AT, 210285_X_AT, 200014_S_AT, 203137_AT, 209520_S_AT, 217854_S_AT, 201054_AT, 203664_S_AT, 211932_AT | 980 | 48 | 16792 | 4.283673 | 0.140789 |

| | | | | | | | | | | |
|---|---|---|---|---|---|---|---|---|---|---|
| GOTERM_BP_DIRECT | GO:0042795~snRNA tra | 12 | 1.125704 | 0.002286 | 201521_S_AT, 209519_AT, 211297_S_AT, 219273_AT, 218616_AT, 202634_AT, 203198_AT, 203941_AT, 213887_S_AT, 209511_AT, 209520_S_AT, 217854_S_AT, 203664_S_AT, 214446_AT | 980 | 70 | 16792 | 2.937376 | 4.101806 |
| GOTERM_BP_DIRECT | GO:0006367~transcriptio | 18 | 1.688555 | 0.007444 | 212385_AT, 204093_AT, 209101_AT, 213468_AT, 203565_S_AT, 211297_S_AT, 207304_AT, 202634_AT, 221803_S_AT, 203198_AT, 213887_S_AT, 213696_S_AT, 221618_S_AT, 209511_AT, 207978_S_AT, 215424_S_AT, 221938_X_AT, 217854_S_AT, 203664_S_AT | 980 | 152 | 16792 | 2.029108 | 12.77802 |
| GOTERM_BP_DIRECT | GO:0008543~fibroblast g | 11 | 1.031895 | 0.020579 | 208351_S_AT, 208229_AT, 200017_AT, 201521_S_AT, 209101_AT, 200927_S_AT, 209519_AT, 202634_AT, 213887_S_AT, 209511_AT, 209520_S_AT, 217854_S_AT, 203664_S_AT | 980 | 82 | 16792 | 2.298556 | 31.64459 |
| GOTERM_CC_DIRECT | GO:0005665~DNA-direc | 4 | 0.375235 | 0.073175 | 213887_S_AT, 209511_AT, 217854_S_AT, 202634_AT, 203664_S_AT | 1004 | 18 | 18224 | 4.033643 | 68.05265 |
| Annotation Cluster 13 | Enrichment Score: 3.093284945768214 | | | | | | | | | |
| Category | Term | Count | % | PValue | Genes | List Total | Pop Hits | Pop Total | Fold Enrich | FDR |

| | | | | | | | | | | |
|---|---|---|---|---|---|---|---|---|---|---|
| UP_SEQ_FEATURE | domain:PCI | 7 | 0.65666 | 2.72E-04 | 202753_AT, 202467_S_AT, 202141_S_AT, 200647_X_AT, 202232_S_AT, 202143_S_AT, 208697_S_AT, 202352_S_AT, 202142_AT, 215230_X_AT | 1029 | 19 | 20063 | 7.183315 | 0.475049 |
| SMART | SM00088:PINT | 6 | 0.562852 | 8.46E-04 | 202753_AT, 202467_S_AT, 200647_X_AT, 202232_S_AT, 208697_S_AT, 202352_S_AT, 215230_X_AT | 504 | 16 | 10057 | 7.482887 | 1.145882 |
| GOTERM_CC_DIRECT | GO:0016282~eukaryotic | 6 | 0.562852 | 9.44E-04 | 217719_AT, 200005_AT, 200023_S_AT, 200647_X_AT, 202232_S_AT, 208697_S_AT, 215230_X_AT | 1004 | 15 | 18224 | 7.260558 | 1.408581 |
| GOTERM_CC_DIRECT | GO:0033290~eukaryotic | 6 | 0.562852 | 9.44E-04 | 217719_AT, 200005_AT, 200023_S_AT, 200647_X_AT, 202232_S_AT, 208697_S_AT, 215230_X_AT | 1004 | 15 | 18224 | 7.260558 | 1.408581 |
| INTERPRO | IPR000717:Proteasome | 6 | 0.562852 | 0.001666 | 202753_AT, 202467_S_AT, 200647_X_AT, 202232_S_AT, 208697_S_AT, 202352_S_AT, 215230_X_AT | 1008 | 17 | 18559 | 6.498249 | 2.791789 |
| Annotation Cluster 14 | Enrichment Score: 2.836978222032817 | | | | | | | | | |
| Category | Term | Count | % | PValue | Genes | List Total | Pop Hits | Pop Total | Fold Enrich | FDR |

| | | | | | | | | | | |
|---|---|---|---|---|---|---|---|---|---|---|
| GOTERM_MF_DIRECT | GO:0019003~GDP bindi | 14 | 1.313321 | 8.41E-06 | 208734_X_AT, 215772_X_AT, 214835_S_AT, 203401_AT, 206113_S_AT, 207495_AT, 200927_S_AT, 218669_AT, 212459_X_AT, 208732_AT, 204593_S_AT, 221960_S_AT, 208733_AT, 204472_AT, 214435_X_AT, 205461_AT, 200833_S_AT, 217852_S_AT, 201179_S_AT | 971 | 54 | 16881 | 4.507266 | 0.013491 |
| GOTERM_MF_DIRECT | GO:0003924~GTPase ac | 31 | 2.908068 | 3.16E-05 | 208640_AT, 206113_S_AT, 209313_AT, 221737_AT, 218238_AT, 208724_S_AT, 221987_S_AT, 208733_AT, 214435_X_AT, 205461_AT, 201288_AT, 213603_S_AT, 201096_S_AT, 209316_S_AT, 211622_S_AT, 217852_S_AT, 217595_AT, 211750_X_AT, 208734_X_AT, 201921_AT, 204115_AT, 218156_S_AT, 206917_AT, 200927_S_AT, 207495_AT, 208732_AT, 203105_S_AT, 207419_S_AT, 212214_AT, 219357_AT, 221960_S_AT, 204472_AT, 212724_AT, 212213_X_AT, 207791_S_AT, 200833_S_AT, 200744_S_AT, 201179_S_AT | 971 | 234 | 16881 | 2.303164 | 0.050603 |
| UP_KEYWORDS | GTP-binding | 35 | 3.283302 | 1.30E-04 | 208640_AT, 206113_S_AT, 221737_AT, 209313_AT, 218669_AT, 200947_S_AT, 211849_S_AT, 218238_AT, 208724_S_AT, 200946_X_AT, 208733_AT, 214435_X_AT, 205461_AT, 213603_S_AT, 201096_S_AT, 209316_S_AT, 201308_S_AT, 211622_S_AT, 217852_S_AT, 217595_AT, 211750_X_AT, 208734_X_AT, 215772_X_AT, 214835_S_AT, 219622_AT, 206917_AT, 200927_S_AT, 207495_AT, 212459_X_AT, 208732_AT, 203105_S_AT, 207419_S_AT, 212214_AT, 219357_AT, 221960_S_AT, 204472_AT, 204207_S_AT, 212724_AT, 212213_X_AT, 203267_S_AT, 207791_S_AT, 217874_AT, 200833_S_AT, 202810_AT, 201179_S_AT | 1035 | 343 | 20581 | 2.029084 | 0.184993 |

| | | | | | | | | | | |
|---|---|---|---|---|---|---|---|---|---|---|
| UP_KEYWORDS | Prenylation | 21 | 1.969981 | 3.04E-04 | 206113_S_AT, 208640_AT, 201707_AT, 204528_S_AT, 218669_AT, 208752_X_AT, 208724_S_AT, 217785_S_AT, 208733_AT, 200881_S_AT, 214435_X_AT, 205461_AT, 213603_S_AT, 200880_AT, 213864_S_AT, 208734_X_AT, 201921_AT, 204115_AT, 219622_AT, 207495_AT, 200927_S_AT, 208732_AT, 207419_S_AT, 41657_AT, 221960_S_AT, 212724_AT, 207791_S_AT, 212967_X_AT, 201706_S_AT, 209157_AT, 200833_S_AT | 1035 | 168 | 20581 | 2.485628 | 0.431453 |
| GOTERM_MF_DIRECT | GO:0005525~GTP bindi | 40 | 3.752345 | 3.98E-04 | 200947_S_AT, 208724_S_AT, 200946_X_AT, 213603_S_AT, 201308_S_AT, 217595_AT, 207495_AT, 208732_AT, 203105_S_AT, 221960_S_AT, 204207_S_AT, 207791_S_AT, 217874_AT, 200833_S_AT, 209313_AT, 221737_AT, 206113_S_AT, 208640_AT, 218669_AT, 211849_S_AT, 218238_AT, 201577_AT, 221987_S_AT, 208733_AT, 214435_X_AT, 205461_AT, 201096_S_AT, 209316_S_AT, 211622_S_AT, 217852_S_AT, 208734_X_AT, 215794_X_AT, 211750_X_AT, 215772_X_AT, 218156_S_AT, 214835_S_AT, 219622_AT, 206917_AT, 200927_S_AT, 212459_X_AT, 207419_S_AT, 212214_AT, 219357_AT, 209536_S_AT, 204472_AT, 212213_X_AT, 212724_AT, 202461_AT, 203267_S_AT, 202810_AT, 201179_S_AT | 971 | 384 | 16881 | 1.810955 | 0.635987 |
| UP_SEQ_FEATURE | nucleotide phosphate-bi | 30 | 2.814259 | 0.001339 | 208640_AT, 221737_AT, 206113_S_AT, 218669_AT, 218238_AT, 208724_S_AT, 208733_AT, 214435_X_AT, 205461_AT, 213603_S_AT, 201096_S_AT, 209316_S_AT, 201308_S_AT, 211622_S_AT, 217852_S_AT, 217595_AT, 208734_X_AT, 211750_X_AT, 219622_AT, 206917_AT, 207495_AT, 200927_S_AT, 208732_AT, 203105_S_AT, 207419_S_AT, 212214_AT, 219357_AT, 221960_S_AT, 204472_AT, 212213_X_AT, 212724_AT, 203267_S_AT, 207791_S_AT, 200833_S_AT, 202810_AT, 201179_S_AT | 1029 | 310 | 20063 | 1.886862 | 2.319619 |

| | | | | | | | | | | |
|---|---|---|---|---|---|---|---|---|---|---|
| INTERPRO | IPR005225:Small GTP-b | 20 | 1.876173 | 0.001824 | 206113_S_AT, 208640_AT, 218669_AT, 218238_AT, 208724_S_AT, 208733_AT, 214435_X_AT, 205461_AT, 201096_S_AT, 213603_S_AT, 211622_S_AT, 217852_S_AT, 208734_X_AT, 219622_AT, 200927_S_AT, 207495_AT, 208732_AT, 207419_S_AT, 221960_S_AT, 204472_AT, 203267_S_AT, 212724_AT, 207791_S_AT, 200833_S_AT, 202810_AT | 1008 | 167 | 18559 | 2.204995 | 3.052477 |
| UP_SEQ_FEATURE | short sequence motif:Effe | 13 | 1.219512 | 0.005484 | 208734_X_AT, 219622_AT, 206113_S_AT, 208640_AT, 207495_AT, 200927_S_AT, 218669_AT, 208732_AT, 207419_S_AT, 208724_S_AT, 221960_S_AT, 208733_AT, 212724_AT, 214435_X_AT, 205461_AT, 207791_S_AT, 200833_S_AT, 213603_S_AT | 1029 | 101 | 20063 | 2.509588 | 9.181428 |
| INTERPRO | IPR027417:P-loop conta | 64 | 6.003752 | 0.014198 | 203504_S_AT, 206544_X_AT, 203505_AT, 213829_X_AT, 213468_AT, 209549_S_AT, 203270_AT, 208724_S_AT, 204823_AT, 216466_AT, 213603_S_AT, 201308_S_AT, 217595_AT, 203196_AT, 207495_AT, 208732_AT, 203105_S_AT, 204699_S_AT, 204033_AT, 221960_S_AT, 201699_AT, 201872_S_AT, 207791_S_AT, 203515_S_AT, 200833_S_AT, 215524_X_AT, 211824_X_AT, 201459_AT, 217707_X_AT, 221737_AT, 209313_AT, 203059_S_AT, 206113_S_AT, 208640_AT, 208967_S_AT, 212257_S_AT, 218669_AT, 202348_S_AT, 218238_AT, 205429_S_AT, 213253_AT, 220060_S_AT, 208733_AT, 211212_S_AT, 201241_AT, 214435_X_AT, 205461_AT, 201096_S_AT, 209316_S_AT, 211622_S_AT, 217852_S_AT, 208734_X_AT, 204128_S_AT, 205091_X_AT, 219622_AT, 206917_AT, 208719_S_AT, 200927_S_AT, | 1008 | 879 | 18559 | 1.340557 | 21.55975 |

| | | | | | | | | | | |
|---|---|---|---|---|---|---|---|---|---|---|
| UP_SEQ_FEATURE | lipid moiety-binding regi | 12 | 1.125704 | 0.022556 | 208734_X_AT, 201921_AT, 219622_AT, 206113_S_AT, 208640_AT, 200927_S_AT, 218669_AT, 208732_AT, 207419_S_AT, 208724_S_AT, 221960_S_AT, 208733_AT, 214435_X_AT, 207791_S_AT, 205461_AT, 200833_S_AT, 213603_S_AT | 1029 | 108 | 20063 | 2.166397 | 32.93701 |
| INTERPRO | IPR001806:Small GTPas | 14 | 1.313321 | 0.039048 | 208734_X_AT, 219622_AT, 206113_S_AT, 208640_AT, 207495_AT, 200927_S_AT, 208732_AT, 218669_AT, 207419_S_AT, 208724_S_AT, 221960_S_AT, 208733_AT, 204472_AT, 212724_AT, 214435_X_AT, 207791_S_AT, 205461_AT, 200833_S_AT, 213603_S_AT | 1008 | 139 | 18559 | 1.854416 | 49.15495 |
| GOTERM_BP_DIRECT | GO:0007264~small GTP | 20 | 1.876173 | 0.128876 | 208734_X_AT, 219622_AT, 208640_AT, 206113_S_AT, 204336_S_AT, 207495_AT, 200927_S_AT, 218669_AT, 207099_S_AT, 208732_AT, 217457_S_AT, 207419_S_AT, 208724_S_AT, 215724_AT, 221960_S_AT, 208733_AT, 204472_AT, 212724_AT, 207791_S_AT, 205461_AT, 200833_S_AT, 213603_S_AT, 201096_S_AT, 211622_S_AT, 217852_S_AT | 980 | 246 | 16792 | 1.393065 | 91.98967 |
| Annotation Cluster 15 | Enrichment Score: 2.715901561841804 | | | | | | | | | |
| Category | Term | Count | % | PValue | Genes | List Total | Pop Hits | Pop Total | Fold Enrich | FDR |

| | | | | | | | | | | |
|---|---|---|---|---|---|---|---|---|---|---|
| GOTERM_MF_DIRECT | GO:0008353~RNA polyr | 9 | 0.844278 | 9.83E-07 | 203213_AT, 203198_AT, 208351_S_AT, 207319_S_AT, 204093_AT, 213468_AT, 219273_AT, 211297_S_AT, 203565_S_AT, 203214_X_AT | 971 | 16 | 16881 | 9.779158 | 0.001576 |
| UP_KEYWORDS | DNA-directed RNA polyr | 12 | 1.125704 | 2.23E-06 | 214729_AT, 205628_AT, 205053_AT, 218258_AT, 202634_AT, 208361_S_AT, 203898_AT, 213887_S_AT, 206654_S_AT, 209511_AT, 205264_AT, 217854_S_AT, 203664_S_AT | 1035 | 39 | 20581 | 6.118469 | 0.003171 |
| GOTERM_BP_DIRECT | GO:0006370~7-methylgu | 11 | 1.031895 | 1.17E-05 | 201521_S_AT, 204093_AT, 213468_AT, 209519_AT, 203565_S_AT, 211297_S_AT, 211849_S_AT, 202634_AT, 213887_S_AT, 204207_S_AT, 209511_AT, 209520_S_AT, 217854_S_AT, 203664_S_AT | 980 | 33 | 16792 | 5.711565 | 0.021417 |

| | | | | | | | | | | |
|---|---|---|---|---|---|---|---|---|---|---|
| GOTERM_BP_DIRECT | GO:0006283~transcriptio | 16 | 1.500938 | 1.96E-05 | 203103_S_AT, 204128_S_AT, 208619_AT, 200017_AT, 202467_S_AT, 202141_S_AT, 210257_X_AT, 204093_AT, 201652_AT, 213468_AT, 215997_S_AT, 203565_S_AT, 211297_S_AT, 202634_AT, 202142_AT, 213887_S_AT, 209511_AT, 202143_S_AT, 217854_S_AT, 202213_S_AT, 202214_S_AT, 203664_S_AT | 980 | 74 | 16792 | 3.704799 | 0.035802 |
| GOTERM_BP_DIRECT | GO:0006362~transcriptio | 10 | 0.938086 | 3.52E-05 | 214729_AT, 213887_S_AT, 218258_AT, 209511_AT, 205264_AT, 204093_AT, 213468_AT, 211297_S_AT, 203565_S_AT, 217854_S_AT, 202634_AT | 980 | 30 | 16792 | 5.711565 | 0.064462 |
| GOTERM_BP_DIRECT | GO:0006363~terminatior | 10 | 0.938086 | 4.71E-05 | 214729_AT, 213887_S_AT, 218258_AT, 209511_AT, 205264_AT, 204093_AT, 213468_AT, 211297_S_AT, 203565_S_AT, 217854_S_AT, 202634_AT | 980 | 31 | 16792 | 5.527321 | 0.086153 |
| GOTERM_BP_DIRECT | GO:0006361~transcriptio | 10 | 0.938086 | 8.11E-05 | 214729_AT, 213887_S_AT, 218258_AT, 209511_AT, 205264_AT, 204093_AT, 213468_AT, 211297_S_AT, 203565_S_AT, 217854_S_AT, 202634_AT | 980 | 33 | 16792 | 5.192331 | 0.148274 |

| | | | | | | | | | | |
|---|---|---|---|---|---|---|---|---|---|---|
| GOTERM_MF_DIRECT | GO:0003899~DNA-direc | 10 | 0.938086 | 2.36E-04 | 214729_AT, 203898_AT, 213887_S_AT, 206654_S_AT, 218258_AT, 209511_AT, 205264_AT, 217854_S_AT, 202634_AT, 208361_S_AT, 203664_S_AT | 971 | 38 | 16881 | 4.575045 | 0.378589 |
| KEGG_PATHWAY | hsa00240:Pyrimidine me | 19 | 1.782364 | 3.34E-04 | 214729_AT, 209773_S_AT, 205628_AT, 215165_X_AT, 205053_AT, 208828_AT, 219553_AT, 218258_AT, 203270_AT, 202634_AT, 208361_S_AT, 201577_AT, 213887_S_AT, 206654_S_AT, 202589_AT, 209511_AT, 201695_S_AT, 203939_AT, 217854_S_AT, 202706_S_AT, 203664_S_AT | 505 | 101 | 6879 | 2.562513 | 0.437531 |
| GOTERM_MF_DIRECT | GO:0001056~RNA polyr | 7 | 0.65666 | 3.62E-04 | 203898_AT, 213887_S_AT, 206654_S_AT, 218258_AT, 209511_AT, 217854_S_AT, 202634_AT, 208361_S_AT | 971 | 18 | 16881 | 6.760899 | 0.578854 |
| GOTERM_CC_DIRECT | GO:0005666~DNA-direc | 7 | 0.65666 | 4.00E-04 | 203898_AT, 213887_S_AT, 206654_S_AT, 218258_AT, 209511_AT, 217854_S_AT, 202634_AT, 208361_S_AT | 1004 | 19 | 18224 | 6.687356 | 0.598494 |

| | | | | | | | | | | |
|---|---|---|---|---|---|---|---|---|---|---|
| KEGG_PATHWAY | hsa00230:Purine metabo | 27 | 2.532833 | 4.22E-04 | 214729_AT, 205628_AT, 219553_AT, 203401_AT, 203059_S_AT, 208967_S_AT, 209549_S_AT, 217445_S_AT, 202634_AT, 210250_X_AT, 220606_S_AT, 201577_AT, 205501_AT, 206654_S_AT, 208758_AT, 203939_AT, 203741_S_AT, 203664_S_AT, 209773_S_AT, 208828_AT, 205053_AT, 218258_AT, 208361_S_AT, 213887_S_AT, 209511_AT, 203058_S_AT, 201695_S_AT, 203816_AT, 217854_S_AT, 202168_AT, 202144_S_AT | 505 | 176 | 6879 | 2.089705 | 0.552294 |
| GOTERM_CC_DIRECT | GO:0005736~DNA-direc | 6 | 0.562852 | 4.44E-04 | 214729_AT, 213887_S_AT, 218258_AT, 209511_AT, 205264_AT, 217854_S_AT, 202634_AT | 1004 | 13 | 18224 | 8.377567 | 0.664394 |
| GOTERM_MF_DIRECT | GO:0001054~RNA polym | 5 | 0.469043 | 0.003697 | 214729_AT, 213887_S_AT, 218258_AT, 209511_AT, 217854_S_AT, 202634_AT | 971 | 12 | 16881 | 7.243821 | 5.769072 |
| KEGG_PATHWAY | hsa03020:RNA polymera | 8 | 0.750469 | 0.007312 | 214729_AT, 213887_S_AT, 206654_S_AT, 218258_AT, 209511_AT, 217854_S_AT, 202634_AT, 208361_S_AT, 203664_S_AT | 505 | 32 | 6879 | 3.405446 | 9.178282 |
| GOTERM_BP_DIRECT | GO:0032481~positive re | 9 | 0.844278 | 0.00885 | 203898_AT, 213887_S_AT, 208643_S_AT, 208642_S_AT, 206654_S_AT, 218258_AT, 209511_AT, 209239_AT, 217854_S_AT, 202634_AT, 208361_S_AT | 980 | 51 | 16792 | 3.02377 | 15.01166 |

| | | | | | | | | | | |
|---|---|---|---|---|---|---|---|---|---|---|
| GOTERM_CC_DIRECT | GO:0005675~holo TFIIH | 4 | 0.375235 | 0.019665 | 204093_AT, 213468_AT, 211297_S_AT, 203565_S_AT | 1004 | 11 | 18224 | 6.600507 | 25.78712 |
| GOTERM_BP_DIRECT | GO:0050434~positive re | 6 | 0.562852 | 0.024634 | 203198_AT, 213887_S_AT, 200634_AT, 209511_AT, 217854_S_AT, 202634_AT, 203664_S_AT | 980 | 29 | 16792 | 3.545109 | 36.64198 |
| GOTERM_MF_DIRECT | GO:0008094~DNA-depe | 6 | 0.562852 | 0.034325 | 217707_X_AT, 206544_X_AT, 204093_AT, 212257_S_AT, 213468_AT, 206542_S_AT, 211297_S_AT, 203565_S_AT, 216880_AT | 971 | 32 | 16881 | 3.259719 | 42.89777 |
| GOTERM_BP_DIRECT | GO:0006383~transcriptio | 6 | 0.562852 | 0.040714 | 203898_AT, 213887_S_AT, 218258_AT, 209511_AT, 217854_S_AT, 202634_AT, 208361_S_AT | 980 | 33 | 16792 | 3.115399 | 53.25784 |
| BIOCARTA | h_ptc1Pathway:Sonic He | 4 | 0.375235 | 0.067385 | 203213_AT, 204093_AT, 211297_S_AT, 203565_S_AT, 203214_X_AT | 146 | 11 | 1625 | 4.047323 | 58.82685 |
| GOTERM_CC_DIRECT | GO:0005665~DNA-direc | 4 | 0.375235 | 0.073175 | 213887_S_AT, 209511_AT, 217854_S_AT, 202634_AT, 203664_S_AT | 1004 | 18 | 18224 | 4.033643 | 68.05265 |
| GOTERM_MF_DIRECT | GO:0001055~RNA polyn | 3 | 0.281426 | 0.109273 | 213887_S_AT, 209511_AT, 217854_S_AT, 202634_AT | 971 | 10 | 16881 | 5.215551 | 84.37595 |

| | | | | | | | | | | |
|---|---|---|---|---|---|---|---|---|---|---|
| GOTERM_BP_DIRECT | GO:0045815~positive re | 7 | 0.65666 | 0.15167 | 214729_AT, 208496_X_AT, 213887_S_AT, 218258_AT, 209511_AT, 205264_AT, 217854_S_AT, 202634_AT | 980 | 62 | 16792 | 1.934562 | 95.06872 |
| GOTERM_BP_DIRECT | GO:0035019~somatic st | 7 | 0.65666 | 0.177076 | 204270_AT, 213887_S_AT, 209511_AT, 202526_AT, 203075_AT, 217854_S_AT, 202634_AT, 203664_S_AT | 980 | 65 | 16792 | 1.845275 | 97.17285 |
| KEGG_PATHWAY | hsa04623:Cytosolic DN | 7 | 0.65666 | 0.32715 | 213887_S_AT, 206654_S_AT, 218258_AT, 209239_AT, 217854_S_AT, 202634_AT, 208361_S_AT | 505 | 64 | 6879 | 1.489882 | 99.44705 |
| KEGG_PATHWAY | hsa03022:Basal transcri | 5 | 0.469043 | 0.421611 | 221618_S_AT, 204093_AT, 213468_AT, 211297_S_AT, 203565_S_AT | 505 | 45 | 6879 | 1.513531 | 99.92399 |
| Annotation Cluster 16 | Enrichment Score: 2.663421381764644 | | | | | | | | | |
| Category | Term | Count | % | PValue | Genes | List Total | Pop Hits | Pop Total | Fold Enrich | FDR |
| UP_KEYWORDS | Nucleotide-binding | 144 | 13.50844 | 1.29E-08 | 203504_S_AT, 204283_AT, 200729_S_AT, 207721_X_AT, 203505_AT, 204641_AT, 213829_X_AT, 209549_S_AT, 201805_AT, 213468_AT, 200604_S_AT, 218535_S_AT, 200946_X_AT, 211089_S_AT, 210349_AT, 209799_AT, 203856_AT, 213603_S_AT, 201308_S_AT, 207334_S_AT, 208826_X_AT, 203213_AT, 208351_S_AT, 208642_S_AT, 203196_AT, 202695_S_AT, 205219_S_AT, 211499_S_AT, 208878_S_AT, 203198_AT, 209346_S_AT, 221960_S_AT, 201503_AT, 203515_S_AT, 217874_AT, 215195_AT, 202693_S_AT, 217356_S_AT, 215524_X_AT, 65521_AT, 200877_AT, 206686_AT, 210024_S_AT, 211824_X_AT, 217707_X_AT, 206113_S_AT, 221737_AT, 212257_S_AT, 217445_S_AT, 211080_S_AT, 211297_S_AT, 218669_AT, 211849_S_AT, 202348_S_AT, 218238_AT, 205501_AT, 211077_S_AT, 213253_AT, | 1035 | 1788 | 20581 | 1.601478 | 1.83E-05 |

| | | | | | | | | | | |
|---|---|---|---|---|---|---|---|---|---|---|
| UP_KEYWORDS | ATP-binding | 105 | 9.849906 | 2.63E-05 | 203504_S_AT, 204283_AT,<br>200729_S_AT, 203505_AT, 204641_AT,<br>213829_X_AT, 209549_S_AT,<br>213468_AT, 201805_AT, 218535_S_AT,<br>200946_X_AT, 210349_AT,<br>211089_S_AT, 209799_AT, 203856_AT,<br>207334_S_AT, 203213_AT,<br>208351_S_AT, 208642_S_AT,<br>203196_AT, 202695_S_AT,<br>211499_S_AT, 205219_S_AT,<br>208878_S_AT, 203198_AT,<br>209346_S_AT, 201503_AT,<br>203515_S_AT, 215195_AT,<br>202693_S_AT, 217356_S_AT,<br>215524_X_AT, 65521_AT, 206686_AT,<br>210024_S_AT, 211824_X_AT,<br>200877_AT, 217707_X_AT,<br>212257_S_AT, 211080_S_AT,<br>217445_S_AT, 211297_S_AT,<br>202348_S_AT, 211077_S_AT,<br>213253_AT, 202742_S_AT, 219487_AT,<br>204300_AT, 205214_AT, 209464_AT,<br>218658_S_AT, 203741_S_AT,<br>201623_S_AT, 208643_S_AT,<br>208693_S_AT, 201461_S_AT,<br>216855_S_AT, 203214_X_AT, | 1035 | 1391 | 20581 | 1.501026 | 0.037316 |
| UP_KEYWORDS | Kinase | 61 | 5.722326 | 1.50E-04 | 217503_AT, 204641_AT, 211121_S_AT,<br>209549_S_AT, 201805_AT,<br>209615_S_AT, 210984_X_AT,<br>203270_AT, 210379_S_AT,<br>208876_S_AT, 200604_S_AT,<br>218535_S_AT, 211089_S_AT,<br>210349_AT, 209799_AT, 203856_AT,<br>205393_S_AT, 207334_S_AT,<br>203626_S_AT, 214738_S_AT,<br>202246_S_AT, 203213_AT,<br>208351_S_AT, 217808_S_AT,<br>202695_S_AT, 213116_AT,<br>211499_S_AT, 205219_S_AT,<br>208878_S_AT, 205411_AT, 203198_AT,<br>207143_AT, 209622_AT, 209346_S_AT,<br>41657_AT, 209714_S_AT, 203515_S_AT,<br>202693_S_AT, 215195_AT,<br>217356_S_AT, 208229_AT, 206686_AT,<br>210626_AT, 219553_AT, 203401_AT,<br>213688_AT, 203059_S_AT,<br>207319_S_AT, 208967_S_AT,<br>211080_S_AT, 211297_S_AT,<br>206348_S_AT, 201577_AT,<br>211077_S_AT, 202742_S_AT,<br>209139_S_AT, 202126_AT, 206923_AT,<br>205214_AT, 209464_AT, 203265_S_AT, | 1035 | 735 | 20581 | 1.650322 | 0.212771 |
| GOTERM_MF_DIRECT | GO:0005524~ATP bindir | 116 | 10.8818 | 6.82E-04 | 203504_S_AT, 204283_AT,<br>200729_S_AT, 203505_AT, 204641_AT,<br>213829_X_AT, 209549_S_AT,<br>201805_AT, 213468_AT, 218535_S_AT,<br>200946_X_AT, 210349_AT,<br>211089_S_AT, 209799_AT, 203856_AT,<br>207334_S_AT, 203213_AT,<br>208351_S_AT, 208642_S_AT,<br>203196_AT, 214007_S_AT,<br>202695_S_AT, 205219_S_AT,<br>203926_X_AT, 211499_S_AT,<br>208878_S_AT, 203198_AT,<br>209346_S_AT, 201503_AT,<br>203515_S_AT, 209157_AT, 215195_AT,<br>202693_S_AT, 217356_S_AT,<br>215524_X_AT, 65521_AT, 206686_AT,<br>210024_S_AT, 211824_X_AT,<br>200877_AT, 217707_X_AT,<br>212257_S_AT, 214008_AT,<br>211080_S_AT, 217445_S_AT,<br>211297_S_AT, 202348_S_AT,<br>211077_S_AT, 213253_AT,<br>202742_S_AT, 219487_AT, 204300_AT,<br>205214_AT, 209464_AT, 218658_S_AT,<br>203741_S_AT, 215772_X_AT,<br>201623_S_AT, 208643_S_AT, | 971 | 1495 | 16881 | 1.34895 | 1.088484 |

| | | | | | | | | | | |
|---|---|---|---|---|---|---|---|---|---|---|
| SMART | SM00220:S_TKc | 33 | 3.095685 | 0.001038 | 217503_AT, 207319_S_AT, 204641_AT, 211080_S_AT, 209615_S_AT, 210379_S_AT, 211297_S_AT, 208876_S_AT, 211077_S_AT, 202742_S_AT, 206923_AT, 202126_AT, 210349_AT, 211089_S_AT, 205214_AT, 209799_AT, 209464_AT, 203856_AT, 205393_S_AT, 207334_S_AT, 202246_S_AT, 214738_S_AT, 203213_AT, 208351_S_AT, 203265_S_AT, 205394_AT, 202695_S_AT, 201461_S_AT, 213116_AT, 211499_S_AT, 203214_X_AT, 208854_S_AT, 205411_AT, 208878_S_AT, 203198_AT, 207143_AT, 209622_AT, 41657_AT, 215195_AT, 202693_S_AT, 206474_AT, 206248_AT | 504 | 359 | 10057 | 1.834245 | 1.404092 |
| UP_KEYWORDS | Serine/threonine-protein | 35 | 3.283302 | 0.001509 | 217503_AT, 207319_S_AT, 204641_AT, 211080_S_AT, 209615_S_AT, 210379_S_AT, 211297_S_AT, 208876_S_AT, 218535_S_AT, 211077_S_AT, 202742_S_AT, 206923_AT, 202126_AT, 210349_AT, 211089_S_AT, 205214_AT, 209799_AT, 209464_AT, 203856_AT, 205393_S_AT, 207334_S_AT, 202246_S_AT, 214738_S_AT, 203213_AT, 208351_S_AT, 203265_S_AT, 205394_AT, 202695_S_AT, 201461_S_AT, 213116_AT, 211499_S_AT, 203214_X_AT, 208854_S_AT, 205411_AT, 208878_S_AT, 203198_AT, 207143_AT, 209622_AT, 201234_AT, 41657_AT, 215195_AT, 202693_S_AT, 206474_AT, 206248_AT | 1035 | 393 | 20581 | 1.770931 | 2.122273 |
| GOTERM_BP_DIRECT | GO:0006468~protein ph | 43 | 4.033771 | 0.002282 | 217503_AT, 204641_AT, 209549_S_AT, 213468_AT, 201805_AT, 209615_S_AT, 203565_S_AT, 210379_S_AT, 208876_S_AT, 218535_S_AT, 210349_AT, 211089_S_AT, 209799_AT, 204220_AT, 203856_AT, 207334_S_AT, 214738_S_AT, 202246_S_AT, 208351_S_AT, 202695_S_AT, 213116_AT, 205411_AT, 208878_S_AT, 207143_AT, 203198_AT, 204883_S_AT, 41657_AT, 215195_AT, 202693_S_AT, 206686_AT, 211080_S_AT, 219273_AT, 211297_S_AT, 213756_S_AT, 211077_S_AT, 202742_S_AT, 209139_S_AT, 206923_AT, 202126_AT, 205214_AT, 209464_AT, 203075_AT, 204884_S_AT, 204093_AT, 202118_S_AT, 201461_S_AT, 208854_S_AT, 205188_S_AT, 201234_AT, 203816_AT, 206474_AT, 206248_AT | 980 | 456 | 16792 | 1.615772 | 4.094036 |

| | | | | | | | | | | |
|---|---|---|---|---|---|---|---|---|---|---|
| GOTERM_MF_DIRECT | GO:0004672~protein kin | 35 | 3.283302 | 0.00294 | 206686_AT, 217503_AT, 207319_S_AT, 204641_AT, 201805_AT, 213468_AT, 211080_S_AT, 210984_X_AT, 209615_S_AT, 211297_S_AT, 208876_S_AT, 206348_S_AT, 218535_S_AT, 206923_AT, 202126_AT, 211089_S_AT, 210349_AT, 205214_AT, 209799_AT, 209464_AT, 203856_AT, 205393_S_AT, 202246_S_AT, 214738_S_AT, 203213_AT, 203265_S_AT, 205394_AT, 202695_S_AT, 201461_S_AT, 213116_AT, 211499_S_AT, 203214_X_AT, 208854_S_AT, 205411_AT, 208878_S_AT, 203198_AT, 209622_AT, 201234_AT, 41657_AT, 215195_AT, 202693_S_AT, 206474_AT, 221957_AT, 206248_AT | 971 | 359 | 16881 | 1.694933 | 4.61422 |
| UP_SEQ_FEATURE | binding site:ATP | 45 | 4.221388 | 0.003002 | 217503_AT, 204641_AT, 210984_X_AT, 209615_S_AT, 210379_S_AT, 208876_S_AT, 218535_S_AT, 210349_AT, 211089_S_AT, 209799_AT, 203856_AT, 205393_S_AT, 207334_S_AT, 202246_S_AT, 214738_S_AT, 203213_AT, 208351_S_AT, 202695_S_AT, 213116_AT, 211499_S_AT, 205411_AT, 208878_S_AT, 203198_AT, 207143_AT, 209622_AT, 41657_AT, 203515_S_AT, 215195_AT, 202693_S_AT, 217356_S_AT, 208229_AT, 206686_AT, 219553_AT, 203401_AT, 207319_S_AT, 211080_S_AT, 211297_S_AT, 206348_S_AT, 201577_AT, 211077_S_AT, 202742_S_AT, 206923_AT, 202126_AT, 205214_AT, 209464_AT, 203265_S_AT, 205394_AT, 201461_S_AT, 208854_S_AT, 203214_X_AT, 201234_AT, 209536_S_AT, 200737_AT, 221957_AT, 206474_AT, 206248_AT | 1029 | 558 | 20063 | 1.572385 | 5.12821 |
| INTERPRO | IPR008271:Serine/threo | 30 | 2.814259 | 0.003368 | 217503_AT, 207319_S_AT, 204641_AT, 211080_S_AT, 209615_S_AT, 210379_S_AT, 211297_S_AT, 208876_S_AT, 211077_S_AT, 202742_S_AT, 206923_AT, 202126_AT, 210349_AT, 211089_S_AT, 205214_AT, 209799_AT, 209464_AT, 203856_AT, 205393_S_AT, 207334_S_AT, 214738_S_AT, 202246_S_AT, 203213_AT, 208351_S_AT, 203265_S_AT, 205394_AT, 202695_S_AT, 213116_AT, 201461_S_AT, 203214_X_AT, 208878_S_AT, 203198_AT, 207143_AT, 209622_AT, 41657_AT, 215195_AT, 202693_S_AT, 206474_AT, 206248_AT | 1008 | 312 | 18559 | 1.770356 | 5.567504 |

| | | | | | | | | | | |
|---|---|---|---|---|---|---|---|---|---|---|
| UP_SEQ_FEATURE | nucleotide phosphate-bi | 70 | 6.566604 | 0.005926 | 200840_AT, 217503_AT, 200729_S_AT, 206544_X_AT, 204641_AT, 213829_X_AT, 209549_S_AT, 213468_AT, 209615_S_AT, 210984_X_AT, 203270_AT, 210379_S_AT, 208876_S_AT, 211089_S_AT, 210349_AT, 200079_S_AT, 209799_AT, 203856_AT, 205393_S_AT, 207334_S_AT, 214738_S_AT, 202246_S_AT, 203213_AT, 208351_S_AT, 200727_S_AT, 202695_S_AT, 213116_AT, 211499_S_AT, 205219_S_AT, 208878_S_AT, 205411_AT, 203198_AT, 207143_AT, 209622_AT, 204033_AT, 41657_AT, 201699_AT, 203515_S_AT, 202693_S_AT, 215195_AT, 217356_S_AT, 215524_X_AT, 208229_AT, 211824_X_AT, 201459_AT, 206686_AT, 217707_X_AT, 203401_AT, 203059_S_AT, 207319_S_AT, 208967_S_AT, 212257_S_AT, 211080_S_AT, 217445_S_AT, 211297_S_AT, 202348_S_AT, 206348_S_AT, 211077_S_AT, | 1029 | 994 | 20063 | 1.373068 | 9.885012 |
| GOTERM_MF_DIRECT | GO:0004674~protein ser | 34 | 3.189493 | 0.010334 | 217503_AT, 207319_S_AT, 204641_AT, 211080_S_AT, 209615_S_AT, 210379_S_AT, 219273_AT, 211297_S_AT, 208876_S_AT, 206348_S_AT, 218535_S_AT, 211077_S_AT, 202742_S_AT, 206923_AT, 202126_AT, 211089_S_AT, 205214_AT, 209799_AT, 209464_AT, 203856_AT, 205393_S_AT, 202246_S_AT, 214738_S_AT, 203213_AT, 208351_S_AT, 205394_AT, 202118_S_AT, 202695_S_AT, 201461_S_AT, 213116_AT, 211499_S_AT, 203214_X_AT, 208854_S_AT, 205411_AT, 208878_S_AT, 203198_AT, 209622_AT, 201234_AT, 41657_AT, 215195_AT, 202693_S_AT, 206474_AT, 221957_AT, 206248_AT | 971 | 376 | 16881 | 1.572063 | 15.35031 |
| UP_SEQ_FEATURE | active site:Proton accept | 49 | 4.596623 | 0.012528 | 217503_AT, 204641_AT, 210984_X_AT, 209615_S_AT, 210379_S_AT, 208876_S_AT, 218535_S_AT, 211089_S_AT, 210349_AT, 208758_AT, 209799_AT, 203856_AT, 205393_S_AT, 207334_S_AT, 202246_S_AT, 214738_S_AT, 202862_AT, 203213_AT, 208351_S_AT, 208369_S_AT, 202695_S_AT, 213116_AT, 211499_S_AT, 205411_AT, 208878_S_AT, 207143_AT, 203198_AT, 209622_AT, 41657_AT, 200966_X_AT, 202693_S_AT, 215195_AT, 202144_S_AT, 208229_AT, 210153_S_AT, 207319_S_AT, 214687_X_AT, 211080_S_AT, 211297_S_AT, 210250_X_AT, 211077_S_AT, 201007_AT, 202742_S_AT, 209448_AT, 210154_AT, 202126_AT, 206923_AT, 205214_AT, 209464_AT, 200978_AT, 203265_S_AT, 205394_AT, 207180_S_AT, 201461_S_AT, 203328_X_AT, 203214_X_AT, 208854_S_AT, 209512_AT, 210027_S_AT, 202266_AT, 206474_AT, 206248_AT | 1029 | 672 | 20063 | 1.421698 | 19.81135 |

| | | | | | | | | | | |
|---|---|---|---|---|---|---|---|---|---|---|
| UP_SEQ_FEATURE | domain:Protein kinase | 37 | 3.470919 | 0.013388 | 208229_AT, 217503_AT, 207319_S_AT, 204641_AT, 211080_S_AT, 210984_X_AT, 209615_S_AT, 210379_S_AT, 211297_S_AT, 208876_S_AT, 218535_S_AT, 211077_S_AT, 202742_S_AT, 206923_AT, 202126_AT, 211089_S_AT, 210349_AT, 205214_AT, 209799_AT, 209464_AT, 203856_AT, 205393_S_AT, 207334_S_AT, 202246_S_AT, 214738_S_AT, 203213_AT, 208351_S_AT, 203265_S_AT, 205394_AT, 202695_S_AT, 201461_S_AT, 213116_AT, 211499_S_AT, 203214_X_AT, 208854_S_AT, 205411_AT, 208878_S_AT, 203198_AT, 207143_AT, 209622_AT, 201234_AT, 41657_AT, 215195_AT, 202693_S_AT, 206474_AT, 206248_AT | 1029 | 478 | 20063 | 1.509226 | 21.02512 |
| INTERPRO | IPR017441:Protein kinas | 31 | 2.908068 | 0.026648 | 208229_AT, 217503_AT, 207319_S_AT, 210984_X_AT, 209615_S_AT, 210379_S_AT, 211297_S_AT, 208876_S_AT, 211077_S_AT, 202742_S_AT, 206923_AT, 210349_AT, 211089_S_AT, 205214_AT, 209799_AT, 209464_AT, 203856_AT, 205393_S_AT, 207334_S_AT, 202246_S_AT, 203213_AT, 208351_S_AT, 203265_S_AT, 205394_AT, 202695_S_AT, 213116_AT, 201461_S_AT, 211499_S_AT, 203214_X_AT, 205411_AT, 208878_S_AT, 208854_S_AT, 203198_AT, 207143_AT, 41657_AT, 215195_AT, 202693_S_AT, 206474_AT, 206248_AT | 1008 | 381 | 18559 | 1.498065 | 36.78786 |
| INTERPRO | IPR000719:Protein kinas | 36 | 3.377111 | 0.055406 | 208229_AT, 217503_AT, 207319_S_AT, 204641_AT, 210984_X_AT, 211080_S_AT, 209615_S_AT, 210379_S_AT, 211297_S_AT, 208876_S_AT, 211077_S_AT, 202742_S_AT, 206923_AT, 202126_AT, 210349_AT, 211089_S_AT, 205214_AT, 209799_AT, 209464_AT, 203856_AT, 205393_S_AT, 207334_S_AT, 202246_S_AT, 214738_S_AT, 203213_AT, 208351_S_AT, 203265_S_AT, 205394_AT, 202695_S_AT, 201461_S_AT, 213116_AT, 211499_S_AT, 203214_X_AT, 208854_S_AT, 205411_AT, 208878_S_AT, 203198_AT, 207143_AT, 209622_AT, 201234_AT, 41657_AT, 215195_AT, 202693_S_AT, 206474_AT, 206248_AT | 1008 | 487 | 18559 | 1.36103 | 62.0139 |

| | | | | | | | | | | |
|---|---|---|---|---|---|---|---|---|---|---|
| INTERPRO | IPR011009:Protein kinas | 37 | 3.470919 | 0.100006 | 208229_AT, 217503_AT, 207319_S_AT, 204641_AT, 211080_S_AT, 210984_X_AT, 209615_S_AT, 210379_S_AT, 211297_S_AT, 208876_S_AT, 218535_S_AT, 211077_S_AT, 202742_S_AT, 206923_AT, 202126_AT, 211089_S_AT, 210349_AT, 205214_AT, 209799_AT, 209464_AT, 203856_AT, 205393_S_AT, 207334_S_AT, 202246_S_AT, 214738_S_AT, 203213_AT, 208351_S_AT, 203265_S_AT, 205394_AT, 202695_S_AT, 201461_S_AT, 213116_AT, 211499_S_AT, 203214_X_AT, 208854_S_AT, 205411_AT, 208878_S_AT, 203198_AT, 207143_AT, 209622_AT, 201234_AT, 41657_AT, 215195_AT, 202693_S_AT, 206474_AT, 206248_AT | 1008 | 531 | 18559 | 1.282925 | 83.29252 |
| GOTERM_BP_DIRECT | GO:0046777~protein au | 14 | 1.313321 | 0.204345 | 208229_AT, 217503_AT, 204641_AT, 211080_S_AT, 210984_X_AT, 209615_S_AT, 201461_S_AT, 208876_S_AT, 208878_S_AT, 208854_S_AT, 205411_AT, 209622_AT, 41657_AT, 210349_AT, 205214_AT, 209464_AT, 203856_AT | 980 | 172 | 16792 | 1.394684 | 98.47392 |
| Annotation Cluster 17 | Enrichment Score: 2.1232808514361574 | | | | | | | | | |
| Category | Term | Count | % | PValue | Genes | List Total | Pop Hits | Pop Total | Fold Enrich | FDR |
| UP_KEYWORDS | Iron-sulfur | 11 | 1.031895 | 7.49E-04 | 207071_S_AT, 211752_S_AT, 205628_AT, 205083_AT, 200793_S_AT, 208909_AT, 218597_S_AT, 213829_X_AT, 214205_X_AT, 213468_AT, 209080_X_AT, 221094_S_AT | 1035 | 60 | 20581 | 3.645588 | 1.057926 |
| UP_SEQ_FEATURE | metal ion-binding site:Iro | 6 | 0.562852 | 0.001291 | 207071_S_AT, 211752_S_AT, 205628_AT, 200793_S_AT, 213829_X_AT, 213468_AT | 1029 | 17 | 20063 | 6.881495 | 2.236228 |

| | | | | | | | | | | |
|---|---|---|---|---|---|---|---|---|---|---|
| UP_KEYWORDS | 4Fe-4S | 6 | 0.562852 | 0.036425 | 207071_S_AT, 211752_S_AT, 205628_AT, 200793_S_AT, 213829_X_AT, 213468_AT | 1035 | 37 | 20581 | 3.224599 | 40.95677 |
| GOTERM_MF_DIRECT | GO:0051539~4 iron, 4 su | 6 | 0.562852 | 0.091285 | 207071_S_AT, 211752_S_AT, 205628_AT, 200793_S_AT, 213829_X_AT, 213468_AT | 971 | 42 | 16881 | 2.483596 | 78.46755 |
| Annotation Cluster 18 | Enrichment Score: 2.1188921350642107 | | | | | | | | | |
| Category | Term | Count | % | PValue | Genes | List Total | Pop Hits | Pop Total | Fold Enrich | FDR |
| KEGG_PATHWAY | hsa05212:Pancreatic ca | 15 | 1.407129 | 2.01E-04 | 208351_S_AT, 208640_AT, 203132_AT, 210984_X_AT, 205397_X_AT, 207419_S_AT, 207143_AT, 205015_S_AT, 215037_S_AT, 214435_X_AT, 213603_S_AT, 202526_AT, 207334_S_AT, 203075_AT, 209239_AT, 205398_S_AT, 202246_S_AT | 505 | 65 | 6879 | 3.143488 | 0.263108 |
| BIOCARTA | h_raccycdPathway:Influe | 7 | 0.65666 | 0.027879 | 207143_AT, 208351_S_AT, 208640_AT, 203132_AT, 209615_S_AT, 209239_AT, 202246_S_AT | 146 | 27 | 1625 | 2.885591 | 30.20872 |
| KEGG_PATHWAY | hsa05220:Chronic myelo | 10 | 0.938086 | 0.078567 | 207143_AT, 208351_S_AT, 215037_S_AT, 201218_AT, 203132_AT, 202226_S_AT, 202526_AT, 207334_S_AT, 209239_AT, 202246_S_AT | 505 | 72 | 6879 | 1.891914 | 65.81409 |
| Annotation Cluster 19 | Enrichment Score: 2.1075005385402457 | | | | | | | | | |
| Category | Term | Count | % | PValue | Genes | List Total | Pop Hits | Pop Total | Fold Enrich | FDR |

| | | | | | | | | | | |
|---|---|---|---|---|---|---|---|---|---|---|
| SMART | SM00320:WD40 | 26 | 2.439024 | 0.001836 | 203103_S_AT, 202190_AT, 203721_S_AT, 221531_AT, 220762_S_AT, 203163_AT, 220258_S_AT, 200813_S_AT, 209974_S_AT, 211684_S_AT, 211318_S_AT, 65591_AT, 218564_AT, 214336_S_AT, 219193_AT, 201558_AT, 201457_X_AT, 218512_AT, 32723_AT, 210935_S_AT, 202031_S_AT, 211547_S_AT, 218278_AT, 221207_S_AT, 209592_S_AT, 221938_X_AT, 203409_AT, 219109_AT, 200744_S_AT, 218585_S_AT | 504 | 267 | 10057 | 1.943122 | 2.471915 |
| UP_SEQ_FEATURE | repeat:WD 3 | 26 | 2.439024 | 0.003022 | 203103_S_AT, 202190_AT, 203721_S_AT, 221531_AT, 220762_S_AT, 203163_AT, 220258_S_AT, 200813_S_AT, 209974_S_AT, 211684_S_AT, 211318_S_AT, 65591_AT, 218564_AT, 214336_S_AT, 219193_AT, 201558_AT, 201457_X_AT, 218512_AT, 32723_AT, 210935_S_AT, 202031_S_AT, 211547_S_AT, 218278_AT, 221207_S_AT, 209592_S_AT, 221938_X_AT, 203409_AT, 219109_AT, 200744_S_AT, 218585_S_AT | 1029 | 269 | 20063 | 1.884524 | 5.162261 |
| UP_KEYWORDS | WD repeat | 26 | 2.439024 | 0.004172 | 203103_S_AT, 202190_AT, 203721_S_AT, 221531_AT, 220762_S_AT, 203163_AT, 220258_S_AT, 200813_S_AT, 209974_S_AT, 211684_S_AT, 211318_S_AT, 65591_AT, 218564_AT, 214336_S_AT, 219193_AT, 201558_AT, 201457_X_AT, 218512_AT, 32723_AT, 210935_S_AT, 202031_S_AT, 211547_S_AT, 218278_AT, 221207_S_AT, 209592_S_AT, 221938_X_AT, 203409_AT, 219109_AT, 200744_S_AT, 218585_S_AT | 1035 | 281 | 20581 | 1.839895 | 5.76358 |

| | | | | | | | | | | |
|---|---|---|---|---|---|---|---|---|---|---|
| INTERPRO | IPR020472:G-protein be | 13 | 1.219512 | 0.0044 | 203103_S_AT, 202190_AT, 214336_S_AT, 221531_AT, 201558_AT, 201457_X_AT, 203163_AT, 218512_AT, 32723_AT, 210935_S_AT, 200813_S_AT, 211547_S_AT, 209974_S_AT, 211318_S_AT, 65591_AT, 219109_AT, 200744_S_AT | 1008 | 93 | 18559 | 2.573679 | 7.215016 |
| UP_SEQ_FEATURE | repeat:WD 1 | 26 | 2.439024 | 0.004626 | 203103_S_AT, 202190_AT, 203721_S_AT, 221531_AT, 220762_S_AT, 203163_AT, 220258_S_AT, 200813_S_AT, 209974_S_AT, 211684_S_AT, 211318_S_AT, 65591_AT, 218564_AT, 214336_S_AT, 219193_AT, 201558_AT, 201457_X_AT, 218512_AT, 32723_AT, 210935_S_AT, 202031_S_AT, 211547_S_AT, 218278_AT, 221207_S_AT, 209592_S_AT, 221938_X_AT, 203409_AT, 219109_AT, 200744_S_AT, 218585_S_AT | 1029 | 278 | 20063 | 1.823514 | 7.799756 |
| UP_SEQ_FEATURE | repeat:WD 2 | 26 | 2.439024 | 0.004626 | 203103_S_AT, 202190_AT, 203721_S_AT, 221531_AT, 220762_S_AT, 203163_AT, 220258_S_AT, 200813_S_AT, 209974_S_AT, 211684_S_AT, 211318_S_AT, 65591_AT, 218564_AT, 214336_S_AT, 219193_AT, 201558_AT, 201457_X_AT, 218512_AT, 32723_AT, 210935_S_AT, 202031_S_AT, 211547_S_AT, 218278_AT, 221207_S_AT, 209592_S_AT, 221938_X_AT, 203409_AT, 219109_AT, 200744_S_AT, 218585_S_AT | 1029 | 278 | 20063 | 1.823514 | 7.799756 |

| | | | | | | | | | | |
|---|---|---|---|---|---|---|---|---|---|---|
| INTERPRO | IPR001680:WD40 repea | 26 | 2.439024 | 0.005973 | 203103_S_AT, 202190_AT, 203721_S_AT, 221531_AT, 220762_S_AT, 203163_AT, 220258_S_AT, 200813_S_AT, 209974_S_AT, 211684_S_AT, 211318_S_AT, 65591_AT, 218564_AT, 214336_S_AT, 219193_AT, 201558_AT, 201457_X_AT, 218512_AT, 32723_AT, 210935_S_AT, 202031_S_AT, 211547_S_AT, 218278_AT, 221207_S_AT, 209592_S_AT, 221938_X_AT, 203409_AT, 219109_AT, 200744_S_AT, 218585_S_AT | 1008 | 268 | 18559 | 1.78621 | 9.673786 |
| UP_SEQ_FEATURE | repeat:WD 4 | 24 | 2.251407 | 0.006199 | 203103_S_AT, 202190_AT, 203721_S_AT, 221531_AT, 220762_S_AT, 203163_AT, 220258_S_AT, 200813_S_AT, 209974_S_AT, 211684_S_AT, 211318_S_AT, 65591_AT, 219193_AT, 214336_S_AT, 201558_AT, 201457_X_AT, 218512_AT, 32723_AT, 210935_S_AT, 211547_S_AT, 221207_S_AT, 218278_AT, 209592_S_AT, 203409_AT, 221938_X_AT, 219109_AT, 200744_S_AT, 218585_S_AT | 1029 | 255 | 20063 | 1.835065 | 10.31762 |
| INTERPRO | IPR015943:WD40/YVTN | 30 | 2.814259 | 0.007726 | 203103_S_AT, 202190_AT, 203721_S_AT, 221531_AT, 203163_AT, 213030_S_AT, 220762_S_AT, 220258_S_AT, 200813_S_AT, 215324_AT, 211684_S_AT, 209974_S_AT, 211318_S_AT, 65591_AT, 207290_AT, 218564_AT, 214336_S_AT, 219193_AT, 208619_AT, 201558_AT, 201457_X_AT, 219688_AT, 218512_AT, 32723_AT, 210935_S_AT, 202031_S_AT, 211547_S_AT, 218278_AT, 221207_S_AT, 209592_S_AT, 203409_AT, 221938_X_AT, 219109_AT, 200744_S_AT, 218585_S_AT | 1008 | 331 | 18559 | 1.668735 | 12.34024 |

| | | | | | | | | | | |
|---|---|---|---|---|---|---|---|---|---|---|
| INTERPRO | IPR019775:WD40 repea | 18 | 1.688555 | 0.008864 | 203103_S_AT, 202190_AT, 214336_S_AT, 203721_S_AT, 219193_AT, 201558_AT, 221531_AT, 220762_S_AT, 203163_AT, 218512_AT, 32723_AT, 210935_S_AT, 200813_S_AT, 211547_S_AT, 211318_S_AT, 65591_AT, 209592_S_AT, 203409_AT, 200744_S_AT, 219109_AT, 218585_S_AT | 1008 | 166 | 18559 | 1.99645 | 14.0314 |
| INTERPRO | IPR017986:WD40-repea | 28 | 2.626642 | 0.009071 | 203103_S_AT, 202190_AT, 203721_S_AT, 221531_AT, 220762_S_AT, 203163_AT, 220258_S_AT, 200813_S_AT, 209974_S_AT, 211684_S_AT, 211318_S_AT, 65591_AT, 218564_AT, 214336_S_AT, 219193_AT, 208619_AT, 201558_AT, 201457_X_AT, 219688_AT, 218512_AT, 32723_AT, 210935_S_AT, 202031_S_AT, 211547_S_AT, 218278_AT, 221207_S_AT, 209592_S_AT, 221938_X_AT, 203409_AT, 219109_AT, 200744_S_AT, 218585_S_AT | 1008 | 306 | 18559 | 1.684731 | 14.33613 |
| UP_SEQ_FEATURE | repeat:WD 5 | 22 | 2.06379 | 0.010532 | 203103_S_AT, 202190_AT, 203721_S_AT, 221531_AT, 203163_AT, 220762_S_AT, 220258_S_AT, 200813_S_AT, 211684_S_AT, 209974_S_AT, 65591_AT, 219193_AT, 214336_S_AT, 201457_X_AT, 218512_AT, 32723_AT, 210935_S_AT, 211547_S_AT, 221207_S_AT, 218278_AT, 203409_AT, 221938_X_AT, 219109_AT, 200744_S_AT, 218585_S_AT | 1029 | 237 | 20063 | 1.809901 | 16.92394 |

| | | | | | | | | | | |
|---|---|---|---|---|---|---|---|---|---|---|
| UP_SEQ_FEATURE | repeat:WD 6 | 18 | 1.688555 | 0.023997 | 203103_S_AT, 202190_AT, 214336_S_AT, 203721_S_AT, 219193_AT, 221531_AT, 220762_S_AT, 203163_AT, 218512_AT, 220258_S_AT, 32723_AT, 210935_S_AT, 200813_S_AT, 211547_S_AT, 211684_S_AT, 65591_AT, 218278_AT, 200744_S_AT, 219109_AT, 218585_S_AT | 1029 | 196 | 20063 | 1.790593 | 34.64694 |
| UP_SEQ_FEATURE | repeat:WD 7 | 11 | 1.031895 | 0.246681 | 203103_S_AT, 219193_AT, 211684_S_AT, 221531_AT, 65591_AT, 218512_AT, 210935_S_AT, 219109_AT, 200744_S_AT, 200813_S_AT, 218585_S_AT, 211547_S_AT | 1029 | 151 | 20063 | 1.420353 | 99.29922 |
| Annotation Cluster 20 | Enrichment Score: 2.103308326132548 | | | | | | | | | |
| Category | Term | Count | % | PValue | Genes | List Total | Pop Hits | Pop Total | Fold Enrich | FDR |
| GOTERM_BP_DIRECT | GO:1901998~toxin trans | 9 | 0.844278 | 9.09E-04 | 200873_S_AT, 217726_AT, 200877_AT, 219861_AT, 200881_S_AT, 200812_AT, 220953_S_AT, 207495_AT, 200880_AT, 208696_AT | 980 | 36 | 16792 | 4.283673 | 1.650475 |
| GOTERM_BP_DIRECT | GO:1904851~positive re | 5 | 0.469043 | 0.001142 | 200873_S_AT, 200877_AT, 200812_AT, 220258_S_AT, 208696_AT | 980 | 9 | 16792 | 9.519274 | 2.069129 |

| | | | | | | | | | | |
|---|---|---|---|---|---|---|---|---|---|---|
| GOTERM_BP_DIRECT | GO:1904874~positive re | 6 | 0.562852 | 0.001225 | 200873_S_AT, 200877_AT, 201459_AT, 200812_AT, 208696_AT, 201614_S_AT | 980 | 15 | 16792 | 6.853878 | 2.216941 |
| GOTERM_BP_DIRECT | GO:0032212~positive re | 8 | 0.750469 | 0.002087 | 200873_S_AT, 208351_S_AT, 200877_AT, 200812_AT, 204641_AT, 220258_S_AT, 209464_AT, 211080_S_AT, 208696_AT | 980 | 32 | 16792 | 4.283673 | 3.749873 |
| INTERPRO | IPR027413:GroEL-like e | 5 | 0.469043 | 0.007268 | 200873_S_AT, 200877_AT, 219487_AT, 200812_AT, 208696_AT | 1008 | 15 | 18559 | 6.137235 | 11.65141 |
| GOTERM_BP_DIRECT | GO:1904871~positive re | 4 | 0.375235 | 0.008867 | 200873_S_AT, 200877_AT, 200812_AT, 208696_AT | 980 | 8 | 16792 | 8.567347 | 15.03741 |
| INTERPRO | IPR002423:Chaperonin | 5 | 0.469043 | 0.009281 | 200873_S_AT, 200877_AT, 219487_AT, 200812_AT, 208696_AT | 1008 | 16 | 18559 | 5.753658 | 14.64427 |
| INTERPRO | IPR027409:GroEL-like a | 5 | 0.469043 | 0.009281 | 200873_S_AT, 200877_AT, 219487_AT, 200812_AT, 208696_AT | 1008 | 16 | 18559 | 5.753658 | 14.64427 |
| INTERPRO | IPR002194:Chaperonin | 4 | 0.375235 | 0.010455 | 200873_S_AT, 200877_AT, 200812_AT, 208696_AT | 1008 | 9 | 18559 | 8.182981 | 16.34515 |
| GOTERM_CC_DIRECT | GO:0002199~zona pellu | 4 | 0.375235 | 0.010872 | 200873_S_AT, 200877_AT, 200812_AT, 208696_AT | 1004 | 9 | 18224 | 8.067286 | 15.13798 |

| | | | | | | | | | | |
|---|---|---|---|---|---|---|---|---|---|---|
| GOTERM_CC_DIRECT | GO:0005832~chaperoni | 4 | 0.375235 | 0.010872 | 200873_S_AT, 200877_AT, 200812_AT, 208696_AT | 1004 | 9 | 18224 | 8.067286 | 15.13798 |
| INTERPRO | IPR017998:Chaperone t | 4 | 0.375235 | 0.024243 | 200873_S_AT, 200877_AT, 200812_AT, 208696_AT | 1008 | 12 | 18559 | 6.137235 | 34.08151 |
| INTERPRO | IPR027410:TCP-1-like c | 4 | 0.375235 | 0.024243 | 200873_S_AT, 200877_AT, 200812_AT, 208696_AT | 1008 | 12 | 18559 | 6.137235 | 34.08151 |
| GOTERM_BP_DIRECT | GO:0007339~binding of | 4 | 0.375235 | 0.334549 | 200873_S_AT, 200877_AT, 200812_AT, 208696_AT | 980 | 35 | 16792 | 1.958251 | 99.94198 |
| Annotation Cluster 21 | Enrichment Score: 2.085631829512401 | | | | | | | | | |
| Category | Term | Count | % | PValue | Genes | List Total | Pop Hits | Pop Total | Fold Enrich | FDR |
| GOTERM_BP_DIRECT | GO:0006294~nucleotide | 10 | 0.938086 | 2.60E-05 | 200017_AT, 208619_AT, 210257_X_AT, 204093_AT, 213468_AT, 215997_S_AT, 203565_S_AT, 211297_S_AT, 201222_S_AT, 203409_AT, 202214_S_AT, 202213_S_AT, 209375_AT | 980 | 29 | 16792 | 5.908515 | 0.047575 |
| GOTERM_BP_DIRECT | GO:0000715~nucleotide | 9 | 0.844278 | 2.90E-05 | 200017_AT, 208619_AT, 202467_S_AT, 210257_X_AT, 202141_S_AT, 201652_AT, 201222_S_AT, 215997_S_AT, 202142_AT, 202143_S_AT, 203409_AT, 202214_S_AT, 202213_S_AT, 209375_AT | 980 | 23 | 16792 | 6.70488 | 0.052984 |

| | | | | | | | | | | |
|---|---|---|---|---|---|---|---|---|---|---|
| GOTERM_BP_DIRECT | GO:0070911~global gen | 9 | 0.844278 | 3.89E-04 | 200017_AT, 208619_AT, 210257_X_AT, 200739_S_AT, 213468_AT, 201222_S_AT, 215997_S_AT, 203409_AT, 209096_AT, 202213_S_AT, 202214_S_AT, 209375_AT | 980 | 32 | 16792 | 4.819133 | 0.709015 |
| GOTERM_BP_DIRECT | GO:0000717~nucleotide | 7 | 0.65666 | 0.001287 | 200017_AT, 208619_AT, 210257_X_AT, 213468_AT, 201222_S_AT, 215997_S_AT, 203409_AT, 202213_S_AT, 202214_S_AT, 209375_AT | 980 | 22 | 16792 | 5.451948 | 2.328201 |
| GOTERM_CC_DIRECT | GO:0031465~Cul4B-RIN | 4 | 0.375235 | 0.001529 | 208619_AT, 210257_X_AT, 215997_S_AT, 203409_AT, 218585_S_AT, 202213_S_AT, 202214_S_AT | 1004 | 5 | 18224 | 14.52112 | 2.270905 |
| KEGG_PATHWAY | hsa03420:Nucleotide ex | 11 | 1.031895 | 0.001783 | 204128_S_AT, 208619_AT, 208828_AT, 210257_X_AT, 204093_AT, 213468_AT, 201222_S_AT, 215997_S_AT, 203565_S_AT, 211297_S_AT, 203409_AT, 202214_S_AT, 202213_S_AT, 209375_AT | 505 | 47 | 6879 | 3.188077 | 2.314204 |

| | | | | | | | | | | |
|---|---|---|---|---|---|---|---|---|---|---|
| GOTERM_BP_DIRECT | GO:0042769~DNA dama | 7 | 0.65666 | 0.016776 | 208692_AT, 204128_S_AT, 200017_AT, 208619_AT, 200881_S_AT, 210257_X_AT, 200880_AT, 215997_S_AT, 218585_S_AT, 202213_S_AT, 202214_S_AT | 980 | 36 | 16792 | 3.331746 | 26.6217 |
| GOTERM_BP_DIRECT | GO:0070914~UV-damag | 4 | 0.375235 | 0.022913 | 208619_AT, 210257_X_AT, 215997_S_AT, 203409_AT, 202213_S_AT, 202214_S_AT, 209375_AT | 980 | 11 | 16792 | 6.230798 | 34.56479 |
| GOTERM_BP_DIRECT | GO:0006289~nucleotide | 7 | 0.65666 | 0.030322 | 204883_S_AT, 208619_AT, 204884_S_AT, 219502_AT, 213468_AT, 201222_S_AT, 203409_AT, 209375_AT | 980 | 41 | 16792 | 2.925436 | 43.07231 |
| GOTERM_BP_DIRECT | GO:1901990~regulation | 3 | 0.281426 | 0.043534 | 208619_AT, 213468_AT, 209375_AT | 980 | 6 | 16792 | 8.567347 | 55.70928 |
| GOTERM_BP_DIRECT | GO:0006296~nucleotide | 6 | 0.562852 | 0.061931 | 204128_S_AT, 200017_AT, 208619_AT, 210257_X_AT, 213468_AT, 215997_S_AT, 203409_AT, 202213_S_AT, 202214_S_AT | 980 | 37 | 16792 | 2.778599 | 68.95548 |

| | | | | | | | | | | |
|---|---|---|---|---|---|---|---|---|---|---|
| GOTERM_BP_DIRECT | GO:0033683~nucleotide | 6 | 0.562852 | 0.068049 | 204128_S_AT, 200017_AT, 208619_AT, 210257_X_AT, 213468_AT, 215997_S_AT, 203409_AT, 202213_S_AT, 202214_S_AT | 980 | 38 | 16792 | 2.705478 | 72.45802 |
| UP_KEYWORDS | Xeroderma pigmentosun | 3 | 0.281426 | 0.07178 | 213468_AT, 203409_AT, 209375_AT | 1035 | 9 | 20581 | 6.628341 | 65.27623 |
| GOTERM_BP_DIRECT | GO:0006293~nucleotide | 4 | 0.375235 | 0.120421 | 208619_AT, 210257_X_AT, 213468_AT, 215997_S_AT, 203409_AT, 202213_S_AT, 202214_S_AT | 980 | 21 | 16792 | 3.263751 | 90.44101 |
| GOTERM_BP_DIRECT | GO:0006295~nucleotide | 4 | 0.375235 | 0.147479 | 208619_AT, 210257_X_AT, 213468_AT, 215997_S_AT, 203409_AT, 202213_S_AT, 202214_S_AT | 980 | 23 | 16792 | 2.979947 | 94.60337 |
| GOTERM_BP_DIRECT | GO:0035518~histone H2 | 3 | 0.281426 | 0.152293 | 208619_AT, 210257_X_AT, 215997_S_AT, 203409_AT, 202213_S_AT, 202214_S_AT | 980 | 12 | 16792 | 4.283673 | 95.13447 |
