## Supplementary material for "Targeting dormant ovarian cancer cells *in vitro* and in an *in vivo* model of platinum resistance": Table S5

| Table S5. Functional annotations for genes with higher expression in cells with low spheroid-forming capacity | | | | | | | | | | |
|---|---|---|---|---|---|---|---|---|---|---|
| Annotation Cluster 1 | Enrichment Score: 5.637675676584239 | | | | | | | | | |
| Category | Term | Count | % | PValue | Genes | List Total | Pop Hits | Pop Total | ld Enrichme | FDR |
| UP_KEYWORDS | Metal-binding | 268 | 24.05745 | 1.81E-10 | 206836_AT, 215552_S_AT, 204161_S_AT, 206078_AT, 221092_AT, 202197_AT, 221789_X_AT, 222152_AT, 221287_AT, 219951_S_AT, 206605_AT, 210391_AT, 210944_S_AT, 201549_X_AT, 206683_AT, 205624_AT, 206166_S_AT, 205998_X_AT, 209211_AT, 210336_X_AT, 202702_AT, 210800_AT, 207129_AT, 217922_AT, 209976_S_AT, 202326_AT, 207058_S_AT, 210694_S_AT, 220055_AT, 207625_S_AT, 207186_S_AT, 211002_S_AT, 216026_S_AT, 212816_S_AT, 219102_AT, 218173_S_AT, 206301_AT, 206502_S_AT, 217428_S_AT, 41113_AT, 215461_AT, 204685_S_AT, 203160_S_AT, 203948_S_AT, 203704_S_AT, 212359_S_AT, 217594_AT, 217242_AT, 203765_AT, 206718_AT, 202871_AT, 202340_X_AT, 201975_AT, 201285_AT, 216780_AT, 205178_S_AT, 212369_AT, 222104_X_AT, 215911_X_AT, 210579_S_AT, 31835_AT, 211202_S_AT, 213832_AT, 203739_AT, | 1060 | 3640 | 20581 | 1.429532 | 2.59E-07 |
| UP_KEYWORDS | Zinc-finger | 140 | 12.56732 | 3.66E-07 | 215552_S_AT, 202197_AT, 221092_AT, 205181_AT, 212080_AT, 221287_AT, 219951_S_AT, 210391_AT, 201549_X_AT, 206683_AT, 219086_AT, 209211_AT, 210336_X_AT, 213204_AT, 202702_AT, 207058_S_AT, 220618_S_AT, 210694_S_AT, 221645_S_AT, 220113_X_AT, 220055_AT, 214861_AT, 219676_AT, 207625_S_AT, 207186_S_AT, 211002_S_AT, 204523_AT, 216026_S_AT, 209959_AT, 220668_S_AT, 218173_S_AT, 218524_AT, 206301_AT, 206502_S_AT, 203956_AT, 218803_AT, 41113_AT, 215461_AT, 213639_S_AT, 207120_AT, 203160_S_AT, 201368_AT, 203704_S_AT, 212359_S_AT, 217594_AT, 217242_AT, 212485_AT, 220891_AT, 202340_X_AT, 202871_AT, 214751_AT, 201975_AT, 212753_AT, 210266_S_AT, 201285_AT, 216780_AT, 212369_AT, 222104_X_AT, 205178_S_AT, 210579_S_AT, 211202_S_AT, 216350_S_AT, 214253_S_AT, 203739_AT, 213081_AT, | 1060 | 1781 | 20581 | 1.526247 | 5.22E-04 |
| UP_KEYWORDS | Zinc | 172 | 15.43986 | 1.31E-06 | 215552_S_AT, 204161_S_AT, 221092_AT, 202197_AT, 221287_AT, 219951_S_AT, 210391_AT, 201549_X_AT, 206683_AT, 205624_AT, 206166_S_AT, 209211_AT, 210336_X_AT, 210800_AT, 202702_AT, 207129_AT, 202326_AT, 207058_S_AT, 210694_S_AT, 220055_AT, 211002_S_AT, 207186_S_AT, 207625_S_AT, 216026_S_AT, 218173_S_AT, 206301_AT, 206502_S_AT, 41113_AT, 215461_AT, 203160_S_AT, 203704_S_AT, 212359_S_AT, 217594_AT, 217242_AT, 206718_AT, 202340_X_AT, 202871_AT, 201975_AT, 201285_AT, 216780_AT, 205178_S_AT, 222104_X_AT, 212369_AT, 210579_S_AT, 31835_AT, 211202_S_AT, 213832_AT, 214253_S_AT, 203739_AT, 206119_AT, 219504_S_AT, 218543_S_AT, 218478_S_AT, 211143_X_AT, 207394_AT, 211917_S_AT, 202051_S_AT, 215978_X_AT, 217593_AT, 215636_AT, 201694_S_AT, 204453_AT, 207202_S_AT, 213269_AT, | 1060 | 2348 | 20581 | 1.422299 | 0.001863 |
| GOTERM_MF_DIRECT | GO:0046872~metal ion binding | 161 | 14.45242 | 3.23E-05 | 206836_AT, 209915_S_AT, 204161_S_AT, 202197_AT, 221092_AT, 206078_AT, 205181_AT, 221287_AT, 201904_S_AT, 208709_S_AT, 219951_S_AT, 212876_AT, 214981_AT, 206792_X_AT, 206683_AT, 219086_AT, 219786_AT, 206166_S_AT, 201906_S_AT, 209211_AT, 208357_X_AT, 210336_X_AT, 220858_AT, 202702_AT, 210800_AT, 208222_AT, 207058_S_AT, 221645_S_AT, 220113_X_AT, 220055_AT, 219676_AT, 207625_S_AT, 218742_AT, 204523_AT, 216910_AT, 220668_S_AT, 212816_S_AT, 218524_AT, 216638_S_AT, 205938_AT, 206301_AT, 206502_S_AT, 217428_S_AT, 41113_AT, 204685_S_AT, 213639_S_AT, 207120_AT, 206455_S_AT, 201368_AT, 203948_S_AT, 203704_S_AT, 205338_S_AT, 217242_AT, 212485_AT, 220891_AT, 214751_AT, 201975_AT, 212753_AT, 207998_S_AT, 201285_AT, 206759_AT, 216780_AT, 212369_AT, 210304_AT, 222104_X_AT, | 967 | 2069 | 16881 | 1.358429 | 0.051835 |

| | | | | | | | | | | |
|---|---|---|---|---|---|---|---|---|---|---|
| GOTERM_MF_DIRECT | GO:0008270~zinc ion binding | 84 | 7.540395 | 0.023069 | 215552_S_AT, 221064_S_AT, 204088_AT, 212080_AT, 208709_S_AT, 49327_AT, 202051_S_AT, 210391_AT, 214082_AT, 201549_X_AT, 215848_AT, 201694_S_AT, 205624_AT, 215636_AT, 204453_AT, 207202_S_AT, 212704_AT, 203985_AT, 213204_AT, 219228_AT, 202702_AT, 207129_AT, 202326_AT, 207058_S_AT, 220618_S_AT, 210736_X_AT, 210694_S_AT, 209705_AT, 207115_X_AT, 202193_AT, 214861_AT, 202032_S_AT, 211002_S_AT, 207186_S_AT, 216026_S_AT, 209054_S_AT, 209959_AT, 214703_S_AT, 218173_S_AT, 210159_S_AT, 212982_AT, 203956_AT, 218803_AT, 202423_AT, 217250_S_AT, 215461_AT, 209205_S_AT, 218836_AT, 203160_S_AT, 222030_AT, 212760_AT, 212359_S_AT, 217594_AT, 206718_AT, 202340_X_AT, 202871_AT, 201975_AT, 207251_AT, 222058_AT, 210341_AT, 210266_S_AT, 212753_AT, 201285_AT, 220605_S_AT, 205178_S_AT, 202486_AT, 208279_S_AT, | 967 | 1169 | 16881 | 1.254401 | 31.2108 |
| Annotation Cluster 2 | Enrichment Score: 4.233492463191242 | | | | | | | | | |
| Category | Term | Count | % | PValue | Genes | List Total | Pop Hits | Pop Total | ld Enrichme | FDR |
| UP_KEYWORDS | Cell junction | 67 | 6.014363 | 4.27E-07 | 208823_S_AT, 209915_S_AT, 210251_S_AT, 210782_X_AT, 217419_X_AT, 215810_X_AT, 211451_S_AT, 219039_AT, 206166_S_AT, 212285_S_AT, 220858_AT, 213939_S_AT, 207217_S_AT, 204524_AT, 202149_AT, 213845_AT, 210736_X_AT, 221857_S_AT, 221526_X_AT, 209925_AT, 220492_S_AT, 203016_S_AT, 205417_S_AT, 209638_X_AT, 204730_AT, 206189_AT, 212982_AT, 209873_S_AT, 209586_S_AT, 204447_AT, 206490_AT, 204685_S_AT, 212518_AT, 204765_AT, 204990_S_AT, 221347_AT, 221244_S_AT, 202178_AT, 201839_S_AT, 40225_AT, 202761_S_AT, 217820_S_AT, 202871_AT, 206705_AT, 200931_S_AT, 206489_S_AT, 210094_S_AT, 201015_S_AT, 221088_S_AT, 200606_AT, 207011_S_AT, 205245_AT, 214081_AT, 216024_AT, 206735_AT, 205230_AT, 32029_AT, 207436_X_AT, 218874_S_AT, 209793_AT, 46665_AT, | 1060 | 675 | 20581 | 1.927222 | 6.09E-04 |
| GOTERM_CC_DIRECT | GO:0030054~cell junction | 45 | 4.039497 | 2.76E-04 | 208823_S_AT, 210251_S_AT, 209915_S_AT, 210782_X_AT, 204088_AT, 217419_X_AT, 211451_S_AT, 219039_AT, 206166_S_AT, 212285_S_AT, 213939_S_AT, 207217_S_AT, 213845_AT, 210736_X_AT, 221526_X_AT, 209925_AT, 220492_S_AT, 209638_X_AT, 204730_AT, 212982_AT, 206189_AT, 204447_AT, 206490_AT, 204685_S_AT, 204990_S_AT, 221347_AT, 204765_AT, 202178_AT, 203623_AT, 217820_S_AT, 206705_AT, 206489_S_AT, 210094_S_AT, 221088_S_AT, 200606_AT, 205245_AT, 214530_X_AT, 220804_S_AT, 206735_AT, 205230_AT, 209793_AT, 46665_AT, 210181_S_AT, 217096_AT, 217637_AT, 217280_X_AT, 206844_AT, 210835_S_AT, 216651_S_AT, 215865_AT | 1009 | 459 | 18224 | 1.77073 | 0.404525 |
| UP_KEYWORDS | Synapse | 33 | 2.962298 | 0.001687 | 208823_S_AT, 209915_S_AT, 210782_X_AT, 217419_X_AT, 217820_S_AT, 206705_AT, 211451_S_AT, 219039_AT, 206489_S_AT, 221088_S_AT, 212285_S_AT, 216024_AT, 206735_AT, 213845_AT, 210736_X_AT, 205230_AT, 46665_AT, 209793_AT, 210181_S_AT, 217096_AT, 220492_S_AT, 205417_S_AT, 217637_AT, 219760_AT, 209638_X_AT, 217280_X_AT, 204730_AT, 212982_AT, 206189_AT, 204447_AT, 210835_S_AT, 216651_S_AT, 206490_AT, 204685_S_AT, 215865_AT, 221347_AT | 1060 | 357 | 20581 | 1.79476 | 2.377822 |
| Annotation Cluster 3 | Enrichment Score: 3.612286442678076 | | | | | | | | | |
| Category | Term | Count | % | PValue | Genes | List Total | Pop Hits | Pop Total | ld Enrichme | FDR |

| | | | | | | | | | | |
|---|---|---|---|---|---|---|---|---|---|---|
| UP_KEYWORDS | Transcription | 175 | 15.70916 | 1.22E-06 | 215552_S_AT, 221092_AT, 210391_AT, 201997_S_AT, 201549_X_AT, 206683_AT, 209211_AT, 210336_X_AT, 204947_AT, 210977_S_AT, 207267_S_AT, 32259_AT, 207058_S_AT, 209290_S_AT, 220055_AT, 207186_S_AT, 207625_S_AT, 214948_S_AT, 207700_S_AT, 218173_S_AT, 215881_X_AT, 202616_S_AT, 213743_AT, 206502_S_AT, 41113_AT, 213920_AT, 203704_S_AT, 217242_AT, 217877_S_AT, 202340_X_AT, 207672_AT, 203693_S_AT, 216780_AT, 208574_AT, 50221_AT, 222104_X_AT, 212369_AT, 207084_AT, 219735_S_AT, 211202_S_AT, 32029_AT, 203739_AT, 204980_AT, 219504_S_AT, 221230_S_AT, 218441_S_AT, 203752_S_AT, 210835_S_AT, 211143_X_AT, 207394_AT, 202360_AT, 215404_X_AT, 215978_X_AT, 216109_AT, 217593_AT, 201694_S_AT, 204453_AT, 207202_S_AT, 213269_AT, 203985_AT, 202611_S_AT, 219228_AT, 220844_AT, 204524_AT, 202485_S_AT, | 1060 | 2398 | 20581 | 1.416934 | 0.001737 |
| UP_KEYWORDS | Transcription regulation | 169 | 15.17056 | 3.13E-06 | 215552_S_AT, 221092_AT, 210391_AT, 201997_S_AT, 201549_X_AT, 206683_AT, 209211_AT, 210336_X_AT, 204947_AT, 210977_S_AT, 207267_S_AT, 207058_S_AT, 32259_AT, 209290_S_AT, 220055_AT, 207186_S_AT, 207625_S_AT, 214948_S_AT, 207700_S_AT, 218173_S_AT, 215881_X_AT, 202616_S_AT, 213743_AT, 206502_S_AT, 41113_AT, 213920_AT, 203704_S_AT, 217242_AT, 202340_X_AT, 217877_S_AT, 207672_AT, 203693_S_AT, 216780_AT, 208574_AT, 50221_AT, 222104_X_AT, 212369_AT, 207084_AT, 219735_S_AT, 211202_S_AT, 32029_AT, 203739_AT, 204980_AT, 219504_S_AT, 221230_S_AT, 203752_S_AT, 210835_S_AT, 211143_X_AT, 207394_AT, 202360_AT, 215404_X_AT, 215978_X_AT, 216109_AT, 217593_AT, 201694_S_AT, 204453_AT, 207202_S_AT, 213269_AT, 202611_S_AT, 203985_AT, 219228_AT, 220844_AT, 204524_AT, 202485_S_AT, | 1060 | 2332 | 20581 | 1.40708 | 0.004457 |
| UP_KEYWORDS | Nucleus | 324 | 29.08438 | 9.08E-05 | 215552_S_AT, 221092_AT, 212455_AT, 221217_S_AT, 222152_AT, 220079_S_AT, 205271_S_AT, 201085_S_AT, 212027_AT, 210391_AT, 201997_S_AT, 201549_X_AT, 202256_AT, 206683_AT, 204558_AT, 213694_AT, 213235_AT, 207855_S_AT, 209211_AT, 210336_X_AT, 204947_AT, 210977_S_AT, 220370_S_AT, 214047_S_AT, 201817_AT, 205933_AT, 207267_S_AT, 32259_AT, 202326_AT, 207058_S_AT, 209290_S_AT, 220055_AT, 209256_S_AT, 219977_AT, 207625_S_AT, 207186_S_AT, 214048_AT, 200686_S_AT, 216026_S_AT, 214948_S_AT, 201696_AT, 205890_S_AT, 208382_S_AT, 212816_S_AT, 202127_AT, 210737_AT, 207700_S_AT, 218173_S_AT, 209873_S_AT, 202616_S_AT, 202809_S_AT, 213743_AT, 206502_S_AT, 41113_AT, 206815_AT, 204129_AT, 203160_S_AT, 213920_AT, 201536_AT, 203704_S_AT, 40640_AT, 217594_AT, 217242_AT, 206718_AT, 202761_S_AT, 202871_AT, | 1060 | 5244 | 20581 | 1.199618 | 0.129304 |
| UP_KEYWORDS | DNA-binding | 141 | 12.65709 | 2.70E-04 | 214646_AT, 208006_AT, 215552_S_AT, 221092_AT, 205181_AT, 212080_AT, 201085_S_AT, 210391_AT, 201997_S_AT, 204739_AT, 206683_AT, 204558_AT, 203882_AT, 209211_AT, 210336_X_AT, 204947_AT, 210977_S_AT, 214047_S_AT, 205933_AT, 213660_S_AT, 213359_AT, 221645_S_AT, 209290_S_AT, 220055_AT, 219676_AT, 214048_AT, 204523_AT, 216026_S_AT, 214948_S_AT, 207684_AT, 208382_S_AT, 220668_S_AT, 209959_AT, 214060_AT, 218524_AT, 202616_S_AT, 206502_S_AT, 41113_AT, 213639_S_AT, 207120_AT, 214404_X_AT, 213920_AT, 201368_AT, 203704_S_AT, 208239_AT, 217242_AT, 217877_S_AT, 202340_X_AT, 203693_S_AT, 207672_AT, 214751_AT, 211105_S_AT, 210266_S_AT, 216780_AT, 208574_AT, 78495_AT, 50221_AT, 212369_AT, 207084_AT, 220804_S_AT, 219735_S_AT, 212431_AT, 216350_S_AT, 203739_AT, 213081_AT, 202519_AT, 204980_AT, | 1060 | 2050 | 20581 | 1.335445 | 0.383453 |

| Category | Term | Count | % | PValue | Genes | List Total | Pop Hits | Pop Total | Fold Enrichment | FDR |
|---|---|---|---|---|---|---|---|---|---|---|
| GOTERM_BP_DIRECT | GO:0006351~transcription, DNA-templated | 144 | 12.92639 | 4.93E-04 | 210499_S_AT, 215552_S_AT, 207144_S_AT, 205181_AT, 210539_AT, 210391_AT, 201997_S_AT, 201549_X_AT, 206683_AT, 200704_AT, 210336_X_AT, 204947_AT, 210977_S_AT, 207267_S_AT, 32259_AT, 207058_S_AT, 213359_AT, 221645_S_AT, 220113_X_AT, 209290_S_AT, 220055_AT, 214861_AT, 219676_AT, 207625_S_AT, 207186_S_AT, 218259_AT, 204523_AT, 207684_AT, 209959_AT, 218173_S_AT, 218524_AT, 207700_S_AT, 209638_X_AT, 213927_AT, 215881_X_AT, 203221_AT, 202775_S_AT, 202616_S_AT, 206502_S_AT, 213743_AT, 41113_AT, 213639_S_AT, 207120_AT, 214404_X_AT, 213920_AT, 208099_X_AT, 208239_AT, 217242_AT, 217877_S_AT, 202340_X_AT, 203693_S_AT, 214751_AT, 212753_AT, 210266_S_AT, 216836_S_AT, 216780_AT, 208574_AT, 78495_AT, 50221_AT, 40837_AT, 212369_AT, 220804_S_AT, 219735_S_AT, | 945 | 1955 | 16792 | 1.308839 | 0.894007 |
| GOTERM_MF_DIRECT | GO:0003700~transcription factor activity, sequence-specific DNA binding | 78 | 7.001795 | 0.001671 | 208006_AT, 215552_S_AT, 207144_S_AT, 221092_AT, 205181_AT, 203577_AT, 205514_AT, 212080_AT, 207090_X_AT, 202474_S_AT, 210391_AT, 201549_X_AT, 218312_S_AT, 206683_AT, 203882_AT, 217593_AT, 201694_S_AT, 207202_S_AT, 209211_AT, 210336_X_AT, 204947_AT, 210977_S_AT, 209454_S_AT, 208216_AT, 207058_S_AT, 218639_S_AT, 221645_S_AT, 209290_S_AT, 220055_AT, 219676_AT, 207625_S_AT, 211002_S_AT, 204523_AT, 207684_AT, 208137_X_AT, 202152_X_AT, 207757_AT, 201369_S_AT, 218524_AT, 207767_S_AT, 202616_S_AT, 213980_S_AT, 206502_S_AT, 210669_AT, 41113_AT, 205039_S_AT, 209205_S_AT, 213639_S_AT, 215620_AT, 212847_AT, 201368_AT, 216960_S_AT, 209863_S_AT, 212863_X_AT, 203704_S_AT, 208239_AT, 217242_AT, 217877_S_AT, 203693_S_AT, 211105_S_AT, | 967 | 961 | 16881 | 1.416912 | 2.645573 |
| GOTERM_CC_DIRECT | GO:0005634~nucleus | 339 | 30.43088 | 0.003816 | 215552_S_AT, 221092_AT, 212455_AT, 221217_S_AT, 222152_AT, 205271_S_AT, 218079_S_AT, 210391_AT, 210944_S_AT, 201997_S_AT, 201549_X_AT, 202256_AT, 206683_AT, 204558_AT, 206166_S_AT, 213694_AT, 207855_S_AT, 210336_X_AT, 204947_AT, 210977_S_AT, 220370_S_AT, 214047_S_AT, 201817_AT, 209275_S_AT, 205933_AT, 207267_S_AT, 202326_AT, 207058_S_AT, 209290_S_AT, 220055_AT, 209256_S_AT, 219977_AT, 207625_S_AT, 218319_AT, 207186_S_AT, 214048_AT, 200686_S_AT, 216026_S_AT, 208845_AT, 214948_S_AT, 201696_AT, 205890_S_AT, 208382_S_AT, 212816_S_AT, 210737_AT, 207700_S_AT, 218173_S_AT, 215881_X_AT, 209873_S_AT, 202616_S_AT, 202809_S_AT, 213743_AT, 206502_S_AT, 41113_AT, 217939_S_AT, 206815_AT, 204129_AT, 204990_S_AT, 203160_S_AT, | 1009 | 5415 | 18224 | 1.130717 | 5.451939 |
| GOTERM_MF_DIRECT | GO:0003677~DNA binding | 113 | 10.14363 | 0.043011 | 210499_S_AT, 215552_S_AT, 212080_AT, 201085_S_AT, 210391_AT, 201997_S_AT, 201549_X_AT, 204739_AT, 204558_AT, 204947_AT, 202702_AT, 222046_AT, 205933_AT, 213660_S_AT, 221645_S_AT, 220113_X_AT, 209290_S_AT, 220055_AT, 204523_AT, 216026_S_AT, 214948_S_AT, 207684_AT, 213313_AT, 208382_S_AT, 209959_AT, 218524_AT, 202616_S_AT, 207120_AT, 216113_AT, 201368_AT, 217877_S_AT, 202340_X_AT, 202871_AT, 203693_S_AT, 207672_AT, 214751_AT, 211105_S_AT, 222047_S_AT, 212231_AT, 210266_S_AT, 216780_AT, 78495_AT, 212369_AT, 212431_AT, 218621_AT, 211202_S_AT, 216350_S_AT, 213081_AT, 202519_AT, 204980_AT, 221230_S_AT, 215822_X_AT, 217563_AT, 212293_AT, 208599_AT, 218441_S_AT, 214514_AT, 211143_X_AT, 207394_AT, 212375_AT, 214715_X_AT, 208224_AT, 212152_X_AT, 212376_S_AT, 205514_AT, 213266_AT, 208835_S_AT, | 967 | 1674 | 16881 | 1.178405 | 50.57531 |
| Annotation Cluster 4 | Enrichment Score: 3.272681408073182 | | | | | | | | | |
| Category | Term | Count | % | PValue | Genes | List Total | Pop Hits | Pop Total | Fold Enrichment | FDR |

| | | | | | | | | | | |
|---|---|---|---|---|---|---|---|---|---|---|
| UP_KEYWORDS | Transferase | 128 | 11.49013 | 1.38E-05 | 215299_X_AT, 219639_X_AT, 210981_S_AT, 206078_AT, 202894_AT, 203615_X_AT, 212080_AT, 205271_S_AT, 221164_X_AT, 211034_S_AT, 212876_AT, 207133_X_AT, 203398_S_AT, 203671_AT, 208222_AT, 203127_S_AT, 201817_AT, 213700_S_AT, 32259_AT, 207058_S_AT, 202326_AT, 218017_S_AT, 219365_S_AT, 214165_S_AT, 220113_X_AT, 202193_AT, 213670_X_AT, 209467_S_AT, 216026_S_AT, 202127_AT, 220668_S_AT, 218173_S_AT, 207700_S_AT, 221696_S_AT, 213927_AT, 206301_AT, 218652_S_AT, 214114_X_AT, 203229_S_AT, 214322_AT, 203397_S_AT, 214100_X_AT, 216185_AT, 202849_X_AT, 202676_X_AT, 219815_AT, 40225_AT, 217594_AT, 212403_AT, 220891_AT, 207312_AT, 203669_S_AT, 38447_AT, 204060_S_AT, 215654_AT, 212380_AT, 205370_X_AT, 216836_S_AT, 211424_X_AT, 216451_AT, 220737_AT, | 1060 | 1708 | 20581 | 1.455066 | 0.019678 |
| INTERPRO | IPR000719:Protein kinase, catalytic domain | 49 | 4.398564 | 4.95E-05 | 208823_S_AT, 220958_AT, 210981_S_AT, 202894_AT, 206078_AT, 213792_S_AT, 215404_X_AT, 205271_S_AT, 221287_AT, 206223_AT, 205486_AT, 212252_AT, 208222_AT, 204524_AT, 207073_AT, 216765_AT, 203709_AT, 219365_S_AT, 202193_AT, 209467_S_AT, 220030_AT, 202127_AT, 221696_S_AT, 214339_S_AT, 206301_AT, 213927_AT, 203229_S_AT, 214322_AT, 221244_S_AT, 202849_X_AT, 202178_AT, 203139_AT, 40225_AT, 216310_AT, 207312_AT, 204060_S_AT, 38447_AT, 216836_S_AT, 204890_S_AT, 207011_S_AT, 216451_AT, 220737_AT, 215296_AT, 215326_AT, 212757_S_AT, 32029_AT, 207187_AT, 202454_S_AT, 203379_AT, 208820_AT, 217849_S_AT, 215234_AT, 212293_AT, 204267_X_AT, 202951_AT, 210582_S_AT, 202281_AT, 215638_AT | 1010 | 487 | 18559 | 1.848844 | 0.083156 |
| INTERPRO | IPR017441:Protein kinase, ATP binding site | 41 | 3.680431 | 5.31E-05 | 221244_S_AT, 208823_S_AT, 202849_X_AT, 210981_S_AT, 202178_AT, 206078_AT, 202894_AT, 213792_S_AT, 203139_AT, 215404_X_AT, 205271_S_AT, 206223_AT, 216310_AT, 207312_AT, 38447_AT, 204060_S_AT, 204890_S_AT, 216836_S_AT, 205486_AT, 216451_AT, 212252_AT, 208222_AT, 220737_AT, 204524_AT, 207073_AT, 216765_AT, 203709_AT, 215326_AT, 215296_AT, 212757_S_AT, 32029_AT, 207187_AT, 202193_AT, 209467_S_AT, 208820_AT, 203379_AT, 217849_S_AT, 214339_S_AT, 213927_AT, 206301_AT, 215234_AT, 212293_AT, 204267_X_AT, 202951_AT, 210582_S_AT, 203229_S_AT, 214322_AT | 1010 | 381 | 18559 | 1.977389 | 0.08916 |
| INTERPRO | IPR011009:Protein kinase-like domain | 52 | 4.667864 | 5.75E-05 | 208823_S_AT, 220958_AT, 210981_S_AT, 202894_AT, 206078_AT, 213792_S_AT, 215404_X_AT, 205271_S_AT, 221287_AT, 206223_AT, 207133_X_AT, 205486_AT, 212252_AT, 208222_AT, 204524_AT, 207073_AT, 216765_AT, 203709_AT, 219365_S_AT, 202193_AT, 220030_AT, 209467_S_AT, 202127_AT, 221696_S_AT, 214339_S_AT, 213927_AT, 206301_AT, 203229_S_AT, 214322_AT, 221244_S_AT, 202849_X_AT, 202178_AT, 203139_AT, 40225_AT, 216310_AT, 207312_AT, 204060_S_AT, 38447_AT, 216836_S_AT, 204890_S_AT, 207011_S_AT, 216451_AT, 220737_AT, 215326_AT, 215296_AT, 212757_S_AT, 215588_X_AT, 32029_AT, 207187_AT, 202454_S_AT, 203379_AT, 208820_AT, 217849_S_AT, 215234_AT, 212293_AT, 202642_S_AT, 204267_X_AT, 202951_AT, 210582_S_AT, 202281_AT, 215638_AT | 1010 | 531 | 18559 | 1.799459 | 0.09652 |

| | | | | | | | | | | |
|---|---|---|---|---|---|---|---|---|---|---|
| GOTERM_BP_DIRECT | GO:0006468~protein phosphorylation | 47 | 4.219031 | 8.79E-05 | 202360_AT, 208823_S_AT, 220958_AT, 210981_S_AT, 206078_AT, 215404_X_AT, 205271_S_AT, 221287_AT, 206223_AT, 207133_X_AT, 210975_X_AT, 205486_AT, 212252_AT, 208222_AT, 204524_AT, 207073_AT, 219365_S_AT, 203709_AT, 202193_AT, 209467_S_AT, 202127_AT, 214339_S_AT, 206301_AT, 213927_AT, 213743_AT, 213980_S_AT, 214114_X_AT, 203229_S_AT, 221244_S_AT, 206455_S_AT, 202849_X_AT, 202676_X_AT, 212863_X_AT, 202178_AT, 203139_AT, 40225_AT, 216310_AT, 207312_AT, 210133_AT, 204890_S_AT, 216836_S_AT, 207011_S_AT, 216451_AT, 220737_AT, 215296_AT, 32029_AT, 215588_X_AT, 207187_AT, 203379_AT, 217849_S_AT, 215234_AT, 212293_AT, 204267_X_AT, 202951_AT, 210582_S_AT, 202281_AT | 945 | 456 | 16792 | 1.831486 | 0.159863 |
| UP_SEQ_FEATURE | domain:Protein kinase | 46 | 4.129264 | 8.88E-05 | 208823_S_AT, 220958_AT, 210981_S_AT, 206078_AT, 202894_AT, 213792_S_AT, 215404_X_AT, 205271_S_AT, 221287_AT, 206223_AT, 205486_AT, 212252_AT, 208222_AT, 204524_AT, 207073_AT, 216765_AT, 219365_S_AT, 203709_AT, 202193_AT, 220030_AT, 209467_S_AT, 202127_AT, 221696_S_AT, 214339_S_AT, 206301_AT, 213927_AT, 203229_S_AT, 214322_AT, 221244_S_AT, 202849_X_AT, 202178_AT, 203139_AT, 40225_AT, 216310_AT, 207312_AT, 38447_AT, 204060_S_AT, 204890_S_AT, 216836_S_AT, 216451_AT, 215296_AT, 215326_AT, 32029_AT, 212757_S_AT, 215588_X_AT, 202454_S_AT, 208820_AT, 217849_S_AT, 215234_AT, 212293_AT, 204267_X_AT, 202951_AT, 210582_S_AT, 202281_AT, 215638_AT | 1043 | 478 | 20063 | 1.85115 | 0.159779 |
| UP_KEYWORDS | Serine/threonine-protein kinase | 39 | 3.500898 | 1.48E-04 | 221244_S_AT, 208823_S_AT, 202849_X_AT, 220958_AT, 210981_S_AT, 202676_X_AT, 202178_AT, 206078_AT, 203139_AT, 205271_S_AT, 40225_AT, 206223_AT, 216310_AT, 207312_AT, 207133_X_AT, 38447_AT, 204060_S_AT, 210975_X_AT, 205486_AT, 216451_AT, 212252_AT, 208222_AT, 220737_AT, 204524_AT, 207073_AT, 216765_AT, 203709_AT, 215326_AT, 215296_AT, 215588_X_AT, 212757_S_AT, 32029_AT, 202193_AT, 209467_S_AT, 202127_AT, 203379_AT, 217849_S_AT, 214339_S_AT, 213927_AT, 215234_AT, 212293_AT, 214114_X_AT, 204267_X_AT, 202951_AT, 210582_S_AT, 203229_S_AT, 202281_AT, 214322_AT | 1060 | 393 | 20581 | 1.926782 | 0.211173 |
| UP_KEYWORDS | ATP-binding | 103 | 9.245961 | 1.68E-04 | 210981_S_AT, 206078_AT, 202894_AT, 221287_AT, 205271_S_AT, 211207_S_AT, 210539_AT, 204558_AT, 216409_AT, 213204_AT, 208222_AT, 213700_S_AT, 203727_AT, 220920_AT, 215873_X_AT, 202193_AT, 209467_S_AT, 208382_S_AT, 202127_AT, 221696_S_AT, 213927_AT, 206301_AT, 214114_X_AT, 203229_S_AT, 204685_S_AT, 214322_AT, 202849_X_AT, 202676_X_AT, 208099_X_AT, 40225_AT, 207312_AT, 38447_AT, 204060_S_AT, 212441_AT, 216836_S_AT, 216451_AT, 220737_AT, 203082_AT, 215911_X_AT, 204497_AT, 209474_S_AT, 215326_AT, 32029_AT, 215588_X_AT, 221225_AT, 206317_S_AT, 202454_S_AT, 214594_X_AT, 208820_AT, 212293_AT, 210197_AT, 202951_AT, 202804_AT, 202307_S_AT, 213485_S_AT, 212375_AT, 208823_S_AT, 220958_AT, 214328_S_AT, 211970_X_AT, 213792_S_AT, 212376_S_AT, 207561_S_AT, 215404_X_AT, | 1060 | 1391 | 20581 | 1.437708 | 0.238748 |

| | | | | | | | | | | |
|---|---|---|---|---|---|---|---|---|---|---|
| GOTERM_MF_DIRECT | GO:0004672~protein kinase activity | 38 | 3.411131 | 3.90E-04 | 221244_S_AT, 208823_S_AT, 220958_AT, 202676_X_AT, 202178_AT, 206078_AT, 203577_AT, 203139_AT, 205271_S_AT, 221287_AT, 40225_AT, 216310_AT, 207312_AT, 38447_AT, 210975_X_AT, 216836_S_AT, 207011_S_AT, 205486_AT, 222104_X_AT, 220737_AT, 204524_AT, 207073_AT, 216765_AT, 203709_AT, 219365_S_AT, 215326_AT, 32029_AT, 202193_AT, 209467_S_AT, 220030_AT, 202454_S_AT, 208820_AT, 202127_AT, 203379_AT, 221696_S_AT, 217849_S_AT, 214339_S_AT, 213927_AT, 215234_AT, 212293_AT, 214114_X_AT, 204267_X_AT, 210582_S_AT, 203229_S_AT, 202281_AT, 215638_AT | 967 | 359 | 16881 | 1.847825 | 0.623118 |
| UP_SEQ_FEATURE | binding site:ATP | 49 | 4.398564 | 4.39E-04 | 208823_S_AT, 210981_S_AT, 202894_AT, 206078_AT, 213792_S_AT, 215404_X_AT, 205271_S_AT, 206223_AT, 205486_AT, 212252_AT, 208222_AT, 204524_AT, 202726_AT, 207073_AT, 216765_AT, 203709_AT, 202193_AT, 209467_S_AT, 220030_AT, 202127_AT, 221696_S_AT, 214339_S_AT, 206301_AT, 213927_AT, 203229_S_AT, 214322_AT, 221244_S_AT, 202849_X_AT, 202178_AT, 218231_AT, 203139_AT, 218844_AT, 216310_AT, 207312_AT, 204060_S_AT, 38447_AT, 204890_S_AT, 216836_S_AT, 216451_AT, 220737_AT, 215296_AT, 215326_AT, 215588_X_AT, 212757_S_AT, 32029_AT, 207187_AT, 202454_S_AT, 203379_AT, 208820_AT, 217849_S_AT, 215234_AT, 212293_AT, 210197_AT, 204267_X_AT, 202951_AT, 210582_S_AT, 215638_AT | 1043 | 558 | 20063 | 1.68917 | 0.788105 |
| UP_KEYWORDS | Kinase | 60 | 5.385996 | 4.87E-04 | 208823_S_AT, 220958_AT, 210981_S_AT, 202894_AT, 206078_AT, 213792_S_AT, 214928_AT, 215404_X_AT, 205271_S_AT, 206223_AT, 207133_X_AT, 210975_X_AT, 205486_AT, 208222_AT, 212252_AT, 219632_S_AT, 204524_AT, 207073_AT, 213700_S_AT, 216765_AT, 203709_AT, 219365_S_AT, 202193_AT, 220030_AT, 209467_S_AT, 202127_AT, 221696_S_AT, 218942_AT, 214339_S_AT, 213927_AT, 206301_AT, 214114_X_AT, 203229_S_AT, 212518_AT, 214322_AT, 221244_S_AT, 202849_X_AT, 202676_X_AT, 202178_AT, 203139_AT, 218231_AT, 40225_AT, 202743_AT, 216310_AT, 207312_AT, 204060_S_AT, 38447_AT, 213396_S_AT, 216836_S_AT, 204890_S_AT, 216451_AT, 220737_AT, 215326_AT, 215296_AT, 212757_S_AT, 215588_X_AT, 32029_AT, 207187_AT, 202454_S_AT, 203379_AT, 208820_AT, 217849_S_AT, 215234_AT, 212293_AT, 210197_AT, 204267_X_AT, 213177_AT, 202951_AT, 210582_S_AT, 202281_AT, | 1060 | 735 | 20581 | 1.584983 | 0.691603 |
| UP_SEQ_FEATURE | active site:Proton acceptor | 56 | 5.02693 | 5.94E-04 | 215299_X_AT, 208823_S_AT, 210981_S_AT, 202894_AT, 206078_AT, 213792_S_AT, 203615_X_AT, 215404_X_AT, 205271_S_AT, 208709_S_AT, 213279_AT, 49327_AT, 206223_AT, 207064_S_AT, 203608_AT, 205486_AT, 208222_AT, 212252_AT, 204524_AT, 207073_AT, 216765_AT, 203709_AT, 209467_S_AT, 220030_AT, 202127_AT, 221696_S_AT, 214339_S_AT, 213927_AT, 206301_AT, 220224_AT, 203229_S_AT, 214322_AT, 221244_S_AT, 202849_X_AT, 203948_S_AT, 202178_AT, 203139_AT, 40225_AT, 216310_AT, 207312_AT, 204060_S_AT, 38447_AT, 212380_AT, 216836_S_AT, 204890_S_AT, 220605_S_AT, 216451_AT, 220737_AT, 204573_AT, 215326_AT, 215296_AT, 215588_X_AT, 212757_S_AT, 32029_AT, 207187_AT, 202454_S_AT, 203379_AT, 208820_AT, 217849_S_AT, 215234_AT, 212293_AT, 204267_X_AT, 202951_AT, 202281_AT, 215638_AT | 1043 | 672 | 20063 | 1.602988 | 1.064517 |

| | | | | | | | | | | |
|---|---|---|---|---|---|---|---|---|---|---|
| GOTERM_MF_DIRECT | GO:0004674~protein serine/threonine kinase activity | 38 | 3.411131 | 9.29E-04 | 221244_S_AT, 208823_S_AT, 220958_AT, 202676_X_AT, 206078_AT, 202178_AT, 203139_AT, 205271_S_AT, 40225_AT, 206223_AT, 216310_AT, 207312_AT, 207133_X_AT, 204060_S_AT, 210975_X_AT, 205486_AT, 212252_AT, 208222_AT, 216451_AT, 220737_AT, 204524_AT, 207073_AT, 216765_AT, 203709_AT, 215326_AT, 215296_AT, 215588_X_AT, 212757_S_AT, 32029_AT, 202193_AT, 209467_S_AT, 202127_AT, 203379_AT, 217849_S_AT, 214339_S_AT, 213927_AT, 215234_AT, 212293_AT, 213743_AT, 214114_X_AT, 204267_X_AT, 202951_AT, 210582_S_AT, 203229_S_AT, 202281_AT, 214322_AT | 967 | 376 | 16881 | 1.76428 | 1.479397 |
| UP_SEQ_FEATURE | nucleotide phosphate-binding region:ATP | 73 | 6.552962 | 0.002667 | 212375_AT, 208823_S_AT, 210981_S_AT, 202894_AT, 206078_AT, 213792_S_AT, 212376_S_AT, 207561_S_AT, 215404_X_AT, 205271_S_AT, 206223_AT, 204558_AT, 38069_AT, 205486_AT, 213204_AT, 208222_AT, 212252_AT, 208128_X_AT, 204524_AT, 207073_AT, 216765_AT, 203727_AT, 203709_AT, 202193_AT, 220030_AT, 209467_S_AT, 208382_S_AT, 202127_AT, 221696_S_AT, 214339_S_AT, 213927_AT, 206301_AT, 203694_S_AT, 215840_AT, 217250_S_AT, 203229_S_AT, 214322_AT, 221244_S_AT, 202849_X_AT, 204873_AT, 202178_AT, 203139_AT, 218231_AT, 213077_AT, 220777_AT, 216310_AT, 218844_AT, 207312_AT, 204060_S_AT, 38447_AT, 212441_AT, 216836_S_AT, 204890_S_AT, 205887_X_AT, 216451_AT, 208021_S_AT, 203082_AT, 220737_AT, 202486_AT, 213647_AT, 215326_AT, 215296_AT, 32029_AT, 212757_S_AT, 215588_X_AT, 31861_AT, 207187_AT, | 1043 | 994 | 20063 | 1.412694 | 4.698039 |
| INTERPRO | IPR008271:Serine/threonine-protein kinase, active site | 30 | 2.692998 | 0.003505 | 221244_S_AT, 208823_S_AT, 202849_X_AT, 210981_S_AT, 206078_AT, 202178_AT, 203139_AT, 205271_S_AT, 40225_AT, 216310_AT, 207312_AT, 38447_AT, 204060_S_AT, 212252_AT, 208222_AT, 216451_AT, 220737_AT, 204524_AT, 207073_AT, 216765_AT, 203709_AT, 215296_AT, 212757_S_AT, 32029_AT, 209467_S_AT, 202127_AT, 203379_AT, 217849_S_AT, 213927_AT, 215234_AT, 212293_AT, 204267_X_AT, 202951_AT, 203229_S_AT, 202281_AT, 214322_AT | 1010 | 312 | 18559 | 1.766851 | 5.726778 |
| GOTERM_MF_DIRECT | GO:0005524~ATP binding | 108 | 9.694794 | 0.008839 | 210981_S_AT, 206078_AT, 202894_AT, 221287_AT, 205271_S_AT, 211207_S_AT, 210539_AT, 207133_X_AT, 204558_AT, 216409_AT, 213204_AT, 208222_AT, 213700_S_AT, 203727_AT, 219365_S_AT, 220920_AT, 215873_X_AT, 202193_AT, 209467_S_AT, 208382_S_AT, 202127_AT, 221696_S_AT, 213927_AT, 206301_AT, 214114_X_AT, 203229_S_AT, 204685_S_AT, 214322_AT, 202849_X_AT, 202676_X_AT, 208099_X_AT, 40225_AT, 207312_AT, 38447_AT, 204060_S_AT, 212441_AT, 216836_S_AT, 216451_AT, 220737_AT, 203082_AT, 215911_X_AT, 204497_AT, 209474_S_AT, 215326_AT, 32029_AT, 215588_X_AT, 221225_AT, 206317_S_AT, 202454_S_AT, 214594_X_AT, 208820_AT, 212293_AT, 210197_AT, 202951_AT, 213485_S_AT, 202804_AT, 202307_S_AT, 212375_AT, 208823_S_AT, 220958_AT, 214328_S_AT, 211970_X_AT, 213792_S_AT, 212376_S_AT, | 967 | 1495 | 16881 | 1.261114 | 13.26547 |

| | | | | | | | | | | |
|---|---|---|---|---|---|---|---|---|---|---|
| UP_KEYWORDS | Nucleotide-binding | 115 | 10.32316 | 0.00979 | 210981_S_AT, 206078_AT, 202894_AT, 221789_X_AT, 221287_AT, 205271_S_AT, 211207_S_AT, 210539_AT, 204558_AT, 216409_AT, 213204_AT, 208222_AT, 209315_AT, 213700_S_AT, 203727_AT, 220920_AT, 215873_X_AT, 202193_AT, 209467_S_AT, 208845_AT, 208382_S_AT, 202127_AT, 221696_S_AT, 213927_AT, 206301_AT, 214114_X_AT, 203229_S_AT, 204685_S_AT, 214322_AT, 202849_X_AT, 202676_X_AT, 208099_X_AT, 40225_AT, 207312_AT, 38447_AT, 204060_S_AT, 212441_AT, 216836_S_AT, 216451_AT, 220737_AT, 203082_AT, 215911_X_AT, 204497_AT, 209474_S_AT, 215326_AT, 215588_X_AT, 32029_AT, 221225_AT, 206317_S_AT, 202454_S_AT, 214594_X_AT, 208820_AT, 212293_AT, 210197_AT, 213433_AT, 202951_AT, 213485_S_AT, 202804_AT, 202307_S_AT, 212375_AT, 208823_S_AT, 220958_AT, 214328_S_AT, 211970_X_AT, | 1060 | 1788 | 20581 | 1.248794 | 13.08017 |
| GOTERM_BP_DIRECT | GO:0046777~protein autophosphorylation | 18 | 1.615799 | 0.016941 | 221244_S_AT, 32029_AT, 212757_S_AT, 202894_AT, 213792_S_AT, 203139_AT, 215404_X_AT, 209467_S_AT, 206223_AT, 208820_AT, 204060_S_AT, 214339_S_AT, 213927_AT, 215234_AT, 216836_S_AT, 204851_S_AT, 203229_S_AT, 208222_AT, 212252_AT, 214322_AT, 204524_AT | 945 | 172 | 16792 | 1.859579 | 26.71841 |
| SMART | SM00220:S_TKc | 34 | 3.052065 | 0.025783 | 221244_S_AT, 208823_S_AT, 202849_X_AT, 210981_S_AT, 202178_AT, 206078_AT, 203139_AT, 205271_S_AT, 221287_AT, 40225_AT, 216310_AT, 207312_AT, 38447_AT, 204060_S_AT, 212252_AT, 208222_AT, 216451_AT, 220737_AT, 204524_AT, 207073_AT, 216765_AT, 203709_AT, 219365_S_AT, 215326_AT, 215296_AT, 212757_S_AT, 32029_AT, 209467_S_AT, 202127_AT, 203379_AT, 217849_S_AT, 214339_S_AT, 213927_AT, 215234_AT, 212293_AT, 204267_X_AT, 202951_AT, 203229_S_AT, 202281_AT, 214322_AT | 652 | 359 | 10057 | 1.460849 | 30.32528 |
| Annotation Cluster 5 | Enrichment Score: 2.3275745759815414 | | | | | | | | | |
| Category | Term | Count | % | PValue | Genes | List Total | Pop Hits | Pop Total | ld Enrichme | FDR |
| UP_SEQ_FEATURE | zinc finger region:PHD-type 2 | 10 | 0.897666 | 1.41E-05 | 201549_X_AT, 211202_S_AT, 209705_AT, 40446_AT, 214861_AT, 202423_AT, 217250_S_AT, 212080_AT, 207186_S_AT, 209054_S_AT, 218173_S_AT | 1043 | 30 | 20063 | 6.411953 | 0.025428 |
| UP_SEQ_FEATURE | zinc finger region:PHD-type 1 | 10 | 0.897666 | 3.31E-05 | 201549_X_AT, 211202_S_AT, 209705_AT, 40446_AT, 214861_AT, 202423_AT, 217250_S_AT, 212080_AT, 207186_S_AT, 209054_S_AT, 218173_S_AT | 1043 | 33 | 20063 | 5.829048 | 0.059531 |

| | | | | | | | | | | |
|---|---|---|---|---|---|---|---|---|---|---|
| UP_KEYWORDS | Chromatin regulator | 26 | 2.333932 | 0.007283 | 203160_S_AT, 212375_AT, 214911_S_AT, 212152_X_AT, 212376_S_AT, 212080_AT, 202474_S_AT, 213202_AT, 201549_X_AT, 211310_AT, 37652_AT, 211202_S_AT, 202326_AT, 32259_AT, 209705_AT, 207115_X_AT, 40446_AT, 216326_S_AT, 214861_AT, 207186_S_AT, 209054_S_AT, 218173_S_AT, 208685_X_AT, 220667_AT, 212571_AT, 202642_S_AT, 217250_S_AT, 205039_S_AT, 202423_AT, 202455_AT | 1060 | 287 | 20581 | 1.758944 | 9.891386 |
| INTERPRO | IPR011011:Zinc finger, FYVE/PHD-type | 16 | 1.436266 | 0.009821 | 211202_S_AT, 205230_AT, 209705_AT, 202197_AT, 40446_AT, 214861_AT, 212080_AT, 207186_S_AT, 217096_AT, 220668_S_AT, 209054_S_AT, 218173_S_AT, 201549_X_AT, 203651_AT, 210266_S_AT, 217250_S_AT, 202423_AT | 1010 | 141 | 18559 | 2.085134 | 15.27608 |
| UP_SEQ_FEATURE | zinc finger region:PHD-type 3 | 4 | 0.359066 | 0.012734 | 201549_X_AT, 211202_S_AT, 212080_AT, 209054_S_AT, 218173_S_AT | 1043 | 10 | 20063 | 7.694343 | 20.61755 |
| INTERPRO | IPR001965:Zinc finger, PHD-type | 11 | 0.987433 | 0.022475 | 201549_X_AT, 211202_S_AT, 210266_S_AT, 209705_AT, 40446_AT, 214861_AT, 202423_AT, 217250_S_AT, 212080_AT, 207186_S_AT, 209054_S_AT, 218173_S_AT | 1010 | 89 | 18559 | 2.271098 | 31.7361 |
| INTERPRO | IPR019787:Zinc finger, PHD-finger | 10 | 0.897666 | 0.027253 | 201549_X_AT, 211202_S_AT, 210266_S_AT, 209705_AT, 40446_AT, 202423_AT, 217250_S_AT, 212080_AT, 207186_S_AT, 209054_S_AT, 218173_S_AT | 1010 | 79 | 18559 | 2.325981 | 37.12974 |
| SMART | SM00249:PHD | 11 | 0.987433 | 0.06108 | 201549_X_AT, 211202_S_AT, 210266_S_AT, 209705_AT, 40446_AT, 214861_AT, 202423_AT, 217250_S_AT, 212080_AT, 207186_S_AT, 209054_S_AT, 218173_S_AT | 652 | 89 | 10057 | 1.906442 | 58.18146 |
| INTERPRO | IPR019786:Zinc finger, PHD-type, conserved site | 8 | 0.718133 | 0.070878 | 201549_X_AT, 211202_S_AT, 210266_S_AT, 209705_AT, 40446_AT, 217250_S_AT, 207186_S_AT, 209054_S_AT, 218173_S_AT | 1010 | 67 | 18559 | 2.194059 | 70.90952 |
