## Supplementary material for "Targeting dormant ovarian cancer cells *in vitro* and in an *in vivo* model of platinum resistance": Table S6

| Table S6. Average photon flux measurements | | | | | | | | |
|---|---|---|---|---|---|---|---|---|
| **Arm** | **Treatment (Tx)** | **Before Tx** | **Primary Tx: carbo x1** | **Primary Tx: carbo x2** | **Primary Tx: carbo x3** | **Tx break** | **Secondary Tx week 1** | **Secondary Tx week 2** |
| **1** | Carbo-carbo | $4.5x10^7$ | $1.0x10^8$ | $5.1x10^7$ | $1.7x10^7$ | $4.8x10^7$ | $5.4x10^7$ | $4.2x10^7$ |
| **2** | Carbo-UCN-01 | $2.3x10^7$ | $3.9x10^7$ | $1.9x10^7$ | $7.0x10^6$ | $4.6x10^6$ | $7.5x10^6$ | $9.3x10^5$ |
| **3** | Carbo-Oltipraz | $1.6x10^7$ | $2.7x10^7$ | $2.3x10^6$ | $7.0x10^5$ | $5.9x10^5$ | $8.6x10^5$ | $5.5x10^6$ |
| **4** | Carbo-None | $5.2x10^7$ | $5.7x10^7$ | $2.2x10^7$ | $8.8x10^6$ | $4.7x10^7$ | $3.1x10^7$ | $2.1x10^8$ |
| **5** | None | $2.2x10^7$ | $6.3x10^7$ | $6.0x10^7$ | $4.1x10^7$ | $6.8x10^7$ | $2.6x10^8$ | $3.0x10^8$ |
