## Supplementary figures and images for "Targeting dormant ovarian cancer cells *in vitro* and in an *in vivo* model of platinum resistance"

### Figure S1

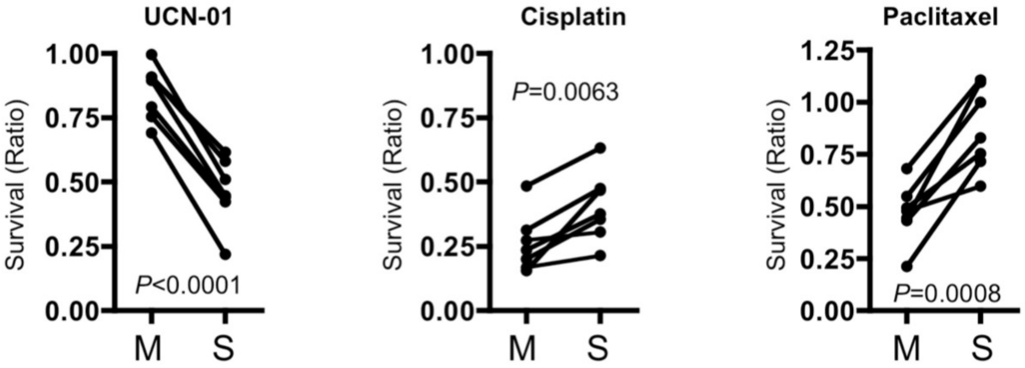
